# A cinnamyl alcohol dehydrogenase-like scaffold organizes monoterpenoid indole alkaloid biosynthesis

**DOI:** 10.64898/2026.04.16.718883

**Authors:** Di Gao, Scott Galeung Alexander Mann, Binbin Chen, Yuanwei Gou, Cong Chen, Chong Qiao, Jorge Jonathan Oswaldo Garza-Garcia, Mohammadamin Shahsavarani, Xiaojing Jiang, Hannah Caroline Tran, Jingfei Bao, Mathew Bailey Richardson, Jianing Li, Jacob Owen Perley, Jaewook Hwang, Feng Dong, Chang Dong, Lei Huang, Vincenzo De Luca, Yajie Wang, Yang Qu, Jiazhang Lian

## Abstract

Biosynthesis of ~3,000 monoterpenoid indole alkaloids (MIAs), including the anticancer drug vinblastine, involves the highly unstable intermediate strictosidine aglycone. Its formation by strictosidine β-glucosidase (SGD) and subsequent conversion by geissoschizine synthase (GS) occur in spatially separated compartments, representing a major biosynthesis bottleneck. Here we discover VinBLAST, a cinnamyl alcohol dehydrogenase-like protein repurposed as a scaffold for efficient processing of this labile intermediate. VinBLAST physically mediates SGD and GS interaction in the nucleus and allosterically enhances GS catalytic efficiency. VinBLAST homologues from diverse plant families enhance biosynthesis of several representative MIAs, with the production of catharanthine increased to ~160 mg L^-1^ in yeast, nearly 1,000-fold higher than previous studies. Our discovery provides a missing link in organizing MIA biosynthesis and enables scalable bioproduction of geissoschizine-derived therapeutics.

---

Catharanthine and vindoline are the immediate precursors of the iconic anticancer monoterpenoid indole alkaloid (MIA) vinblastine in *Catharanthus roseus* (Madagascar periwinkle). The elucidation of their complete ~30-step specialized biosynthetic pathways marks a major milestone in the field of plant secondary metabolism *(1-3)*. This achievement enables their *de novo* biosynthesis in yeast cell factories. However, current production remains at the microgram-per-liter scale, far below the gram-per-liter threshold required for large-scale biomanufacturing *(4-6)*. The low yields likely stem from the lengthy pathway and intricate compartmentalization of MIA biosynthesis *(7, 8)*. Strictosidine, the universal precursor to over 3,000 MIAs, is synthesized in the vacuole and subsequently relocated to the nucleus. There strictosidine β-glucosidase (SGD) removes its glucose moiety to generate strictosidine aglycone, a highly labile intermediate *(9, 10)* (Fig. 1A). In MIA-producing plant families (Apocynaceae, Loganiaceae, Gelsemiaceae, and Rubiaceae within the order of Gentianales), numerous cinnamyl alcohol dehydrogenase (CAD)-like reductases have evolved. These enzymes reduce strictosidine aglycone into stable isomers, enabling further MIA diversification *(11-13)*. For vinblastine, the reduction is performed by the CAD-like geissoschizine synthase (GS), a cytosolic homodimeric enzyme. The resultant geissoschizine is a key branch point enabling the biosynthesis of hundreds of MIAs *(14-16)* (Fig. 1B). Through six and thirteen additional enzymatic steps, geissoschizine is converted into catharanthine and vindoline *(3, 17)*, which are subsequently coupled to form vinblastine *(18)*.

**Fig. 1.**
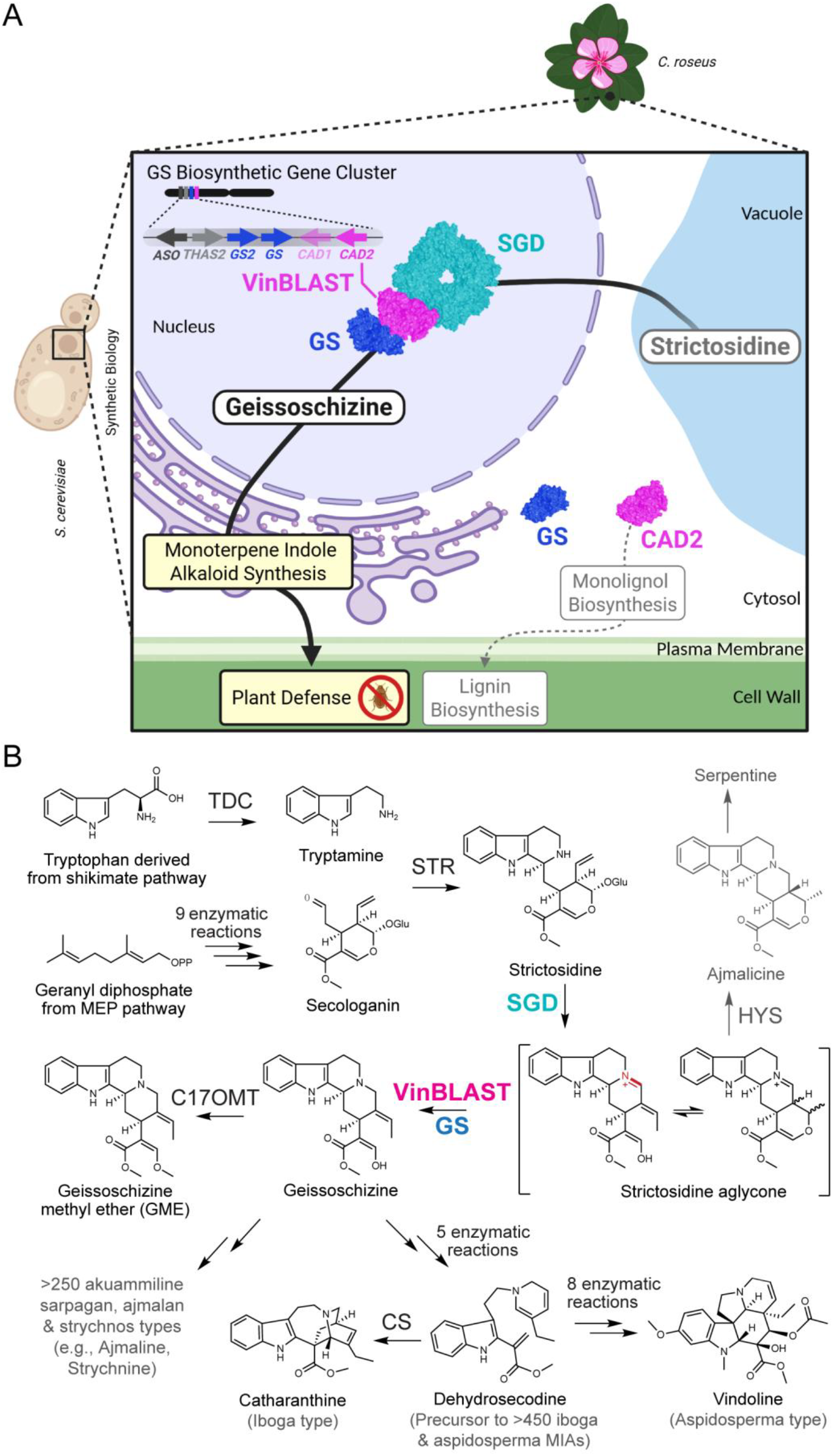
VinBLAST functions as a scaffold tethering SGD and GS in the nucleus, a critical interaction for MIA biosynthesis. **(A)** A proposed model for the scaffolding role of VinBLAST (magenta) in *C. roseus* and bioengineered yeast cells (created with BioRender.com). Evolved from a CAD enzyme, VinBLAST interacts with SGD (cyan) and tethers GS (blue) to SGD in the nucleus. This overcomes the inefficient transport of the unstable intermediate strictosidine aglycone between GS and SGD, enabling its direct conversion to geissoschizine within the nucleus. (**B)** The biosynthetic pathways of GME, catharanthine, vindoline, and other geissoschizine-derived MIAs. TDC: tryptophan decarboxylase; STR: strictosidine synthase; SGD: strictosidine β-glucosidase; GS: geissoschizine synthase; HYS: heteroyohimbine synthase; C17OMT: C17-enol *O*-methyltransferase from *Mitragyna speciosa*; CS: catharanthine synthase.

Despite these milestones, a puzzling question remains regarding in vivo processing of the labile strictosidine aglycone. SGD contains a bipartite nuclear localization signal (NLS) at its *C*-terminus and is thus sequestered in the nucleus *(10)*. Fluorescent protein tagging and bimolecular fluorescence complementation (BiFC) have shown that several CAD-like reductases such as heteroyohimbine synthase (HYS) preferentially localize to the nucleus and physically interacts with SGD. In contrast, although GS displays both cytosolic and nuclear localization, BiFC experiments have not detected an interaction between GS and SGD *(15)*. This observation suggests the involvement of an unidentified factor that facilitates the processing and transfer of strictosidine aglycone from SGD to GS and downstream enzymes.

Here we identify and characterize the **Vin**ca alkaloid **B**iosynthesis **L**ocalizing and **A**ctivating **S**caffold **T**ether (VinBLAST), the missing component that tethers GS and SGD in the nucleus. Molecular dynamics (MD) simulations followed by experimental validations elucidate molecular mechanisms for the SGD-VinBLAST-GS interactions. VinBLAST scaffolds the association between SGD and GS, while simultaneously reshaping the substrate access tunnel at the VinBLAST-GS interface and substantially increasing GS’s catalytic rate. Silencing *VinBLAST* decreases catharanthine and vindoline levels by ~90% in *C. roseus*, demonstrating its essential role in planta. In yeast cell factories, expressing *VinBLAST* increases catharanthine production by nearly three orders of magnitude, promising industrial biomanufacturing of vinblastine and other geissoschizine-derived pharmaceuticals. We identified a large group of VinBLAST homologues found beyond MIA-producing plant families, suggesting CAD-like proteins may have other uncharacterized functions across the plant kingdom. These findings reveal a critical role of non-catalytic scaffolding proteins in plant specialized metabolism.

## VinBLAST is an MIA synthesis activator

HYS and GS are the two primary CAD-like reductases that compete for the strictosidine aglycone in *C. roseus* leaves (Fig. 1B), where *HYS* expression is about 1/3 of *GS* expression (fig. S1). Although HYS interacts and colocalizes with SGD in the nucleus, GS-derived MIAs exceeds HYS-derived MIAs by over 12 folds *(19, 20)*. This disparity further implies that an additional factor may facilitate SGD-GS substrate channeling in planta. Our recent genomic analysis revealed clustering of CAD-like reductases in *C. roseus*, including a GS biosynthetic gene cluster containing *GS*, 8-hydroxygeraniol oxidoreductase *(8HGO) (21)*, and *O*-acetylstemmadenine oxidase *(ASO) (2)*, all involved in vinblastine biosynthesis, along with their homologous genes such as *GS2* and *THAS2* (Fig. 1A) *(22)*. We hypothesize that a component of this cluster may function as the endogenous link between SGD and GS.

Co-expression analysis indicated that two CAD-like genes *(CAD1* and *CAD2) (2)*, located immediately next to *GS* in the cluster, exhibited strong expression correlation to *STR, GS*, and *GO* (encoding geissoschizine oxidase) (Fig. 2A). This pattern prompted us to test their functions via virus-induced gene silencing (VIGS) experiments. We also targeted three additional homologous genes *CAD3*–*CAD5*, although not in the cluster *(2)*. *CAD1* and *CAD2* share 82% nucleotide identity, making co-silencing likely (fig. S2). Silencing *CAD3*–*CAD5* (~70% nucleotide identity with *CAD2)* had no effect on MIA profiles (fig. S3). However, silencing *CAD2* led to a striking decrease in catharanthine and vindoline levels, by 93.5% and 83.7%, respectively, and a substantial increase in HYS-derived MIAs, ajmalicine and serpentine, by 6.1- and 3.7-fold, respectively (Fig. 2B). This mirrored our earlier VIGS-GS results, where geissoschizine flux was redirected to HYS-derived products *(14)*. Compared with the empty vector (EV) control, VIGS-CAD2 plants showed an 88.7% decrease in *CAD2* transcripts, a modest 38.5% decrease in *CAD1*, and slightly increased *GS* transcript levels (Fig. 2C).

**Fig. 2.**
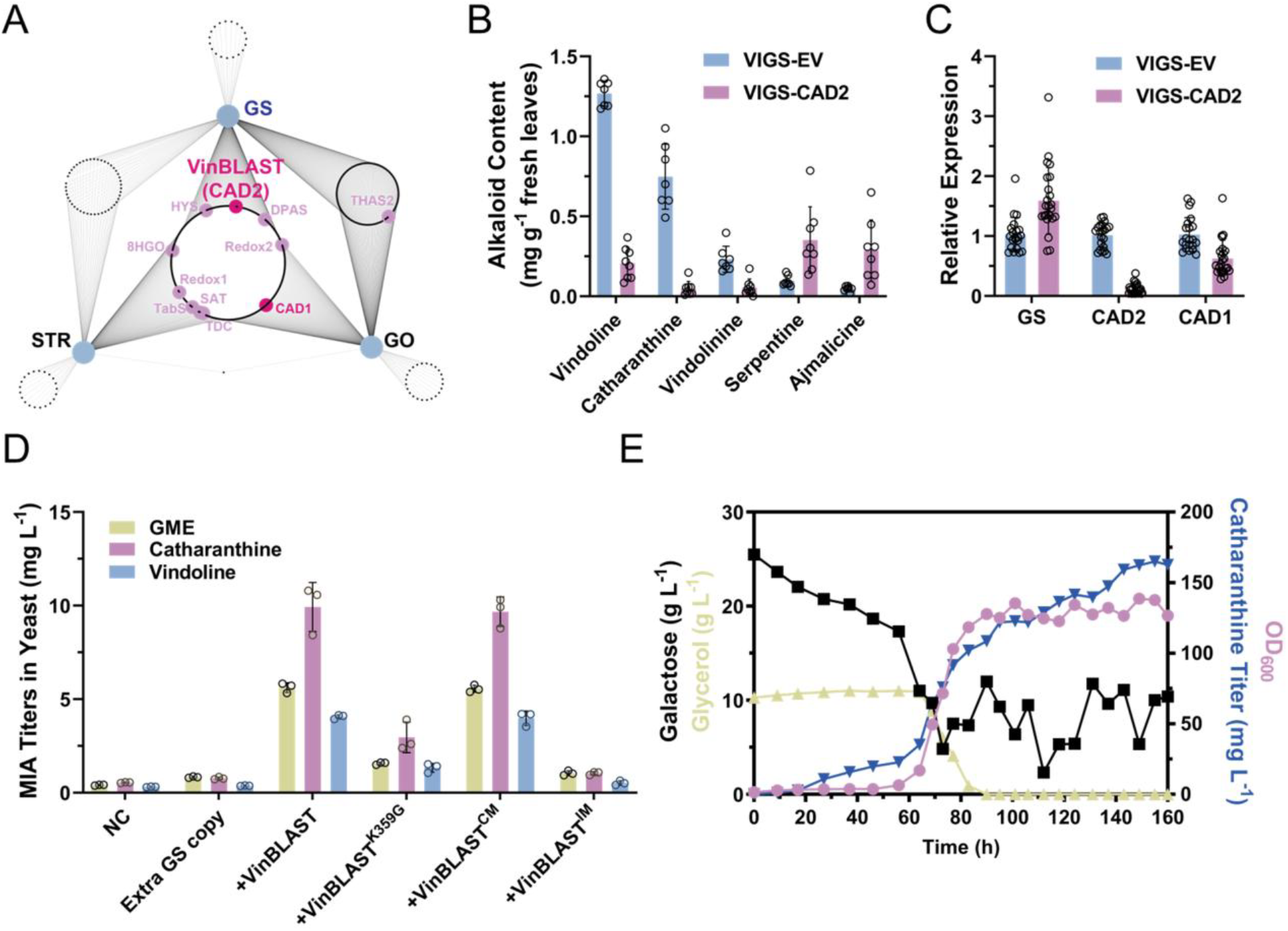
VinBLAST is essential for efficient MIA biosynthesis in *C. roseus* and yeast cell factories. **(A)** Weighted gene co-expression network analysis (WGCNA). A large part of MIA biosynthetic genes in *C. roseus* leaf and root tissues were clustered in the same module (blue module in Data S2). Co-expression correlations (r ≥ 0.9) between genes in the blue module and three bait genes *(STR, GS*, and *GO)* are shown. Enlarged light blue dots indicate the baits. Each dot means a gene, and the central circle of black dots indicates genes co-expressed with all the baits. Data S2 presents a list of genes co-expressed with one or more baits. VinBLAST and several key enzymes involved in catharanthine and vindoline biosynthesis co-expressed with all baits are indicated in enlarged magenta and purple dots, respectively. DPAS: dihydroprecondylocarpine synthase; Redox1/2: stemmadenine forming reductases; SAT: stemmadenine *O*-acetyltransferase; TabS: tabersonine synthase; 8HGO: 8-hydroxygeraniol oxidoreductase. **(B)** MIA contents following VIGS of *CAD2 (VinBLAST)* in *C. roseus* leaves. The results represent the mean ± s.d. of 7 or 8 biological replicates. The figure with statistical analysis is provided in Data S1. **(C)** The relative expression levels of *GS, CAD2 (VinBLAST)*, and *CAD1* in leaves of VIGS-CAD2 plants compared with the empty vector (EV) control. The results represent the mean ± s.d. of 7 or 8 biological replicates, each with 3 technical replicates. The figure with statistical analysis is provided in Data S1. **(D)** MIA titers in engineered *S. cerevisiae* strains producing GME (yellow), catharanthine (purple), and vindoline (blue). Negative Control (NC) refers to the base strains harboring the full biosynthetic pathways with a single *GS* copy, which were further engineered to include an extra copy of *GS, VinBLAST, VinBLAST*^*K359G*^, *VinBLAST*^*CM*^, and *VinBLAST*^*IM*^, respectively. VinBLAST^K359G^: the VinBLAST-SGD interaction mutant; VinBLAST^CM^: the catalytic mutant VinBLAST^C51A,H56A^; VinBLAST^IM^: the VinBLAST-GS interaction mutant VinBLAST^M298E,V299E^. Detailed strain information is provided in table S1. The results represent the mean ± s.d. of three biological replicates. The figure with statistical analysis is provided in Data S1. **(E)** Fed-batch fermentation profiles for strain CA02, which produces 164.9 mg L^-1^ catharanthine from simple carbon sources (glycerol and galactose). Galactose was continuously fed into the bioreactor to maintain its concentration at approximately 8 g L^-1^. Samples were taken every 5-10 hours to measure OD_600_ (purple), glycerol concentration (yellow), galactose concentration (black), and catharanthine titer (blue).

These findings suggested CAD2 as the potential link between SGD and GS. We next integrated a single copy of *CAD2* expression cassette into our *de novo* MIA-producing *S. cerevisiae* strains, which produced geissoschizine methyl ether (GME, corynanthe type), catharanthine (iboga type), and vindoline (aspidosperma type), one-, six-, and 13-step downstream of geissoschizine (Fig. 1B and fig. S4). This resulted in substantial increases in GME (13.7-fold), catharanthine (18.4-fold), and vindoline (13.1-fold) titers in 24-well plate cultures (Fig. 2D and table S1). Similarly, introducing *CAD2* into the catharanthine-producing *Pichia pastoris* strain CAN19 *(5)* yielded a 10.2-fold boost in producing catharanthine (fig. S5), confirming CAD2 function in a different chassis. In *S. cerevisiae*, further increasing *GS* and *CAD2* copy numbers only gave modest titer increase up to 36%, suggesting that a single copy of *CAD2* is sufficient to relieve the geissoschizine production bottleneck (fig. S6). In fed-batch fermentation, our *S. cerevisiae* strain CA02 produced 164.9 mg L^-1^ catharanthine from simple carbon sources (Fig. 2E). Based on these and subsequent results, we designate CAD2 as the **Vin**ca alkaloid **B**iosynthesis **L**ocalizing and **A**ctivating **S**caffold **T**ether (VinBLAST).

### VinBLAST links GS and SGD in the nucleus

To investigate the mechanism of this enhancement, we first conducted BiFC experiments to assess protein-protein interactions among SGD, GS, and VinBLAST in vivo. As expected, SGD exhibited strong self-interaction in the yeast nucleus, consistent with its known oligomeric structure; GS alone did not interact with SGD and mainly resided in the cytosol. When expressed alone, VinBLAST primarily localized to the cytosol. Strikingly, VinBLAST exhibited interaction with SGD in the nucleus and interacted with GS in the cytosol as well (Fig. 3A and figs. S7 to S9). When untagged VinBLAST was co-expressed in GS-SGD BiFC assays, we observed clear GS-SGD interaction in the nucleus, confirming that VinBLAST indeed is a previously unrecognized scaffolding protein that mediates the physical association between GS and SGD (Fig. 3B and fig. S9).

**Fig. 3.**
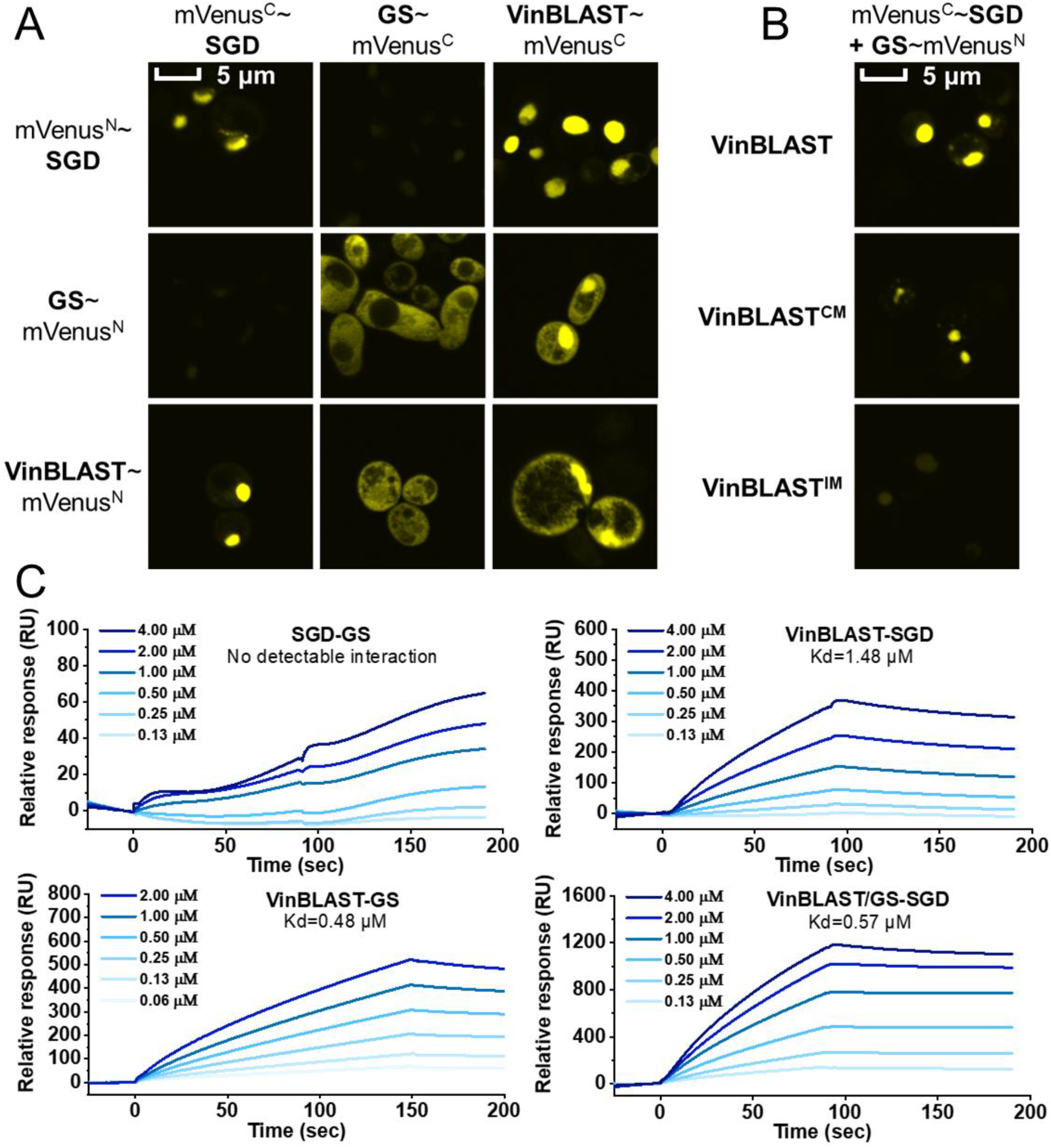
VinBLAST acts as a scaffold by dimerizing with GS, enabling the interaction between GS and SGD. **(A)** BiFC experiments in *S. cerevisiae*, indicating self-interactions of VinBLAST homodimer, GS homodimer, and SGD oligomer, and demonstrating the interaction of VinBLAST with GS in the cytosol and with SGD in the nucleus. Notably, GS alone does not interact with SGD. **(B)** VinBLAST-mediated GS and SGD association in the nucleus. GS and SGD interact in the yeast nucleus only after co-expression with untagged VinBLAST. Disruption of VinBLAST’s catalytic activity (VinBLAST^CM^) does not impact its scaffolding function, whereas disruption of VinBLAST dimerization (VinBLAST^IM^) abolishes the GS-SGD interaction. The mVenus fragments were fused to the *N*-terminus of SGD as well as *C*-termini of VinBLAST and GS. The detection of yellow fluorescence indicates protein-protein interaction. The images were acquired using a confocal laser scanning microscope at 600× magnification, with a 5-μm scale bar displayed in the upper left corner. The detailed imaging results are provided in figs. S8 and S9. **(C)** SPR assays to quantify the interactions among GS, VinBLAST, and SGD. While GS showed no detectable binding to SGD, interactions were detected for VinBLAST-SGD and VinBLAST-GS, with a Kd of ~1.48 μM and ~0.48 μM, respectively. Notably, VinBLAST and GS mixture further strengthened their interaction with SGD (Kd ≈ 0.57 μM). The purified proteins were injected over the immobilized ligand surface at gradient concentrations (0.06-4 μM).

We next employed surface plasmon resonance (SPR) assays to quantify the binding affinities among SGD, VinBLAST, and GS in vitro. While GS showed no detectable binding to SGD, VinBLAST interacted with SGD with a dissociation constant (Kd) of approximately 1.48 μM (Fig. 3C). Notably, a 1:1 mixture of VinBLAST and GS further strengthened their interaction with SGD (Kd ≈ 0.57 μM, Fig. 3C and fig. S10), supporting the conclusion that VinBLAST enhances the effective association between GS and SGD.

VinBLAST shares >70% amino acid identity with diverse plant CADs and reduces cinnamyl and coniferyl aldehydes to their corresponding alcohols (fig. S11). However, its catalytic efficiency is 144-fold lower than that of another *C. roseus* leaf CAD which may be physiologically responsible for lignin biosynthesis (fig. S12). Additionally, VinBLAST was inactive towards strictosidine aglycone (fig. S11). To decouple VinBLAST’s scaffolding role from its catalytic function, we mutated two key residues (C51A and H56A) required for NADPH binding and constructed the catalytic mutant (VinBLAST^CM^, VinBLAST^C51A,H56A^) with abolished CAD activity (fig. S11D). However, its capacity to interact with SGD or GS remained unchanged, as demonstrated by both BiFC (Fig. 3B) and yeast two-hybrid (Y2H, figs. S13 and S14). Expressing VinBLAST^CM^ in *S. cerevisiae* still resulted in comparable GME, catharanthine, and vindoline production levels as VinBLAST (Fig. 2D). These results confirm that the role of VinBLAST in scaffolding GS and SGD as well as enhancing MIA biosynthesis is independent of its CAD catalytic function.

To obtain direct evidence for the linkage function of VinBLAST, we investigated the interaction interface between VinBLAST and SGD. Molecular dynamics (MD) simulation identified K359 of VinBLAST as a critical residue for SGD binding (figs. S15 to S17). BiFC (fig. S18), Y2H (fig. S19), and SPR (fig. S20) assays confirmed that VinBLAST^K359G^ weakened, if not completely abolished, interaction with SGD while retaining its ability to bind GS. Furthermore, by replacing VinBLAST with VinBLAST^K359G^ in the engineered yeast strains, production-promoting effects on GME, catharanthine, and vindoline were dropped to 27.9%, 29.8%, and 32.8%, respectively, of the wild-type levels. (Fig. 2D). Together, these results demonstrate that VinBLAST-mediated physical linkage of GS and SGD is a critical driver of MIA biosynthesis.

We found that VinBLAST’s scaffolding function could be partially mimicked by artificially tethering GS and SGD using two independent self-assembling protein tag systems: the Regulatory Interaction and Anchoring Domain (RIAD) and Dimerization Domain (RIDD) from protein kinase A, and the SpyTag/SpyCatcher system derived from the *Streptococcus pyogenes* FbaB protein. By tagging GS and SGD with these interacting pairs, we evaluated MIA production in yeasts, with GME and vindoline titers increased by approximately 1.8-to 3.7-fold relative to the baseline strains (figs. S21 to S23). Intriguingly, even the best-performing artificial scaffolding strategies did not match the level of enhancement achieved by VinBLAST (figs. S24 and S25), suggesting that its functional role extends beyond simple tethering of GS and SGD.

### VinBLAST boosts GS catalytic activity

To investigate the effect of VinBLAST on GS activity, we performed coupled in vitro assays (figs. S26-28). VinBLAST, VinBLAST^CM^, and VinBLAST^K359G^ all exhibited comparable GS-enhancing activity. Kinetic analysis further revealed that VinBLAST enhanced the GS catalytic rate *(V*_*max*_) by 24.3-fold without affecting the substrate binding affinity *(K*m). These results strongly suggest altered GS active site architecture in the VinBLAST-GS complex.

To probe the molecular mechanism underlying enhanced GS activity, we performed MD simulations of VinBLAST-GS complex. Given the conserved β-sheet-stabilized homodimer architecture shared among GS, HYS, and related CADs *(16)*, we hypothesized that VinBLAST-GS might form an analogous heterodimer through similar structure motif. Native polyacrylamide gel electrophoresis confirmed the heterodimer formation (fig. S29). To validate the hypothesis, we first simulated the GS-GS homodimer interaction (fig. S30), identifying I301 as a critical interfacial residue, forming reciprocal amide backbone interactions across the β-sheet interface.

Substituting I301 with glutamate (I301E, GS interaction mutant GS^IM^) disrupted the dimer interface, increasing the β-sheet separation from ≈5 Å to ≈10 Å in silico (figs. S30F and S30G). This abolished GS self-interaction in BiFC assay and eliminated its catalytic activity both in vitro and in vivo (figs. S31 and S32), demonstrating the essential role of GS dimerization in catalysis. MD simulations revealed that dimer disruption exposed the active site to solvent, thermodynamically disfavoring substrate binding. Over 1,000 ns, snapshot showed complete substrate dissociation from the monomeric GS active site (fig. S33), explaining the loss of catalytic activity in GS^IM^ compared with the WT homodimer.

Further MD simulations of the VinBLAST-GS heterodimer identified M298 and V299 as key residues mediating heterodimerization via amide backbone interactions (Figs. 4A, 4B and fig. S34). To validate this, we constructed a VinBLAST-GS interaction mutant VinBLAST^IM^ (VinBLAST^M298E,V299E^) to disrupt heterodimer formation with GS^IM^ through electrostatic repulsion. BiFC and Y2H showed weakened interaction between VinBLAST^IM^ and GS^IM^ as well as wild-type GS (Fig. 3B and figs. S9, S13, and S31). These observations were corroborated by in vitro pull-down assays: His-tagged VinBLAST and VinBLAST^CM^ successfully retained non-tagged GS on affinity columns, whereas VinBLAST^IM^ failed (fig. S35). Accordingly, the production-promoting effects of VinBLAST^IM^ on GME, catharanthine, and vindoline were dropped to 18.7%, 10.5%, and 12.8%, respectively, of the wild-type levels (Fig. 2C). These findings suggest that VinBLAST forms a heterodimer with GS to both facilitate scaffolding and enhance catalytic activity.

**Fig. 4.**
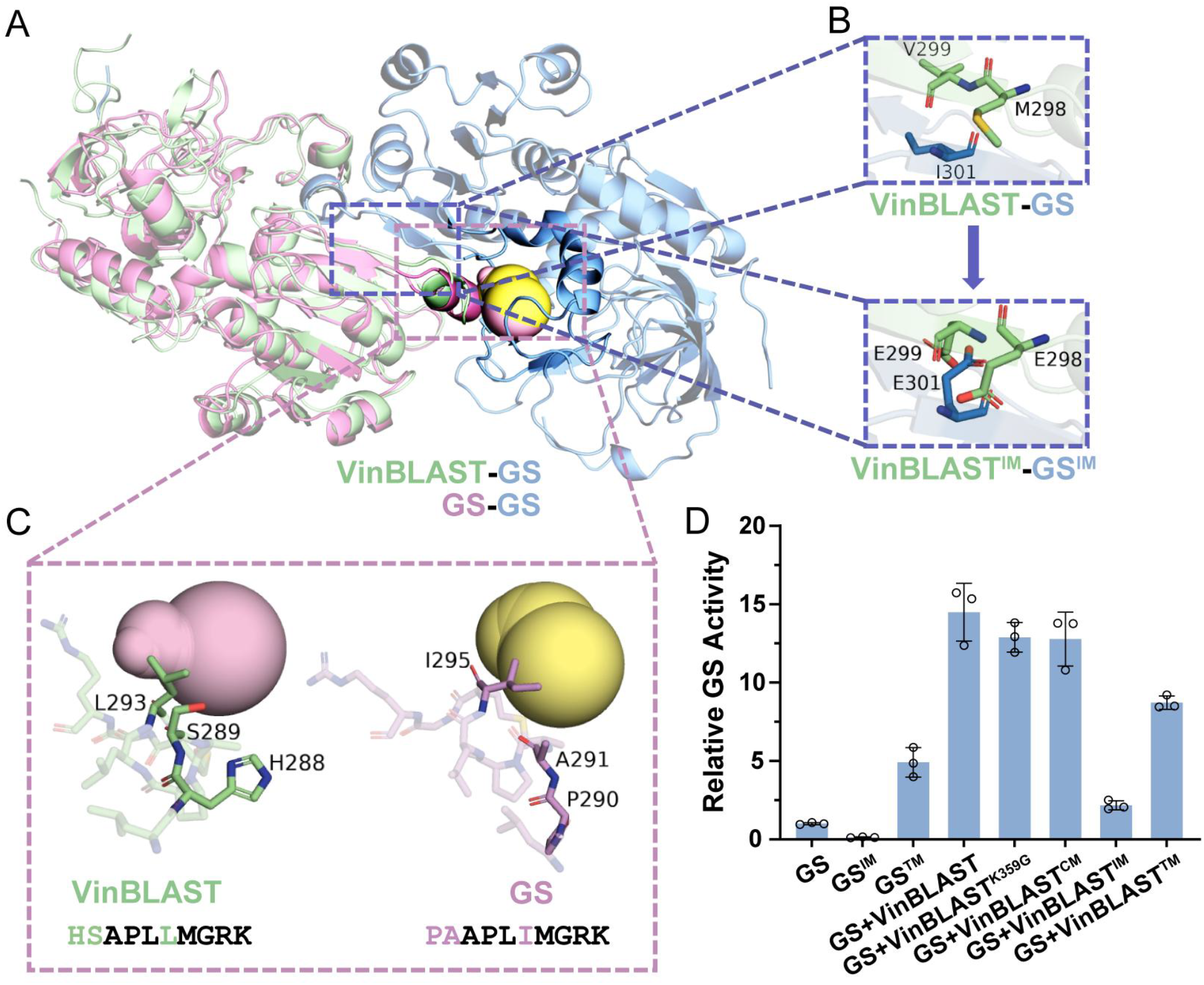
Dimerization between VinBLAST and GS reshapes the substrate tunnel at their heterodimer interface. **(A)** Overlay schematic diagram comparing the substrate tunnel in the VinBLAST-GS heterodimer (purple spheres) with that in the GS-GS homodimer (yellow spheres). The X-ray crystal structure (PDB ID: 8A3N) is selected as the template for GS homodimer and the binary structure of VinBLAST-GS heterodimer is predicted by AlphaFold3. Residues involved in tunnel formation are highlighted. The VinBLAST monomer is colored cyan, while the two monomers of GS are depicted in purple and blue, respectively. **(B)** VinBLAST and GS dimer interface. Upper dashed rectangle: structural details of interacting β-sheet pair at the VinBLAST-GS interface; bottom dashed rectangle: structural details of interacting β-sheet pair at the VinBLAST^IM^-GS^IM^ interface; VinBLAST^IM^: the interaction mutant VinBLAST^M298E,V299E^; GS^IM^: the interaction mutant GS^I301E^. **(C)** Close-up of amino acid residues from VinBLAST and GS participating in substrate tunnel formation. Key segments in VinBLAST: HSAPLLMGRK (288-297), shown in cyan sticks; GS: PAAPLIMGRK (290-299), shown in purple sticks. Different residues between the segments are highlighted. **(D)** Effects of VinBLAST on GS in vitro activity. The activity of GS homodimer (set as 1) was compared with GS mutant as well as that with the addition of VinBLAST or VinBLAST mutants. GS™: GS tunnel mutant GS^P290H,A291S,I295L^; VinBLAST™: VinBLAST tunnel mutant VinBLAST^H288P,S289A,L293I^. The results represent the mean ± s.d. of three technical replicates. The figure with statistical analysis is provided in Data S1.

### VinBLAST-GS reshapes the substrate tunnel

To investigate the mechanism of the rate accelerating upon VinBLAST-GS dimerization, we used MD simulations (figs. S36 to S38) to obtain the dynamic information of 4,21-dehydrogeissoschizine, the direct GS substrate among the various strictosidine aglycone isoforms in equilibrium, at the active site of GS (Figs. 4A and 4C). The near-attack conformation (NAC) frequency analysis *(23, 24)* revealed the VinBLAST-GS heterodimer exhibited higher occurrence (41.6%) compared with the GS-GS homodimer (23.0% and 26.5% at individual sites) (fig. S37). This increased NAC frequency suggests more efficient attainment of transition-state geometry, aligning with the observed rate acceleration in kinetic assays.

The heterodimer’s enhanced catalytic efficiency may suggest its substrate tunnel enable more efficient substrate entry than the homodimer. To test whether the tunnel structure alone could enhance catalytic performance, we engineered a GS tunnel mutant (GS™, GS^P290H, A291S, I295L^) to mimic the tunnel architecture observed in VinBLAST-GS heterodimer (Fig. 4C). GS™ exhibited a 5.3-fold increase in *V*_*max*_ (Fig. 4D and fig. S28). Expressing GS™ in *S. cerevisiae* also resulted in 2.0~2.7-fold increase in MIA production relative to the wild-type GS, whose production-promoting effect was further increased to 4.9~7.5-fold by tethering GS™ and SGD using artificial protein scaffolds (fig. S39). Conversely, we engineered a reciprocal VinBLAST tunnel mutant (VinBLAST™, VinBLAST^H288P, S289A, L293I^) to adopt a GS-like tunnel structure.

VinBLAST™ modestly decreased the heterodimer activity by 24.9% (Fig. 4D and fig. S28). These results confirm that VinBLAST-GS interaction reshapes the substrate tunnel and the VinBLAST-GS tunnel architecture is crucial for enhancing GS catalytic efficiency.

### VinBLAST is conserved beyond Gentianales

As geissoschizine is a central precursor to over 700 MIAs in nature, VinBLAST homologues are likely widespread among MIA-producing plant families. Syntenic analysis confirmed the presence of conserved VinBLAST-GS gene clusters in MIA-producing species in Gentianales, including *Rauvolfia tetraphylla* (Apocynaceae), *Gelsemium sempervirens* (Gelsemiaceae), and *Mitragyna speciosa* (Rubiaceae) (Fig. 5A). The gene cluster is also found in non-MIA producing Gentianales species, such as *Calotropis gigantea* (Apocynaceae) and *Eustoma grandiflorum* (Gentianaceae) and in *Nepeta mussinii* (Lamiales) and *Solanum lycopersicum* (Solanales) (Fig. 5A). The synteny can even be traced back in *Vitis vinefera* (grapevine), an early diverging core eudicot that split from other lineages ~148 million years ago, suggesting this ancient synthetic region served as an evolutionary hotspot for emergence of CAD-like reductases, including GS and VinBLAST, across many plant lineages *(22)*.

**Fig. 5.**
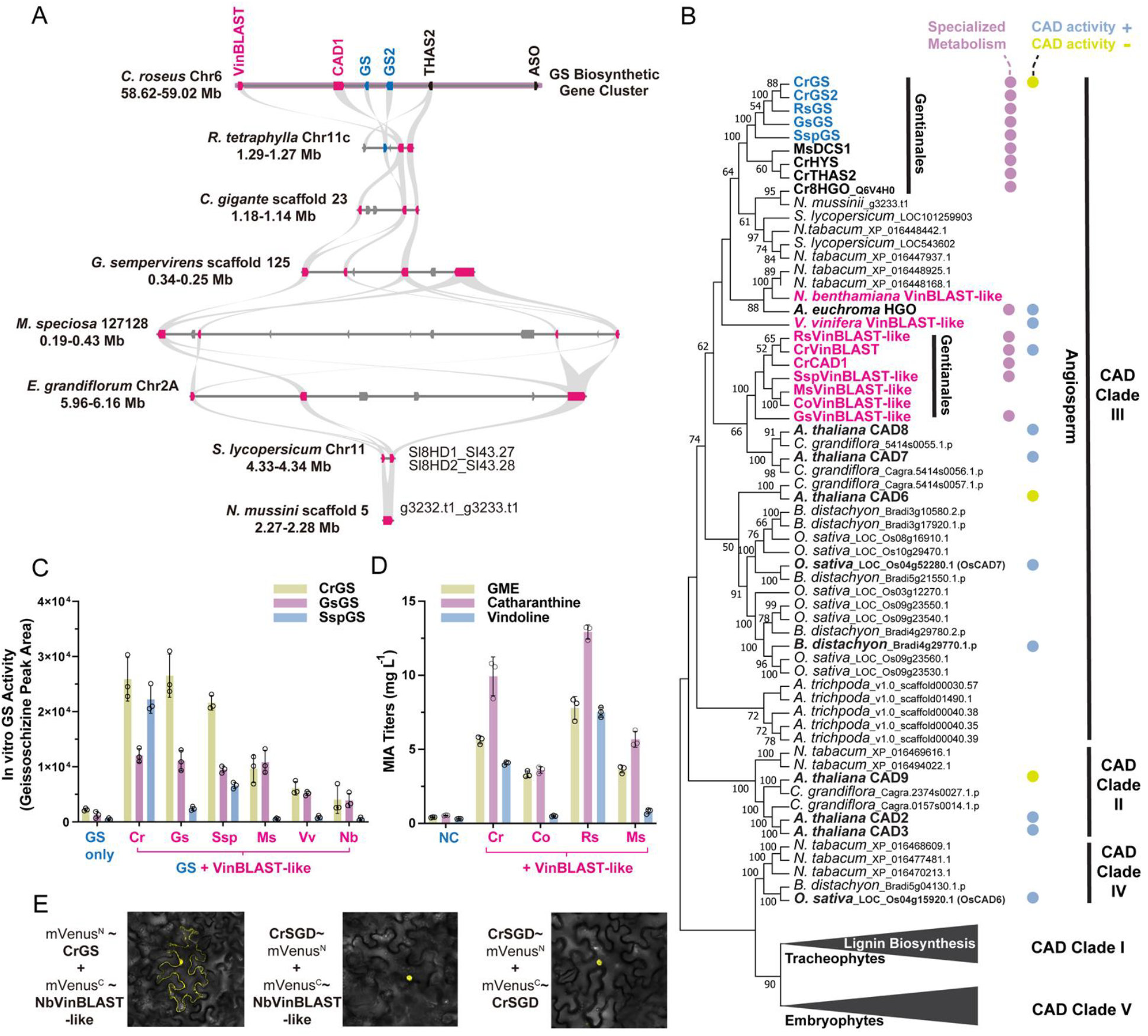
VinBLAST activity extends beyond Gentianales. **(A)** Synteny analysis. The *C. roseus* GS biosynthetic gene cluster (magenta shade) is syntenic to a CAD-rich genomic locus found in both MIA-producing *(Rauvolfia tetraphyllam, Gelsemium sempervirens*, and *Mitragyna speciosa)* and non-producing species *(Calotropis gigantea*, and *Eustoma grandiflora)* in the order of Genianales, as well as in species outside of this order *(Solanum lycopersicum* in Solanales and *Nepeta Mussinii* in Lamiales). Each box represents a gene locus, with arrows indicating transcriptional direction. Ribbons connect syntenic gene pairs (grey), with GS and VinBLAST homologues shown in blue and magenta, respectively. **(B)** Phylogenetic analysis. VinBLAST and GS homologues are placed within CAD clade III, with the full phylogenetic tree provided in fig. S40. CAD clade III comprises only angiosperms, including early diverging *Amborella trichopoda*, monocots, and eudicots. Members with demonstrated in vitro CAD activities are marked with blue dots, while those experimentally shown to lack CAD activity marked with yellow dots. Proteins implicated in specialized metabolism are marked with purple dots. Unlike the CAD clade I, which contains tracheophyte members with genetically and biochemically supported roles in lignin biosynthesis, the lignin-related functions of CADs in other clades have not been explicitly demonstrated, despite some in vitro evidence. **(C)** In vitro activities of CrGS *(Catharanthus roseus*; Apocynaceae family), GsGS *(Gelsemium sempervirens*; Gelsemiaceae family), and SspGS *(Strychnos spinosa*; Loganicaceae family) when paired with VinBLAST homologues from *Cr, Gs, Ssp, Ms (Mitragyna speciosa;* Rubiaceae family), *Vv (Vitis vinifera;* Vitales order), and *Nb (Nicotiana benthamiana;* Solanales order). Yellow represents CrGS, purple represents GsGS, and blue represents SspGS. The results represent the mean ± s.d. of three technical replicates. The figure with statistical analysis is provided in Data S1. **(D)** MIA titers of GME, catharanthine, and vindoline in engineered *S. cerevisiae* strains. Negative controls (NC) strains lacking VinBLAST expression are compared with strains expressing a single copy of VinBLAST homologues from *Cr, Co (Cephalanthus occidentalis;* Rubiaceae family), *Rs (Rauvolfia serpentina;* Apocynaceae family), and *Ms* species. Detailed strain information is provided in table S1. Yellow represents the GME-producing strains, purple represents the catharanthine-producing strains, and blue represents the vindoline-producing strains. The results represent the mean ± s.d. of three biological replicates. The figure with statistical analysis is provided in Data S1. **(E)** BiFC experiments in *N. benthamiana* leaf epidermis. NbVinBLAST and CrGS interact mainly in the cytosol (also diffusing into the nucleus), NbVinBLAST and CrSGD interact in the nucleus, and CrSGD monomers interact in the nucleus. Images show overlays of yellow fluorescence and transmitted light microscopy. The detailed imaging results are provided in fig. S41.

A phylogenetic analysis placed VinBLAST, GS, and their homologues in CAD clade III comprising angiosperms *(25)* (Fig. 5B and fig. S40). In contrast to several clade I CADs, whose roles in lignin biosynthesis are well supported by genetic and biochemical evidence, the in vivo function of CADs in other clades are largely unknown, although in vitro CAD activity has been documented for some members *(26-32)*. Given the high sequence similarity of VinBLAST (>70% amino acid identity) to bona fide and putative CADs, its activity in scaffolding SGD and GS or boosting GS activity may be found in other homologous CADs in clade III.

To validate this hypothesis, we conducted in vitro enzyme assays and in vivo yeast fermentation tests, pairing VinBLAST homologues from both MIA producing and non-producing species and GS homologues from three plant families (table S3). All tested VinBLAST homologues enhanced GS in vitro activity (Fig. 5C). For instance, *Strychnos spinosa* (Loganiaceae) VinBLAST enhanced its own SspGS in vitro activity by 13.2-fold, while *C. roseus* VinBLAST boosted SspGS activity by an impressive 44.5-fold. Remarkably, VinBLAST homologues from grapevine and *Nicotiana benthamiana* (tobacco) (Fig. 5B), distantly related species not known for MIA production, enhanced *C. roseus* GS activity by 2.7- and 1.8-fold, respectively. In *S. cerevisiae*, VinBLAST homologues increased the production of GME, catharanthine, and vindoline to varying degrees, with peak improvements of 19.0-fold, 23.9-fold, and 24.2-fold, respectively (Fig. 5D). BiFC experiment further supported the scaffolding function of the *N. benthamiana* VinBLAST homologue, showing its interaction with CrGS in the cytosol and CrSGD in the nucleus (Fig. 5E and fig. S41). With the widespread VinBLAST activity in plants and the superior performance of those from MIA-producing species, our results support that VinBLAST evolves from ancestral CADs to acquire a specialized, indispensable scaffolding role in MIA biosynthesis.

## Conclusions

Here we report the discovery and functional characterization of VinBLAST, a lignin biosynthesis related enzyme (CAD) repurposed as a scaffold and activator essential for MIA biosynthesis. Although VinBLAST retains detectable CAD activity, its catalytic efficiency is two orders of magnitude lower than that of a clade I CAD from *C. roseus* leaves (fig. S12), making it less likely to be a major player for lignin biosynthesis. Instead, VinBLAST controls MIA metabolic flux in planta and enables efficient production by physically tethering SGD and GS. This tethering facilitates processing of the unstable strictosidine aglycone intermediate while allosterically enhancing GS enzymatic activity. We show that the geissoschizine biosynthesis-enhancing activity is conserved among a group of CAD-like proteins from diverse MIA producing families, and is also present in distantly related species such as grapevine and tobacco. As a pivotal branch point in MIA biosynthesis, geissoschizine gives rises to major MIA classes (aspidosperma, sarpagan, iboga, akuammiline, and derivative bis-MIAs) *(33)*. These MIAs include well-known and extensively studied MIA pharmaceuticals: vinblastine and derivatives (anticancer), ajmaline (antiarrhythmic), ibogaine (psychoactive), strychnine (curare poison), gelsemine (glycine receptor agonist), conolidine (non-opioid painkiller), and voacamine (phytocannabinoid). This breadth highlights the potential of VinBLAST for scalable production of all geissoschizine-derived therapeutics.

At the metabolic organization level, protein scaffolds have been recently reported in several specialized plant metabolic pathways. For example, a cellulose synthase-like protein has dual functions as a scaffold and a cholesterol glucuronosyltransferase in steroidal glycoalkaloid biosynthesis in tomato *(34, 35)*. Similarly, a non-catalytic scaffold protein is identified in paclitaxel biosynthesis *(36)*. Extensive studies on protein–protein interactions and metabolon formation in flavonoid biosynthesis, particularly among cytochrome P450 monooxygenases and dioxygenases, further reveal that plant specialized metabolism is far more modular and spatially organized than previously understood *(37, 38)*. Geissoschizine is a central precursor for multiple major MIA classes and is further processed by numerous downstream enzymes. It is therefore plausible that interactions among SGD, GS, and VinBLAST serve as an anchoring module for larger, modular MIA metabolons.

The discovery and characterization of VinBLAST led to the construction of a yeast cell factory for *de novo* production of catharanthine at an impressive titer of ~160 mg L^-1^. Our study sheds new light on the molecular organization of MIA biosynthesis and hidden functions of diverse CAD-like reductases in nature. Moreover, we establish VinBLAST as a previously unrecognized, indispensable protein scaffold enabling efficient MIA biosynthesis in microbial cell factories.

## Acknowledgments

We appreciate Prof. Huimin Zhao from the University of Illinois at Urbana-Champaign for insightful discussion and suggestions. We also would like to thank iBioFoundry and Core Facility at ZJU-Hangzhou Global Scientific and Technological Innovation Center for analytical support.

## Author contributions

Conceptualization, supervision, and funding acquisition: JL, YQ, and YW BiFC experiments: DG, CC, and HCT

In vitro assays and pull-down experiments: SGAM and JJOG-G

Homology modeling and MD simulations: BC, YG, and MBR

VIGS experiments: MS

Synteny and co-expression analysis: CC

*P. pastoris* experiments: XJ

Y2H experiments and bioreactor fermentation: DG, YG, and JB

Cloning and strain construction: DG, SGAM, YG, CC, JJOG-G, XJ, HCT, JB, JNL, JOP, JH, and FD

SPR experiments: CQ

Writing: DG, BC, CD, LH, VDL, YW, YQ, and JL

## Data, code, and materials availability

All data are available in the main text or the supplementary materials. No new code was generated in the course of this study. Correspondence and requests for materials should be addressed to JL and YQ under materials transfer agreements.

## Supplementary Materials

Materials and Methods

Supplementary Text

Figs. S1 to S44

Tables S1 to S10

References*(39-55)*

Data S1 to S3

MDAR Reproducibility Checklist

