## Supplementary Materials for "A cinnamyl alcohol dehydrogenase-like scaffold organizes monoterpenoid indole alkaloid biosynthesis"

**The PDF file includes:**

Materials and Methods
Supplementary Text
Figs. S1 to S44
Tables S1 to S10

**Other Supplementary Material for this manuscript includes the following:**

Data S1 to S3
MDAR Reproducibility Checklist

### Materials and Methods

#### Chemicals

Geissoschizine methyl ether (GME) standard was purchased from MedChemExpress (New Jersey, USA). Catharanthine and vindoline standards were purchased from Yuanye Bio-Technology Co. Ltd (Shanghai, China). Vindolinine, ajmalicine, serpentine, tryptamine, and secologanin standards were purchased from Sigma-Aldrich (Millipore Sigma, Burlington, MA, USA). Vindorosine and epi-vindolinine standards were previously purified from *Catharanthus roseus* plant (*19*). Strictosidine aglycone was produced as previously described with modifications (*14*). Briefly, 20 mg of tryptamine was dissolved in 2 mL methanol, and 10 mg of secologanin was dissolved in 1 mL methanol. These were added to 100 mL of 20 mM Tris buffer (pH 7.5) along with 2 mg of purified recombinant STR, and the reaction was incubated at 30°C for 1 h. Then, 0.5 mg of purified recombinant SGD was added, and the reaction was continued at 30°C for 20 min. During this step, strictosidine aglycone, which has poor water solubility, precipitated from the solution along with proteins. After centrifugation at 7,000 g for 10 min, the pellet was dissolved in methanol, centrifuged to remove impurity, and quantified by LC-MS/MS using an ajmalicine standard curve. This yielded 2.7 mg of strictosidine aglycone for in vitro assays. All the other chemicals were obtained from Sigma-Aldrich, unless specifically indicated above. For molecular biology, all restriction enzymes, T4 DNA ligase, Q5 polymerase, and NEBuilder® HiFi DNA Assembly mixture were purchased from New England Biolabs (Beijing, China) and ThermoFisher (Waltham, MA, USA).

#### Cloning

VinBLAST and GS homologues were amplified from cDNAs of respective plant species. *MsVinBLAST* and *NbVinBLAST* were cloned in pDEST17 vector using Gateway® BP and LR clonase II according to manufacturer's protocol (ThermoFisher, Waltham, MA, USA). All remaining genes were cloned into pET-30b(+) vector by restriction/ligation (*Bam*HI/*Sac*I; *Bam*HI/*Sal*I) or Gibson Assembly. *VvVinBLAST* was synthesized and subcloned in pET-28a(+) between *EcoR*I/*Not*I sites (Twist Bioscience, CA, USA). To create non-histagged GS and VinBLAST proteins, these genes were amplified and cloned into pET-30b(+) digested with *Nde*I/*Sal*I. These vectors were transformed to *E. coli* BL21(DE3) cells for recombinant protein expression and purification. All plasmids used for the yeast BiFC assay were constructed using the pESC-URA/HIS/LEU/TRP series of vectors. Specifically, the mVenus$^{C}$ or mVenus$^{N}$ fragment and the relevant gene were assembled under the control of the *GAL10* promoter via one-step, multi-fragment Gibson assembly. A flexible linker was introduced between the gene and the mVenus$^{C/N}$ tag. To create BiFC vectors for tobacco transient expression, the TMV promoter and mVenus$^{N}$ or mVenus$^{C}$ fragments were amplified from pGTQL1211YN or pGTQL1221YC. The fragments were assembled using Gibson Assembly into pGTQL1211 digested with *Xho*I/*Spe*I, to generate VinBLAST and GS with *N*-terminal mVenus fusion. For SGD with *C*-terminal mVenus$^{C}$ fusion, the gene was directly gateway-cloned into pGTQL1221YC. These vectors were mobilized to *Agrobacterium tumefaciens* strain LBA4404 for transient expression in tobacco leaves. For VIGS of *CrCAD1-5*, the gene fragments were amplified from *C. roseus* cDNA and cloned into pTRV2 vector within *EcoR*I site. For all the genomic modifications in *S. cerevisiae*, the exogenous genes were subcloned into the multiple cloning sites (MCSs) of pESC series vectors (pESC-HIS, pESC-TRP, pESC-LEU, and pESC-URA) by restriction/ligation (*Bam*HI/*Xho*I; *Bam*HI/*Sal*I; *Not*I) or Gibson Assembly. New cDNA sequences obtained in this study were deposited to NCBI Genbank with the following accession numbers: PV770557-PV770565 (table S3). *CAD1-5* nucleotide sequences were provided in table S4. Recombinant plasmids constructed in this study were listed in table S5. The complete sequences of all yeast BiFC plasmids are provided in table S6. Primers used in this study were listed in table S7.

Construction of genome-edited *S. cerevisiae* strains

All genomic modifications in *S. cerevisiae* strains were performed using the CRISPR/Cas9 system. Exogenous gene cassettes, flanked by 40 bp homology arms matching the integration sites, were amplified via polymerase chain reaction (PCR). For sgRNA plasmid construction, spacer sequences designed using Benchling (https://benchling.com) were cloned into the *Bsa*I restriction sites of pRS423-SpSgH and/or pRS426-SpSgH. Subsequently, 800 ng of sgRNA plasmids and 800 ng of donor DNA fragments (homology arm-containing cassettes) were co-transformed into *S. cerevisiae* competent cells expressing Cas9, employing the LiAc/ssDNA/PEG method (*39*). Transformants were selected on SED-URA/G418 or SED-HIS/G418 agar plates. Positive clones were initially screened by colony PCR and further verified by DNA sequencing. All genome-edited *S. cerevisiae* strains constructed in this study are summarized in table S1, while the corresponding genomic integration sites are provided in table S8.

Yeast 24-well plates fermentation

For yeast strains harboring the full MIA biosynthetic pathway, single colonies were picked from fresh YPD agar plates and inoculated into 3 mL YPD medium. Following 24 h of cultivation at 30°C with shaking at 250 rpm, 30 μL of the seed culture was transferred to 24-well plates containing 3 mL fresh YPD medium and grown under identical conditions for an additional 24 h. Cells were then harvested by centrifugation at 4,000 rpm for 5 min and washed twice with sterile water to eliminate residual glucose. The collected cells were resuspended in 3 mL fresh YP medium supplemented with 2% (w/v) galactose to induce exogenous gene expression, followed by incubation at 30°C with 250 rpm shaking. After fermentation, the broth was extracted with an equal volume of ethyl acetate. The organic (upper) phase was filtered through a 0.22 µm membrane and subjected to qualitative and quantitative analysis.

Yeast fed-batch fermentation

Fed-batch fermentation was conducted using a T&J Intelli-Ferm B series fermentation system equipped with two 5-L glass bioreactors from T&J Bio-engineering Co., LTD (Shanghai, China), with the fermentation medium containing 10 g $L^{-1}$ yeast extract, 20 g $L^{-1}$ peptone A, 10 g $L^{-1}$ glycerol. The feeding solution consisted of 400 g $L^{-1}$ galactose as carbon source, 36% acetic acid and base solution (100 mL water with 200 mL ammonium hydroxide) for pH adjustment, 100 g $L^{-1}$ yeast extract and 200 g $L^{-1}$ peptone A as nitrogen source, and the antifoam agent. The pH was maintained at 6.0 throughout the fermentation process, with dissolved oxygen controlled at 30% saturation. Fermentation was initiated at an $OD_{600}$ of 1.5, followed by continuous galactose feeding. Temperature was maintained at 30°C, and the stirring speed was dynamically adjusted in response to real-time dissolved oxygen levels through a cascaded control strategy to maintain optimal oxygen transfer efficiency. During cultivation, samples were periodically collected for analysis, and the feeding rate was adjusted based on real-time monitoring of cell growth and metabolic activity to optimize production efficiency.

Virus-induced gene silencing experiments

The VIGS experiments were carried out as described previously (*40*) with a phytoene desaturase (PDS) silencing indicator. In brief, cells from overnight cultures of *A. tumefaciens* carrying pTRV1 and various pTRV2 vectors grown at 28°C shaking incubator were collected by centrifugation and resuspended in infection buffer (10 mM MES pH 5.6, 10 mM $MgCl_2$, 0.2 mM acetosyringone) to $OD_{600}$ of 1.5. The suspensions were incubated at 28°C for 2.5 h, then equal pTRV1 and pTRV2 suspensions were mixed just before the infection. A sharpened toothpick dipped in the mixed suspension was used to pierce just underneath the apical meristem of 4-week-old *C. roseus* cv. Pacifica XP with 2-3 pairs of true leaves, then 0.1 mL suspension mixture was used to flood the wound. The infected plants were grown at a glasshouse with 18/6 h photoperiod at 25°C for approximately 4 weeks, when the VIGS-PDS leaves developed white sectors. The developing leaves were cut in half along the main vertical vein. One half of the leaves were used for RNA extraction and qPCR, whereas the other half leaves were ground and extracted with methanol for alkaloid analysis using LC-MS/MS. The sample size ($n$ = 6 or 7 plants per group) was selected based on prior experience and common practice in *C. roseus* VIGS studies to ensure adequate biological replication for assessing variability in silencing efficiency and metabolite production.

cDNA synthesis and qRT-PCR analysis

Plant leaves (10-100 mg) were ground in liquid nitrogen using a small pestle and a microtube. Total RNA was extracted with TRI Reagent™ (ThermoFisher, Waltham, MA, USA) according to the manufacture's protocol. The cDNA was synthesized using the LunaScript® RT SuperMix (New England Biolabs, Ipswich, MA, USA) according to the manufacture's protocol. qRT-PCR experiments were performed using primers 40-45 on an Agilent AriaMx Real-Time PCR System (Santa Clara, CA, USA), employing the SensiFAST SYBR No-ROX qPCR 2X Master Mix (FroggaBio, Concord, Canada) following the manufacturer's protocol. The qRT-PCR (10 µL, 5 ng total RNA) cycles included 40 cycles of 95°C for 10 sec and 58°C for 30 sec. The Ct values and standard ΔΔCt method was used to quantify gene expression levels, which were normalized by using the expression of *C. roseus* 60S ribosomal RNA housekeeping gene (*40*).

Recombinant protein expression and purification

Luria-Bertani (LB) media (200 mL) were first inoculated with 2 mL overnight *E. coli* BL21(DE3) cultures harboring various genes, then grown to $OD_{600}$ 0.6-0.7 at 37°C. The cultures were induced with 0.1 mM IPTG at room temperature (for STR and SGD) or 15°C (for the rest constructs), 200 rpm overnight. The harvested cells were sonicated in lysis buffer (20 mM Tris-HCl pH 7.5, 100 mM NaCl, 10 mM imidazole, 10% (v/v) glycerol). After centrifugation at 10,000 g for 10 min, the supernatants were incubated with 1 mL Ni-NTA resin at 4°C for 20 min. The resins were washed with 20 mL wash buffer (20 mM Tris-HCl pH 7.5, 200 mM NaCl, 30 mM imidazole, 10% (v/v) glycerol). The recombinant proteins were eluted with elution buffer (20 mM Tris-HCl pH 7.5, 100 mM NaCl, 250 mM imidazole, 10% (v/v) glycerol), desalted with a PD-10 desalting column (Citiva, Marlborough, MA, USA) into 20 mM Tris-HCl pH 7.5, 100 mM NaCl, 10% (v/v) glycerol, and stored at -80°C.

In vitro assays

In vitro assays were conducted using 0.4 µg of purified GS, either alone or in combination with 0.4 µg of purified VinBLAST, in a 100 µL reaction containing 1 mM NADPH and 20 mM Tris-HCl at pH 7.5. Strictosidine aglycone (1.5 µM) served as the substrate. Reactions were performed in triplicate and incubated in a 30°C water bath for 25 min. To assess VinBLAST's CAD activity, 100 µL reactions were prepared containing 20 mM Tris-HCl at pH 7.5, 1 mM NADPH, 2 µg of either cinnamyl or coniferyl aldehyde, and 1 µg purified VinBLAST. Reactions were incubated at 30°C for 1 h. Reactions were quenched by adding an equal volume of methanol, followed by centrifugation. The supernatants were analyzed by LC-MS/MS.

VinBLAST-GS kinetics assays

Kinetic reactions (100 µL) were carried out in 20 mM Tris-HCl buffer at pH 7.5, containing 1 mM NADPH, and either 2 µg of purified GS alone or a combination of 0.5 µg GS and 0.5 µg VinBLAST. Strictosidine aglycone was tested at concentrations ranging from 0.1875 µM to 24 µM. To ensure consistent kinetic measurements, reaction mixtures containing all components except the substrate were pre-warmed at 30°C in a water bath. Reactions were initiated by adding 5 µL of strictosidine aglycone in methanol. For GS-only reactions, the incubation time was 5 min; for VinBLAST-GS complex assays, the reaction proceeded for 1 min. Reactions were quenched with an equal volume of methanol and analyzed by LC-MS/MS. Reaction conditions are critical for GS kinetics assays due to the inherent instability of strictosidine aglycone. This substrate gradually precipitates from solution and acts as a non-specific electrophile, which can aggregate and inactivate proteins, particularly problematic at concentrations over 3 µM for durations exceeding 20 min.

In vitro pull-down assays

Purified His-tagged GS or VinBLAST (1 µg) were first incubated with Ni-NTA resin (approximately 1:5 v/v; 50 µL slurry) in sample buffer (20 mM Tris-HCl, pH 7.5, 100 mM NaCl, 10% (v/v) glycerol) for 30 min on ice with gentle agitation. Following this, total *E. coli* lysate (3.5 µg) containing GS or VinBLAST lacking a His-tag was added to the slurry and incubated for an additional 30 min on ice. Then, the resin was pelleted by centrifugation (1,000 rpm), and the supernatant was discarded. The resin was washed twice with sample buffer by resuspension and centrifugation to remove unbound proteins. The washed, protein-bound resin was then directly used in enzymatic assays to estimate pull-down efficiency. For GS activity, a 100 µL reaction was set up in 20 mM Tris-HCl (pH 7.5) containing 1 mM NADPH and 1 µg strictosidine aglycone. For CAD activity, the same buffer conditions were used with 1 µg coniferyl aldehyde as the substrate. The assays were incubated for 1 h at 30°C and quenched by adding an equal volume of methanol. Supernatants were analyzed by LC-MS/MS to quantify product formation.

MD simulations

The X-ray crystal structure (PDB ID: 8A3N) including wide type GS homodimer with NADPH was selected as the template structure in this work. The structure of GS monomer was extracted from it and used as the initial structure for the relevant MD simulations. The binary structure of VinBLAST-GS heterodimer with NADPH was predicted by AlphaFold3. The structure of HYS monomer was extracted from the X-ray crystal structure (PDB ID: 5H83) and subsequently used to replace the VinBLAST moiety in VinBLAST-GS heterodimer, thereby generating HYS-GS heterodimer. Subsequently, the substrate 4,21-dehydrogeissoschizine was docked into the monomer- or dimer-NADPH complex by AutoDock Vina (*41*). The same procedure was applied to all other strictosidine aglycone substrates. Referring to the reaction catalyzed by GS, the closest complex structure was chosen as the initial configuration for the subsequent MD simulations.

The structure of the SGD monomer was extracted from the X-ray crystal structure (PDB ID: 2JF7), and the structure of the VinBLAST monomer with zinc was predicted using AlphaFold3. These two monomers were then combined according to each of the defined potential interfaces using PyMOL, resulting in nine possible SGD-VinBLAST interaction systems. For the seven SGD-VinBLAST systems exhibiting strong PPI interactions, VinBLAST was aligned with GS and subsequently replaced by GS to generate SGD-GS systems.

All simulations were performed with GROMACS-2022.02 (*42*) using an amber99sb-ildn force field (*43*). The force field parameters of all substrates and NADPH were constructed by Gaussian and ACPYPE. To keep the experimental conditions consistent, the temperature of the simulation system was constant at 310 K, and the pressure was stabilized at 1.0 bar, using the Berendsen thermostat and Parrinello-Rahman pressure coupling. Periodic boundary conditions were applied in all simulations. The cut-off of non-bonded van der Waals interaction and long-range electrostatic interaction were both set at a distance of 1.2 nm. The linear constraint solver algorithm (*44*) was used in simulations to constrain bond lengths.

The ternary complex of dimer-NADPH-substrate was first placed in a 12×12×10 $nm^3$ box, then was solvated by water molecules. 0.1 M NaCl was added into the system to maintain electrical neutrality. The system was first energy-minimized and equilibrated for 0.1 ns, then followed by MD simulations in the NVT ensemble (1 ns) and the NPT ensemble (2 ns). A further 100 ns MD production run was then performed to get a stable conformation for the dimer-NADPH-substrate ternary complex. The monomer-NADPH-substrate ternary complex was first placed in a 10×6×6 $nm^3$ box, then underwent the same solvation and neutralization process as mentioned above, followed by 0.1 ns energy minimization, as well as 1 ns NVT and 2 ns NPT equilibration. Finally, a long-term molecular dynamics simulation of 1,000 ns was performed for monomer-NADPH-substrate ternary complex. All input files for MD simulations are provided in Data S3.

LC-MS/MS detection

LC-MS/MS analysis of samples from *de novo* MIA-producing yeast strains was performed using a SHIMADZU Triple Quadrupole LC-MS/MS 8045 system equipped with a HyPURITY C18 column (3.0 µm, 150 × 4.6 mm) to achieve optimal separation of complex metabolite mixtures. The mobile phase consisted of 0.1% formic acid in water (solvent A) and 0.1% formic acid in methanol (solvent B), with a 50-min gradient elution at a flow rate of 0.3 mL $min^{-1}$ programmed as follows: initial 10% B, linear increase to 90% B for 0-25 min, maintained at 90% B until 35 min, then returned to 10% B by 50 min. The analysis was conducted with the column oven set at 30°C, using a atomizing gas flow of 3.0 L $min^{-1}$, collision-induced dissociation (CID) gas pressure of 230 kPa, desolvation line (DL) temperature of 250°C, and heat block temperature of 400°C. Target MIAs were quantified in multiple-reaction monitoring (MRM) mode with specific transitions: catharanthine with a collision energy of 20 eV and an *m/z* transition from 337.10 to 144.15; vindoline with a collision energy of 28 eV and an *m/z* transition from 457.05 to 188.05; geissoschizine methyl ether with a collision energy of 32 eV and an *m/z* transition from 367.20 to 144.10. All data were processed using SHIMADZU LabSolutions software for metabolite quantification and analysis.

The in vitro assay samples were analyzed using a Ultivo Triple Quadrupole LC-MS/MS system from Agilent (Santa Clara, CA, USA), equipped with an Avantor® ACE® UltraCore C18 2.5 Super C18 column (50×3 mm, particle size 2.5 µm) as well as a photodiode array detector and a mass spectrometer. For alkaloid analysis, the following solvent systems were used: Solvent A, methanol:acetonitrile:ammonium acetate (1 M):water at 29:71:2:398; solvent B, methanol: acetonitrile:ammonium acetate (1 M):water at 130:320:0.25:49.7. The following linear elution gradient was used: 0-5.0 min 80% A, 20% B; 5.0-5.8 min 1% A, 99% B; 5.8-8.0 min 80% A, 20% B; the flow during the analysis was constant and 0.6 mL $min^{-1}$. The photodiode array detector range was 200 to 500 nm. The mass spectrometer was operated with the gas temperature at 300°C and gas flow of 10 L $min^{-1}$. Capillary voltage was 4 kV from *m/z* 100 to *m/z* 1,000 with scan time 500 ms, and the fragmentor performed at 135 V with positive polarity.

Bimolecular fluorescence complementation (BiFC)

To perform the yeast BiFC assay, mVenus is divided into two parts, the *N*-terminal fragment ($mVenus^N$) and the *C*-terminal fragment ($mVenus^C$). Neither of these two parts can generate yellow fluorescence when they exist separately. The proteins to be tested were fused to the $mVenus^N$ and $mVenus^C$, respectively. Considering the presence of a *C*-terminal NLS (nucleus localization signal) in SGD, the mVenus fragments were fused to the *N*-terminus of SGD. For both VinBLAST and GS, the mVenus fragments were fused to their *C*-termini. The target protein-coding sequences were amplified and then fused to either the *N*-terminal ($mVenus^N$) or *C*-terminal ($mVenus^C$) fragments of the fluorescent protein mVenus on pESC series vectors. The recombinant plasmids were co-transformed into the wild-type *S. cerevisiae* strain CEN.PK2-1C, and positive clones were selected on SED agar plates lacking appropriate nutrients. The transformed yeast cells were then cultured in liquid medium at 30°C with shaking at 250 rpm for 24 h. The yeast cells were harvested by centrifugation at 4,000 rpm for 5 min, washed twice with sterile water to remove residual glucose, and subsequently transferred into fresh SE medium containing 20 g $L^{-1}$ galactose to induce plasmid expression for 12 h with shaking at 250 rpm. For fluorescence observation, cells were harvested by centrifugation, washed twice with phosphate-buffered saline (PBS), and resuspended in fresh PBS before visualization. Appropriate negative controls, including empty vector (EV) transformations and non-interacting protein pairs, were included in each experiment to validate specific protein-protein interactions. The images were acquired using the confocal laser scanning microscope Olympus FV3000 2.3.1.163. The image size was 1024×1024 pixels; the objective lens was PLAPON 60XO with Zoom×10.0; the sampling speed was 2.0 µs $pixel^{-1}$. The fluorescence of mCherry was observed with detection wavelength of 570-620 nm; the fluorescence of EGFP was observed with detection wavelength of 500-540 nm; and the fluorescence of mVenus was observed with detection wavelength of 530-630 nm.

To perform BiFC in *Nicotiana benthamiana*, single colonies of *A. tumefaciens* strain LBA4404 carrying BiFC vectors were grown overnight at 28°C in LB medium with shaking at 200 rpm. Cells were harvested by centrifugation and resuspended in infiltration buffer (10 mM MES, pH 5.6; 10 mM $MgCl_2$; 0.2 mM acetosyringone) to an $OD_{600}$ of 0.8. The resuspended cultures were incubated at 28°C for 3 h to induce virulence. Equal volumes of the desired BiFC strains were mixed and infiltrated into the abaxial surface of tobacco leaves using a needleless syringe. Following infiltration, plants were maintained in a growth chamber set to 26°C under a 16 h light/8 h dark photoperiod for 3 days to allow protein expression. A 5 by 5 mm section of infiltrated leaf tissue was excised and mounted on a microscope slide for imaging. Fluorescence was detected using a Leica SP8 Confocal Scanning Laser Microscope with a hybrid detector set to an emission bandwidth of 510-540 nm (for mVenus) and a Smart Gain setting of 40.0%. Transmitted light images were captured using a photomultiplier tube (PMT) with a gain of 150.0 V. Sequential scanning between the hybrid detector and PMT trans detector was used to acquire images at a spatial resolution of either 512 by 512 or 1024 by 1024 pixels, with a magnification of 250×. Images were processed using Fiji (version 2.16.0).

Yeast two-hybrid (Y2H) assays

The protein-protein interactions were investigated using the growth-based yeast two-hybrid (Y2H) system. The coding sequences of target proteins were fused to either the transcription activation domain (AD) or DNA-binding domain (BD) of the GAL4 system at their *N*-termini in pGADT7 or pGBKT7 plasmids. The recombinant plasmids were co-transformed into the *S. cerevisiae* strain Y2H-GOLD. To evaluate the strength of protein-protein interactions and minimize the background noise from leaky *HIS3* reporter gene expression, the competitive inhibitor 3-amino-1,2,4-triazole (3-AT) were added at different concentration (0 mM, 5 mM, and 20 mM) into the agar plates. The yeast strains were diluted at different ratios in 10-fold increments, and then 10 µL culture solutions were spotted onto SED agar plates with histidine, tryptophan and leucine deficiencies and incubated at 30°C for 96 h to allow colony formation.

Synteny analysis

To ensure consistency across genomic datasets, coding sequence (genome.cds) files were extracted from genome annotation (genome.gff) and genome sequence (genome.fasta) files using GffRead, which converts genomic features into standardized formats (*45*). Synteny analysis was performed using Python-MCscan (https://github.com/tanghaibao/jcvi/wiki/MCscan-(Python-version)), with the genome.cds and genome.gff files as inputs (*46*). Gene pairs were compared using LAST (https://gitlab.com/mcfrith/last) to filter out tandem duplications and low-confidence alignments, retaining high-quality anchors for clustering into syntenic blocks (*47*). Microsynteny visualization was conducted following the Python-MCscan workflow to map regions of interest. Percentage protein identity of key enzymes in the syntenic regions was provided in table S9. Details on genome information used in this study were provided in table S10.

Co-expression analysis

Co-expression analysis was performed by R package WGCNA (version 1.73, Weighted Gene Co-Expression Network Analysis, https://github.com/Uauy-Lab/WheatHomoeologExpression) (*48*). As most MIA pathway genes were expressed in each *C. roseus* sample, FPKM>20 was set as a threshold to select candidate genes, followed by WGCNA analysis. The module containing the MIA biosynthetic pathway genes (STR, GS, GO, etc.) was further analyzed. STR, GS, and GO were selected as “baits” with correlation threshold ($r > 0.9$) for constructing the list of co-expressed genes. Cytoscape (v.3.9.1) (*49*) was used to visualize the co-expression network.

Phylogenetic analysis

The protein sequences used to construct the phylogenetic tree were retrieved from S. de Vries et al. (*25*), with additional CAD sequences listed in the table S3. Following the approach described in that study, highly variable *N*-terminal regions were trimmed to enable more robust phylogenetic comparisons. The evolutionary history was inferred using the Maximum Likelihood method with the JTT matrix-based model. The tree with the highest log-likelihood value is presented. Branch support values indicate the percentage of replicate trees (from 100 bootstrap replicates) in which the associated taxa clustered together. Initial trees for the heuristic search were generated automatically using the Neighbor-Joining and BioNJ algorithms based on pairwise distances estimated under the JTT model. The topology with the highest log-likelihood score was then selected. The final tree is drawn to scale, with branch lengths representing the number of substitutions per site, as indicated by the scale bar. CAD clade I and IV, based on S. de Vries et al. (*25*) designation were collapsed for simplicity.

Surface plasmon resonance (SPR) assays

The binding affinities of protein-protein interactions were measured on a Biacore 8K instrument (Cytiva) with CM5 sensor chips (Cytiva) at 25°C. The immobilized proteins SGD, VinBLAST, and GS were diluted with 10 mM NaAc (pH 4.5), respectively. The method was performed in multi-cycle mode, with sensor chip surface regeneration between each cycle. The analytes were serially diluted to gradient concentrations with 1x PBS-P+ buffer (20 mM phosphate buffer, 2.7 mM KCl, 137 mM NaCl, and 0.05% (v/v) Tween 20, pH 7.4) and injected at a flow rate of 30 µL $min^{-1}$ over the ligands for 90 sec, followed by dissociation for 90 sec. After dissociation, 10 mM Glycine-HCl (pH 2.5) was injected for 30 sec to remove any non-covalently bound proteins from the chip surface. The binding affinities were analyzed with the software Biacore Insight Evaluation 6.0.7.1750 using the 1:1 binding mode.

### Supplementary Text

Co-silencing of *CAD1* and *CAD2* in *C. roseus* leaf

When we silenced *CAD2* (90% decrease), *CAD1* transcript was mildly affected (38% decrease). However, when we silenced *CAD1*, its transcript was decreased to 38.4%, while *CAD2* transcript levels were also decreased to 43.3%, indicating a degree of co-silencing between the two highly homologous genes (82% identity at nucleic acid level, fig. S2).

The impact of VIGS-CAD1 on MIA production in *C. roseus* was weaker than that observed with VIGS-CAD2. Silencing *CAD2* led to a striking decrease in catharanthine and vindoline levels, by 93.5% and 83.7%, respectively, and a substantial increase in HYS-derived MIAs, ajmalicine and serpentine, by 6.1- and 3.7-fold, respectively (Fig. 2B). Silencing *CAD1* led to a relatively modest decrease in catharanthine and vindoline levels, by 75.0% and 45.7%, respectively, and an increase in ajmalicine and serpentine by 3.1- and 8.5-fold, respectively (fig. S2).

VinBLAST boosts GS catalytic activity

To further investigate the role of VinBLAST in the GS-SGD relationship, we tested GS in vitro activity in the presence of VinBLAST. In the coupled SGD-GS-C17OMT reaction, adding recombinant VinBLAST enhanced GME yield from strictosidine, with optimal enhancement observed at GS:VinBLAST stoichiometries between 2:1 and 1:4 (fig. S26). Although VinBLAST functions as a scaffold linking SGD and GS, the observed enhancement of GS activity by VinBLAST is independent of SGD. When strictosidine aglycone replaced strictosidine as the substrate, co-incubation of recombinant GS and VinBLAST at a 1:1 ratio, with or without SGD, resulted in comparable increase in geissoschizine yield (13.4- and 14.5-fold, respectively). The VinBLAST$^{K359G}$ mutant also exhibited comparable GS-enhancing activity (fig. S27).

In vitro kinetic analysis revealed further details in GS catalysis. We observed no difference in the binding affinities for strictosidine aglycone between the GS homodimer, the VinBLAST-GS complex, and VinBLAST$^{CM}$-GS complex, with comparable $K_m$ values of 0.495 μM, 0.512 μM, and 0.617 μM, respectively (fig. S28 and table S2). In contrast, VinBLAST, and VinBLAST$^{CM}$, markedly enhanced the GS catalytic rate, increasing the $V_{max}$ of GS by 24.3-, and 30.3-fold, respectively. These results strongly suggest altered GS active site architecture in the VinBLAST-GS complex.

Protein-protein interactions among CAD1, GS, and SGD

BiFC assays showed that CAD1 interacted with GS in the cytosol and with SGD in the nucleus, and we also observed CAD1 homodimerization in the cytosol. However, in contrast to CAD2, the presence of CAD1 did not enable detectable interaction between SGD and GS, indicating that CAD1 was not as effective as CAD2 to serve as a scaffold to bridge SGD and GS (fig. S42).

Y2H results showed that CAD1 interacted with both GS and itself (fig. S43). Regarding the interaction with SGD, Y2H revealed a very weak interaction between CAD1 and SGD, which is consistent with the generally weak Y2H signals observed for SGD with other proteins in our study (fig. S14). We interpret these findings to suggest that Y2H lacks sufficient sensitivity to reliably detect the very weak protein-protein interactions involving SGD.

Effect of CAD1 in enhancing MIA production

To evaluate the role of CAD1 in enhancing MIA production, we tested its performance in yeast strains engineered to produce GME, catharanthine, and vindoline. In all yeast strains,

CAD1 was able to improve the MIA titers, yet its improvements were only 27.4%, 7.85%, and 21.3%, respectively, of those achieved by CAD2 (fig. S44). These results indicate that CAD1 has a limited capacity to enhance MIA production when compared with CAD2.

**Fig. S1. The gene expression levels of *HYS*, *GS2*, *GS*, *CAD2* (*VinBLAST*), and *CAD1* in *C. roseus* leaf.** RNAseq data was obtained from BioProject PRJNA888475 from NCBI (*C. roseus* cv. Pacifica XP Burgundy) and analyzed using CLC Genomic Bench version 20.0.4. The vertical axis represents the Transcript per Kilobase per Million mapped reads (TPM) values. The results represent the mean ± s.d. from three biological replicates.

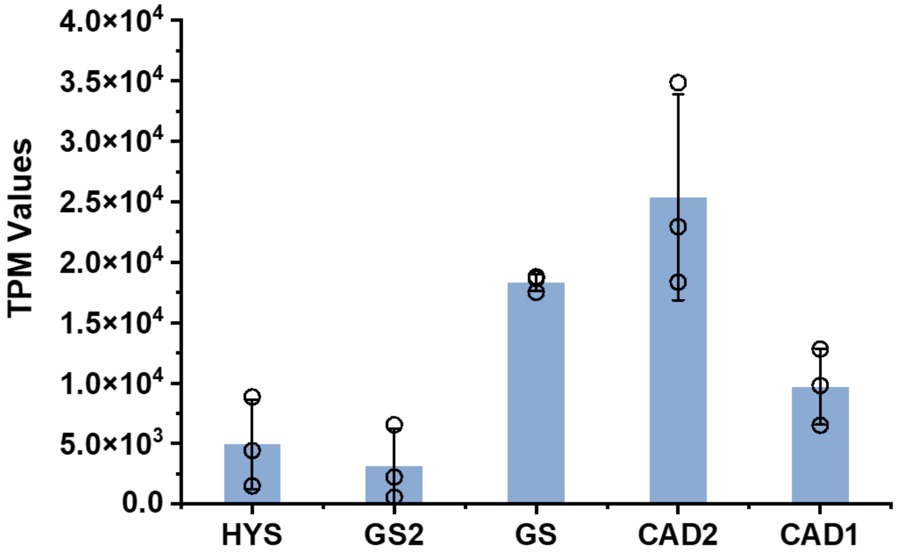

**Fig. S2. Virus-induced gene silencing of *CrCAD1* and *CrCAD2* in *C. roseus* leaf.** In VIGS-CAD1 plants, the transcript level of *CAD1* was decreased to 38.4%, while *CAD2* transcript levels were also decreased to 43.3%, indicating a degree of co-silencing between the two highly homologous genes. In contrast, when *CAD2* was targeted by VIGS, *CAD1* transcript level was less affected. The impact of VIGS-CAD1 on MIA production in *C. roseus* was weaker than that observed with VIGS-CAD2. **(A)** MIA contents following VIGS of *CAD2* (*VinBLAST*) in *C. roseus* leaves. The results represent the mean ± s.d. of 7 or 8 biological replicates. **(B)** MIA contents following VIGS of *CAD1* in *C. roseus* leaves. The results represent the mean ± s.d. of 6 or 7 biological replicates. **(C)** The relative expression levels of *GS*, *CAD2* (*VinBLAST*), and *CAD1* in leaves of VIGS-CAD2 plants compared with the empty vector (EV) control. The results represent the mean ± s.d. of 7 or 8 biological replicates, each with 3 technical replicates. **(D)** The relative expression levels of *GS*, *CAD2* (*VinBLAST*), and *CAD1* in leaves of VIGS-CAD1 plants compared with the empty vector (EV) control. The results represent the mean ± s.d. of 6 or 7 biological replicates, each with 3 technical replicates.

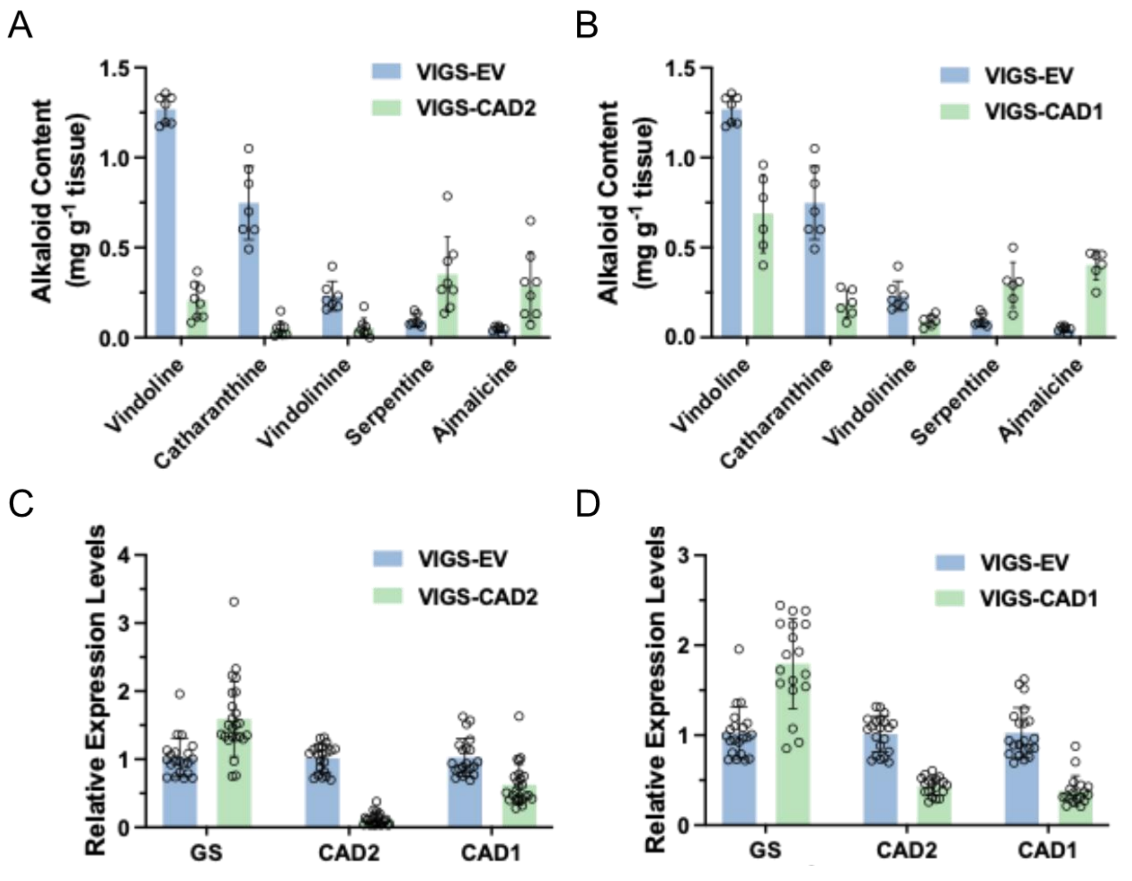

**Fig. S3. Virus-induced gene silencing of *CrCAD3, CrCAD4,* and *CrCAD5* did not alter alkaloid contents in *C. roseus* leaf.** The alkaloid contents (mg $g^{-1}$ fresh leaf) for catharanthine (purple), vindoline (blue), and ajmalicine (cyan) were measured in VIGS-empty vector (EV), VIGS-CAD3, VIGS-CAD4, and VIGS-CAD5 plants. The results represent the mean ± s.d. from seven independent biological replicates.

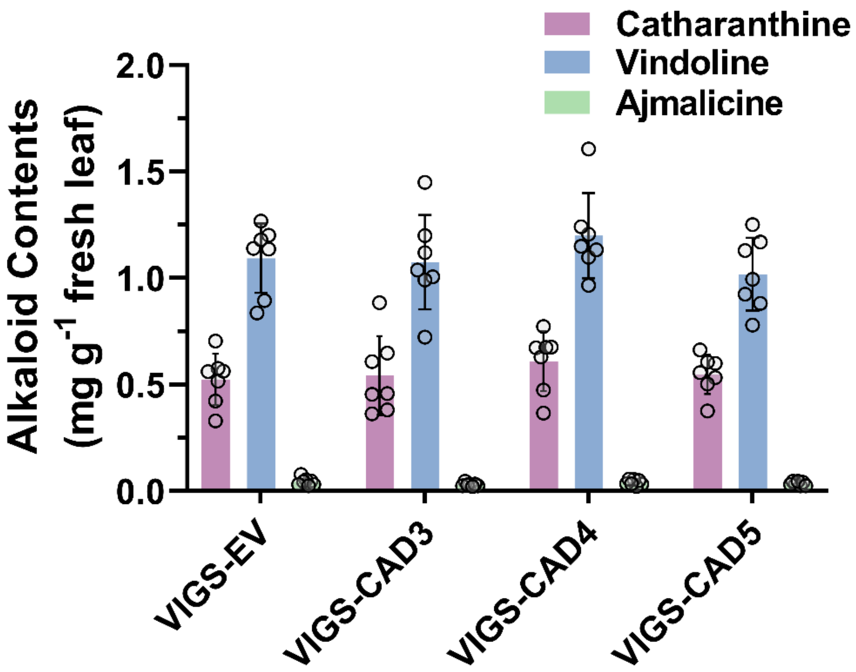

**Fig. S4. The MS/MS spectra confirmed the *de novo* production of geissoschizine methyl ether (GME), catharanthine, and vindoline in *Saccharomyces cerevisiae*.** The MS/MS product ion fingerprints of the fermentation sample of strain GM01 (**A**), CA01 (**B**), VI01 (**C**) were identical with authentic standards.

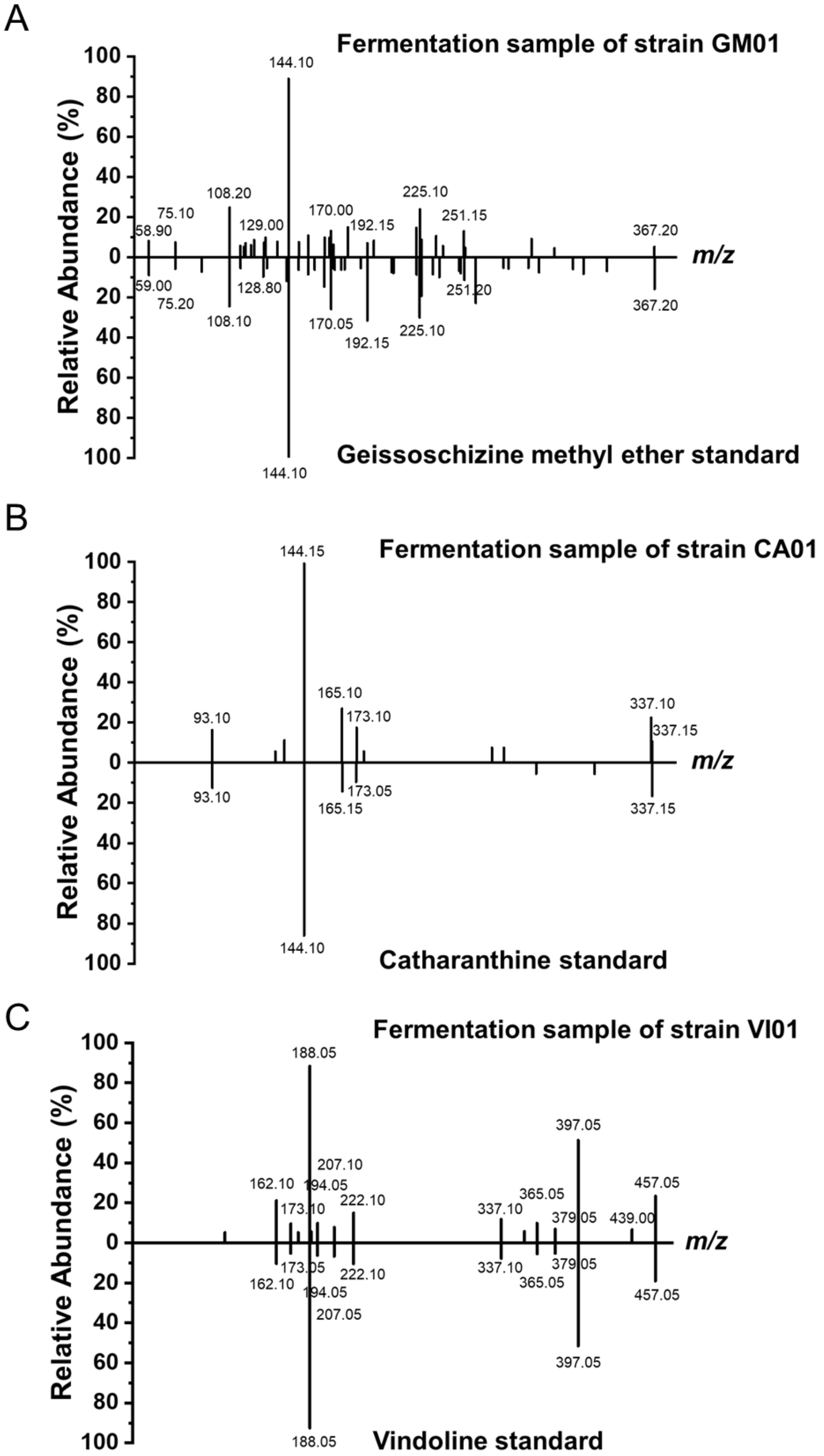

**Fig. S5. *VinBLAST* expression enhanced catharanthine production in *Pichia pastoris*.** The *VinBLAST* expression cassette (*TEF1p-VinBLAST-0547t*) was integrated into the chromosome (Int63) of CAN19 (*5*), a previously engineered *P. pastoris* strain for *de novo* biosynthesis of catharanthine at a titer ~0.155 mg $L^{-1}$. Following the same fermentation conditions as previously described (*5*), the introduction of *VinBLAST* increased the production of catharanthine by 10.2-fold. This result indicated the function of VinBLAST in different chassis cells and the potential for general applications. The results represent the mean ± s.d. of three biological replicates.

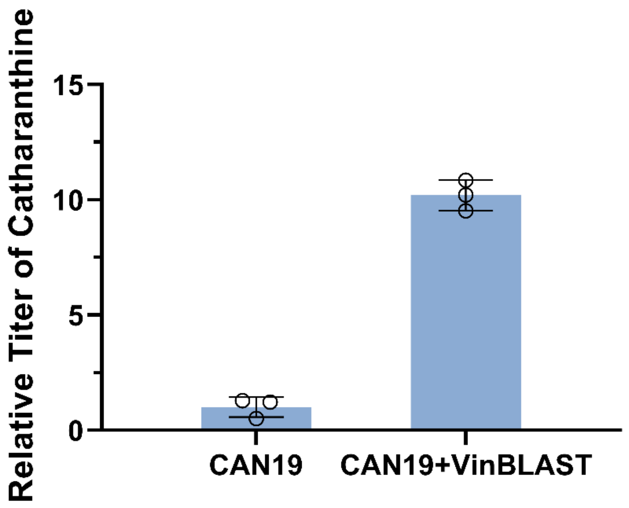

**Fig. S6. A single genomic copy of *VinBLAST* was sufficient to relieve biosynthesis bottleneck in *de novo* geissoschizine methyl ether (GME) production in engineered *S. cerevisiae* strains**. Each strain contains 0-2 genomic copies of GS and/or VinBLAST. Introducing a second copy of both GS and VinBLAST resulted in only a modest 36% increase in GME titer compared to strains carrying a single copy of each enzyme, indicating that a single copy of VinBLAST is sufficient to overcome the major bottleneck in GME biosynthesis. The parental strain was AJM7-ΔHYS (*4*) and the corresponding genomic modifications for each strain were specified underneath. The strain genotypes were listed in table S1. The results represent the mean ± s.d. of three biological replicates.

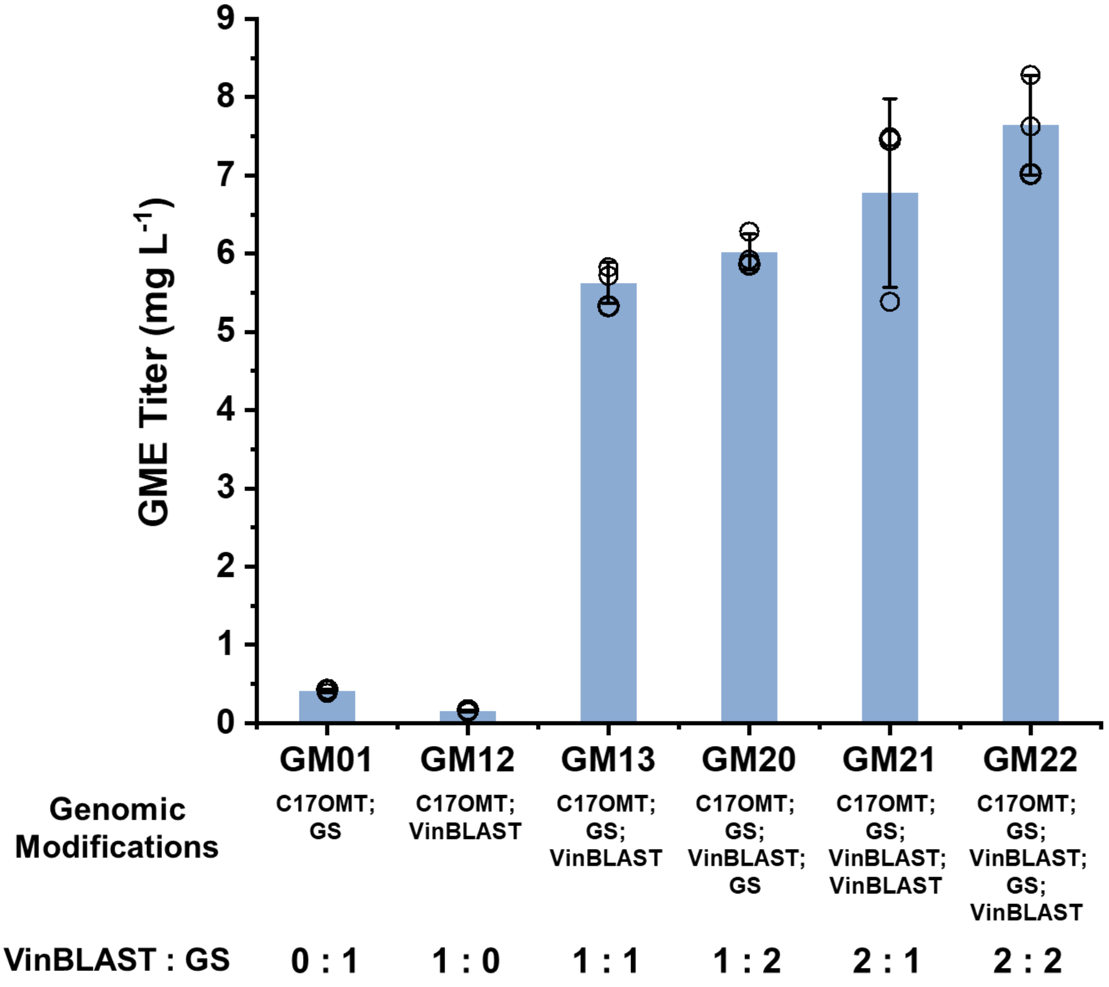

**Fig. S7. Subcellular localization of VinBLAST, GS, and SGD in yeast.** Enhanced green fluorescent protein (EGFP) was fused to either the *N*- or *C*-terminus of VinBLAST (**A** and **B**), GS (**C** and **D**), and SGD (**E** and **F**) to assess localization. Nab2 tagged with mCherry at the *C*-terminus functioned as a nuclear marker. All proteins were expressed in the wild-type CEN.PK2-1C yeast strains. The images were acquired using a confocal laser scanning microscope (Olympus FV3000) at 600× magnification, with a 5 μm scale bar displayed in the upper left corner.

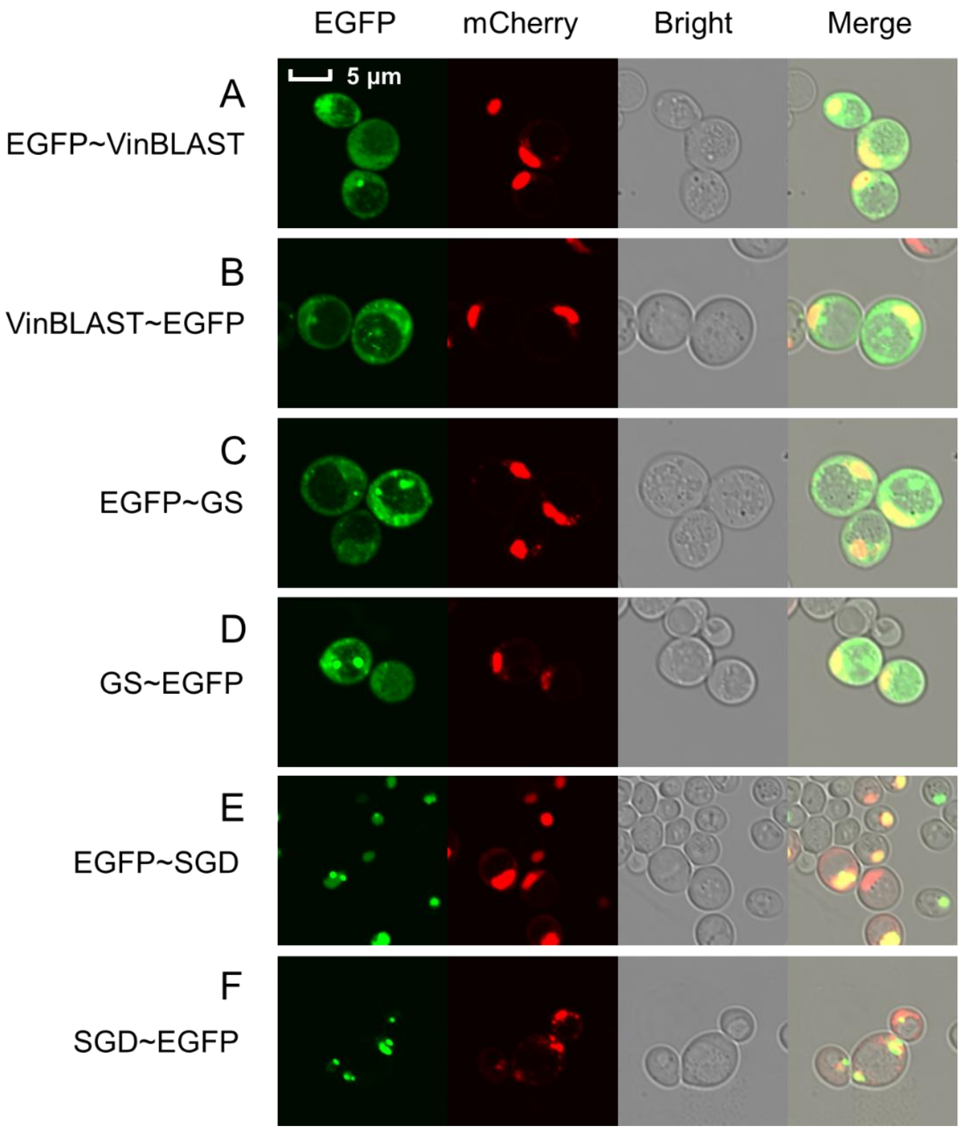

**Fig. S8. The bimolecular fluorescence complementation (BiFC) experiments in yeast demonstrated protein-protein interactions among VinBLAST, GS, and SGD.** The figures showed the interactions of SGD with itself **(A)**, with GS **(B)**, and with VinBLAST **(C)**; the interactions of GS with SGD **(D)**, with itself **(E)**, and with VinBLAST **(F)**; and the interactions of VinBLAST with SGD **(G)**, with GS **(H)**, and with itself **(I)**. The *N*-terminal fragment (mVenus$^{N}$) and the *C*-terminal fragment (mVenus$^{C}$) of mVenus were individually fused to proteins of interest. Yellow fluorescence is generated when the two mVenus fragments are brought into proximity by interaction of the fused proteins. Considering that SGD has a nuclear localization signal (NLS) at the *C*-terminus, the mVenus fragments were fused to the *N*-terminus of SGD. For both VinBLAST and GS, the mVenus fragments were fused to their *C*-termini. Wild-type VinBLAST interacted with GS in the cytosol and with SGD in the nucleus. Nab2 tagged with mCherry at the *C*-terminus functioned as a nuclear marker. All proteins were expressed in the wild-type CEN.PK2-1C yeast strains. The images were acquired using a confocal laser scanning microscope (Olympus FV3000) at 600× magnification, with a 5 µm scale bar displayed in the upper left corner.

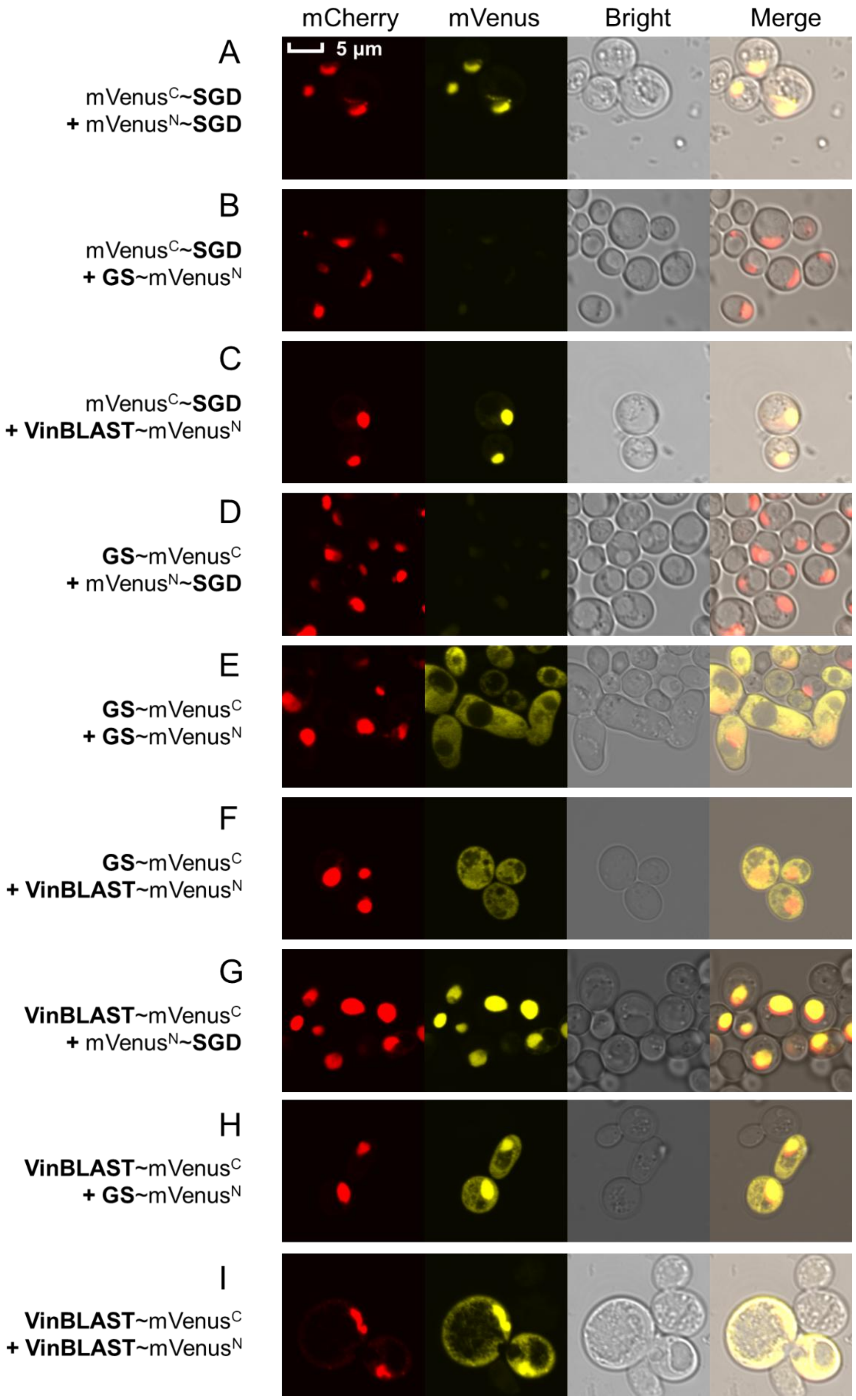
mCherry
mVenus
Bright
Merge
5 µm
A
mVenusC~SGD
+ mVenusN~SGD
B
mVenusC~SGD
+ GS~mVenusN
C
mVenusC~SGD
+ VinBLAST~mVenusN
D
GS~mVenusC
+ mVenusN~SGD
E
GS~mVenusC
+ GS~mVenusN
F
GS~mVenusC
+ VinBLAST~mVenusN
G
VinBLAST~mVenusC
+ mVenusN~SGD
H
VinBLAST~mVenusC
+ GS~mVenusN
I
VinBLAST~mVenusC
+ VinBLAST~mVenusN

**Fig. S9. The BiFC experiments in yeast demonstrated protein-protein interactions among VinBLAST, VinBLAST$^{CM}$, VinBLAST$^{IM}$, GS, and SGD.** The *N*-terminal fragment (mVenus$^{N}$) and the *C*-terminal fragment (mVenus$^{C}$) of mVenus were individually fused to proteins of interest. Yellow fluorescence is generated when the two mVenus fragments are brought into proximity by interaction of the fused proteins. Considering that SGD has a nuclear localization signal (NLS) at the *C*-terminus, the mVenus fragments were fused to the *N*-terminus of SGD. For both VinBLASTs and GS, the mVenus fragments were fused to their *C*-termini. Wild-type VinBLAST interacted with GS in the cytosol and with SGD in the nucleus. Notably, when non-tagged VinBLAST was co-expressed, it enabled GS to interact with SGD in the nucleus. The VinBLAST catalytic mutant (VinBLAST$^{CM}$; C51A and H56A mutations disrupting NADPH binding) retained wild-type activity. In contrast, the interaction mutant (VinBLAST$^{IM}$; M298E and V299E mutations weakening dimer formation) failed to interact with either SGD or GS in BiFC assays, indicating the importance of dimerization in mediating protein-protein interactions. Nab2 tagged with mCherry at the *C*-terminus functioned as a nuclear marker. All proteins were expressed in the wild-type CEN.PK2-1C yeast strains. The images were acquired using a confocal laser scanning microscope (Olympus FV3000) at 600× magnification, with a 5 μm scale bar displayed in the upper left corner. The yellow box indicates BiFC experiments involving the three proteins related to wild-type VinBLAST **(A-C)**, the purple box indicates BiFC experiments involving the three proteins related to VinBLAST$^{CM}$ **(D-F)**, and the blue box indicates BiFC experiments involving the three proteins related to VinBLAST$^{IM}$ **(G-I)**.

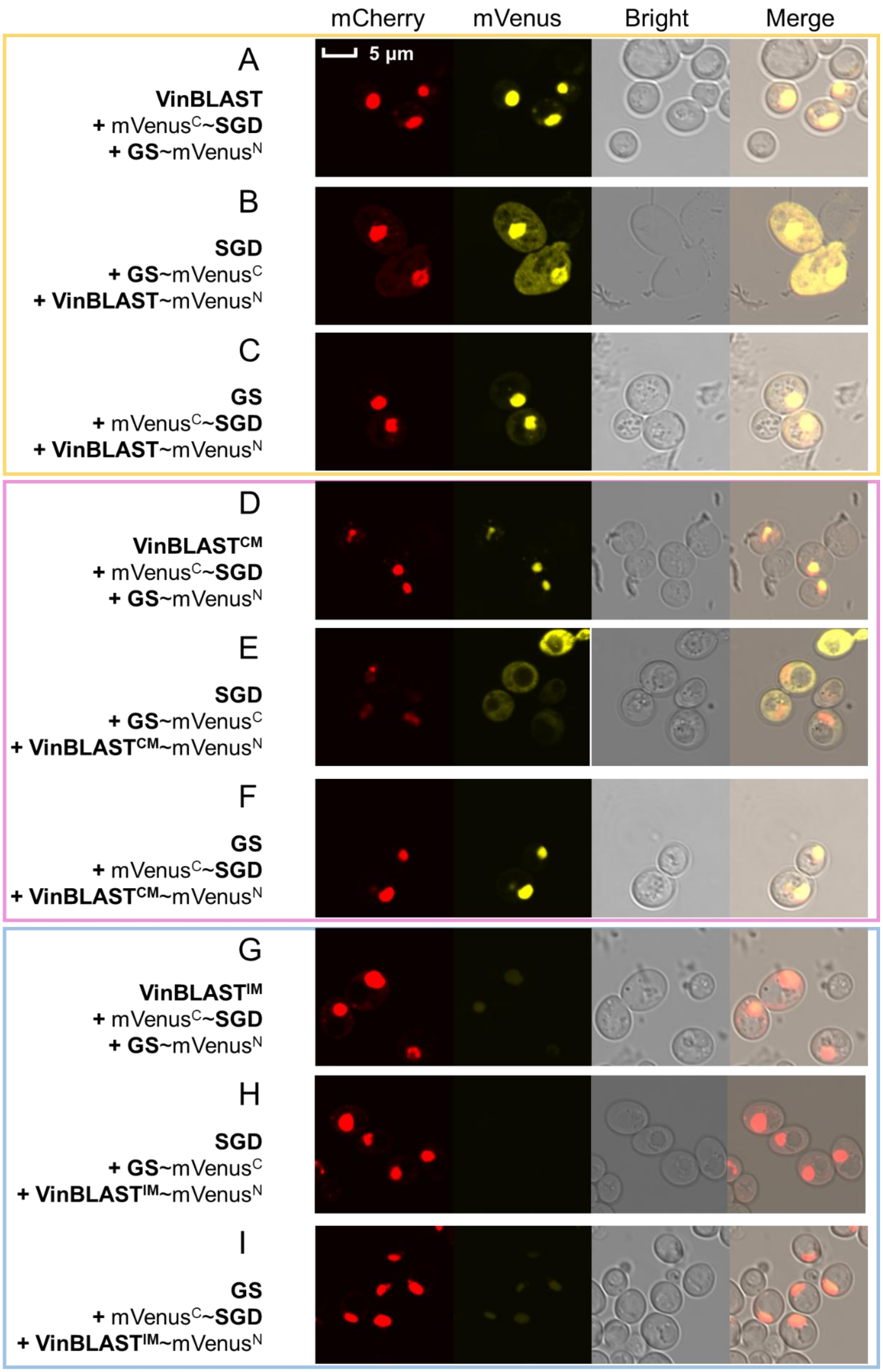
mCherry
mVenus
Bright
Merge
5 μm
A
VinBLAST
+ mVenusC~SGD
+ GS~mVenusN
B
SGD
+ GS~mVenusC
+ VinBLAST~mVenusN
C
GS
+ mVenusC~SGD
+ VinBLAST~mVenusN
D
VinBLASTCM
+ mVenusC~SGD
+ GS~mVenusN
E
SGD
+ GS~mVenusC
+ VinBLASTCM~mVenusN
F
GS
+ mVenusC~SGD
+ VinBLASTCM~mVenusN
G
VinBLASTIM
+ mVenusC~SGD
+ GS~mVenusN
H
SGD
+ GS~mVenusC
+ VinBLASTIM~mVenusN
I
GS
+ mVenusC~SGD
+ VinBLASTIM~mVenusN

**Fig. S10. The Surface plasmon resonance (SPR) assays were performed to investigate the interactions between VinBLAST and GS.** Panels show SPR assays of 16OMT **(A)**, VinBLAST **(B)** binding to VinBLAST, and GS **(C)**, VinBLAST **(D)** binding to GS. VinBLAST or GS were immobilized on the sensor chip, and the analyte proteins were injected over the immobilized ligand surface at gradient concentrations (0.06-2 μM). At the concentrations tested (0.06-2 μM), no binding between 16OMT and VinBLAST **(A)** was detected, as a negative control. Interactions of VinBLAST homodimer **(B)**, GS homodimer **(C)**, and VinBLAST-GS heterodimer **(D)** were detected by SPR. Experiments were repeated twice with similar results.

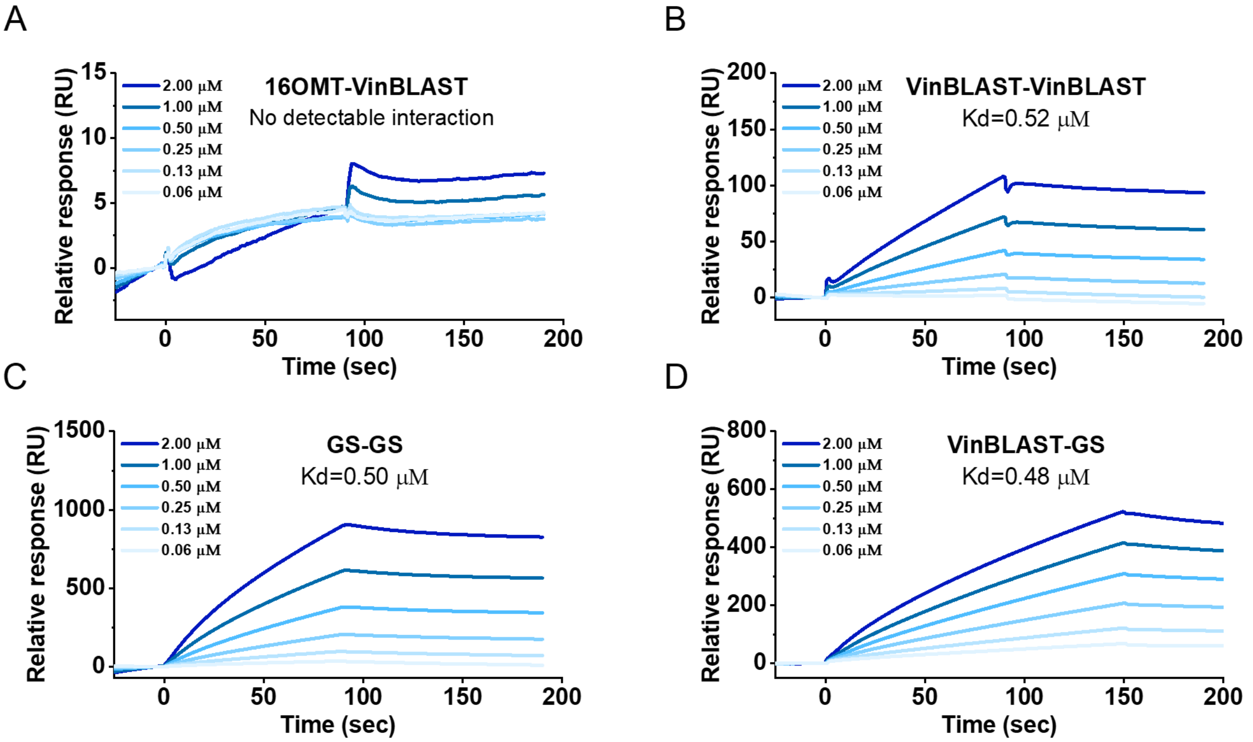

**Fig. S11. VinBLAST retained bona fide cinnamyl alcohol dehydrogenase (CAD) activity while showing no strictosidine aglycone reducing activity. (A)** Purified recombinant VinBLAST reduced both cinnamyl and coniferyl aldehydes to their corresponding alcohols. In contrast, both VinBLAST catalytic mutant (VinBLAST$^{CM}$; C51A and H56A mutations disrupting NADPH binding) and the interaction mutant (VinBLAST$^{IM}$; M298E and V299E mutations weakening dimer formation) completely lost this activity. The results indicated that both NADPH binding and dimerization were essential for CAD enzyme activity. The LC-MS/MS chromatograms show UV absorbance at 280 nm. **(B)** Purified recombinant VinBLAST homologues from *Vitis vinifera*, *Strychnos spinosa*, *Gelsemium sempervirens*, and *N. benthamiana* demonstrated CAD activity, reducing coniferyl aldehyde, whereas CrGS did not. The LC-MS/MS chromatograms show UV absorbance at 280 nm. **(C)** GS reduced strictosidine aglycone generated through SGD-catalyzed hydrolysis of strictosidine. In contrast, CrVinBLAST did not exhibit this reductase activity. **(D)** Structure of VinBLAST-GS heterodimer predicted by AlphaFold3, guiding the design of VinBLAST$^{CM}$, VinBLAST$^{IM}$, and GS$^{IM}$ mutants. Blue: GS; cyan: VinBLAST; the upper left dashed rectangle: structural details of interacting β-sheet pair at the VinBLAST-GS interface; the bottom left dashed rectangle: structural details of interacting β-sheet pair at the VinBLAST$^{IM}$-GS$^{IM}$ interface; the right dashed circle: NADPH binding site in VinBLAST. VinBLAST$^{CM}$: the catalytic mutant VinBLAST$^{C51A,H56A}$; VinBLAST$^{IM}$: the interaction mutant VinBLAST$^{M298E,V299E}$; GS$^{IM}$: the interaction mutant GS$^{I301E}$.

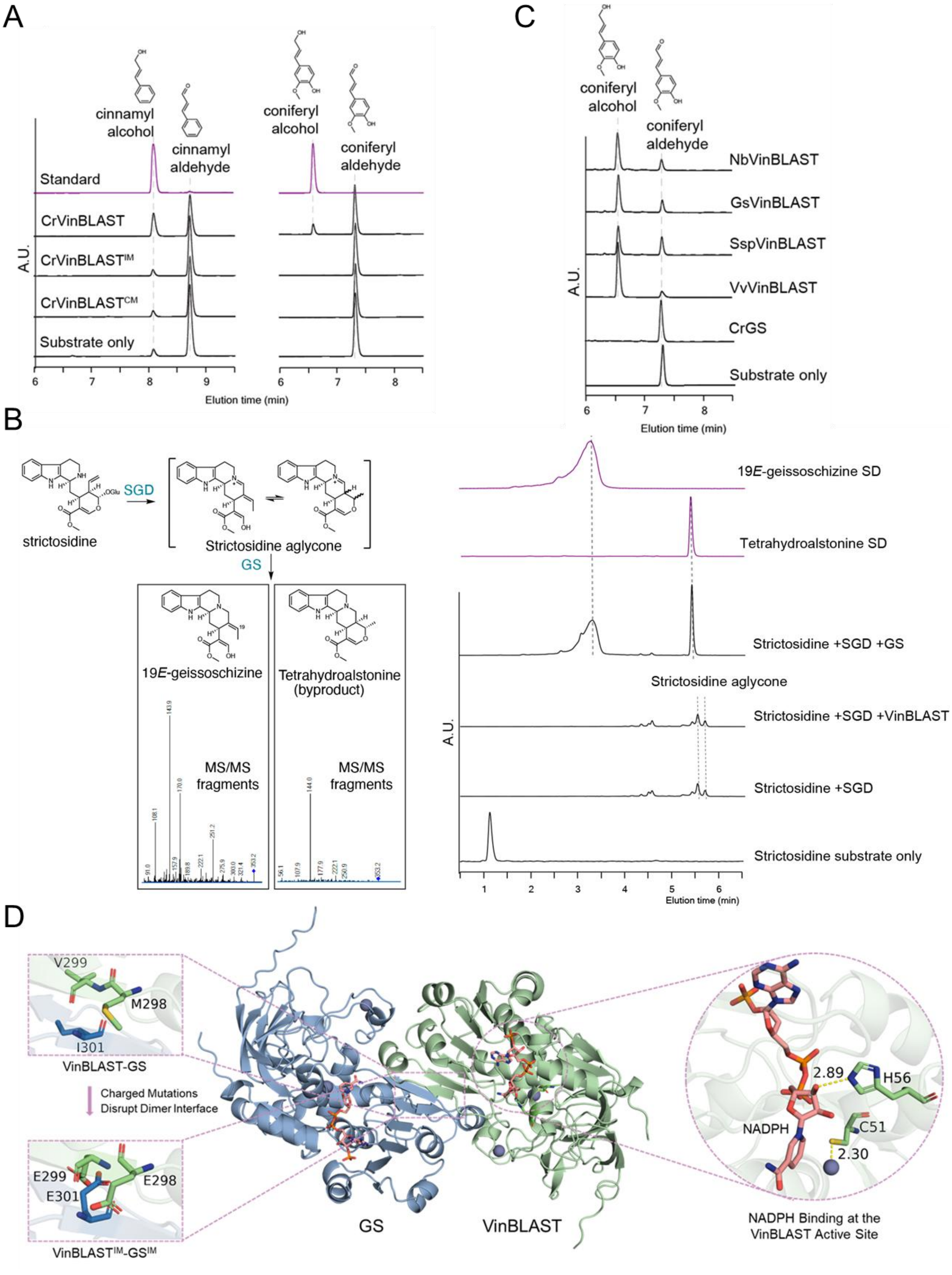

A
cinnamyl alcohol
cinnamyl aldehyde
coniferyl alcohol
coniferyl aldehyde
Standard
CrVinBLAST
CrVinBLAST^IM
CrVinBLAST^CM
Substrate only
A.U.
Elution time (min)
B
strictosidine
SGD
Strictosidine aglycone
GS
19E-geissoschizine
Tetrahydroalstonine (byproduct)
MS/MS fragments
MS/MS fragments
19E-geissoschizine SD
Tetrahydroalstonine SD
Strictosidine +SGD +GS
Strictosidine aglycone
Strictosidine +SGD +VinBLAST
Strictosidine +SGD
Strictosidine substrate only
A.U.
Elution time (min)
C
coniferyl alcohol
coniferyl aldehyde
NbVinBLAST
GsVinBLAST
SspVinBLAST
VvVinBLAST
CrGS
Substrate only
A.U.
Elution time (min)
D
V299
M298
I301
VinBLAST-GS
Charged Mutations Disrupt Dimer Interface
E299
E298
E301
VinBLAST^IM-GS^IM
GS
VinBLAST
2.89
H56
NADPH
C51
2.30
NADPH Binding at the VinBLAST Active Site

**Fig. S12. Cinnamyl alcohol dehydrogenase (CAD) activities of CrC1150, VinBLAST, and CAD1.** Reactions were performed in triplicate using coniferyl aldehyde as the substrate for CrC1150 (Genbank KU865327) homodimer **(A)**, VinBLAST homodimer **(B)**, and CrCAD1 homodimer **(C)**. The results showed that CAD1 exhibited a $V_{max}$ that is 7.0-fold higher than that of VinBLAST, and a $K_m$ value that is 2.8-fold higher. However, we also unexpectedly identified CrC1150, a member of CAD clade I, as the genuine enzyme responsible for coniferyl aldehyde reduction in *C. roseus*. CrC1150 showed superior kinetic parameters for coniferyl aldehyde compared with both CAD2 and CAD1. Notably, CAD clade I includes bona fide cinnamyl alcohol dehydrogenases that are actually involved in lignin biosynthesis, as demonstrated both biochemically and genetically. These findings suggest that neither CAD1 nor VinBLAST (CAD2) serve as the primary lignin biosynthetic CAD in *C. roseus*, consistent with our proposal that VinBLAST functions primarily as a scaffold rather than a canonical CAD enzyme. Data were fitted using the Michaelis-Menten model and non-liner regression in GraphPad Prism version 10.4.2. The results represent the mean of three technical replicates.

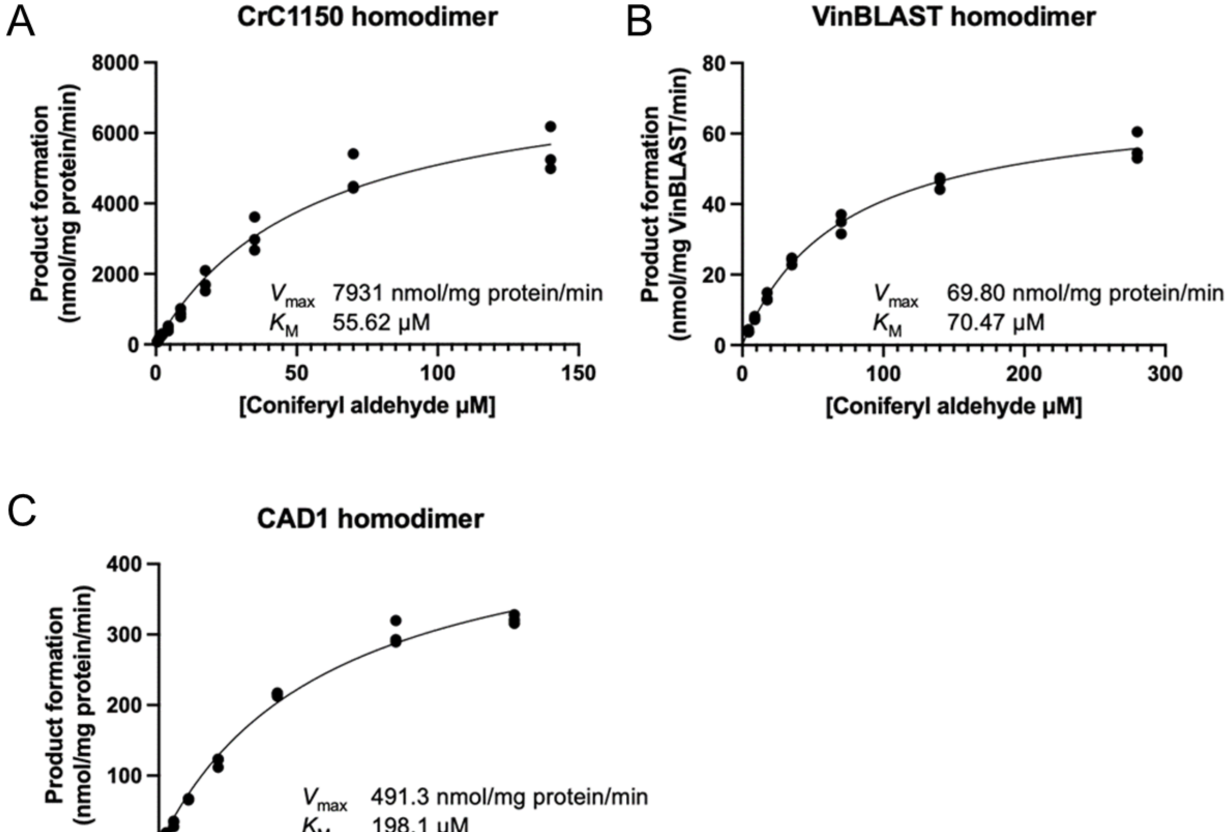

**Fig. S13. The yeast two-hybrid (Y2H) assays to investigate the VinBLASTs and GS interactions.** The Y2H results demonstrated that the heterodimer formed by VinBLAST and GS exhibited stronger binding affinity compared with either the VinBLAST or GS homodimers. The test proteins were fused to the GAL4 transcription activation domain (AD) and GAL4 DNA-binding domain (BD), respectively, at the *N*-terminus, and transformed into the Y2H-GOLD host strain with histidine deficiency. Upon protein-protein interactions, the *GAL* promoter is transcriptionally activated, enabling the yeast strains to express the *HIS3* gene and therefore grow on the histidine-deficient agar plates. Weakly interacting proteins, such as GS with GS, VinBLAST with VinBLAST, VinBLAST$^{IM}$ with GS, and GS$^{IM}$ with VinBLAST/VinBLAST$^{CM}$/VinBLAST$^{IM}$ failed to activate the *GAL* promoter and thus the corresponding strains could not grow. VinBLAST$^{CM}$ represents the catalytic mutant VinBLAST$^{C51A,H56A}$, VinBLAST$^{IM}$ represents the interaction mutant VinBLAST$^{M298E,V299E}$, and GS$^{IM}$ represents the interaction mutant GS$^{I301E}$. 3-AT (3-Amino-1,2,4-triazole) is a common competitive protein inhibitor used in Y2H experiments to suppress the leaky expression of *HIS3* and decrease the background noise. The yeast cultures, diluted at different ratios, were spotted onto SED amino acid-deficient plates containing no 3-AT, 5 mM 3-AT, and 20 mM 3-AT, respectively. The growth of the yeast clones was observed after incubation at 30°C for 96 h.

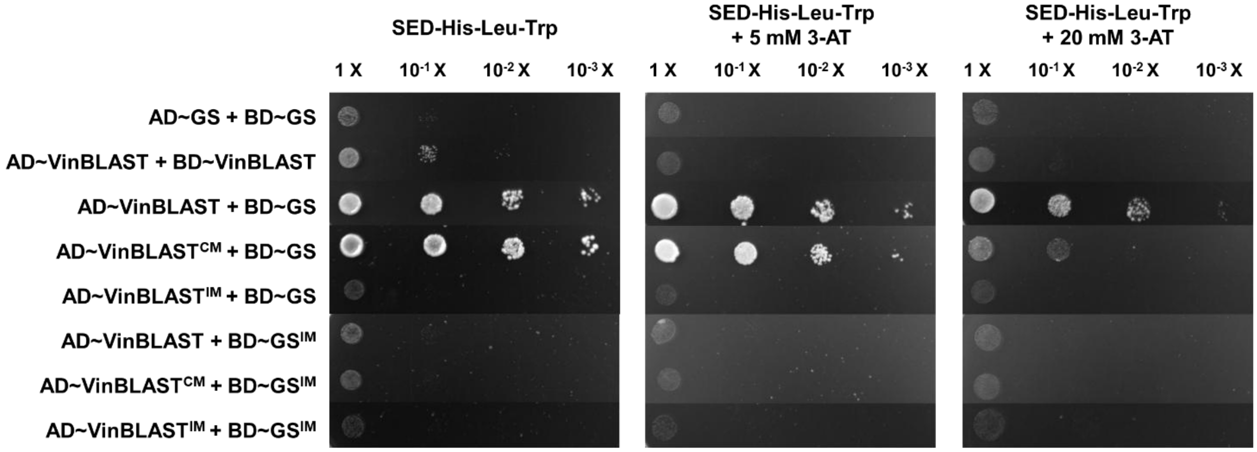

**Fig. S14. The Y2H assays to investigate SGD interactions.** The interaction between VinBLAST and GS was used as a positive control. No strong interactions were detected between SGD and SGD, HYS, GS, or VinBLAST in the Y2H system. Given that SGD is known to form an octamer and therefore must self-associate, the absence of a detectable signal for SGD-SGD in Y2H suggests that this assay has a high response threshold and is not sufficiently sensitive to reliably characterize weak protein-protein interactions involving SGD. The yeast cultures, diluted at different ratios, were spotted onto SED amino acid-deficient plates containing no 3-AT, 5 mM 3-AT, and 20 mM 3-AT, respectively. The growth of the yeast clones was observed after incubation at 30°C for 96 h.

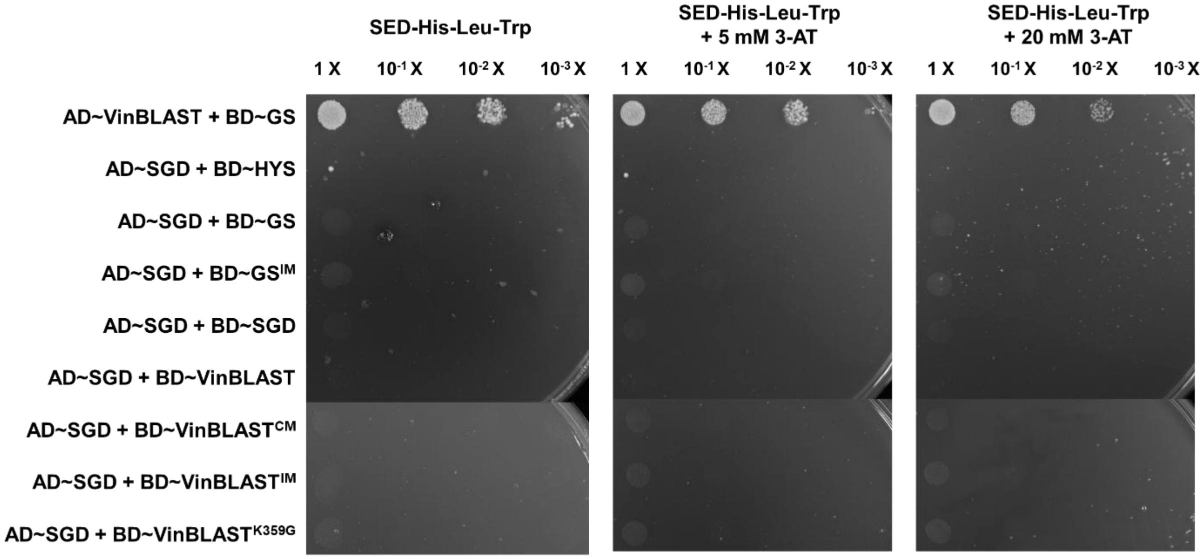

**Fig. S15. Definition of the potential interaction interfaces for SGD and VinBLAST. (A)** Potential interaction interfaces for SGD are colored in blue (interface-A), green (interface-B), and magenta (interface-C). **(B)** Potential interaction interfaces for VinBLAST are colored in orange (interface-A), magenta (interface-B), and purple (interface-C). As neither SGD nor VinBLAST is catalytically active as the monomer form, the interfaces involved in SGD homopolymerization, those responsible for VinBLAST-GS heterodimer formation, as well as the substrate entry/exit interfaces, were excluded. Accordingly, three potential interaction interfaces were defined for each of SGD and VinBLAST.

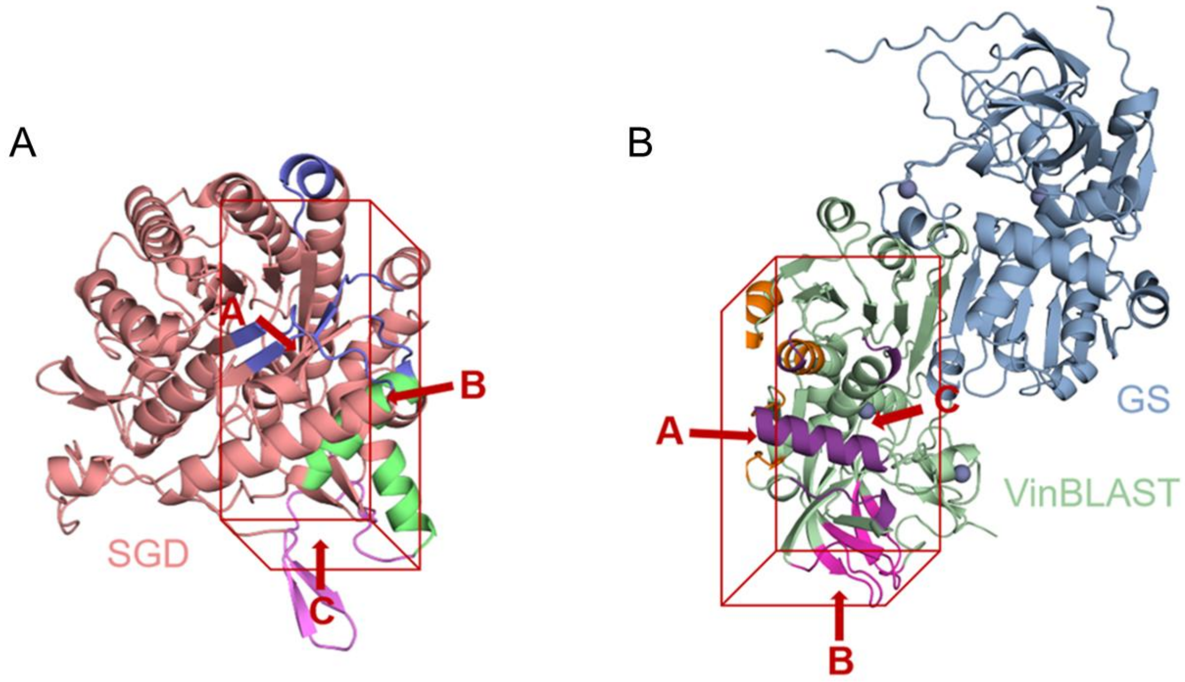

**Fig. S16. Protein-protein interaction (PPI) energy between each monomer of SGD and VinBLAST.** A/A and A/B systems exhibited weaker interactions between SGD and VinBLAST than other systems, inconsistent with experimental observations indicating clear interaction between SGD and VinBLAST, leading to the exclusion of these two configurations of SGD-GS complex for further consideration. Meanwhile, the B/B configuration of the SGD-GS complex failed to reach a stable state; consequently, no energy analysis under equilibrium conditions could be performed.

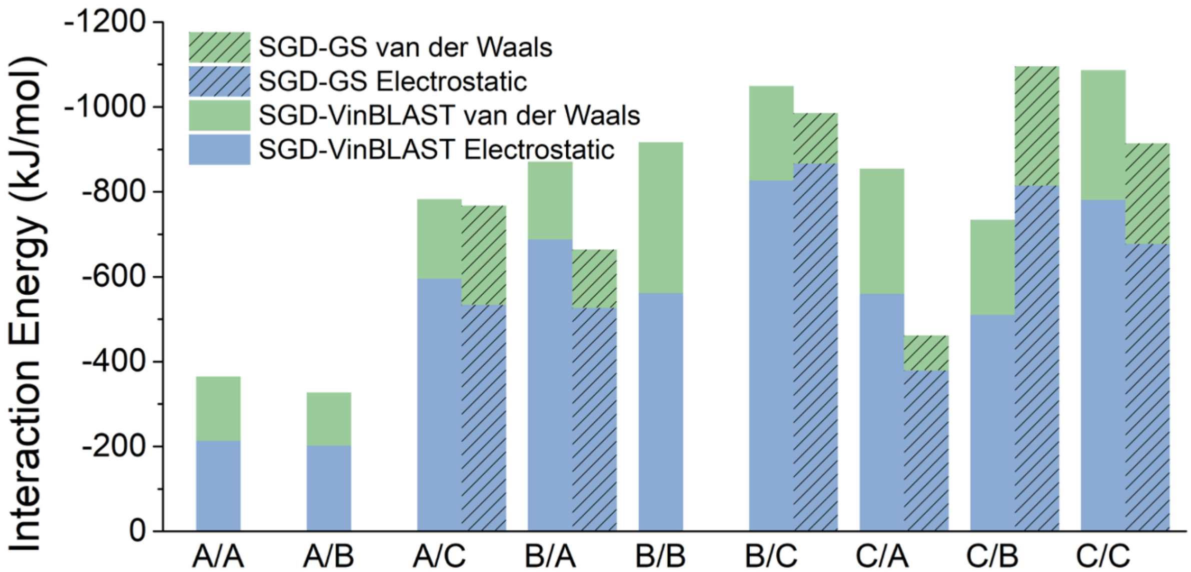

**Fig. S17. Importance of VinBLAST K359 for the formation of SGD and VinBLAST complex.** **(A)** Interaction energy between key binding residues in SGD and VinBLAST. **(B)** Protein-protein interaction energy between each monomer of SGD and VinBLAST. VinBLAST-WT: interaction between SGD and the wild type VinBLAST, VinBLAST$^{K359G}$: interaction between SGD and the K359G mutant of VinBLAST, GS-WT: interaction between SGD and the wild type GS. The K359G mutation decrease the interaction energy between VinBLAST and SGD. **(C)** Structural details of K359 in VinBLAST and its interacting residues (VinBLAST, shown in cyan; SGD, shown in salmon). **(D)** Structural details of responding residue K361 of GS (shown in pink) and its interacting residues (GS, shown in blue).

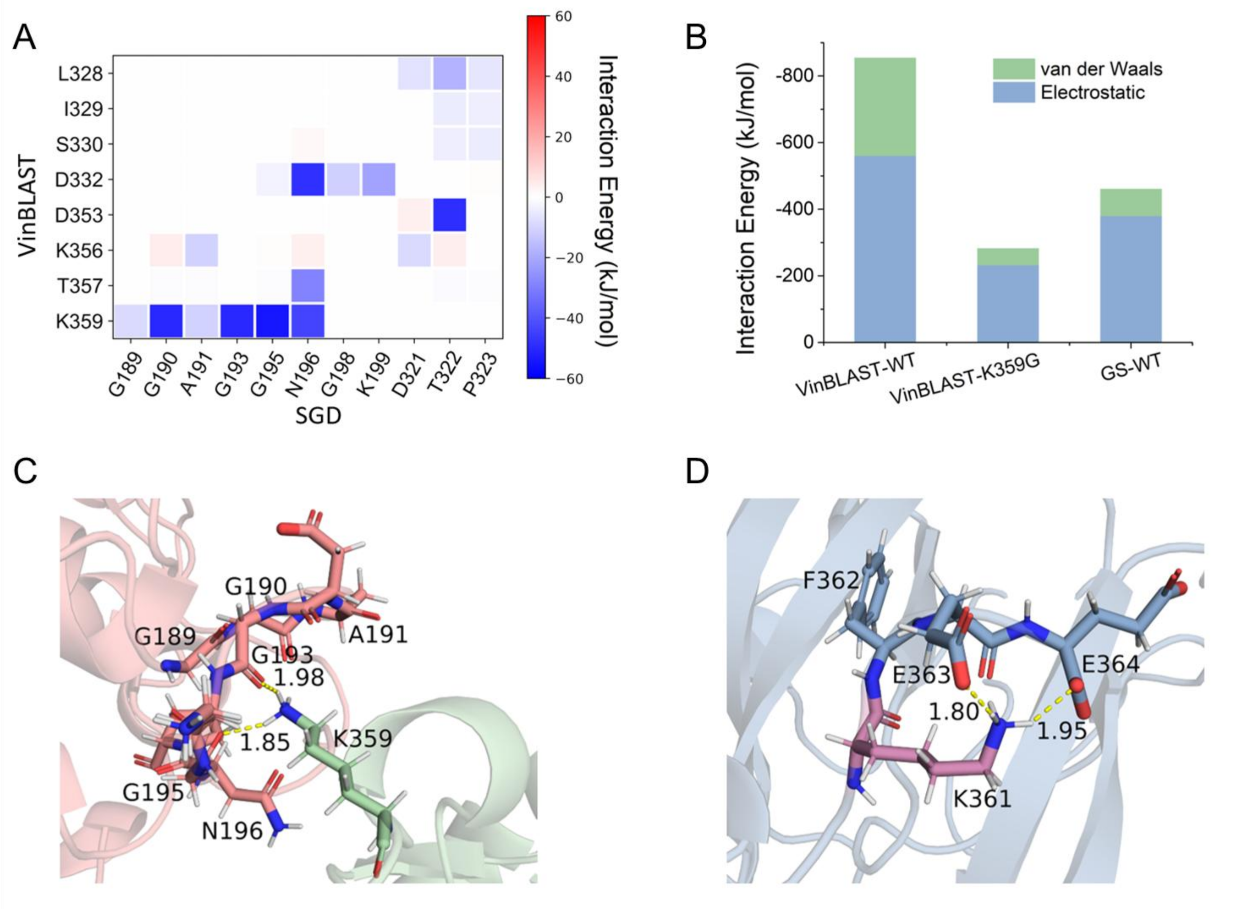

**Fig. S18. The BiFC experiments in yeast demonstrated protein-protein interactions among VinBLAST$^{K359G}$, GS, and SGD.** The *N*-terminal fragment (mVenus$^{N}$) and the *C*-terminal fragment (mVenus$^{C}$) of mVenus were individually fused to proteins of interest. Yellow fluorescence is generated when the two mVenus fragments are brought into proximity by interaction of the fused proteins. Considering that SGD has a nuclear localization signal (NLS) at the *C*-terminus, the mVenus fragments were fused to the *N*-terminus of SGD. For both VinBLAST$^{K359G}$ and GS, the mVenus fragments were fused to their *C*-termini. BiFC assays showed that VinBLAST$^{K359G}$ interacted with GS in the cytosol **(A)**. However, in contrast to wild type VinBLAST, the VinBLAST$^{K359G}$ mutant did not enable detectable interaction with SGD **(B)**. When untagged VinBLAST$^{K359G}$ was constitutively expressed, it was also unable to mediate detectable interaction between SGD and GS **(C)**, indicating that VinBLAST$^{K359G}$ cannot serve as a scaffold to bridge SGD and GS. Nab2 tagged with mCherry at the *C*-terminus functioned as a nuclear marker. All proteins were expressed in the wild-type CEN.PK2-1C yeast strains. The images were acquired using a confocal laser scanning microscope (Olympus FV3000) at 600× magnification, with a 5 μm scale bar displayed in the upper left corner.

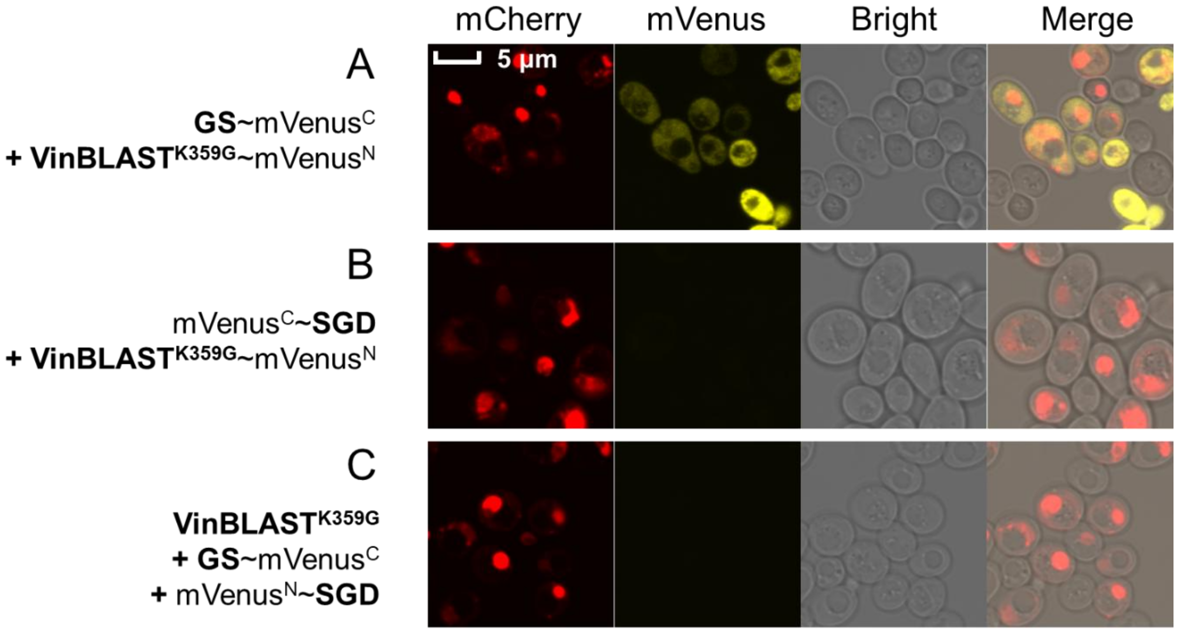

**Fig. S19. The Y2H assays to investigate the VinBLAST$^{K359G}$ and GS interactions.** The test proteins were fused to the GAL4 transcription activation domain (AD) and GAL4 DNA-binding domain (BD), respectively, at the *N*-terminus, and transformed into the Y2H-GOLD host strain with histidine deficiency. Upon protein-protein interactions, the *GAL* promoter is transcriptionally activated, enabling the yeast strains to express the *HIS3* gene and therefore grow on the histidine-deficient agar plates. Weakly interacting proteins failed to activate the *GAL* promoter and thus the corresponding strains could not grow. 3-AT (3-Amino-1,2,4-triazole) is a common competitive protein inhibitor used in Y2H experiments to suppress the leaky expression of *HIS3* and decrease the background noise. The yeast cultures, diluted at different ratios, were spotted onto SED amino acid-deficient plates containing no 3-AT, 5 mM 3-AT, and 20 mM 3-AT, respectively. The growth of the yeast clones was observed after incubation at 30°C for 96 h.

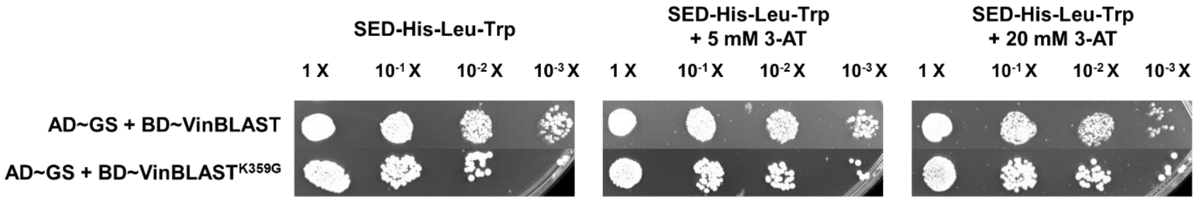

**Fig. S20. The SPR assays to investigate VinBLAST, VinBLAST$^{K359G}$, SGD, and GS interactions.** VinBLAST, VinBLAST$^{K359G}$, SGD, or GS were immobilized on the sensor chip, and the analyte proteins were injected over the immobilized ligand surface at gradient concentrations (0.06-4 μM). At the concentrations tested (0.06-2 μM), no binding between 16OMT and VinBLAST **(A)** or 16OMT and VinBLAST$^{K359G}$ **(B)** were detected, as the negative controls. Interaction between VinBLAST and SGD (Kd=1.48 μM) **(C)** was stronger than the interaction between VinBLAST$^{K359G}$ and SGD (Kd=2.87 μM) **(D).** Meanwhile, VinBLAST and GS (Kd=0.48 μM) **(E)** showed comparable interaction as VinBLAST$^{K359G}$ and GS (Kd=0.48 μM) **(F)**. Experiments were repeated twice with similar results.

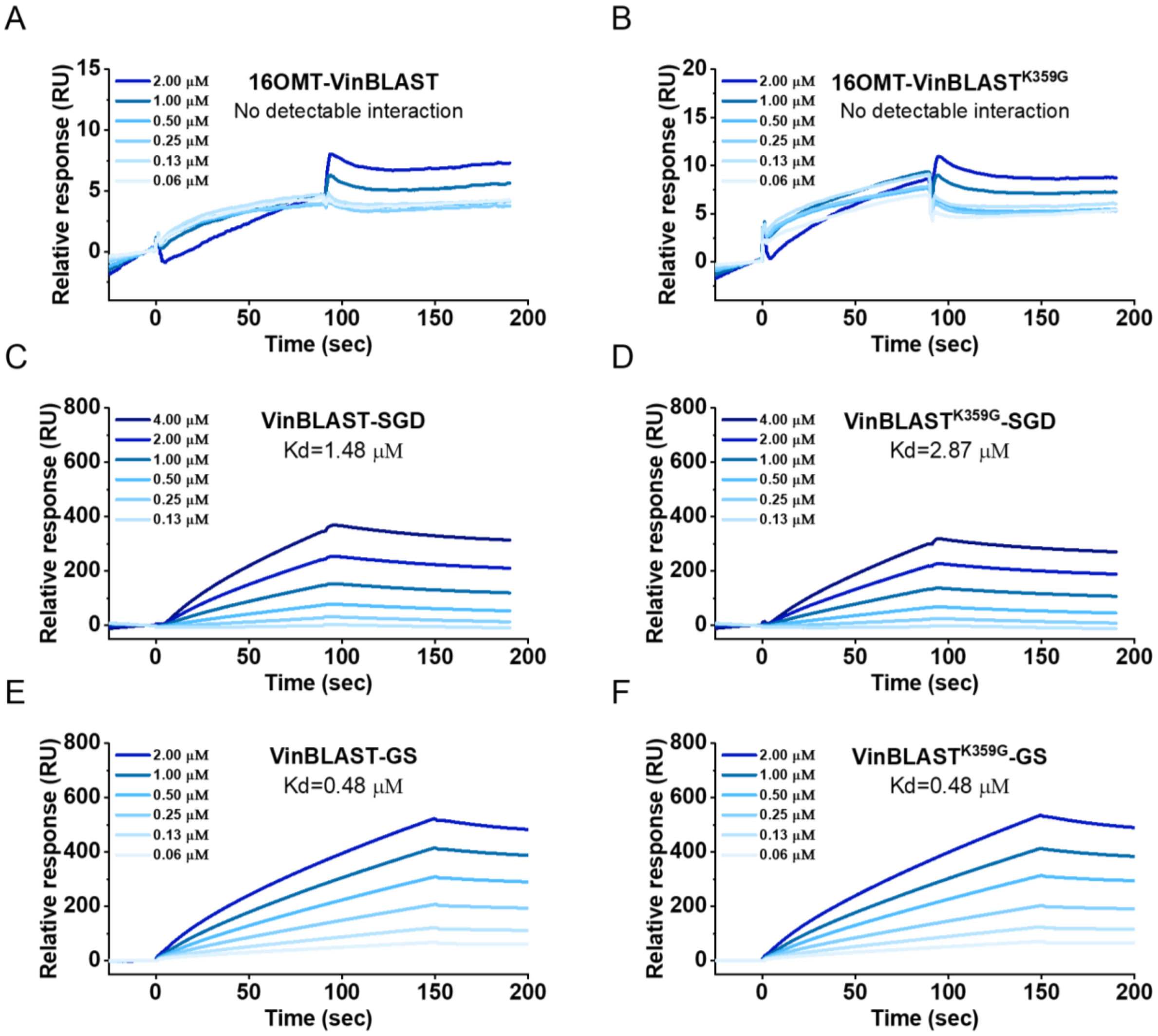

**Fig. S21. Subcellular localization of SGD and its nuclear localization signal (NLS) truncated version tSGD.** While wild-type SGD localized to the nucleus **(A)**, tSGD shifted its subcellular localization to the cytoplasm **(B)**. SGD and tSGD were fused to EGFP at the *N*-terminus. The red fluorescence was Nab2 tagged with mCherry at the *C*-terminus functioned as a nuclear marker. The tagged proteins were all expressed in the wild-type CEN.PK2-1C strain. The images were acquired using a confocal laser scanning microscope (Olympus FV3000) at 600× magnification, with a 5 μm scale bar displayed in the upper left corner.

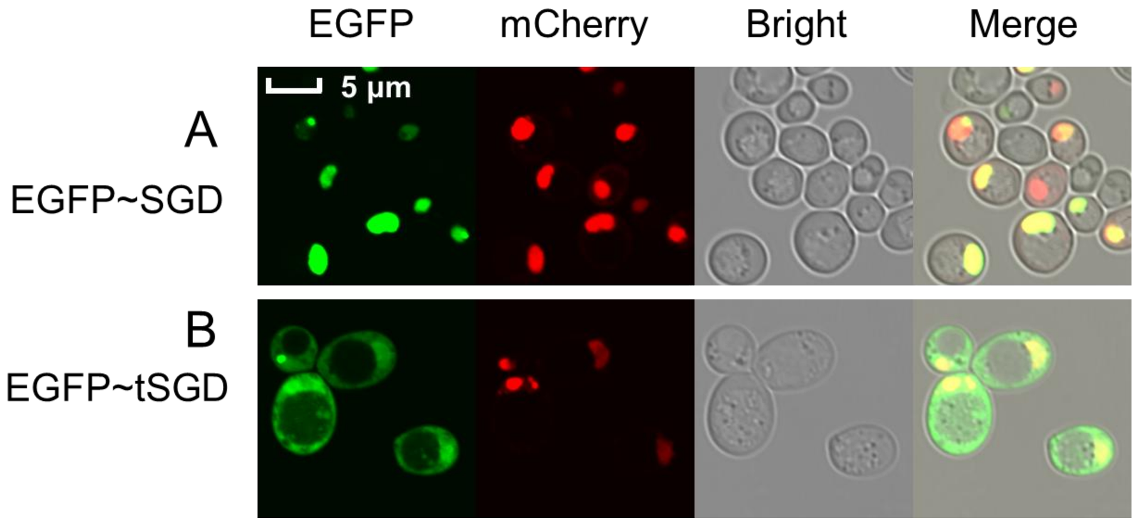

**Fig. S22. Artificial scaffolding improves *de novo* geissoschizine methyl ether (GME) production in engineered *S. cerevisiae* strains.** These strains contain modifications in SGD subcellular localization, GS~SGD fusion, tethering SGD/tSGD and GS with self-interacting peptide tags RIDD/RIAD and SpyTag/SpyCatcher, and GS copy numbers. Among these approaches, artificial scaffolding via self-interacting tags yielded the strongest improvements next to the use of VinBLAST, increasing GME titer by approximately 3.3- to 3.7-fold relative to baseline strains. In comparison, the GS~SGD fusion construct and increasing GS gene copy number resulted in modest enhancements, 1.2-fold and 2.0-fold, respectively. The parental strain was GM02 with SGD removed and the corresponding genomic modifications for each strain were specified underneath. The strain genotypes were listed in table S1. The results represent the mean ± s.d. from three biological replicates.

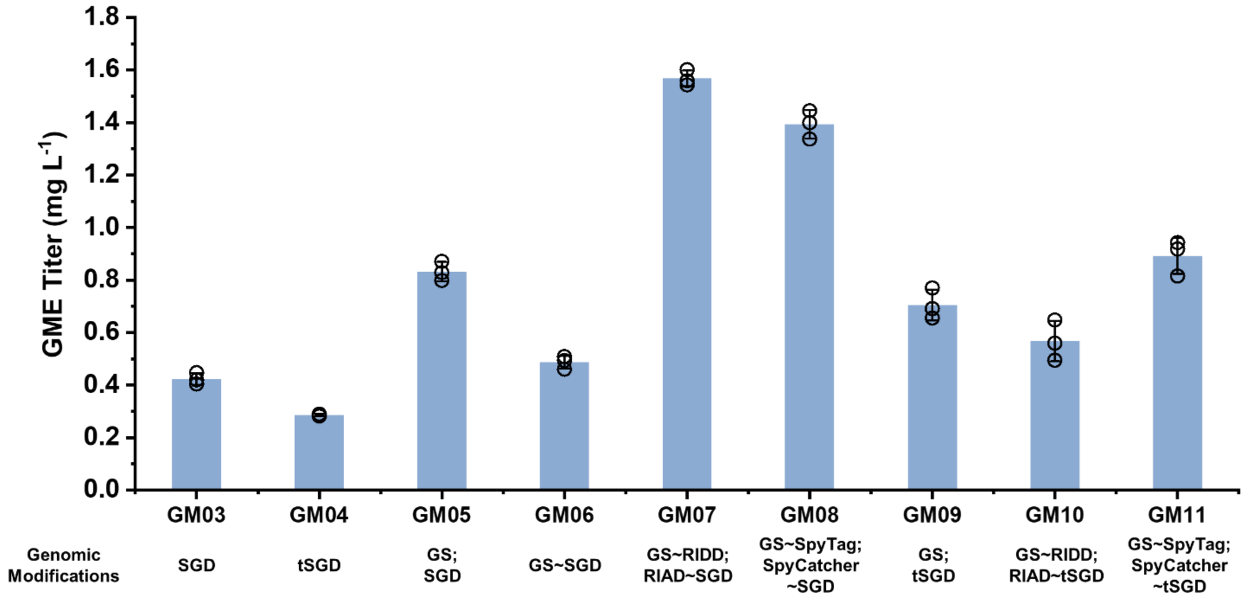

**Fig. S23. Artificial scaffolding improves *de novo* vindoline production in engineered *S. cerevisiae* strains.** These strains contain modifications in SGD subcellular localization, GS~SGD fusion, tethering SGD and GS with self-interacting peptide tags RIDD/RIAD and SpyTag/SpyCatcher, and GS copy numbers. Among these approaches, artificial scaffolding via self-interacting tags yielded the strongest improvements next to the use of VinBLAST, increasing GME titer by approximately 1.7- to 2.3-fold relative to baseline strains. In comparison, the GS~SGD fusion construct and increasing GS gene copy number resulted in modest enhancements, 1.7-fold and 1.6-fold, respectively. The parental strain was VI05 (complete vindoline biosynthetic pathway with SGD removed) and the corresponding genomic modifications for each strain were specified underneath. The strain genotypes were listed in table S1. The results represent the mean ± s.d. from three biological replicates.

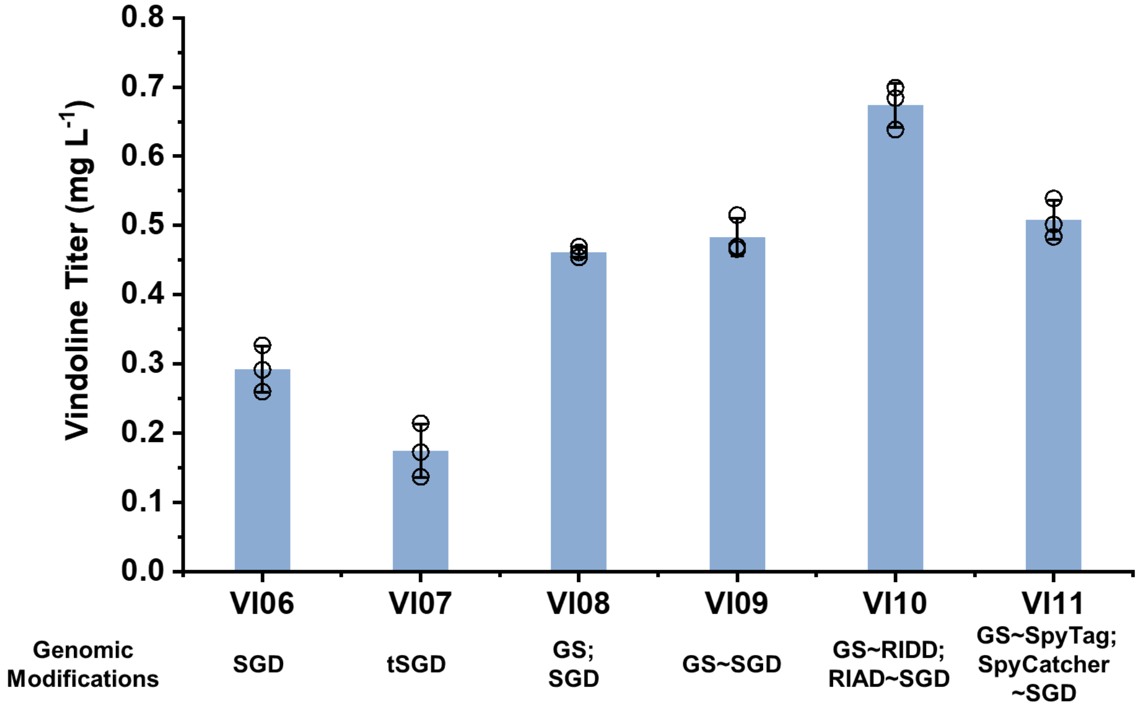

**Fig. S24. VinBLAST outperforms artificial scaffolding for *de novo* biosynthesis of geissoschizine methyl ether (GME) in engineered *S. cerevisiae* strains.** No difference in GME production was observed between wild-type VinBLAST and its catalytic mutant (VinBLAST$^{CM}$; C51A and H56A mutations disrupting NADPH binding), indicating that catalytic activity is not required for enhancement. In contrast, the interaction mutant (VinBLAST$^{IM}$; M298E and V299E mutations weakening dimer formation) showed decreased GS-enhancing activity, suggesting that physical dimerization is essential. The parental strain was AJM7-ΔHYS (*4*) and the corresponding genomic modifications for each strain were specified underneath. The strain genotypes were listed in table S1. The results represent the mean ± s.d. from three biological replicates.

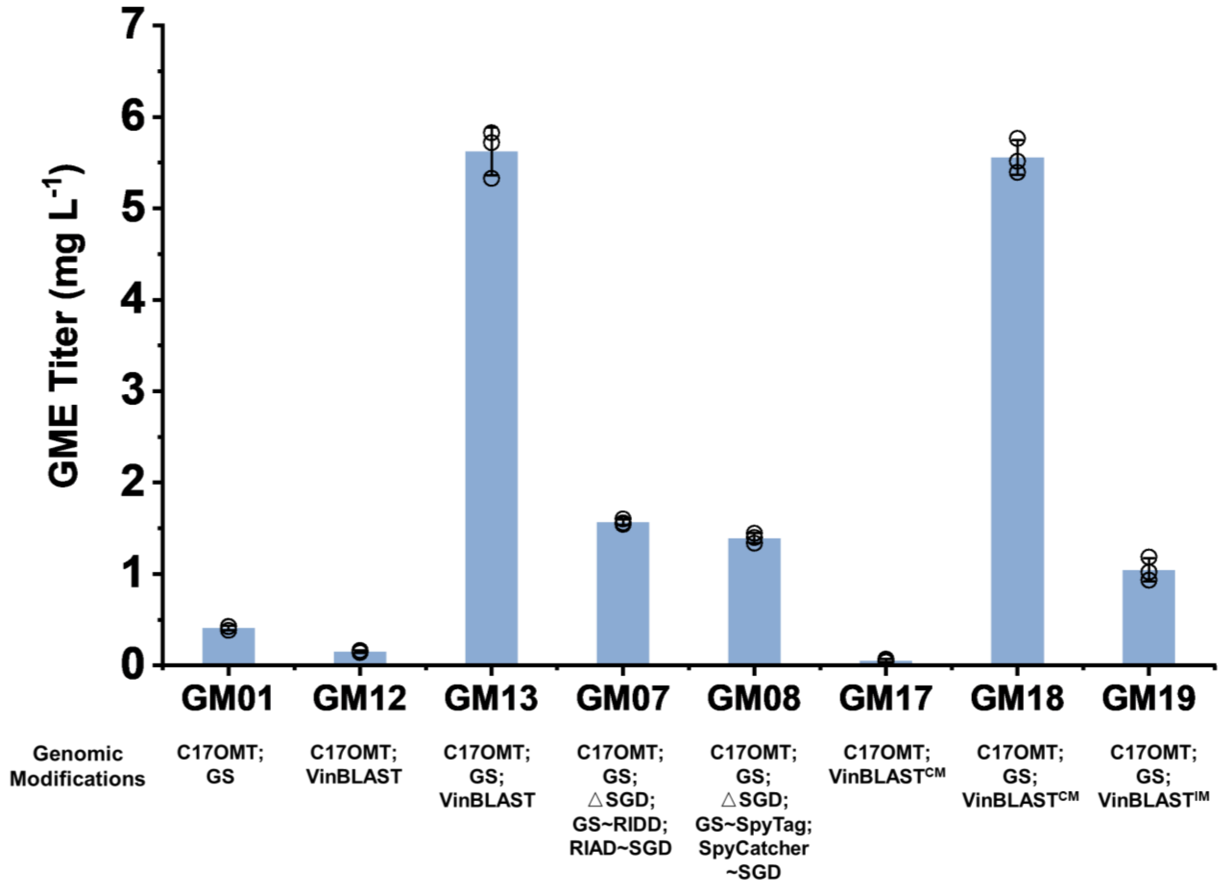

**Fig. S25. VinBLAST outperforms artificial scaffolding for *de novo* biosynthesis of vindoline in engineered *S. cerevisiae* strains.** The parental strain was VIN13 (*4*) and the corresponding genomic modifications for each strain were specified underneath. VinBLAST$^{CM}$ represents the catalytic mutant VinBLAST$^{C51A,H56A}$ and VinBLAST$^{IM}$ represents the interaction mutant VinBLAST$^{M298E,V299E}$. The strain genotypes were listed in table S1. The results represent the mean ± s.d. from three biological triplicates.

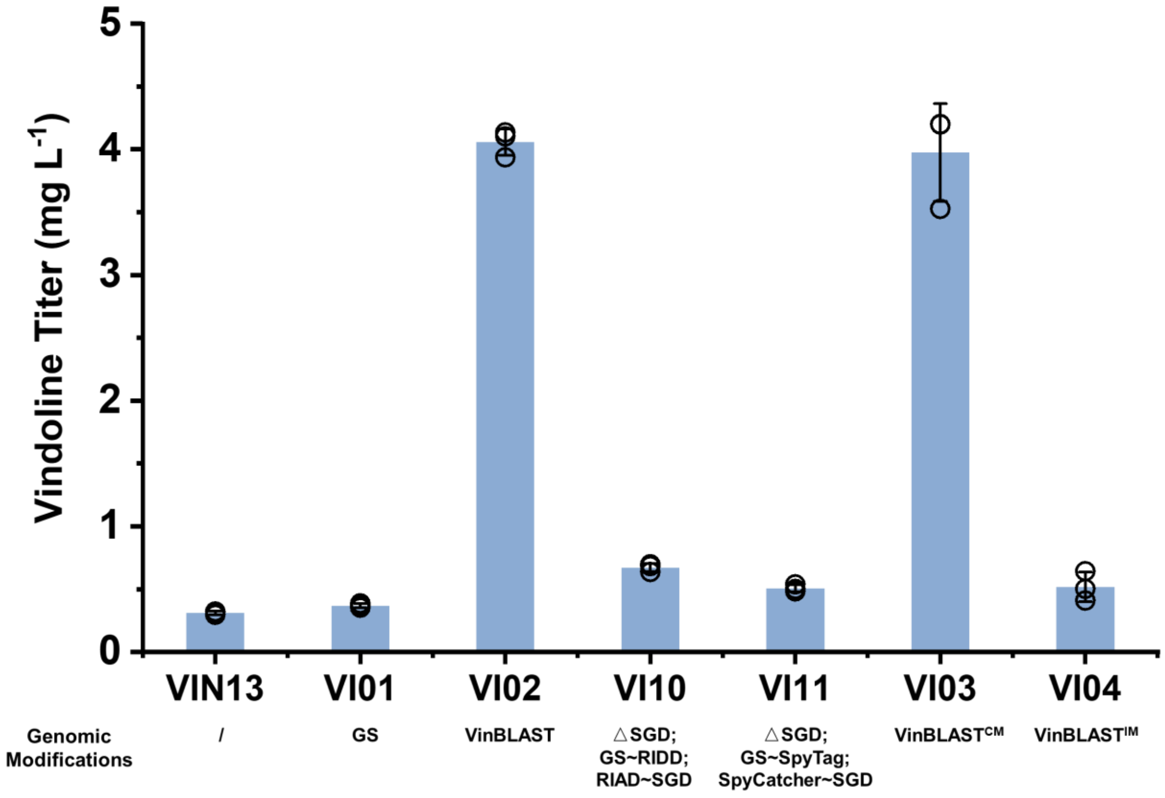

**Fig. S26. Optimal VinBLAST to GS ratio for enhancing activity lied between 4:1 and 1:2.** Various ratios of VinBLAST and GS were tested in in vitro assays containing strictosidine, SGD, and C17OMT to assess the production of geissoschizine methyl ether (GME). In the absence of VinBLAST, only small amounts of GME 19-epimers were detected. Increasing the VinBLAST:GS ratio led to a marked increase in GME production, with maximal enhancement observed within the 4:1 to 1:2 range. At ratios 4:1, the enhancement effect diminished, suggesting a saturation point or potential inhibitory effect of excess VinBLAST. The LC-MS/MS chromatogram showed $[M+H]^+$ ion transitions of 531>352 for strictosidine, 351>170 for strictosidine aglycone, 367>144 for GME.

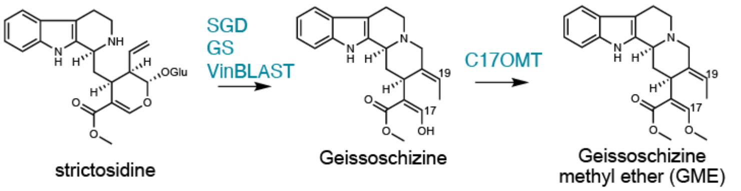


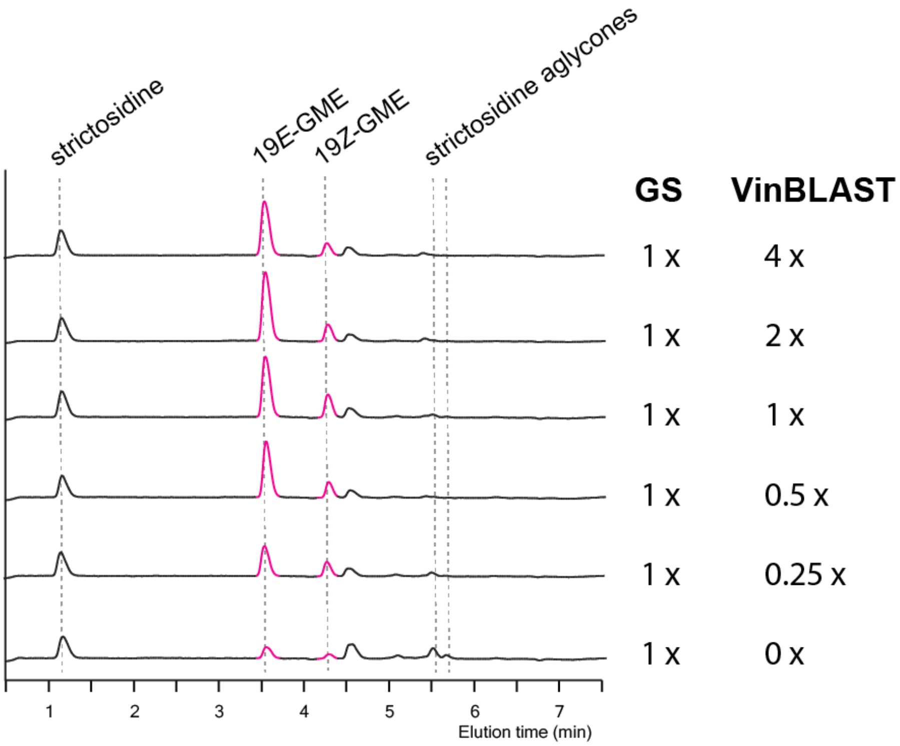

**Fig. S27. The enhanced GS activity by VinBLAST was independent of SGD. (A)** GS activity was evaluated with or without the presence of SGD. The GS+VinBLAST assay contain strictosidine aglycone, GS, and VinBLAST to assess the production of geissoschizine. The GS+VinBLAST+SGD assay contain strictosidine, GS, VinBLAST, and SGD, to assess the production of geissoschizine. The GS+VinBLAST$^{K359G}$ assay contain strictosidine aglycone, GS, and VinBLAST$^{K359G}$ to assess the production of geissoschizine. With the same amount of substrate and each enzyme (recombinant GS and VinBLAST at a 1:1 ratio), it resulted in comparable geissoschizine yield (13~14-fold increase over the control assay without VinBLAST). The results represent the mean ± s.d. of three technical triplicates. **(B)** The representative chromatograms for the bar graph.

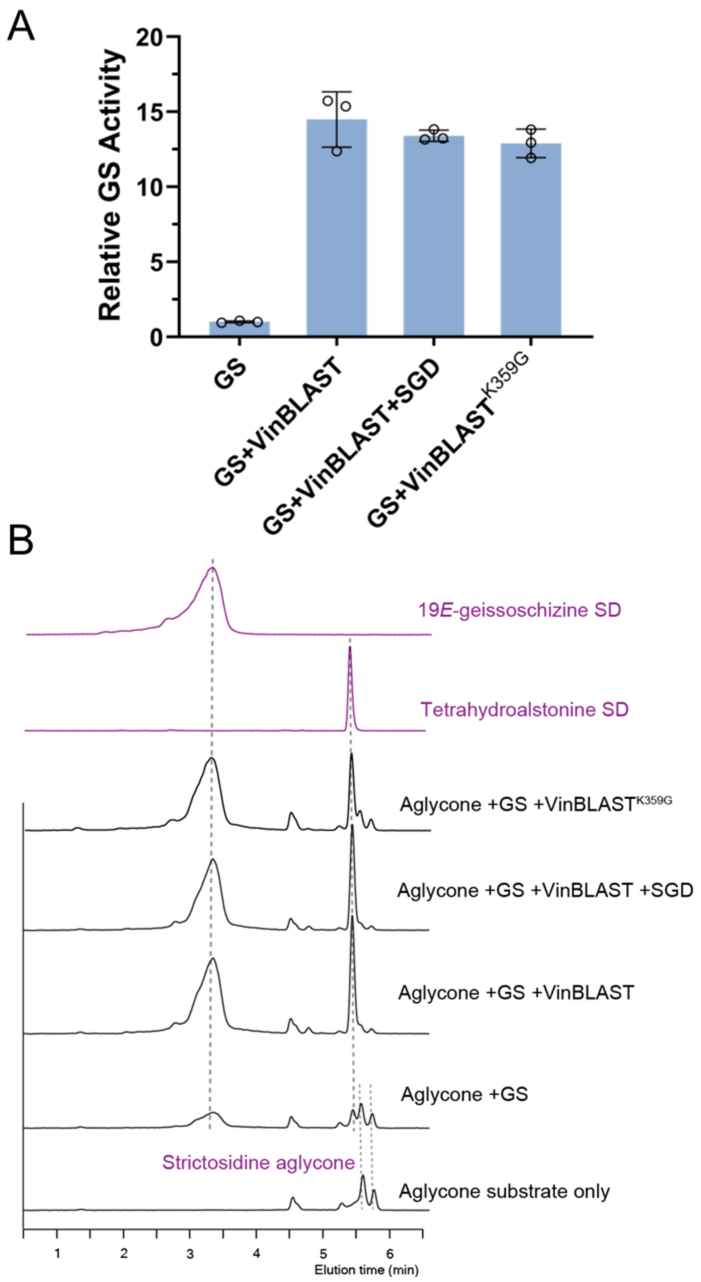

**Fig. S28. Michaelis-Menten kinetics of GS homodimer and heterologous dimers.** Reactions were performed in triplicate using strictosidine aglycone at concentrations ranging from 0.09375 to 24 µM. **(A)** GS homodimer. **(B)** VinBLAST-GS complex. **(C)** VinBLAST$^{CM}$-GS complex. **(D)** VinBLAST$^{IM}$-GS complex. **(E)** GS$^{TM}$ homodimer. **(F)** VinBLAST$^{TM}$-GS complex. Data were fitted using the Michaelis-Menten model and non-liner regression in GraphPad Prism version 10.4.2. The results represent the mean of three technical replicates.

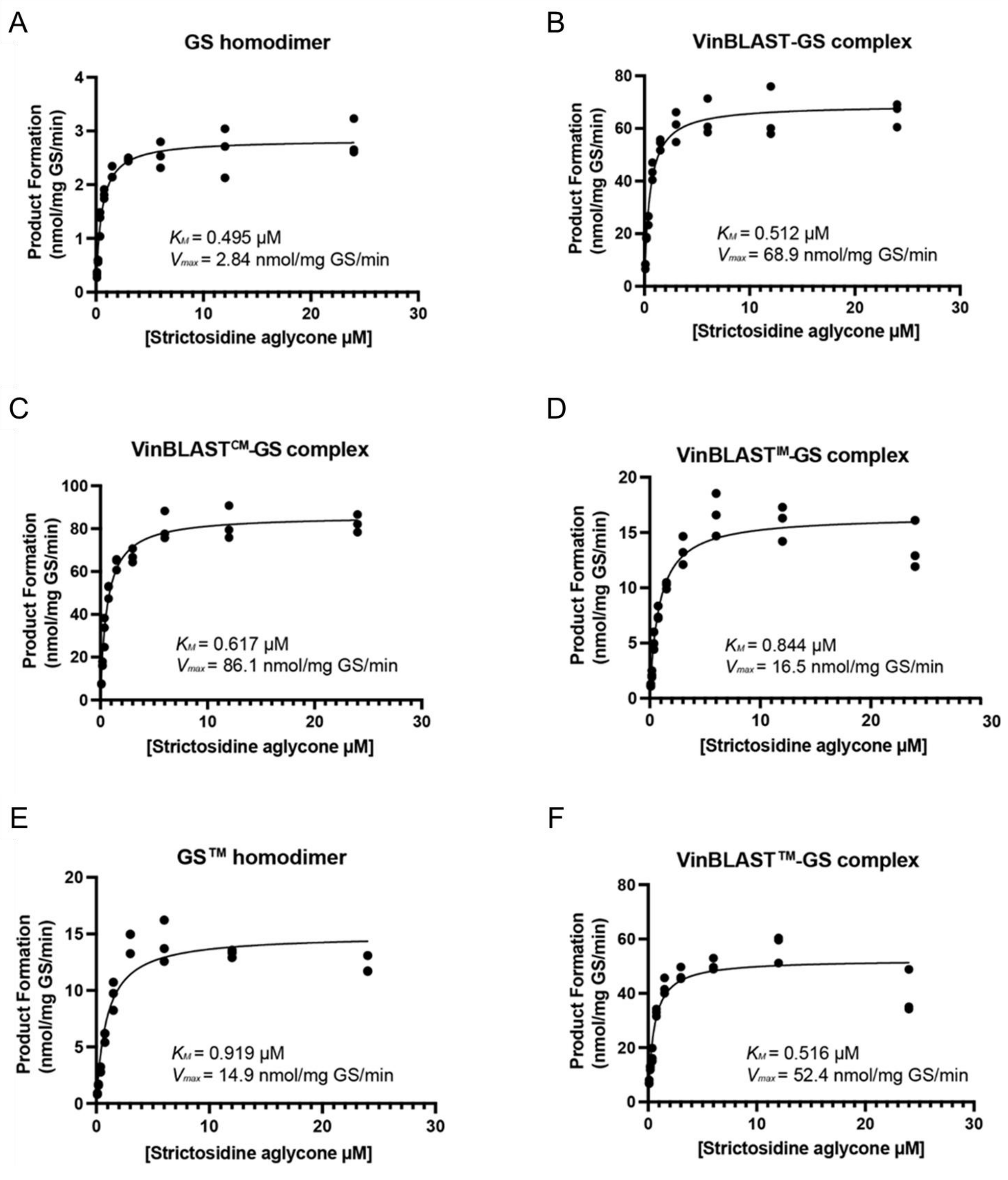

**Fig. S29. Native PAGE analysis indicated the formation of GS and VinBLAST heterodimer.** Purified, *N*-terminal His-tagged recombinant proteins were separated by 4~20% Bis-Tris SDS-PAGE **(A)** and 4~15% Tris-Glycine Native-PAGE **(B)**. Each lane contained 2 µg of each protein. When analyzed individually, both CrGS (theoretical molecular weight for monomer: 45.0 kDa) and CrVinBLAST (theoretical molecular weight for monomer: 43.9 kDa) migrated as homodimers, with no detectable monomeric bands. This is consistent with the crystal structure of GS and the general property of cinnamyl alcohol dehydrogenase (CAD)-like enzymes, which typically function as homodimers. When mixed together, CrGS and CrVinBLAST formed a heterodimer. Notably, no evidence of tetramer formation was observed. Their migration behavior on denaturing SDS-PAGE was consistent with their theoretical monomeric molecular weights.

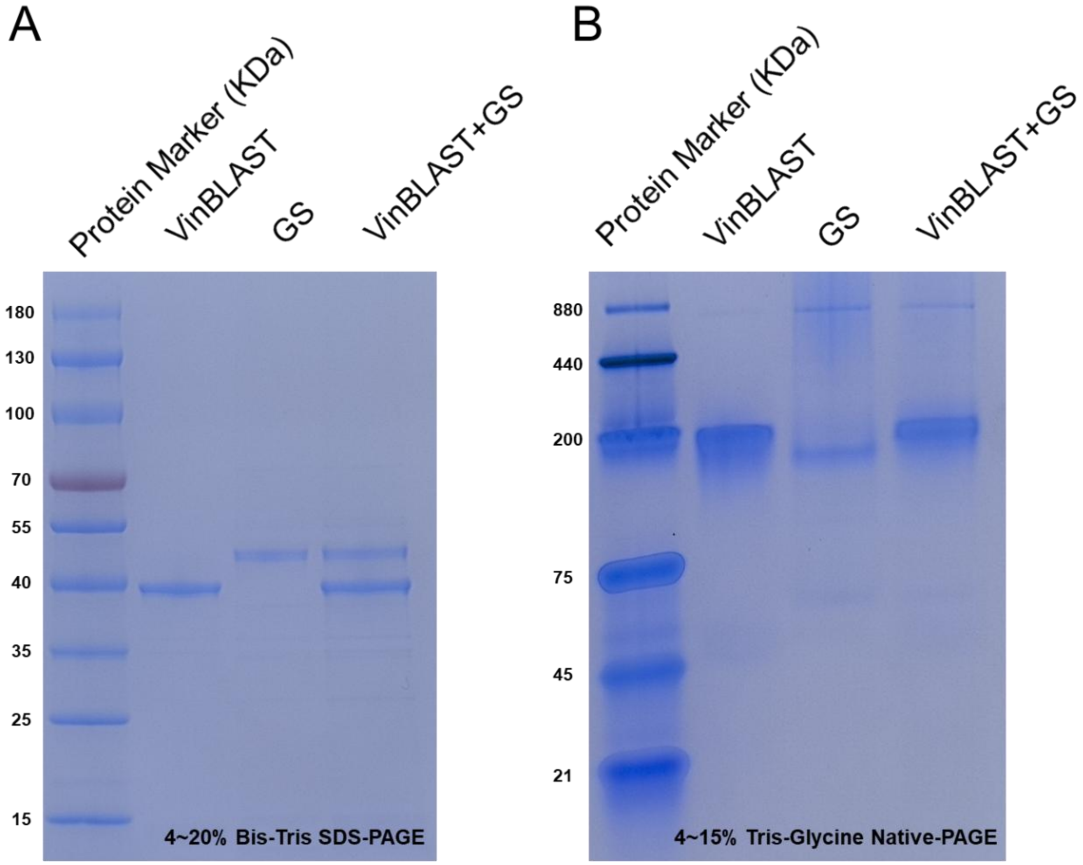

**Fig. S30. Identification of key residues in the dimer interface of GS. (A)** Protein-protein interaction (PPI) energy between each monomer in different dimers. **(B)** Interaction energy between VinBLAST and key binding residues in GS. **(C)** Interaction energy between key binding residues in GS-GS homodimer. **(D)** Interaction energy between key binding residues in $GS^{IM}$-$GS^{IM}$. **(E)** Interaction energy between key binding residues in VinBLAST-GS. **(F)** Distance of one interacting β-sheet pair at the GS-GS homodimer. **(G)** Distance and the structural details of one interacting β-sheet pair at the $GS^{IM}$-$GS^{IM}$.

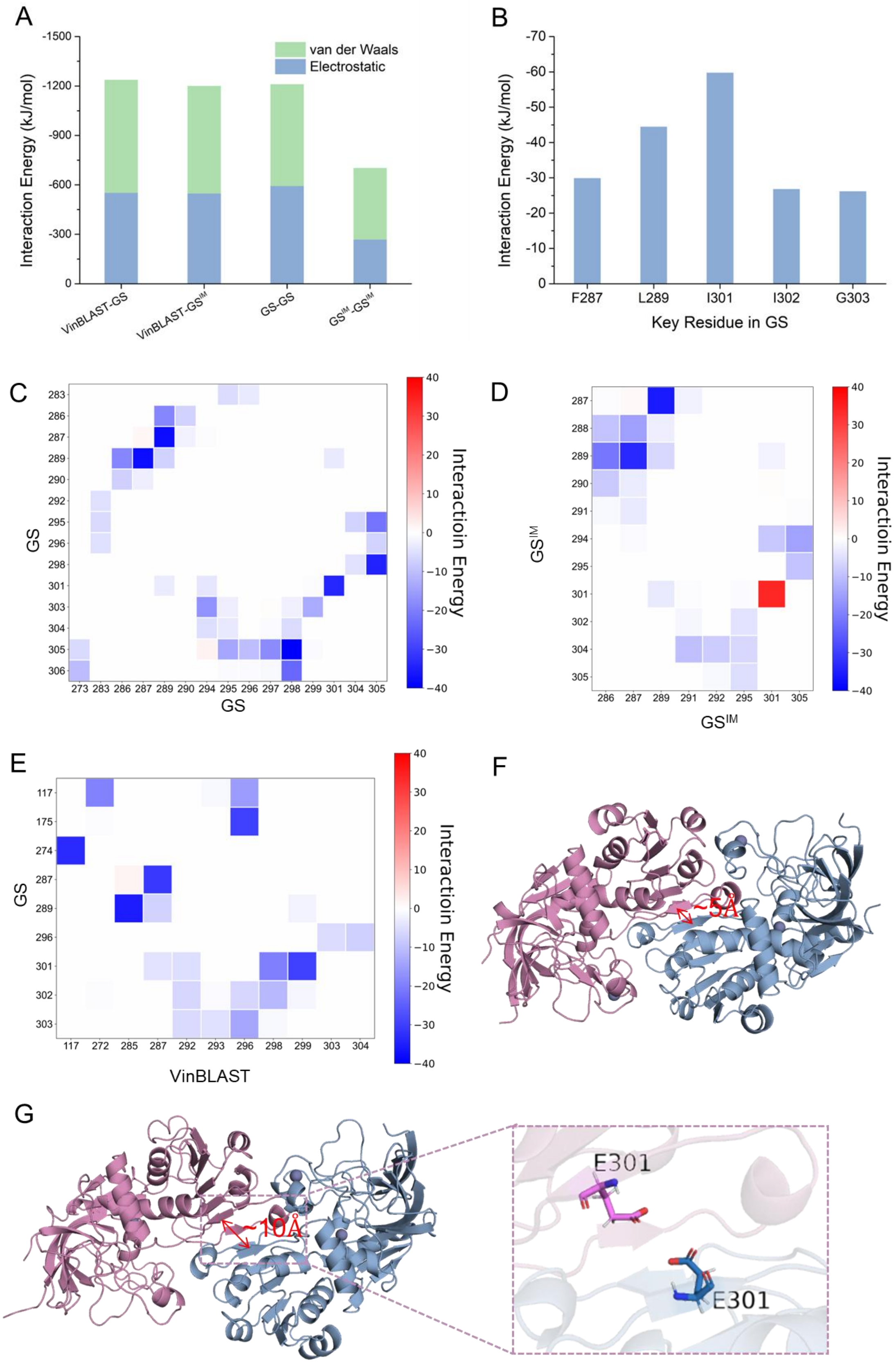
A
van der Waals
Electrostatic
Interaction Energy (kJ/mol)
VinBLAST-GS
VinBLAST-GS^IM
GS-GS
GS^IM-GS^IM
B
Interaction Energy (kJ/mol)
F287
L289
I301
I302
G303
Key Residue in GS
C
GS
Interactioin Energy
D
GS^IM
Interactioin Energy
E
GS
VinBLAST
Interactioin Energy
F
~5Å
G
~10Å
E301
E301

**Fig. S31. The bimolecular fluorescence complementation (BiFC) results of VinBLAST$^{CM}$, VinBLAST$^{IM}$, GS, and GS$^{IM}$ interactions in *S. cerevisiae*.** The *N*-terminal fragment (mVenus$^{N}$) and the *C*-terminal fragment (mVenus$^{C}$) of mVenus were separately fused to proteins of interest. Yellow fluorescence is generated when the two mVenus fragments are brought into proximity by interaction of the fused proteins. For both VinBLAST and GS, the mVenus fragments were fused to their *C*-termini. VinBLAST$^{CM}$ represents the catalytic mutant VinBLAST$^{C51A,H56A}$, VinBLAST$^{IM}$ represents the interaction mutant VinBLAST$^{M298E,V299E}$, and GS$^{IM}$ represents the interaction mutant GS$^{I301E}$. Nab2 tagged with mCherry at the *C*-terminus functioned as a nuclear localization marker (red fluorescence). All proteins were expressed in the wild-type yeast CEN.PK2-1C strain. The images were acquired using a confocal laser scanning microscope (Olympus FV3000) at 600× magnification, with a 5 μm scale bar displayed in the upper left corner.

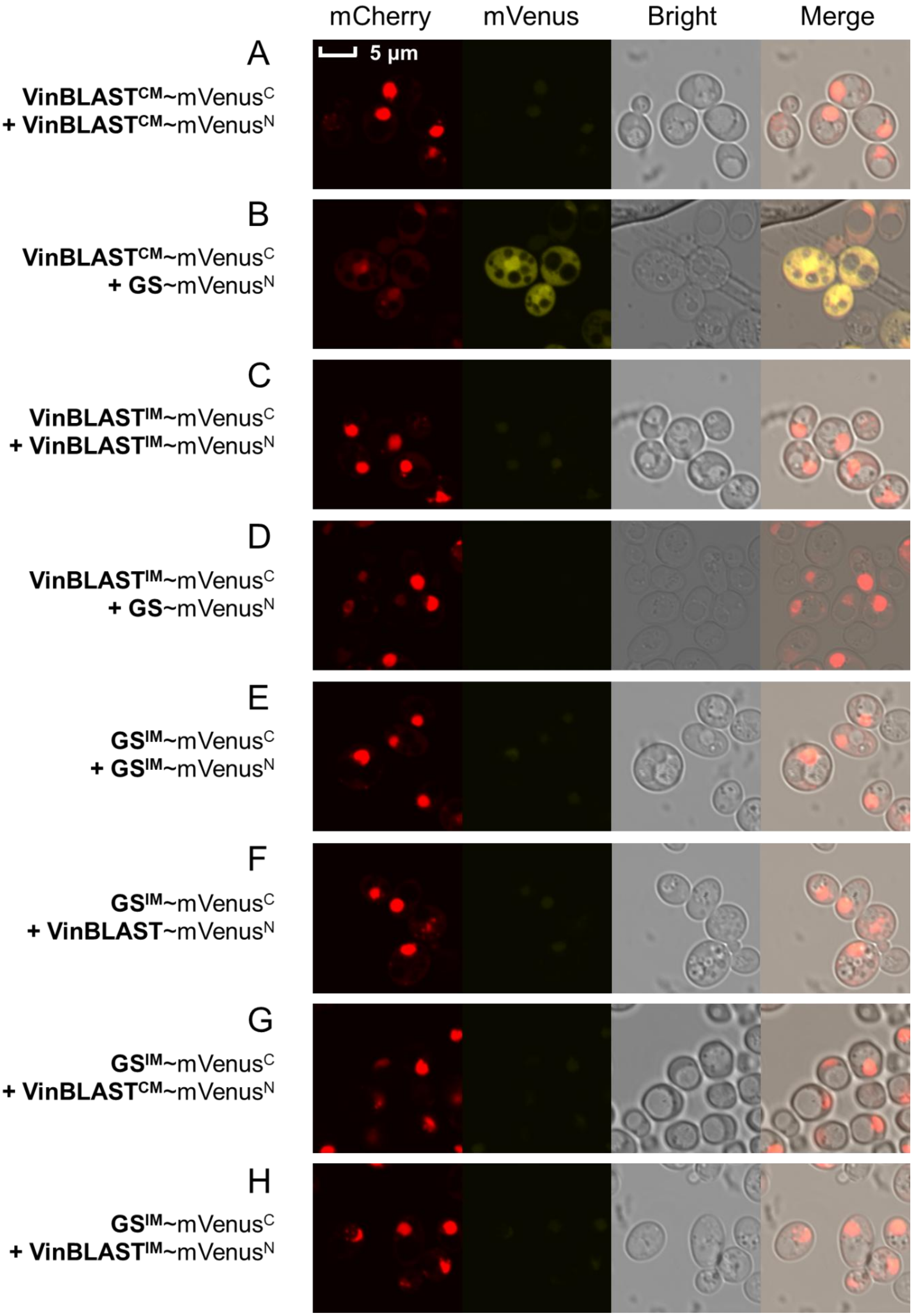
mCherry
mVenus
Bright
Merge
5 µm
A
VinBLAST$^{CM}$~mVenus$^{C}$
+ VinBLAST$^{CM}$~mVenus$^{N}$
B
VinBLAST$^{CM}$~mVenus$^{C}$
+ GS~mVenus$^{N}$
C
VinBLAST$^{IM}$~mVenus$^{C}$
+ VinBLAST$^{IM}$~mVenus$^{N}$
D
VinBLAST$^{IM}$~mVenus$^{C}$
+ GS~mVenus$^{N}$
E
GS$^{IM}$~mVenus$^{C}$
+ GS$^{IM}$~mVenus$^{N}$
F
GS$^{IM}$~mVenus$^{C}$
+ VinBLAST~mVenus$^{N}$
G
GS$^{IM}$~mVenus$^{C}$
+ VinBLAST$^{CM}$~mVenus$^{N}$
H
GS$^{IM}$~mVenus$^{C}$
+ VinBLAST$^{IM}$~mVenus$^{N}$

**Fig. S32. GS$^{IM}$ (GS$^{I301E}$) abolished catalytic activity for *de novo* GME and vindoline production in engineered *S. cerevisiae* strains.** These strains contain modifications in GS$^{IM}$ (GS interaction mutant; I301E). The parental strains were AJM7-ΔHYS (*4*) for GME-producing strains **(A)** and VIN13 (*4*) for vindoline-producing strains **(B)**. The corresponding genomic modifications for each strain were specified underneath. VinBLAST$^{CM}$ represents the catalytic mutant VinBLAST$^{C51A,H56A}$, VinBLAST$^{IM}$ represents the interaction mutant VinBLAST$^{M298E,V299E}$, and GS$^{IM}$ represents the interaction mutant GS$^{I301E}$. The strain genotypes were listed in table S1. The results represent the mean ± s.d. of three biological replicates.

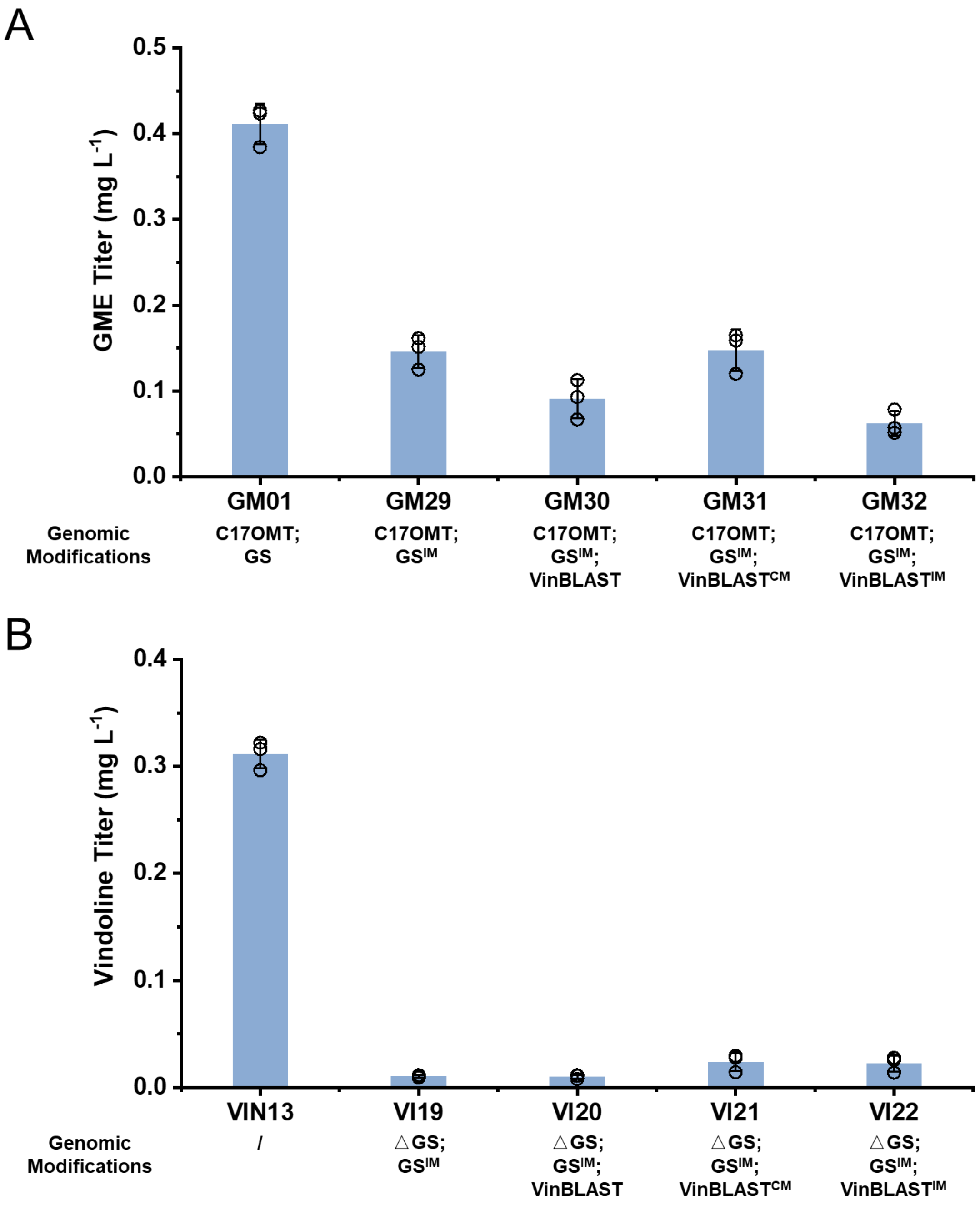

**Fig. S33. Inability of the substrate (4,21-dehydrogeissoschizine) to maintain stable binding at the active site of GS monomer. (A)** The docking result of substrate-NADPH-GS monomer. **(B)** The distance between centroid of the substrate and the catalytic zinc ion during 1,000 ns MD simulations. **(C)** The conformations of substrate-NADPH-GS complex at 0 ns and 1,000 ns, respectively. **(D)** Schematic diagram of the formation of a tunnel by GS-GS homodimer. **(E)** Schematic diagram of the formation of a tunnel by VinBLAST-GS heterodimer. Tunnel is shown as purple surface.

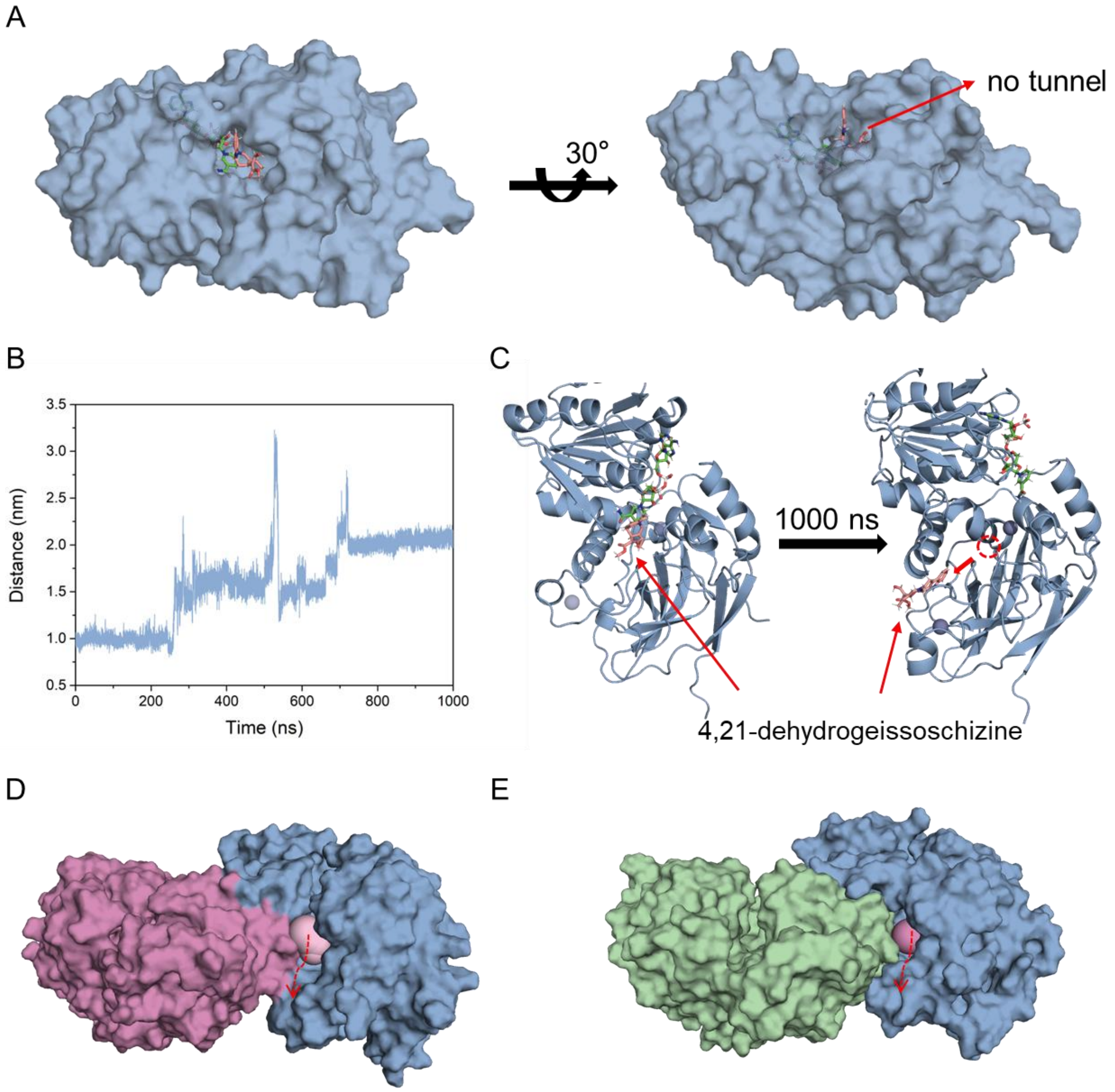

**Fig. S34. Structure of VinBLAST-GS heterodimer.** Key interactions were highlighted in dashed rectangle. Cyan: VinBLAST. Blue: GS.

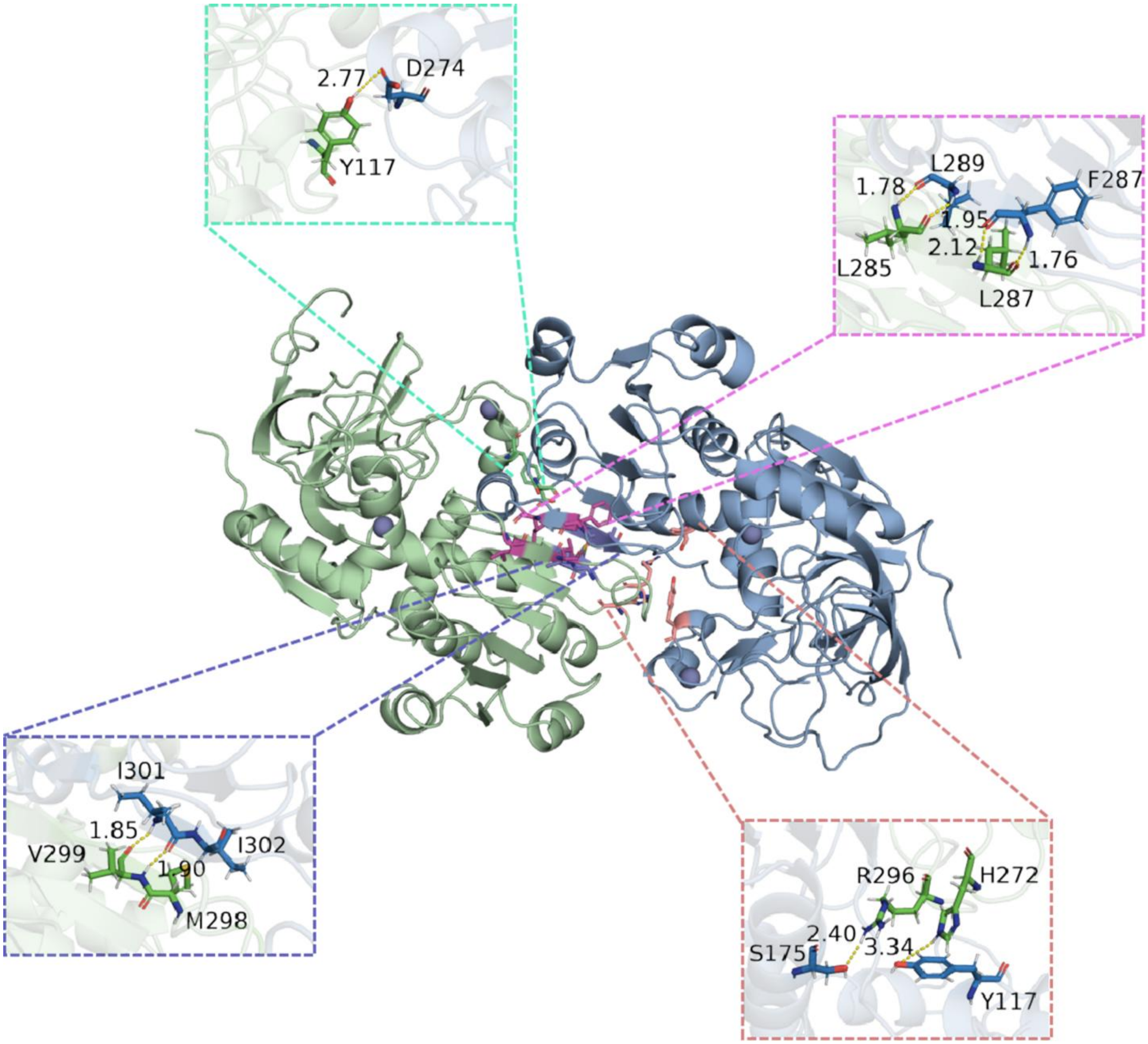

**Fig. S35. Pull-down assays demonstrated interactions between GS and VinBLAST.** Purified 6×His-tagged VinBLAST or its mutants (1 µg), immobilized on Ni-NTA resin, were used to capture the non-tagged GS of interest. Protein-protein interactions were evaluated by testing the GS enzymatic activity of the resin-bound complexes in vitro. VinBLAST and its catalytic mutant (VinBLAST$^{CM}$; C51A and H56A) successfully pulled down GS, while the interaction mutant (VinBLAST$^{IM}$; M298E and V299E) did not. GS activity was assessed by measuring geissoschizine production via LC-MS/MS. Without immobilized VinBLAST, GS lacking a 6×His tag could not bind the resin and showed no detectable activity. Data represent mean ± s.d. of three technical replicates.

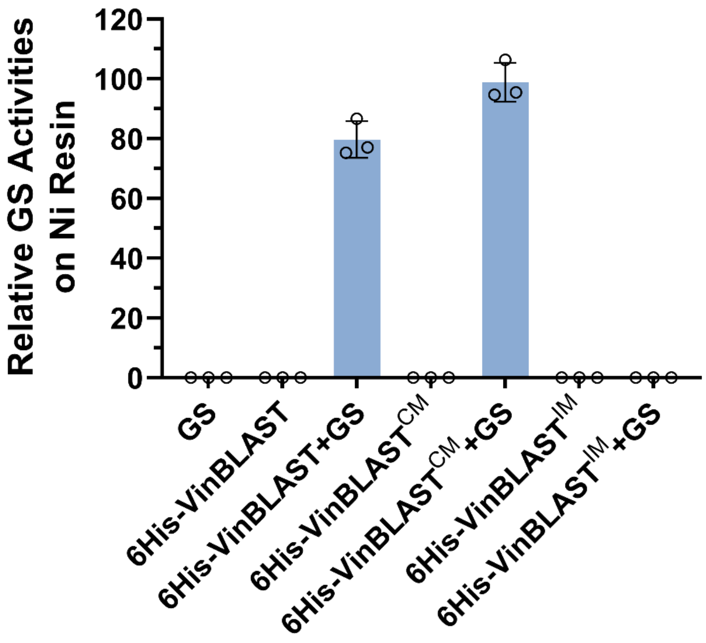

**Fig. S36. Stable binding conformations of various strictosidine aglycones in the active site of VinBLAST-GS heterodimer.** The atoms of substrates involved in the corresponding reaction are highlighted in red circle. **(A)** Binding conformations of 4,21-dehydrogeissoschizine in VinBLAST-GS heterodimer, the active site is highlighted in red rectangle. **(B)** 4,21-dehydrogeissoschizine in the active site of VinBLAST-GS heterodimer; **(C)** 4,21-dehydrodemethylcorynantheine in the active site of VinBLAST-GS heterodimer; **(D)** cathenamine in the active site of VinBLAST-GS heterodimer; **(E)** 19-epicathenamine in the active site of VinBLAST-GS heterodimer; **(F)** cathenamine iminium in the active site of VinBLAST-GS heterodimer; **(G)** 19-epicathenamine iminium in the active site of VinBLAST-GS heterodimer. Only 4,21-dehydrogeissoschizine adopts a conformation as near-attack conformation; in the complexes of substrates 4,21-dehydrodemethylcorynantheine, cathenamine iminium, and 19-epicathenamine iminium, the interatomic distances of key atoms exceed 3 Å, thus hindering the occurrence of the reaction; as for the systems containing cathenamine and 19-epicathenamine, the reactive atoms of the substrates are positioned distant from the protein's active site, thereby impeding the respective reactions.

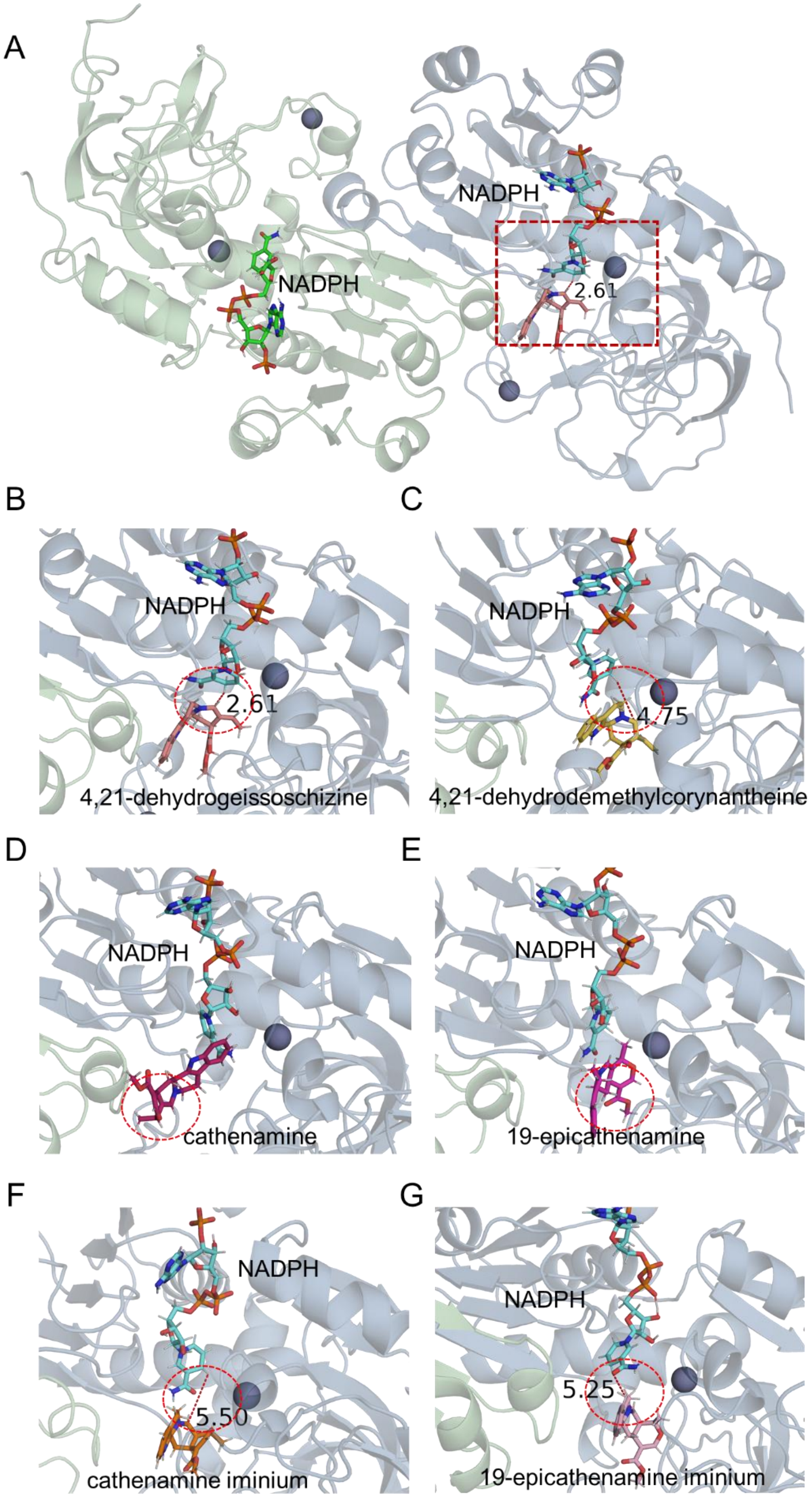
A
NADPH
NADPH
2.61
B
NADPH
2.61
4,21-dehydrogeissoschizine
C
NADPH
4.75
4,21-dehydrodemethylcorynantheine
D
NADPH
cathenamine
E
NADPH
19-epicathenamine
F
NADPH
5.50
cathenamine iminium
G
NADPH
5.25
19-epicathenamine iminium

**Fig. S37. Near-attack conformation (NAC) analysis. (A)** Definition of NAC, conformations where the interatomic distance between reactive carbon and hydrogen atoms (marked as red) ≤ 3.0 Å were designated as NACs. **(B)** Root Mean Square Deviations (RMSDs) of substrate (4,21-dehydrogeissoschizine) during 100 ns MD simulations. **(C)** The interatomic distance between reactive carbon and hydrogen atoms during the MD simulations from 40 to 100 ns. Cyan: substrate in the active site of VinBLAST-GS heterodimer; Blue: substrate in active site 1 of GS-GS homodimer; Purple: substrate in active site 2 of GS-GS homodimer. **(D, E, F)** The most NAC-conformation-representative snapshots were extracted from the 40-100 ns trajectories. Active site of VinBLAST-GS heterodimer **(D)**; Active site 1 of GS-GS homodimer **(E)**; Active site 2 of GS-GS homodimer **(F)**. **(G)** In silico NAC frequencies at the GS active site in VinBLAST-GS heterodimer and in the two monomers of GS-GS homodimers (active site 1 and active site 2).

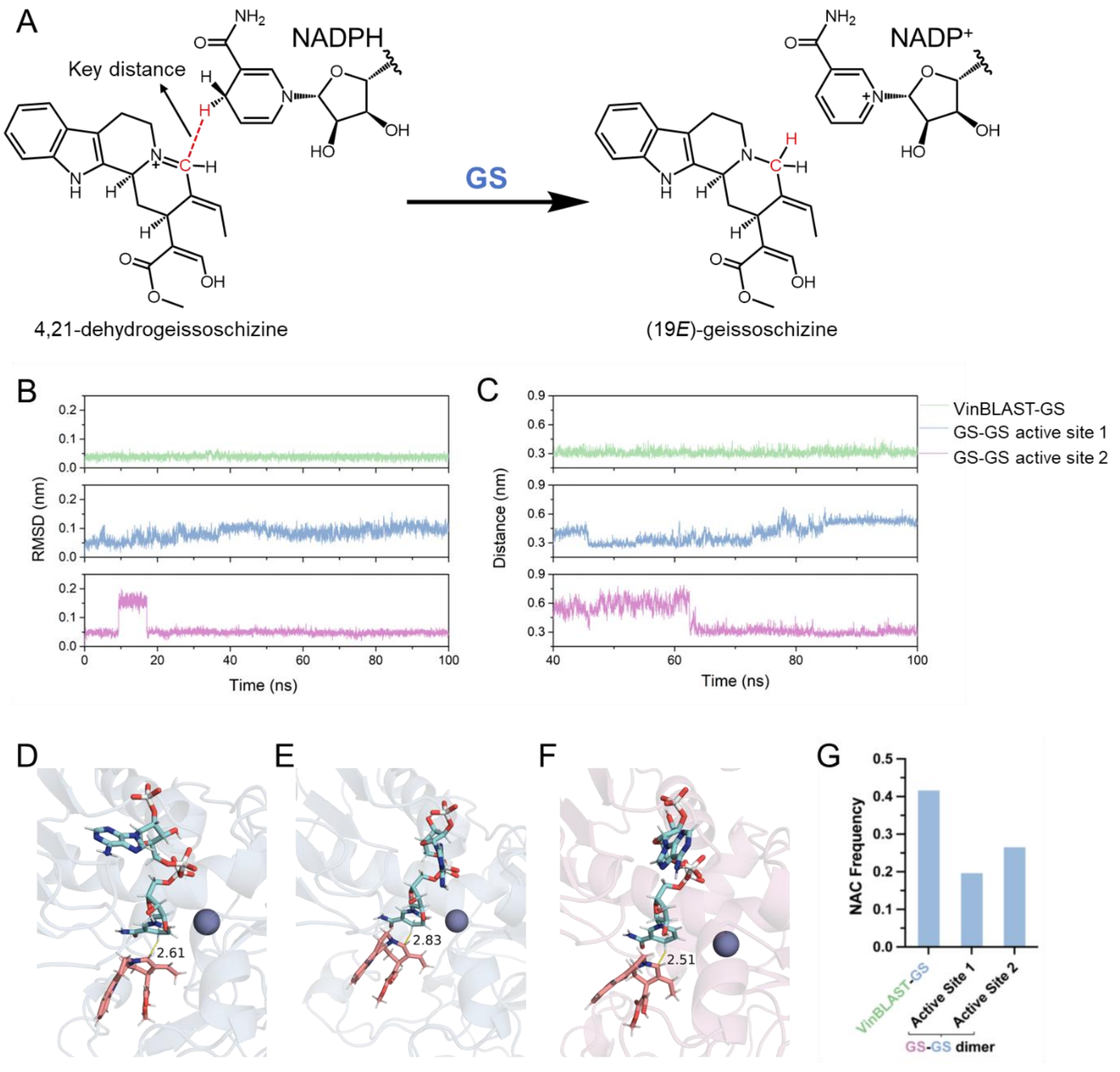

**Fig. S38. MD simulations of HYS-GS heterodimer. (A)** Root Mean Square Deviations (RMSDs) of substrate (4,21-dehydrogeissoschizine) during 100 ns MD simulations. **(B)** The interatomic distance between reactive carbon and hydrogen atoms during the MD simulations from 40 to 100 ns. Definition of NAC is same as VinBLAST-GS heterodimer. **(C)** Schematic diagram of the formation of a tunnel by HYS-GS. Tunnel is shown as pink surface. **(D)** The segments in HYS and VinBLAST participating in substrate tunnel formation. HYS: PAPALLMGGK (284-293), shown in orange sticks; VinBLAST: HSAPLLMGRK (288-297), shown in cyan sticks. The different residues of the segments are highlighted. The lower NAC frequency compared to GS homodimer and VinBLAST-GS heterodimer suggests that it is more difficult for the substrate to adopt the transition-state geometry in HYS-GS heterodimer. Furthermore, the channels in HYS-GS are excessively narrow relative to those in VinBLAST-GS, impeding substrate entry and exit.

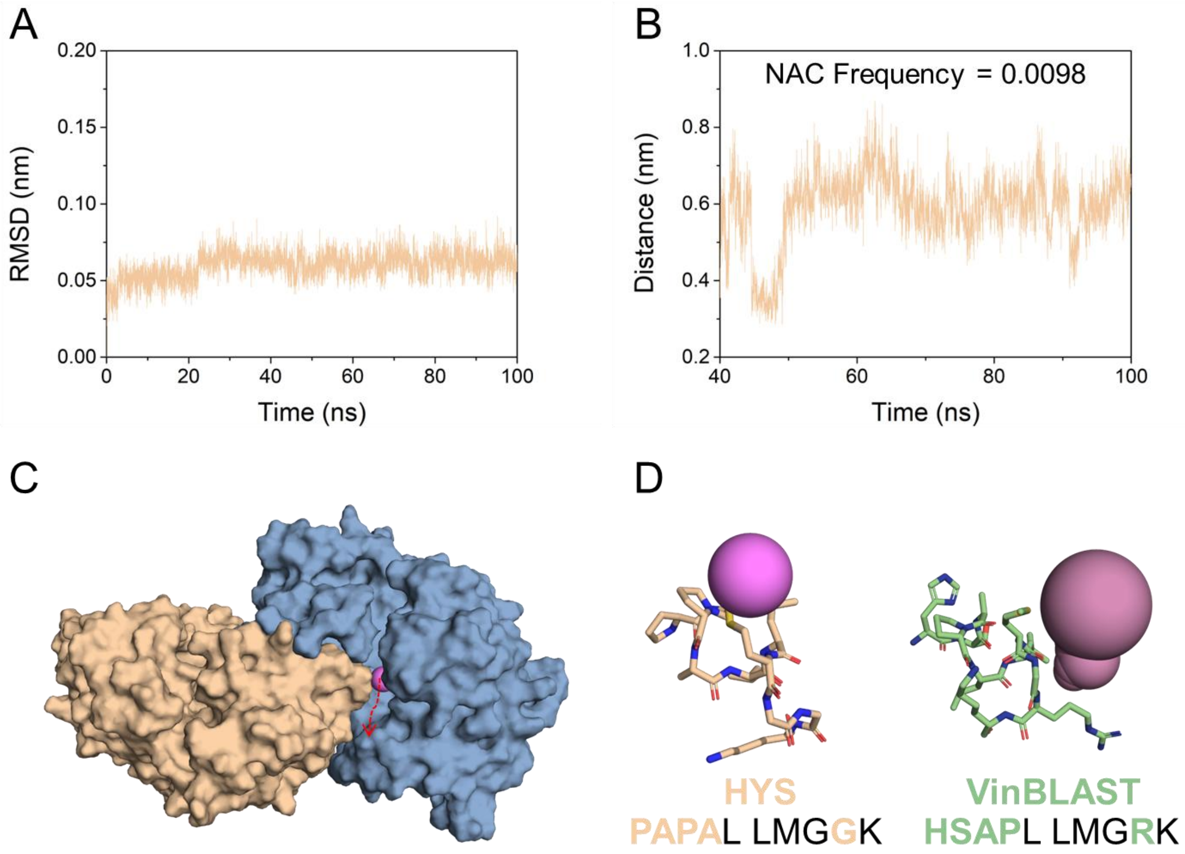

**Fig. S39. GS$^{TM}$ (GS$^{P290H, A291S, I295L}$) improved *de novo* GME and vindoline production in engineered *S. cerevisiae* strains.** These strains contain modifications in GS$^{TM}$ (GS tunnel mutant; P290H, A291S, and I295L), tethering SGD and GS$^{TM}$ with self-interacting peptide tags RIDD/RIAD and SpyTag/SpyCatcher, VinBLAST, and GS copy numbers. The parental strains were AJM7-ΔHYS (*4*) for GME-producing strains **(A)** and VIN13 (*4*) for vindoline-producing strains **(B)**. The corresponding genomic modifications for each strain were specified underneath. The strain genotypes were listed in table S1. The results represent the mean ± s.d. from three biological replicates.

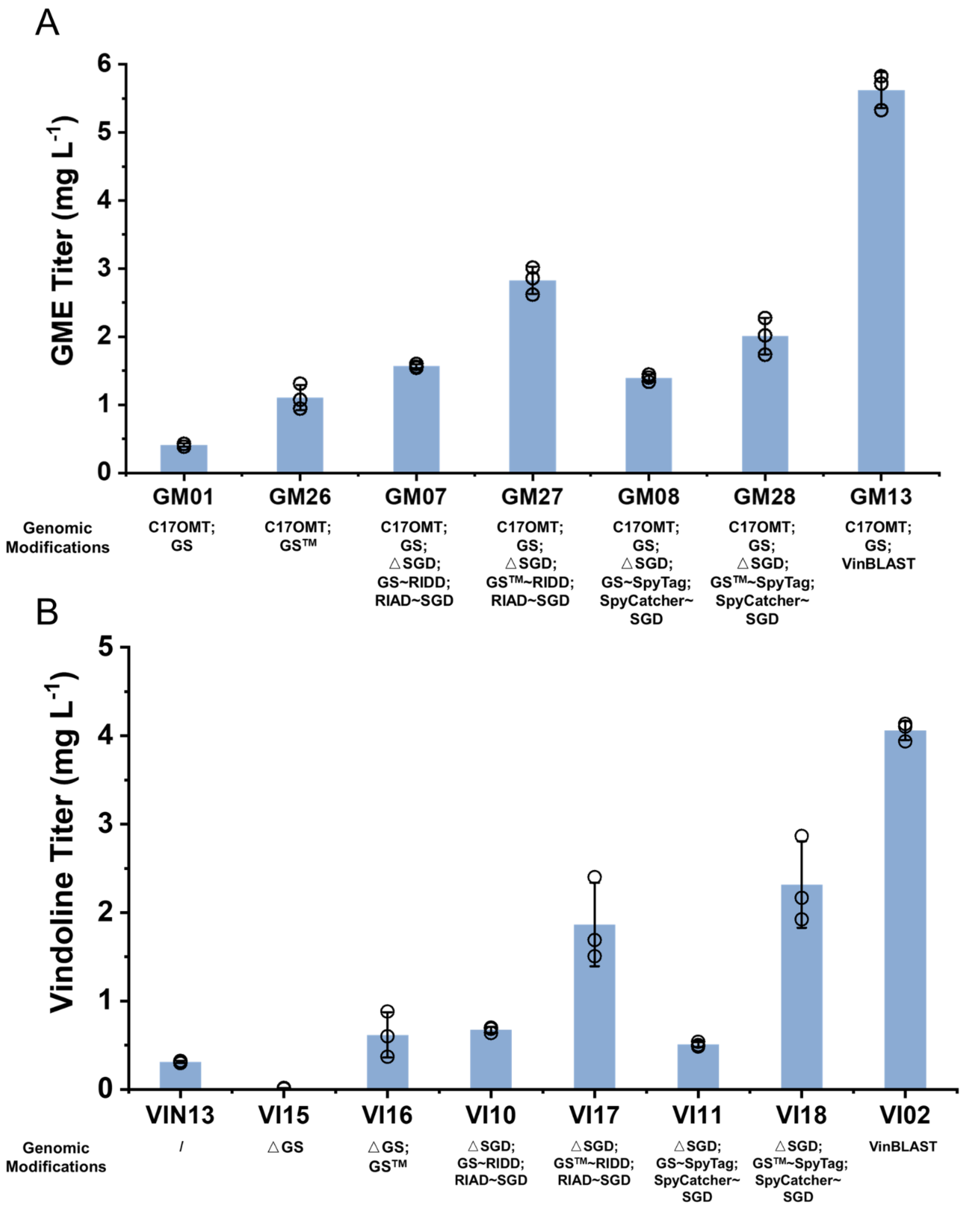

**Fig. S40. Phylogenetic analysis of VinBLAST and GS with diverse CADs and CAD-like enzymes across embryophytes.** A comprehensive phylogenetic tree showing the five CAD clades based on de Vries assignment (*25*). VinBLAST homologues (magenta) and GS homologues (blue) belong to the CAD clade III. CAD clade III comprises only angiosperms, including early diverging *Amborella trichopoda*, monocots, and eudicots. Members with demonstrated in vitro CAD activities are marked with blue dots, while those experimentally shown to lack CAD activity marked with yellow dots. Proteins implicated in specialized metabolism are marked with purple dots. Unlike the CAD clade I, which contains tracheophyte members with genetically and biochemically supported roles in lignin biosynthesis (orange), the lignin-related functions of CADs in other clades have not been explicitly demonstrated, despite some in vitro evidence. Clade I CADs with both biochemical and genetic evidence supporting their involvement in lignin biosynthesis are highlighted in orange. Clade I CAD homologues from MIA-producing species are highlighted in green. In particular, CrC1150 (GenBank: KU865327) exhibited substantially higher CAD activity than VinBLAST (CAD2) in enzymatic assays (fig. S12). The CAD protein sequences (*25*) combined with VinBLAST and GS homologues were aligned by Muscle. Evolutionary analyses were conducted in MEGA11. The evolutionary history was inferred by using the Maximum Likelihood method and JTT matrix-based model. The tree with the highest log likelihood is shown. The percentage of trees in which the associated taxa clustered together is shown next to the branches (100 bootstrap replicates). Initial tree(s) for the heuristic search were obtained automatically by applying Neighbor-Join and BioNJ algorithms to a matrix of pairwise distances estimated using the JTT model, and then selecting the topology with superior log likelihood value.

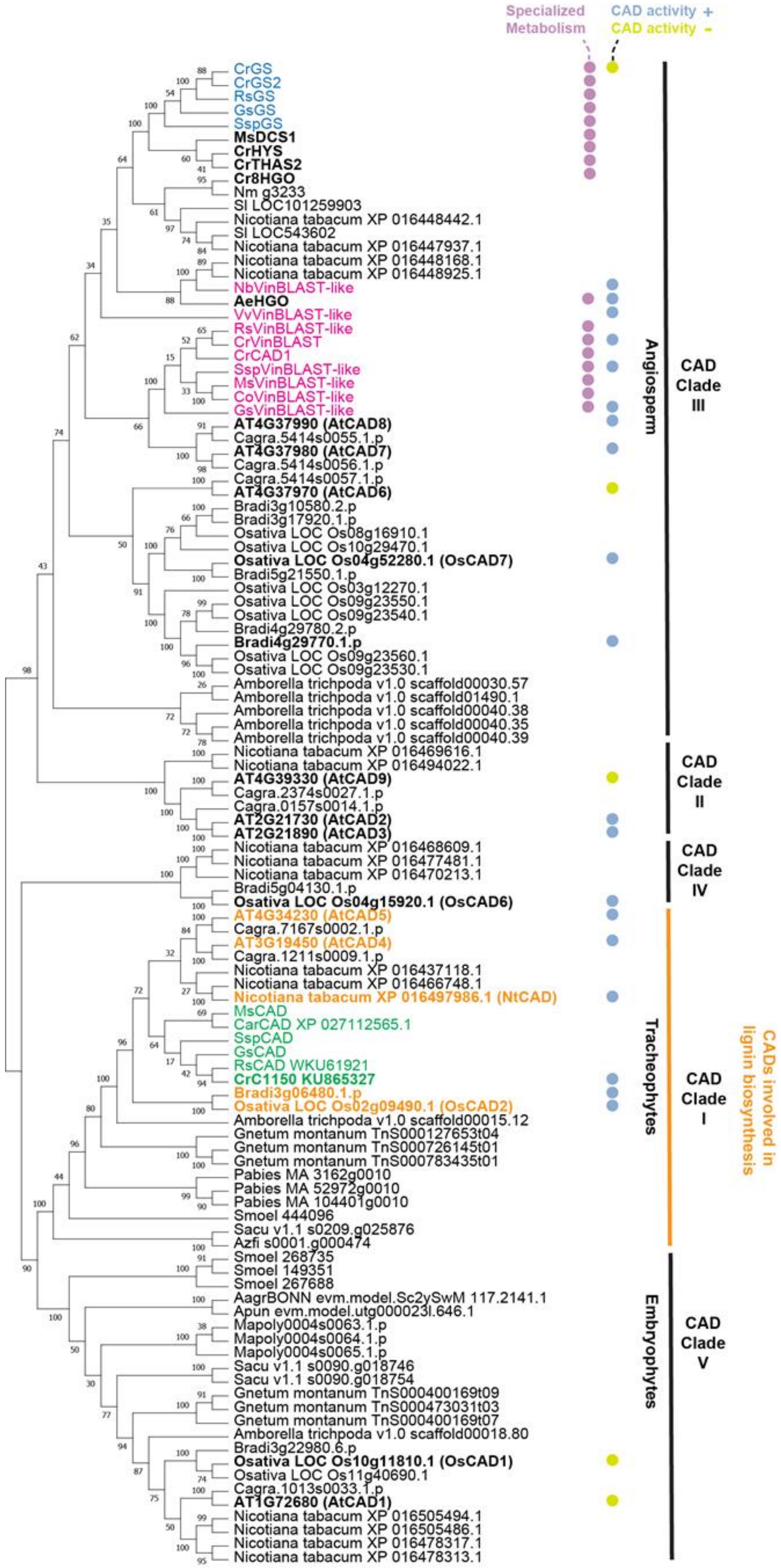
Specialized Metabolism
CAD activity +
CAD activity -
CrGS
CrGS2
RsGS
GsGS
SspGS
MsDCS1
CrHYS
CrTHAS2
Cr8HGO
Nm g3233
Sl LOC101259903
Nicotiana tabacum XP 016448442.1
Sl LOC543602
Nicotiana tabacum XP 016447937.1
Nicotiana tabacum XP 016448168.1
Nicotiana tabacum XP 016448925.1
NbVinBLAST-like
AeHGO
VvVinBLAST-like
RsVinBLAST-like
CrVinBLAST
CrCAD1
SspVinBLAST-like
MsVinBLAST-like
CoVinBLAST-like
GsVinBLAST-like
AT4G37990 (AtCAD8)
Cagra.5414s0055.1.p
AT4G37980 (AtCAD7)
Cagra.5414s0056.1.p
Cagra.5414s0057.1.p
AT4G37970 (AtCAD6)
Bradi3g10580.2.p
Bradi3g17920.1.p
Osativa LOC Os08g16910.1
Osativa LOC Os10g29470.1
Osativa LOC Os04g52280.1 (OsCAD7)
Bradi5g21550.1.p
Osativa LOC Os03g12270.1
Osativa LOC Os09g23550.1
Osativa LOC Os09g23540.1
Bradi4g29780.2.p
Bradi4g29770.1.p
Osativa LOC Os09g23560.1
Osativa LOC Os09g23530.1
Amborella trichpoda v1.0 scaffold00030.57
Amborella trichpoda v1.0 scaffold01490.1
Amborella trichpoda v1.0 scaffold00040.38
Amborella trichpoda v1.0 scaffold00040.35
Amborella trichpoda v1.0 scaffold00040.39
Nicotiana tabacum XP 016469616.1
Nicotiana tabacum XP 016494022.1
AT4G39330 (AtCAD9)
Cagra.2374s0027.1.p
Cagra.0157s0014.1.p
AT2G21730 (AtCAD2)
AT2G21890 (AtCAD3)
Nicotiana tabacum XP 016468609.1
Nicotiana tabacum XP 016477481.1
Nicotiana tabacum XP 016470213.1
Bradi5g04130.1.p
Osativa LOC Os04g15920.1 (OsCAD6)
AT4G34230 (AtCAD5)
Cagra.7167s0002.1.p
AT3G19450 (AtCAD4)
Cagra.1211s0009.1.p
Nicotiana tabacum XP 016437118.1
Nicotiana tabacum XP 016466748.1
Nicotiana tabacum XP 016497986.1 (NtCAD)
MsCAD
CarCAD XP 027112565.1
SspCAD
GsCAD
RsCAD WKU61921
CrC1150 KU865327
Bradi3g06480.1.p
Osativa LOC Os02g09490.1 (OsCAD2)
Amborella trichpoda v1.0 scaffold00015.12
Gnetum montanum TnS000127653t04
Gnetum montanum TnS000726145t01
Gnetum montanum TnS000783435t01
Pabies MA 3162g0010
Pabies MA 52972g0010
Pabies MA 104401g0010
Smoel 444096
Sacu v1.1 s0209.g025876
Azfi s0001.g000474
Smoel 268735
Smoel 149351
Smoel 267688
AagrBONN evm.model.Sc2ySwM 117.2141.1
Apun evm.model.utg000023l.646.1
Mapoly0004s0063.1.p
Mapoly0004s0064.1.p
Mapoly0004s0065.1.p
Sacu v1.1 s0090.g018746
Sacu v1.1 s0090.g018754
Gnetum montanum TnS000400169t09
Gnetum montanum TnS000473031t03
Gnetum montanum TnS000400169t07
Amborella trichpoda v1.0 scaffold00018.80
Bradi3g22980.6.p
Osativa LOC Os10g11810.1 (OsCAD1)
Osativa LOC Os11g40690.1
Cagra.1013s0033.1.p
AT1G72680 (AtCAD1)
Nicotiana tabacum XP 016505494.1
Nicotiana tabacum XP 016505486.1
Nicotiana tabacum XP 016478317.1
Nicotiana tabacum XP 016478313.1
Angiosperm
Tracheophytes
Embryophytes
CAD Clade III
CAD Clade II
CAD Clade IV
CAD Clade I
CAD Clade V
CADs involved in lignin biosynthesis

**Fig. S41. Both *C. roseus* (*Cr*) and *N. benthamiana* (*Nb*) VinBLAST proteins interacted with CrSGD and CrGS in *N. benthamiana* leaf epidermis.** The *N*-terminal fragment (mVenus$^{N}$) and the *C*-terminal fragment (mVenus$^{C}$) of mVenus were individually fused to proteins of interest and transiently expressed in *N. benthamiana* leaf epidermis via *Agrobacterium*-mediated infiltration. Yellow fluorescence is generated when the two mVenus fragments are brought into proximity by interaction of the fused proteins. For both VinBLASTs and GS, mVenus tags were fused to the *N*-termini. For SGD, fusions were made at both *N*- and *C*-termini depending on the construction. Right panels show overlay with brightfield background. **(A)** GFP tagged CrVinBLAST. While CrVinBLAST was predominantly cytosolic, its nuclear localization likely resulted from overexpression and its small size permitting passive nuclear entry. **(B)** Co-expression of BiFC-tagged CrVinBLAST resulted in fluorescence in both the cytosol and nucleus, confirming homodimer formation. **(C, D)** CrGS interacted with both CrVinBLAST and NbVinBLAST in the cytosol and nucleus, indicating conserved interaction capacity. **(E)** Similar nuclear and cytosolic patterns were observed when CrGS and NbVinBLAST were co-expressed in the presence of non-tagged CrSGD. **(F)** CrSGD self-interacted in the nucleus, consistent with its known nuclear localization and oligomeric nature. **(G, H)** NbVinBLAST, despite being derived from a species that does not synthesize iridoids or MIAs, and CrVinBLAST both interacted with CrSGD in the nucleus. **(I, J)** Co-expression of CrGS and CrSGD in *N. benthamiana* showed interaction even in the absence of exogenous VinBLAST, likely due to endogenous NbVinBLAST or other native CADs that could mediate interaction. These results suggest that *N. benthamiana* cells and other plant cells inherently express proteins capable of bridging CrGS and CrSGD, confounding efforts to isolate GS-SGD interactions in this host. Nevertheless, our results demonstrated that CrSGD and CrGS could interact with CADs from both *C. roseus* and *N. benthamiana*, pointing to a conserved mechanism underlying these protein-protein interactions.

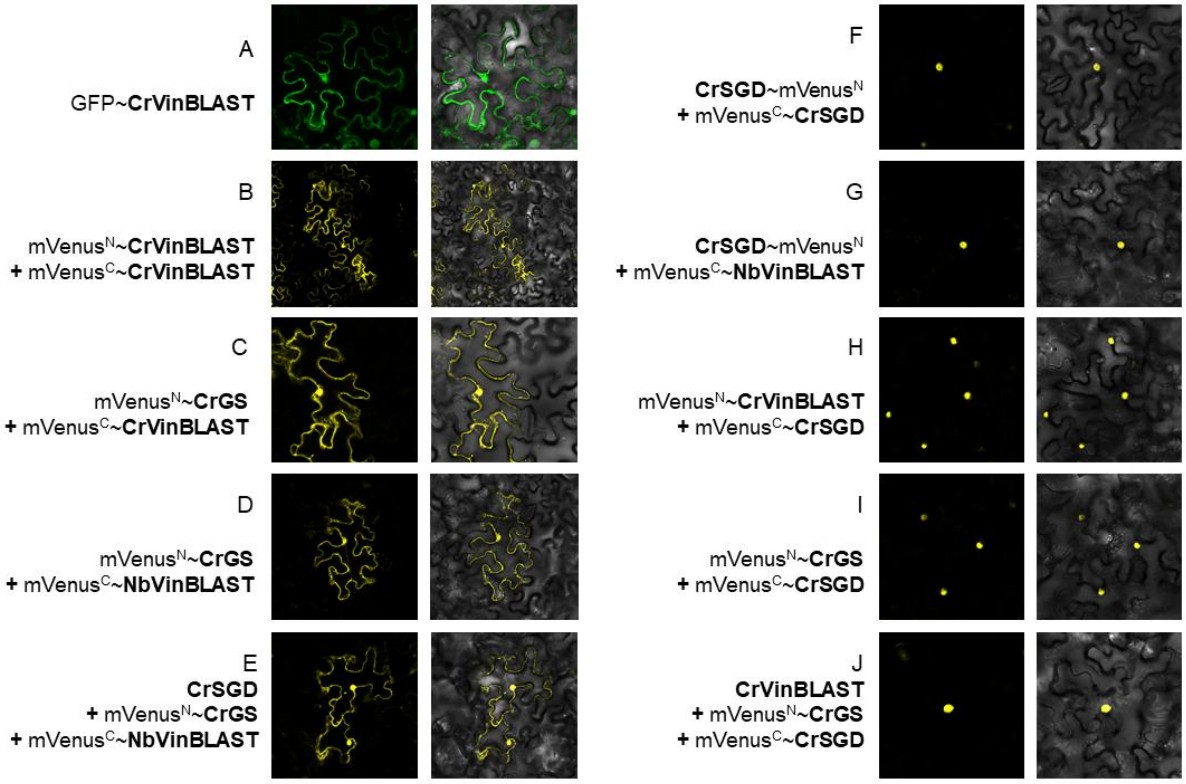

**Fig. S42. The BiFC experiments in yeast demonstrated protein-protein interactions among CAD1, GS, and SGD.** The *N*-terminal fragment (mVenus$^{N}$) and the *C*-terminal fragment (mVenus$^{C}$) of mVenus were individually fused to proteins of interest. Yellow fluorescence is generated when the two mVenus fragments are brought into proximity by interaction of the fused proteins. Considering that SGD has a nuclear localization signal (NLS) at the *C*-terminus, the mVenus fragments were fused to the *N*-terminus of SGD. For both CAD1 and GS, the mVenus fragments were fused to their *C*-termini. BiFC assays showed that CAD1 interacts with GS in the cytosol **(A)** and with SGD in the nucleus **(B)**, and CAD1 homodimerization was also observed in the cytosol **(C)**. However, in contrast to VinBLAST, the presence of CAD1 did not enable detectable interaction between SGD and GS **(D)**, indicating that CAD1 is not as effective as VinBLAST to serve as a scaffold to bridge SGD and GS. Nab2 tagged with mCherry at the *C*-terminus functioned as a nuclear marker. All proteins were expressed in the wild-type CEN.PK2-1C yeast strains. The images were acquired using a confocal laser scanning microscope (Olympus FV3000) at 600× magnification, with a 5 μm scale bar displayed in the upper left corner.

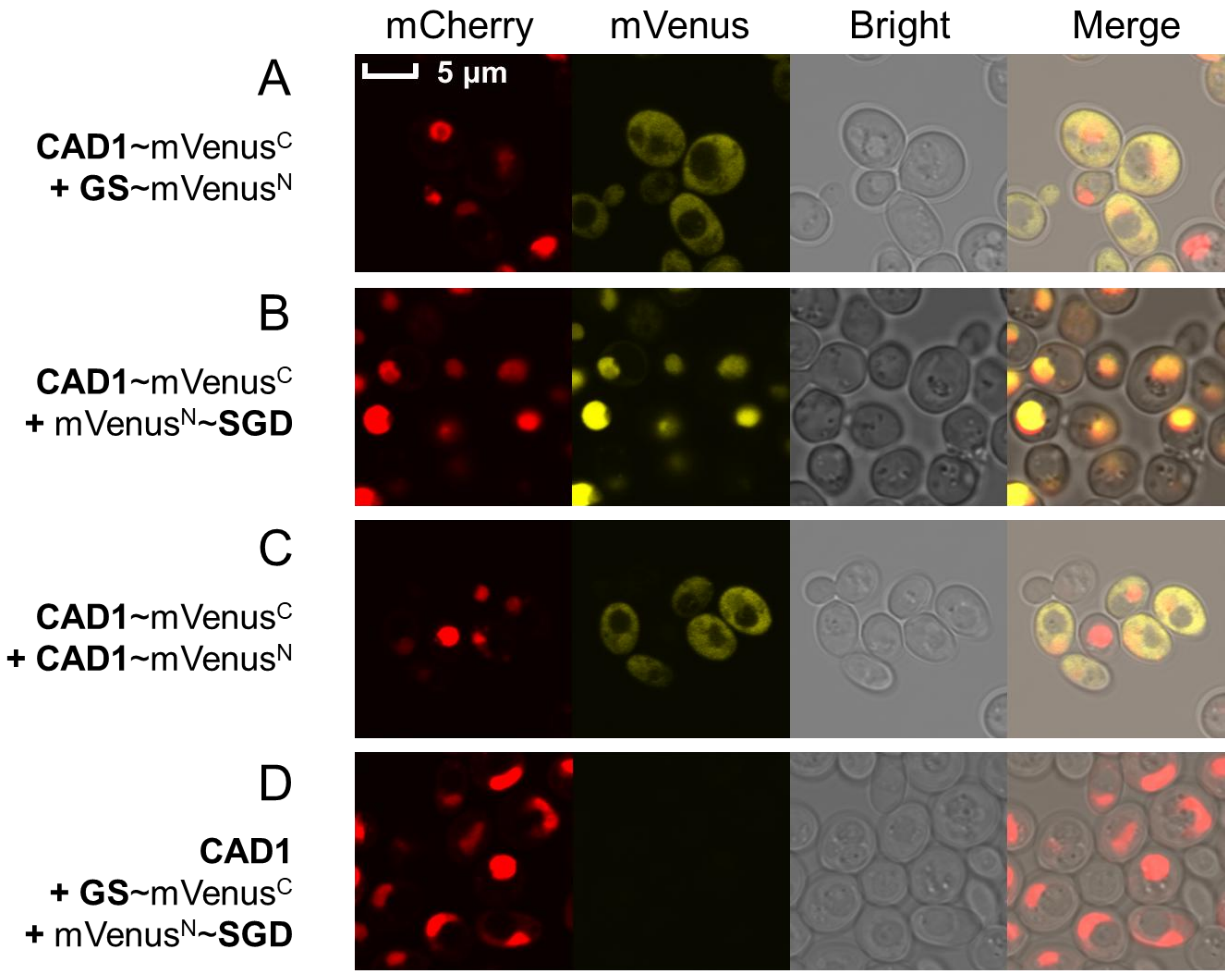

**Fig. S43. The Y2H assays to investigate CAD1, GS, and SGD interactions.** The Y2H results showed that CAD1 interacted with both GS and itself. Regarding the interaction with SGD, Y2H revealed a very weak interaction between CAD1 and SGD, which is consistent with the generally weak Y2H signals observed for SGD with other proteins in our study (fig. S14). We interpret these findings to suggest that Y2H lacks sufficient sensitivity to reliably detect the very weak protein-protein interactions involving SGD. The test proteins were fused to the GAL4 transcription activation domain (AD) and GAL4 DNA-binding domain (BD), respectively, at the *N*-terminus, and transformed into the Y2H-GOLD host strain with histidine deficiency. Upon protein-protein interactions, the *GAL* promoter is transcriptionally activated, enabling the yeast strains to express the *HIS3* gene and therefore grow on the histidine-deficient agar plates. Weakly interacting proteins failed to activate the *GAL* promoter and thus the corresponding strains could not grow. 3-AT (3-Amino-1,2,4-triazole) is a common competitive protein inhibitor used in Y2H experiments to suppress the leaky expression of *HIS3* and decrease the background noise. The yeast cultures, diluted at different ratios, were spotted onto SED amino acid-deficient plates containing no 3-AT, 5 mM 3-AT, and 20 mM 3-AT, respectively. The growth of the yeast clones was observed after incubation at 30°C for 96 h.

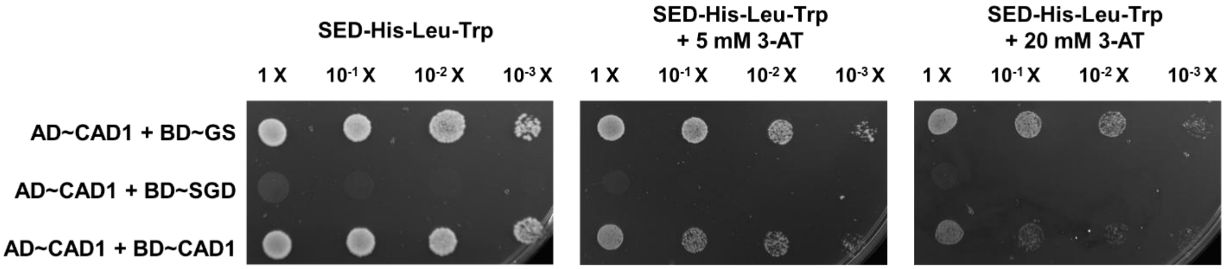

**Fig. S44. MIA titers achieved by CrCAD1 in engineered *S. cerevisiae* strains producing GME, catharanthine, and vindoline.** Negative Control (NC) refers to the base strains harboring the full biosynthetic pathways with a single *GS* copy, which were further engineered to include an extra copy of *VinBLAST* and *CrCAD1*, respectively. In all three strains, the yield improvements conferred by CAD1 were only 27.4%, 7.85%, and 21.3% of those achieved by VinBLAST, respectively. These results indicate that CAD1 has a very limited capacity to enhance MIA production compared to VinBLAST. Yellow, GME; purple, catharanthine; and blue, vindoline. The results represent the mean ± s.d. of three biological replicates.

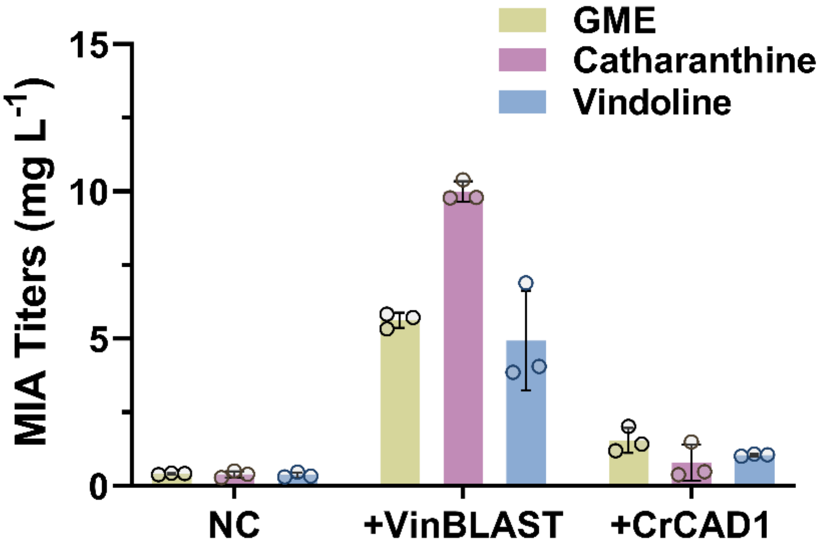

**Table S1.** Yeast strains constructed in this study.

| Strain | Titer (mg L$^{-1}$) | Genotype |
|---|---|---|
| **GME-producing Strains** | | |
| GM01 | 0.41 | AJM7-ΔHYS-IntG4::*C17OMT*-IntG26::*GS* |
| GM02 | 0.01 | GM01-Δ*SGD* |
| GM03 | 0.42 | GM02-IntG17::*SGD* |
| GM04 | 0.29 | GM02-IntG17::*tSGD* |
| GM05 | 0.83 | GM02-IntG16::*GS*-IntG17::*SGD* |
| GM06 | 0.49 | GM02-IntG17::*GS~SGD* |
| GM07 | 1.57 | GM02-IntG16::*GS~RIDD*-IntG17::*RIAD~SGD* |
| GM08 | 1.39 | GM02-IntG16::*GS~SpyTag*-IntG17::*SpyCatcher~SGD* |
| GM09 | 0.71 | GM02-IntG16::*GS*-IntG17::*tSGD* |
| GM10 | 0.57 | GM02-IntG16::*GS~RIDD*-IntG17::*RIAD~tSGD* |
| GM11 | 0.89 | GM02-IntG16::*GS~SpyTag*-IntG17::*SpyCatcher~tSGD* |
| GM12 | 0.41 | AJM7-ΔHYS-IntG4::*C17OMT*-IntG26::*VinBLAST* |
| GM13 | 5.62 | GM01-IntG11::*VinBLAST* |
| GM14 | 1.54 | GM01-IntG11::*CAD1* |
| GM15 | 0.76 | GM01-IntG11::*CAD3* |
| GM16 | 0.31 | GM01-IntG11::*CAD5* |
| GM17 | 0.05 | AJM7-ΔHYS-IntG4::*C17OMT*-IntG26::*VinBLAST*$^{CM}$ (*VinBLAST*$^{C51A,H56A}$) |
| GM18 | 5.56 | GM01-IntG11::*VinBLAST*$^{CM}$ (*VinBLAST*$^{C51A,H56A}$) |
| GM19 | 1.05 | GM01-IntG13::*VinBLAST*$^{IM}$ (*VinBLAST*$^{M298E,V299E}$) |
| GM20 | 6.02 | GM13-IntG21::*GS* |
| GM21 | 6.78 | GM13-IntG22::*VinBLAST* |
| GM22 | 7.64 | GM13-IntG21::*GS*-G22::*VinBLAST* |
| GM23 | 3.32 | GM01-IntG11::*CoVinBLAST* |
| GM24 | 7.79 | GM01-IntG11::*RsVinBLAST* |
| GM25 | 3.65 | GM01-IntG11::*MsVinBLAST* |
| GM26 | 1.11 | AJM7-ΔHYS-IntG4::*C17OMT*-IntG13::*GS*$^{TM}$ (*GS*$^{P290H,A291S,I295L}$) |
| GM27 | 2.83 | GM02-IntG16::*GS*$^{TM}$ (*GS*$^{P290H,A291S,I295L}$)~*RIDD*-IntG17::*RIAD~SGD* |
| GM28 | 2.01 | GM02-IntG16::*GS*$^{TM}$ (*GS*$^{P290H,A291S,I295L}$)~*SpyTag*-IntG17::*SpyCatcher~SGD* |
| GM29 | 0.15 | AJM7-ΔHYS-IntG4::*C17OMT*-IntG13::*GS*$^{IM}$ (*GS*$^{I301E}$) |
| GM30 | 0.09 | GM29-IntG14::*VinBLAST* |
| GM31 | 0.15 | GM29-IntG14::*VinBLAST*$^{CM}$ (*VinBLAST*$^{C51A,H56A}$) |
| GM32 | 0.06 | GM29-IntG14::*VinBLAST*$^{IM}$ (*VinBLAST*$^{M298E,V299E}$) |
| **Catharanthine-producing Strains** | | |
| CA01 | 0.77 | CAT3-IntG13::*GS* |
| CA02 | 9.93 | CAT3-IntG13::*VinBLAST* |
| CA03 | 9.67 | CAT3-IntG13::*VinBLAST*$^{CM}$ (*VinBLAST*$^{C51A,H56A}$) |
| CA04 | 1.04 | CAT3-IntG13::*VinBLAST*$^{IM}$ (*VinBLAST*$^{M298E,V299E}$) |
| CA05 | 3.61 | CAT3-IntG13::*CoVinBLAST* |
| CA06 | 12.92 | CAT3-IntG13::*RsVinBLAST* |
| CA07 | 5.68 | CAT3-IntG13::*MsVinBLAST* |
| **Vindoline-producing Strains** | | |
| VI01 | 0.37 | VIN13-IntG13::*GS* |
| VI02 | 4.06 | VIN13-IntG11::*VinBLAST* |
| VI03 | 3.98 | VIN13-IntG13::*VinBLAST*$^{CM}$ (*VinBLAST*$^{C51A,H56A}$) |
| VI04 | 0.52 | VIN13-IntG13::*VinBLAST*$^{IM}$ (*VinBLAST*$^{M298E,V299E}$) |
| VI05 | 0.00 | VIN13-Δ*SGD* |
| VI06 | 0.29 | VI05-IntG4::*SGD* |
| VI07 | 0.17 | VI05-IntG4::*tSGD* |

| Strain | Titer (mg $L^{-1}$) | Genotype |
|---|---|---|
| VI08 | 0.46 | VI05-IntG4::*SGD*-IntG3::*GS* |
| VI09 | 0.48 | VI05-IntG4::*GS~SGD* |
| VI10 | 0.67 | VI05-IntG4::*GS~RIDD*-IntG3::*RIAD~SGD* |
| VI11 | 0.51 | VI05-IntG4::*GS~SpyTag*-IntG3::*SpyCatcher~SGD* |
| VI12 | 0.50 | VIN13-IntG11::*CoVinBLAST* |
| VI13 | 7.51 | VIN13-IntG11::*RsVinBLAST* |
| VI14 | 0.81 | VIN13-IntG11::*MsVinBLAST* |
| VI15 | 0.01 | VIN13-△*GS* |
| VI16 | 0.62 | VI15-IntG13::$GS^{TM}$ ($GS^{P290H,A291S,I295L}$) |
| VI17 | 1.87 | VI05-IntG4::$GS^{TM}$ ($GS^{P290H,A291S,I295L}$)*~RIDD*-IntG3::*RIAD~SGD* |
| VI18 | 2.32 | VI05-IntG4::$GS^{TM}$ ($GS^{P290H,A291S,I295L}$)*~SpyTag*-IntG3::*SpyCatcher~SGD* |
| VI19 | 0.01 | VI15-IntG13::$GS^{IM}$ ($GS^{I301E}$) |
| VI20 | 0.01 | VI19-IntG14::*VinBLAST* |
| VI21 | 0.02 | VI19-IntG14::$VinBLAST^{CM}$ ($VinBLAST^{C51A,H56A}$) |
| VI22 | 0.02 | VI19-IntG14::$VinBLAST^{IM}$ ($VinBLAST^{M298E,V299E}$) |

**Notes:**

1. The strains AJM7-ΔHYS, CAT3, and VIN13 were constructed in our previous study (*4*).
2. tSGD means the *C*-terminal NLS-truncated version of SGD.
3. VinBLAST without species annotation represents CrVinBLAST, while VinBLAST homologues from other species have been explicitly labeled.
4. The symbol "Δ" represents gene knockout. The symbol "::" represents gene integration. The symbol "~" represents gene fusion expression.

**Table S2.** GS and VinBLAST kinetics parameters. $GS^{TM}$ represents the tunnel mutant $GS^{P290H,A291S,I295L}$, $VinBLAST^{CM}$ represents the catalytic mutant $VinBLAST^{C51A,H56A}$, $VinBLAST^{IM}$ represents the interaction mutant $VinBLAST^{M298E,V299E}$, and $VinBLAST^{TM}$ represents the tunnel mutant $VinBLAST^{H288P,S289A,L293I}$.

| | $K_m$ (µM) | $K_m$ 95% CI* (µM) | $V_{max}$ (nmol/mg GS/min) | $V_{max}$ 95% CI (nmol/mg GS/min) |
|---|---|---|---|---|
| GS homodimer | 0.495 | 0.385-0.633 | 2.84 | 2.68-3.00 |
| $GS^{TM}$ homodimer | 0.919 | 0.647-1.29 | 14.9 | 13.6-16.3 |
| VinBLAST-GS | 0.512 | 0.411-0.637 | 68.9 | 65.5-72.5 |
| $VinBLAST^{CM}$-GS | 0.617 | 0.510-0.744 | 86.1 | 82.4-90.0 |
| $VinBLAST^{IM}$-GS | 0.844 | 0.617-1.15 | 16.5 | 15.3-17.8 |
| $VinBLAST^{TM}$-GS | 0.517 | 0.355-0.734 | 52.4 | 48.0-57.0 |

* Confidence interval. Data were fitted using the Michaelis-Menten model and non-liner regression in GraphPad Prism version 10.4.2.

**Table S3.** List of genes and their corresponding amino acid sequences used in this study.

>CrGS_W8JWW7.1
MAGETTKLDLSVKAVGWGAADASGVLQPIKFYRRVPGERDVKIRVLYSGVCNFDMEMVRNKWGFTRYP
YVFGHETAGEVVEVGSKVEKFKVGDKVAVGCMVGSCGQCYNCQSGMENYCPEPNMADGSVYREQGERS
YGGCSNVMVVDEKFVLRWPENLPQDKGVALLCAGVVVYSPMKHLGLDKPGKHIGVFGLGGLGSVAVKFI
KAFGGKATVISTSRRKEKEAIEEHGADAFVVNTDSEQLKALAGTMDGVVDTTPGGRTPMSLMLNLLKFDG
AVMLVGAPESLFELPAAPLIMGRKKIIGSSTGGLKEYQEMLDFAAKHNIVCDTEVIGIDYLSTAMERIKNLD
VKYRFAIDIGNTLKFEE*

>GsGS_PV770564
MAAKSPEDMHPVKAIGRIAMDSSGVLTPFKFSRRATGEDDVQIKVLYCGVCNFDTDMITNKWGFTRFPYV
FGHETVGEVSEVGSKVKKFKVGDKVAVGCMVGSCRKCKSCINGMENYCPEPEMADGSVYREEAGRSYG
GCSNIMVVNEHFVIRWPEKLPLNAAPLLCAGIVTYSPLRYFGLDKPGMHIGVVGLGGIGHVCVKFIKAFGA
KATVISSSASKEKEAIEKHGADAFLISSNAEQMQAAAGTMDGVIDTTPGGRTPMALMLNLLKTDGTLVVV
GAPEIPFEVPAHPLIIGRKKVVGSSTGGVKEYQEMVDFSAKHNILPDIEVIPIDYLSTAMERIKKSDVKYRFVI
DIANTLKDD*

>SspGS_PV770565
MVGKPAGEDNSVKGIGWGVLDSSGNLVPIKFTRRATGERDVQIKVLYCGICNFDMEMLRNTWGFTRYPY
VFGHETVGEVTEVGNKVQKFRVGDKVAVGCMVGSCRKCKNCENGMENYCREPEMADGSVYRDDAGRS
YGGCSNIMVVDEDFVLRFPDNYPLEAGAPLLCAGVVTYSPLKHFGLDKAGLHIGIVGLGGMGHIAVKLIKA
FGSKATVISSSPSKEKEAIEKHGADAFLVISNAEQMEAATGTMDGVVDTSPGGRASMAVMLNLLKPHGKLI
VIGAPETPYELPLQPLIMGRKSIAGSSTGGVKEYQEVLDFSAKHNIIPDIEVIPIDYLNTAMERIKKVDVKYRF
VIDIGNTLKYE*

>MsC17OMT_WKU61913.1
MQPQRGRKREREREEMESVQSNSSSSDQFAMKGGDDDFSYTKNSTHQRDAIQATKFFIQESIAEKLDVNKF
CGKAFCVADLGCSVGPNTLIAMQNIVEAVELKFKNRKGFHSPTIPEFQVFFNDHTVNDFNTLFRSLPTGHDK
RYYGVGVPGSFYGRLFPCDSIHIMHTSFSTPFLSQVPKEVMDKNSAAWNKGRIHHNHAKADVLKAYEAQH
AEDIDCFLTARAKELVHGGLLMDVTSFRPDGVPHTHVLTNIGMEVLGYCLMDLAKKGLIDEENVDSYNVP
VYLQSPEELKQAVQRNKYFSIEKMESVPMMIDSDVSAKAQQYSLGMRAVMGDVIREQFGAEIVDKLFDLF
KKKLEEHPNFAKGVVLDMFVLLKRNAED*

>RsSGD_Q8GU20.1
MDNTQAEPLVVAIVPKPNASTEHTNSHLIPVTRSKIVVHRRDFPQDFIFGAGGSAYQCEGAYNEGNRGPSIW
DTFTQRSPAKISDGSNGNQAINCYHMYKEDIKIMKQTGLESYRFSISWSRVLPGGRLAAGVNKDGVKFYHD
FIDELLANGIKPSVTLFHWDLPQALEDEYGGFLSHRIVDDFCEYAEFCFWEFGDKIKYWTTFNEPHTFAVNG
YALGEFAPGRGGKGDEGDPAIEPYVVTHNILLAHKAAVEEYRNKFQKCQEGEIGIVLNSMWMEPLSDVQA
DIDAQKRALDFMLGWFLEPLTTGDYPKSMRELVKGRLPKFSADDSEKLKGCYDFIGMNYYTATYVTNAV
KSNSEKLSYETDDQVTKTFERNQKPIGHALYGGWQHVVPWGLYKLLVYTKETYHVPVLYVTESGMVEEN
KTKILLSEARRDAERTDYHQKHLASVRDAIDDGVNVKGYFVWSFFDNFEWNLGYICRYGIIHVDYKSFERY
PKESAIWYKNFIAGKSTTSPAKRRREEAQVELVKRQKT*

>tRsSGD
MDNTQAEPLVVAIVPKPNASTEHTNSHLIPVTRSKIVVHRRDFPQDFIFGAGGSAYQCEGAYNEGNRGPSIW
DTFTQRSPAKISDGSNGNQAINCYHMYKEDIKIMKQTGLESYRFSISWSRVLPGGRLAAGVNKDGVKFYHD
FIDELLANGIKPSVTLFHWDLPQALEDEYGGFLSHRIVDDFCEYAEFCFWEFGDKIKYWTTFNEPHTFAVNG
YALGEFAPGRGGKGDEGDPAIEPYVVTHNILLAHKAAVEEYRNKFQKCQEGEIGIVLNSMWMEPLSDVQA
DIDAQKRALDFMLGWFLEPLTTGDYPKSMRELVKGRLPKFSADDSEKLKGCYDFIGMNYYTATYVTNAV
KSNSEKLSYETDDQVTKTFERNQKPIGHALYGGWQHVVPWGLYKLLVYTKETYHVPVLYVTESGMVEEN
KTKILLSEARRDAERTDYHQKHLASVRDAIDDGVNVKGYFVWSFFDNFEWNLGYICRYGIIHVDYKSFERY
PKESAIWYKNFIAGKSTTSPA*

>RIDD

CGSLRECELYVQKHNIQALLKDSIVQLCTARPERPMAFLREYFERLEKEEAK*

>RIAD
CGLEQYANQLADQIIKEATEGC*

>SpyCatcher
AMVDTLSGLSSEQGQSGDMTIEEDSATHIKFSKRDEDGKELAGATMELRDSSGKTISTWISDGQVKDFYLY
PGKYTFVETAAPDGYEVATAITFTVNEQGQVTVNGKATKGDAHI*

>SpyTag
AHIVMVDAYKPTK*

>Nab2
MSQEQYTENLKVIVAEKLAGIPNFNEDIKYVAEYIVLLIVNGGTVESVVDELASLFDSVSRDTLANVVQTAF
FALEALQQGESAENIVSKIRMMNAQSLGQSDIAQQQQQQQQQQQPDIAQQQPQQQPQQQPQQQPQQQPQQ
QPQQQPQQQPQQQPQLQPLQPQLGTQNAMQTDAPATPSPISAFSGVVNAAAPPQFAPVDNSQRFTQRGGG
AVGKNRRGGRGGNRGGRNNNSTRFNPLAKALGMAGESNMNFTPTKKEGRCRLFPHCPLGRSCPHAHPTK
VCNEYPNCPKPPGTCEFLHPNEDEELMKEMERTREEFQKRKADLLAAKRKPVQTGIVLCKFGALCSNPSCP
FGHPTPANEDAKVIDLMWCDKNLTCDNPECRKAHSSLSKIKEVKPISQKKAAPPPVEKSLEQCKFGTHCTN
KRCKYRHARSHIMCREGANCTRIDCLFGHPINEDCRFGVNCKNIYCLFRHPPGRVLPEKKGAAPNSNVPTN
ERPFALPENAIIENAPPQTSFTHQEQDTEMN*

>mCherry
MEFVSKGEEDNMAIIKEFMRFKVHMEGSVNGHEFEIEGEGEGRPYEGTQTAKLKVTKGGPLPFAWDILSPQ
FMYGSKAYVKHPADIPDYLKLSFPEGFKWERVMNFEDGGVVTVTQDSSLQDGEFIYKVKLRGTNFPSDGP
VMQKKTMGWEASSERMYPEDGALKGEIKQRLKLKDGGHYDAEVKTTYKAKKPVQLPGAYNVNIKLDITS
HNEDYTIVEQYERAEGRHSTGGMDELYK*

>EGFP
MEFVSKGEELFTGVVPILVELDGDVNGHKFSVSGEGEGDATYGKLTLKFICTTGKLPVPWPTLVTTLTYGV
QCFSRYPDHMKQHDFFKSAMPEGYVQERTIFFKDDGNYKTRAEVKFEGDTLVNRIELKGIDFKEDGNILGH
KLEYNYNSHNVYIMADKQKNGIKVNFKIRHNIEDGSVQLADHYQQNTPIGDGPVLLPDNHYLSTQSALSK
DPNEKRDHMVLLEFVTAAGITLGMDELYK*

>mVenusC
MDKQKNGIKANFKIRHNIEDGGVQLADHYQQNTPIGDGPVLLPDNHYLSYQSKLSKDPNEKRDHMVLLEF
VTAAGITLGMDELYK*

>mVenusN
MVSKGEELFTGVVPILVELDGDVNGHKFSVSGEGEGDATYGKLTLKLICTTGKLPVPWPTLVTTLGYGLQC
FARYPDHMKQHDFFKSAMPEGYVQERTIFFKDDGNYKTRAEVKFEGDTLVNRIELKGIDFKEDGNILGHKL
EYNYNSHNVYITADKQKNGIKANFKIRHNIE*

>GAL4 transcription activation domain (AD)
MANFNQSGNIADSSLSFTFTNSSNGPNLITTQTNSQALSQPIASSNVHDNFMNNEITASKIDDGNNSKPLSPG
WTDQTAYNAFGITTGMFNTTTMDDVYNYLFDDEDTPPNPKKE*

> GAL4 DNA-binding domain (BD)
MKLLSSIEQACDICRLKKLKCSKEKPKCAKCLKNNWECRYSPKTKRSPLTRAHLTEVESRLERLEQLFLLIF
PREDLDMILKMDSLQDIKALLTGLFVQDNVNKDAVTDRLASVETDMPLTLRQHRISATSSSEESSNKGQRQ
LTVS*

>CrVinBLAST_CrCAD2_AEO99200.1
MAGKSPEEEHPVKTYGLAAHDSSGVLSPFKFSRRATLEDDVRFKVLYCGICHTDLHFAKNEWGISTYPLVP
GHEIVGEVTEVGGKVTKVKVGDKVGVGCLVGSCRTCDNCRADLENYCPKMVLTYASPNVDGTITYGGYS
NEMVCNEHFIVRFPENLPLDGGAPLLCAGITVYSPMKYYGFAKPGSHIAVNGLGGLGHVAVKFAKAMGAK

VTVISTSEGKKDDALNRLGADAFLLSSNPEALQAATGTFDGILNTISAKHAIIPLLGLLKSHGKLVLLGAPPE
PLDLHSAPLLMGRKMVAGSSIGGLKETQEMLDFAGKHNITADIELISADNINTALERLAKGDVRYRFVLDV
AKTLKAP*

>CrCAD1_AYE56096.1
MAGKSPEEEHPIKAYGWAAHDTSGVLSPFKFSRRATLEDDVRLKVLYCGICHTDLHFLKNEWGFTTYPIVP
GHEIVGIVTEVGKKVTKVKVGDKVGVGGIIGSCRSCDRCSADLEQFCPKQVLAFSMPYFDGTITYGGFSNE
MVCNEHFVLRFPENLPLDAGAPLLCAGITVYSTMKLYGIAKPGKHIGVNGLGGLGHMAVKFAKAWGAKV
TVISTSESKKDEAINRLGADAFLLSSNQEEMQAATDTMDGIIDCVPFKHSLMQLLGLLKFEGKLCVVGGVA
EPLEIYCGPLLTARKMIAGNDLGGLKETQEMIDFAAEHNITADIEVISIDDVNEAMERLEKGDVRYRFVIDIA
NTLKAP*

>CrCAD3_ANQ45232.1
MARKSPEDEHPVKAYGWAVKDGTTGILSPFKFSIRATGDNDVRIKILYCGVCRTDLAATKNAFGFLSYPLV
PGREIVGIVSEIGKNVKKVKVGEKVGVAPHVGSCGKCKSCVNEVENFCPKLIIPYGTPYHDGTICYGGFSNE
TVRDERFVFRFPENLSLPGGAPLVSAGVTTYGALRNNGLDKPGLHVGVVGLGGLGHLAVKFAKALGVKV
TVISTNPSKEHDAINGFGADAFILTHHEEQMKAAMGTLDGILYTVPVVHAIAPLLSLLGSQGKFVLIGAPSQ
LLEVPPIQLLFGGKSIIGSAAGNVKQIQEMLEFAAKHDIIANVEIIQMDYINTAMERLDKGDVRYRFVIDIENS
LTLPSEV*

>CrCAD4_AYE56099.1
MAQTTPNHTQTVSGWAAHDSSGKITPYVFKRRENGVNDVTIEVLYCGMCHTDLHHVRNDWGITMYPVVP
GHEITGIITKVGSNVTNFKVGDKVGVGCLAATCLKCEFCKDSQENYCDQVQFTYNGIFWDGSITYGGYSN
MLVADHRYVVRVPDSLPMDAAAPLLCAGITVYCPMKDENLLDSAGEGKKVGIVGLGGLGHVAVKFAKAF
GHHVTIISTSPSKEKEARERLGVDDFVLSTNKEQMQARKRSLDFILDTVAAKHSLGPILELLKVRGTLSIVGA
PDKPMDLPSFPLIFGKRVVKGSMIGSIEETQEMMDLCGKHNITCDIEMVSTNNINEALDRLAKNDVKYRFVI
DIASKSSNL*

>CrCAD5_AYE56100.1
MAGKSPEEQHPVKAYGWAARDSSGILSPFNFSRRATGDHDVRIKILYCGICHSDLQAAKNEMGFYKYPLVP
GFETVGVASEVGSKVTKVKVGDKVAVGIIVGSCGECNECVNDRDCYCPRMTAAYGSVDHDGTPTYGGFS
NETVVNEKFVFRFPENLSLPGGAPLLNAGITVYSPMRFYGLDKPGMHLGVVGLGGLGHLAVKFGKAFGAK
VTVISTSPSKKDEAINFLGADGFLVSGDAEQMQAAAGTLDGIIDTVPVVHSIETLLWLLKNHSKLVLVGATG
GSFDLPILPLAMGRRTVASSIGGSTKEAQEMLDFAAEHNITANVEIIPMDYVNTAMERVEKSDVRYRFVIDI
GNTLTPPPES*

>RsVinBLAST_PV770557
MAGKSPEEEHPVKAYGVAARDSSGVLSPFKFSRRATLEDDVRLKVLYCGLCHTDIHFLKNEWGFSTYPFVP
GHEVVGEVIEVGSKVTKVKVGDKVAHGGIIGSCRACDNCHADMESYCPKMVMAHGSPNFDGTITYGGFS
NEMVVNEHFVIRYPENLPLAAGAPLLCAGITVYSPMKYYGIAKPGNHIGVNGLGGLGHMAVKFAKALGA
KVTVISSSESKKDDAINHLGADAFLLSKNPEELQAATGTLDGIVDCVSAKHPIIPLLGLLKSHGKLVLVGAP
GEPLELHSAPLLMGRKMIGGSDAGGMKEIQEMVDLAAKHNITADIELVSMDNINTVVERLVKGDVRYRFV
VDVANTLKAP*

>MsVinBLAST_PV770558
MAGKSPEEEHPVKTHGWAARDSSGVLSPFKFSRRATLEDDVRFKVLFCGICHTDIHFLKNEWGFSTYPLVP
GHEIVGEVTEVGSKVTKVKVGDKVGVGCLVGSCRTCENCSANLENYCPKMVLTYATPYFDGTITYGGYSN
EMVCNEHFIIRFPENMPLAGGAPLLCAGVTVYSPMKYYGLAKPGSHIGINGLGGLGHVAVKFAKALGAKV
TVISTSDRKKEEALKHLGADAFLISRNADEMQAAAGTLDGILDCVSAKHALVPLLGLLKYHGKLITVGAPA
EPLELPVAPLIMGRKLVGGSNIGGLAETQEMIDLAAKHNITADIEIVSIDDVNTALERLAKGDVRYRFVIDV
ANTLKAP*

>CoVinBLAST_PV770559
MAGKSPEEEHPVKTHGWAARDSSGVLSPFKFSRRATLEDDVRFKVLFCGVCHTDLHFLKNEWGFSTYPLV
PGHEIVGEVTEVGSKVTKVKVGDKVGVGCLVGSCRTCENCSANLENYCPKMVLTYATPYFDGTITYGGYS
NEMVCNEHFIIRFPENMSLAGGAPLLCAGVTVYSPMKYYGLAKPGSHIGINGLGGLGHVAVKFAKALGAK

VTVISTSDRKKEEALKHLGADAFLISRNADEMQAAAGTLDGILDCVSAKHALVPLLGLLKYHGKLITVGAP
AEPLELPVAPLIMGRKLVGGSNIGGLEETQEMIDLAAKHNITADIEVVSMEDVNTALERLAKGDVRYRFVI
DVANTLKAP*

>GsVinBLAST_PV770560
MAGKSPEDEHPVKTYGWAARDSSGVLSPFKFSRRETLEDDVRLKVLYCGICHTDLHFLKNDWGLSTYPLV
PGHEIVGVVTEVGSKVTKVKVGDKVGVGYLVGSCRTCENCSADLENYCPKMVLTYGTTYFDGTITYGGY
SNEMVCNEHFIIHFPENIPLDAGAPLLCAGITVYSPMKYYGIAKPGSHVGVNGLGGVGHVAVKFAKALGAK
VTVISRSPNKKDEAINHLGADAFLVSRNPDEMQAAMGTMDGILDCVSAKHALLPLIGLLKSHGKLCLIGAP
AEPLELPPSSLIMGRKMVGGSNVGGLKETQEMIEFAAQHNITADVEVVSMDYVNTALERLVKGDVRYRFV
LDVGNTLKAP*

>SspVinBLAST_PV770561
MAGKSPEEVHPVKAYGWAARDTSGVLSPFKFSRRETKEDDVRFKVLYCGVCHTDLHFLKNEWGMTTYPL
VPGHEIVGEVTEVGSKVSKVKVGDKVGVGYFVGSCRSCVNCSADFENYCPKMVLTYGVPYFDGTITYGG
YSNEMVSNEHFVTRFPENMPLDGGAPLLCAGVTVYSALKYYGMAKPGSHIGVNGLGGLGHVAVKFAKAF
GAKVTVISTSPSKKEEALSRLGADAFLVSRNAEEMQAAMGTMDGIIDCVSAKHPLMPLLGLLNYHGKICV
VGAPVEPLELPAGPLLMGRKLVGGSNIGGLKETQEMIEFAAKHNITADVEVIPVDYVNTALERLEKGDVRY
RFVIDVGNTLKAP*

>VvVinBLAST_PV770562
MAKSAEQEHPTKAFGWAATDTSGVLSPFKFSRRETGEKDVRFKVLYCGICHSDLHMAKNDWGTSTYPIVP
GHEIVGIVTEVGSKVGKFKVGDRVGVGCMVGACHSCDSCDNDLENYCPKMVFTYSAPYHDGTTTYGGYS
DVMVAEERYVVRIPDNLPLDAGAPLLCAGITVYSPLQHFGLTKPGMHIGVVGLGGLGHVAVKFAKGFGVK
VTVISTSPSKKKEAIEHLGADYFLVSREPDQMQAAMGTMDGIIDTVSAVHPLLPLIGLLKSQGKLVMVGAP
NRPLELPVFPLIMGRKVVAGSCIGGMKETQEMIDFAGKHNITADVEVIPMDYVNTAMERLEKADVRYRFVI
DIGNTLKSA*

>NbVinBLAST_PV770563
MEKSLEEVHPVKAFGWAAKDTSGLLSPFKFSRRATGDKDVKLKILYCGICHTDLHQVKNEWGFSRYPMVP
GHEIVGEVTEVGSKVDKFKVGEKAGVGCLVGSCRSCENCTNDLENYCFKGIATYCATYLDGTPTYGGYSD
GIVVDEHFVVHVPENMPLAAAAPLLCAGITTYSPLRYFGLDKPGLHIGVVGLGGLGHVGVKFAKALGLKV
TVISTSQSKQKEAIEHLGADSFLISTDPDQLQAAMGTMDGILDTVAATHPVLPLVGLLKTNGKLILLGALEK
PLELPVFPLLMGRKLVAGSGIGGMKETQEMLDFAAKHNITADIEVIPMDYINTAMERLAKADVKYRFVIDV
AKTLKAE*

**Table S4.** List of the CAD1-5 nucleotide sequences tested by VIGS.

>CrCAD1_MH194345
ATGGCCGGAAAATCACCAGAGGAGGAGCACCCAATCAAGGCCTACGGATGGGCTGCTCACGATACAT
CTGGGGTCCTTTCTCCCTTCAAATTCTCCAGGAGGGCAACTCTTGAGGACGATGTCAGGCTCAAGGTGC
TCTATTGTGGAATTTGTCATACAGACCTTCATTTCCTGAAAAATGAGTGGGGCTTTACTACCTATCCTA
TTGTACCAGGCCATGAAATTGTAGGTATAGTTACAGAGGTTGGTAAAAAAGTTACAAAAGTCAAGGTT
GGAGATAAAGTTGGCGTTGGGGGCATTATAGGTTCTTGCCGCAGTTGTGATAGATGTTCTGCAGATTT
GGAGCAGTTTTGTCCCAAACAGGTCTTAGCCTTTTCAATGCCATATTTTGATGGGACCATTACATATGG
AGGCTTTTCAAATGAGATGGTATGCAATGAACATTTTGTTCTTCGTTTCCCAGAGAACCTACCACTTGA
TGCCGGTGCACCATTGCTGTGTGCCGGAATTACTGTGTATAGTACAATGAAACTCTACGGCATTGCCA
AACCGGGAAAGCACATAGGCGTTAACGGTCTTGGTGGGCTTGGGCATATGGCTGTTAAGTTTGCAAAG
GCTTGGGGAGCAAAAGTAACTGTTATCAGTACATCTGAGAGCAAGAAAGACGAAGCTATAAATCGTTT
GGGTGCAGATGCATTTTTGCTGAGCAGTAATCAAGAAGAAATGCAGGCGGCAACAGACACAATGGAC
GGAATAATTGATTGTGTTCCTTTTAAGCACTCGCTTATGCAATTGCTCGGTCTACTCAAGTTTGAAGGA
AAGCTTTGTGTGGTTGGGGGAGTAGCAGAGCCACTAGAGATTTATTGTGGTCCTTTGCTCACTGCAAG
GAAGATGATTGCTGGAAATGACCTTGGAGGATTGAAGGAGACTCAAGAAATGATTGATTTTGCAGCA
GAACACAACATCACTGCAGATATAGAAGTCATTTCCATAGATGATGTCAACGAAGCTATGGAACGCCT
TGAAAAGGGTGATGTTAGATATCGCTTTGTCATTGATATTGCCAACACCTTGAAAGCTCCTTAA

>CrCAD2_MH194346
ATGGCCGGAAAATCACCAGAAGAGGAGCACCCAGTCAAGACCTATGGATTGGCTGCTCATGATTCATC
TGGGGTTTTATCTCCGTTCAAATTCTCCAGGAGGGCAACTCTTGAGGATGATGTGAGGTTCAAGGTGCT
ATATTGTGGGATTTGTCATACTGACCTTCATTTCGCTAAGAATGAGTGGGGTATTTCGACCTATCCTCT
TGTACCAGGACATGAAATCGTAGGGGAAGTTACAGAGGTCGGCGGCAAAGTTACAAAGGTCAAGGTT
GGAGATAAAGTTGGTGTTGGCTGCTTGGTTGGTTCATGCCGCACTTGTGATAATTGTCGTGCAGATCTT
GAGAACTATTGTCCCAAAATGGTGCTAACCTATGCAAGTCCAAACGTTGATGGAACGATTACCTATGG
AGGCTATTCCAATGAGATGGTATGCAATGAACACTTTATTGTTCGTTTCCCAGAGAACCTACCACTTGA
TGGTGGGGCACCATTGCTTTGTGCCGGTATTACTGTGTACAGTCCAATGAAATACTATGGCTTTGCCAA
ACCCGGGAGCCACATAGCTGTTAATGGTCTTGGTGGACTTGGCCATGTGGCTGTTAAGTTTGCAAAGG
CCATGGGAGCAAAAGTGACAGTTATAAGTACATCTGAGGGCAAGAAAGACGATGCCCTCAATCGTTT
GGGTGCAGATGCATTTTTGTTGAGCAGTAATCCAGAAGCACTGCAGGCTGCAACAGGCACATTTGATG
GCATACTTAATACTATTTCTGCTAAGCACGCTATTATCCCATTGCTTGGTCTACTAAAGTCTCATGGCA
AGCTTGTTCTTCTTGGGGCACCCCCGGAACCACTTGATCTTCACTCTGCTCCTTTGCTTATGGGGAGGA
AGATGGTTGCTGGAAGTAGCATTGGAGGATTGAAGGAGACCCAAGAGATGCTTGATTTTGCCGGAAA
GCATAACATTACTGCAGATATAGAACTCATTTCCGCGGACAATATCAACACAGCTTTGGAGCGTCTGG
CCAAGGGTGATGTTAGATATCGCTTTGTCCTTGACGTTGCAAAGACCTTGAAAGCTCCTTAA

>CrCAD3_MH194347.1
ATGGCCAGAAAATCACCAGAAGATGAACATCCCGTGAAGGCTTACGGATGGGCCGTCAAAGATGGAA
CAACTGGAATTCTTTCTCCCTTCAAATTTTCCATAAGGGCAACAGGTGATAATGATGTTCGAATCAAGA
TCCTCTATTGTGGAGTTTGTCGTACCGATCTTGCGGCAACCAAGAACGCATTCGGGTTTCTTTCTTATCC
TCTTGTGCCTGGTAGAGAGATCGTGGGAATAGTGAGCGAGATAGGGAAAAATGTGAAAAAAGTTAAA
GTTGGAGAAAAAGTTGGAGTAGCCCCGCATGTGGGTAGCTGTGGCAAATGCAAGAGTTGTGTGAATG
AGGTGGAGAATTTCTGTCCGAAACTGATCATCCCTTATGGCACCCCATACCACGATGGTACTATTTGCT
ACGGTGGTTTCTCCAACGAGACTGTCAGAGATGAACGCTTTGTTTTTCGTTTTCCTGAAAATCTTTCGC
TGCCTGGCGGAGCTCCCTTGGTTAGTGCTGGGGTTACCACGTACGGTGCATTGAGAAATAATGGCCTC
GACAAGCCCGGATTACACGTGGGAGTCGTCGGTCTAGGTGGATTAGGTCATCTGGCTGTTAAATTTGC
TAAGGCTTTAGGCGTCAAAGTAACTGTTATTAGTACCAATCCTAGCAAGGAGCATGATGCTATAAATG
GTTTCGGTGCTGATGCCTTCATCCTCACCCACCATGAGGAACAAATGAAGGCTGCCATGGGAACTTTA
GATGGAATTCTTTATACAGTGCCTGTTGTTCATGCCATTGCACCATTACTTAGTCTACTGGGAAGTCAA
GGGAAATTTGTGTTGATTGGGGCACCATCTCAATTACTTGAGGTGCCACCTATTCAATTATTATTTGGT
GGAAAATCTATTATTGGAAGTGCGGCTGGAAATGTGAAGCAAATCCAAGAAATGCTTGAATTTGCAGC
AAAACATGATATAATTGCGAATGTTGAGATTATCCAAATGGATTATATAAATACTGCAATGGAACGTC

TAGACAAAGGTGATGTTAGATATCGATTTGTAATTGATATCGAAAACTCTCTCACTCTTCCATCAGAGG
TGTGA

>CrCAD4_MH194348
ATGGCTCAAACAACTCCAAACCATACACAGACAGTTTCAGGCTGGGCTGCTCATGATTCTTCTGGCAA
AATCACTCCTTATGTCTTCAAGAGAAGGGAAAATGGGGTTAACGATGTTACCATAGAGGTTTTGTACT
GTGGGATGTGCCATACTGATCTCCACCATGTCCGTAACGATTGGGGTATTACAATGTACCCTGTTGTAC
CCGGCCATGAAATTACTGGAATTATCACAAAAGTGGGGAGCAATGTGACGAATTTCAAGGTAGGGGA
CAAAGTTGGAGTGGGTTGCTTGGCAGCTACTTGCTTGAAATGTGAGTTCTGCAAAGACTCCCAAGAAA
ACTACTGCGATCAAGTACAGTTCACCTACAATGGCATTTTTTGGGACGGTTCTATCACCTATGGTGGCT
ACTCCAACATGCTTGTTGCAGATCACCGGTACGTGGTGCGTGTGCCGGATAGTCTACCGATGGATGCA
GCGGCACCGCTGTTATGTGCCGGAATAACAGTATACTGCCCCATGAAAGACGAGAACTTGCTGGATTC
AGCCGGAGAAGGGAAGAAAGTAGGCATAGTCGGCCTAGGAGGTCTAGGTCATGTGGCTGTGAAATTT
GCTAAGGCATTTGGTCACCATGTCACTATTATAAGCACATCTCCATCAAAAGAAAAAGAAGCAAGAGA
AAGATTGGGAGTTGATGATTTTGTTCTTAGCACCAATAAAGAACAAATGCAGGCAAGGAAAAGGAGC
CTAGACTTTATTTTGGACACTGTCGCGGCCAAGCACTCTCTTGGACCAATACTTGAGCTCCTCAAAGTT
AGAGGAACTTTGTCCATTGTTGGTGCTCCAGATAAGCCCATGGACCTACCTTCCTTTCCATTGATTTTT
GGGAAGAGAGTGGTGAAAGGAAGTATGATTGGAAGTATTGAAGAAACTCAAGAGATGATGGATTTAT
GTGGTAAGCATAACATCACTTGTGACATTGAAATGGTTTCAACAAATAACATCAATGAAGCTCTTGAT
CGGCTTGCCAAGAATGATGTTAAGTATCGTTTTGTGATCGATATAGCTTCCAAATCATCTAATCTTTAA

>CrCAD5_MH194349
ATGGCTGGAAAATCACCAGAAGAGCAACACCCTGTGAAGGCTTATGGATGGGCAGCTAGAGACTCAT
CTGGAATTCTTTCTCCCTTCAACTTCTCCAGAAGGGCAACCGGGGATCATGATGTCAGAATAAAGATTC
TCTACTGTGGTATTTGCCATTCTGACCTTCAAGCTGCCAAGAACGAAATGGGCTTCTATAAATATCCTC
TTGTGCCCGGGTTTGAGACAGTAGGCGTAGCAAGTGAAGTTGGAAGCAAGGTCACAAAAGTGAAAGT
CGGTGATAAAGTTGCCGTTGGAATCATTGTGGGATCATGTGGAGAATGTAATGAGTGTGTCAATGACC
GTGATTGTTACTGCCCTAGGATGACCGCTGCTTATGGTTCAGTAGACCATGATGGAACTCCCACTTATG
GAGGTTTCTCGAACGAGACAGTAGTAAATGAGAAGTTTGTCTTTCGTTTTCCTGAAAACCTTTCGCTTC
CTGGTGGTGCTCCATTGCTTAATGCTGGAATAACCGTGTACAGTCCCATGAGATTTTATGGCCTGGACA
AGCCAGGAATGCACCTGGGAGTTGTTGGTCTAGGTGGACTTGGTCATTTAGCTGTGAAGTTTGGCAAG
GCTTTTGGGGCCAAAGTCACTGTAATTAGTACCTCTCCAAGCAAGAAAGATGAAGCTATCAATTTTCTT
GGTGCTGATGGCTTCTTGGTCAGCGGTGATGCTGAACAAATGCAGGCTGCTGCTGGAACCTTGGATGG
GATTATTGATACCGTGCCTGTTGTCCATTCTATTGAGACCTTGCTATGGCTTTTGAAGAATCATTCCAA
GCTTGTTTTGGTTGGAGCTACAGGGGGCTCATTCGATTTGCCAATTCTTCCTTTAGCAATGGGCAGAAG
AACTGTGGCTTCAAGCATTGGTGGAAGTACGAAGGAGGCTCAAGAGATGCTGGATTTTGCAGCAGAA
CACAATATCACTGCAAACGTTGAGATTATTCCAATGGACTATGTCAACACAGCAATGGAACGTGTTGA
AAAGAGTGATGTTCGTTATCGATTTGTGATTGATATTGGAAACACCTTAACTCCTCCACCAGAGTCCTA
A

**Table S5.** Plasmids used in this study.

| Plasmid | Description |
|---|---|
| pESC-URA-*GS* | 2μ; *URA3*; *AmpR*; *GAL1p-GS-CYC1t* |
| pESC-URA-*GS*$^{I301E}$ | 2μ; *URA3*; *AmpR*; *GAL1p-GS*$^{I301E}$*-CYC1t* |
| pESC-URA-*GS*$^{P290H,A291S,I295L}$ | 2μ; *URA3*; *AmpR*; *GAL1p-GS*$^{P290H,A291S,I295L}$*-CYC1t* |
| pESC-URA-*MsC17OMT* | 2μ; *URA3*; *AmpR*; *GAL1p-MsC17OMT-CYC1t* |
| pESC-URA-*SGD* | 2μ; *URA3*; *AmpR*; *GAL1p-SGD-CYC1t* |
| pESC-LEU-*SGD* | 2μ; *LEU2*; *AmpR*; *GAL1p-SGD-CYC1t* |
| pESC-LEU-*tSGD* | 2μ; *LEU2*; *AmpR*; *GAL1p-tSGD-CYC1t* |
| pRS423-SpSgΔ*SGD* | 2μ; *HIS3*; *AmpR*; *SNR52p*-SpSgΔ*SGD-SUP4t* |
| pRS423-SpSgΔ*GS* | 2μ; *HIS3*; *AmpR*; *SNR52p*-SpSgΔ*GS-SUP4t* |
| pESC-URA-*GS~SGD* | 2μ; *URA3*; *AmpR*; *GAL1p-GS~SGD-CYC1t* |
| pESC-URA-*GS~RIDD* | 2μ; *URA3*; *AmpR*; *GAL1p-GS~RIDD-CYC1t* |
| pESC-URA-*GS*$^{P290H,A291S,I295L}$*~RIDD* | 2μ; *URA3*; *AmpR*; *GAL1p-GS*$^{P290H,A291S,I295L}$*~RIDD-CYC1t* |
| pESC-URA-*GS~SpyTag* | 2μ; *URA3*; *AmpR*; *GAL1p-GS~SpyTag -CYC1t* |
| pESC-URA-*GS*$^{P290H,A291S,I295L}$*~SpyTag* | 2μ; *URA3*; *AmpR*; *GAL1p-GS*$^{P290H,A291S,I295L}$*~SpyTag -CYC1t* |
| pESC-LEU-*RIAD~SGD* | 2μ; *LEU2*; *AmpR*; *GAL1p-RIAD~SGD-CYC1t* |
| pESC-LEU-*RIAD~tSGD* | 2μ; *LEU2*; *AmpR*; *GAL1p-RIAD~tSGD-CYC1t* |
| pESC-LEU-*SpyCatcher~SGD* | 2μ; *LEU2*; *AmpR*; *GAL1p-SpyCatcher~SGD-CYC1t* |
| pESC-LEU-*SpyCatcher~tSGD* | 2μ; *LEU2*; *AmpR*; *GAL1p-SpyCatcher~tSGD-CYC1t* |
| pESC-URA-*CrVinBLAST* | 2μ; *URA3*; *AmpR*; *GAL1p-CrVinBLAST-CYC1t* |
| pESC-URA-*CrCAD1* | 2μ; *URA3*; *AmpR*; *GAL1p-CrCAD1-CYC1t* |
| pESC-URA-*CrCAD3* | 2μ; *URA3*; *AmpR*; *GAL1p-CrCAD3-CYC1t* |
| pESC-URA-*CrCAD5* | 2μ; *URA3*; *AmpR*; *GAL1p-CrCAD5-CYC1t* |
| pESC-URA-*CrVinBLAST*$^{C51A,H56A}$ | 2μ; *URA3*; *AmpR*; *GAL1p-CrVinBLAST*$^{C51A,H56A}$*-CYC1t* |
| pESC-URA-*CrVinBLAST*$^{M298E,V299E}$ | 2μ; *URA3*; *AmpR*; *GAL1p-CrVinBLAST*$^{M298E,V299E}$*-CYC1t* |
| pESC-URA-*CrVinBLAST*$^{H288P,S289A,L293I}$ | 2μ; *URA3*; *AmpR*; *GAL1p-CrVinBLAST*$^{H288P,S289A,L293I}$*-CYC1t* |
| pESC-URA-*CrVinBLAST*$^{K359G}$ | 2μ; *URA3*; *AmpR*; *GAL1p-CrVinBLAST*$^{K359G}$*-CYC1t* |
| pESC-URA-*CoVinBLAST* | 2μ; *URA3*; *AmpR*; *GAL1p-CoVinBLAST-CYC1t* |
| pESC-URA-*RsVinBLAST* | 2μ; *URA3*; *AmpR*; *GAL1p-RsVinBLAST-CYC1t* |
| pESC-URA-*MsVinBLAST* | 2μ; *URA3*; *AmpR*; *GAL1p-MsVinBLAST-CYC1t* |
| pESC-TRP-*Nab2~mCherry* | 2μ; *TRP1*; *AmpR*; *GAL10p-Nab2~mCherry-ADH1t* |
| pESC-HIS-*EGFP~SGD* | 2μ; *HIS3*; *AmpR*; *GAL10p-EGFP~SGD-ADH1t* |
| pESC-HIS-*EGFP~tSGD* | 2μ; *HIS3*; *AmpR*; *GAL10p-EGFP~tSGD-ADH1t* |
| pESC-URA-*SGD~EGFP* | 2μ; *URA3*; *AmpR*; *GAL10p-SGD~EGFP-ADH1t* |
| pESC-URA-*GS~EGFP* | 2μ; *URA3*; *AmpR*; *GAL10p-GS~EGFP-ADH1t* |
| pESC-URA-*EGFP~GS* | 2μ; *URA3*; *AmpR*; *GAL10p-EGFP~GS-ADH1t* |
| pESC-URA-*CrVinBLAST~EGFP* | 2μ; *URA3*; *AmpR*; *GAL10p-CrVinBLAST~EGFP-ADH1t* |
| pESC-URA-*EGFP~CrVinBLAST* | 2μ; *URA3*; *AmpR*; *GAL10p-EGFP~CrVinBLAST-ADH1t* |
| pESC-HIS-*CrVinBLAST~mVenus*$^{C}$ | 2μ; *HIS3*; *AmpR*; *GAL10p-CrVinBLAST~mVenus*$^{C}$*-ADH1t* |
| pESC-HIS-*CrVinBLAST*$^{C51A,H56A}$*~mVenus*$^{C}$ | 2μ; *HIS3*; *AmpR*; *GAL10p-CrVinBLAST*$^{C51A,H56A}$*~mVenus*$^{C}$*-ADH1t* |
| pESC-HIS-*CrVinBLAST*$^{M298E,V299E}$*~mVenus*$^{C}$ | 2μ; *HIS3*; *AmpR*; *GAL10p-CrVinBLAST*$^{M298E,V299E}$*~mVenus*$^{C}$*-ADH1t* |
| pESC-HIS-*CrCAD1~mVenus*$^{C}$ | 2μ; *HIS3*; *AmpR*; *GAL10p-CrCAD1~mVenus*$^{C}$*-ADH1t* |
| pESC-HIS-*GS~mVenus*$^{C}$ | 2μ; *HIS3*; *AmpR*; *GAL10p-GS~mVenus*$^{C}$*-ADH1t* |
| pESC-HIS-*GS*$^{I301E}$*~mVenus*$^{C}$ | 2μ; *HIS3*; *AmpR*; *GAL10p-GS*$^{I301E}$*~mVenus*$^{C}$*-ADH1t* |
| pESC-HIS-*mVenus*$^{C}$*~SGD* | 2μ; *HIS3*; *AmpR*; *GAL10p-mVenus*$^{C}$*~SGD-ADH1t* |
| pESC-LEU-*CrVinBLAST~mVenus*$^{N}$ | 2μ; *LEU2*; *AmpR*; *GAL10p-CrVinBLAST~mVenus*$^{N}$*-ADH1t* |
| pESC-LEU-*CrVinBLAST*$^{C51A,H56A}$*~mVenus*$^{N}$ | 2μ; *LEU2*; *AmpR*; *GAL10p-CrVinBLAST*$^{C51A,H56A}$*~mVenus*$^{N}$*-ADH1t* |
| pESC-LEU-*CrVinBLAST*$^{M299E,V299E}$*~mVenus*$^{N}$ | 2μ; *LEU2*; *AmpR*; *GAL10p-CrVinBLAST*$^{M299E,V299E}$*~mVenus*$^{N}$*-ADH1t* |
| pESC-LEU-*CrVinBLAST*$^{K359G}$*~mVenus*$^{N}$ | 2μ; *LEU2*; *AmpR*; *GAL10p-CrVinBLAST*$^{K359G}$*~mVenus*$^{N}$*-ADH1t* |
| pESC-LEU-*CrCAD1~mVenus*$^{N}$ | 2μ; *LEU2*; *AmpR*; *GAL10p-CrCAD1~mVenus*$^{N}$*-ADH1t* |
| pESC-LEU-*GS~mVenus*$^{N}$ | 2μ; *LEU2*; *AmpR*; *GAL10p-GS~mVenus*$^{N}$*-ADH1t* |
| pESC-LEU-*GS*$^{I301E}$*~mVenus*$^{N}$ | 2μ; *LEU2*; *AmpR*; *GAL10p-GS*$^{I301E}$*~mVenus*$^{N}$*-ADH1t* |
| pESC-LEU-*mVenus*$^{N}$*~SGD* | 2μ; *LEU2*; *AmpR*; *GAL10p-mVenus*$^{N}$*~SGD-ADH1t* |

| **Plasmid** | **Description** |
|---|---|
| pGADT7-LEU-*AD~GS* | 2µ; *LEU2*; *AmpR*; *ADH1p-AD~GS-ADH1t* |
| pGADT7-LEU-*AD~GS*$^{I301E}$ | 2µ; *LEU2*; *AmpR*; *ADH1p-AD~GS*$^{I301E}$*-ADH1t* |
| pGADT7-LEU-*AD~CrVinBLAST* | 2µ; *LEU2*; *AmpR*; *ADH1p-AD~CrVinBLAST-ADH1t* |
| pGADT7-LEU-*AD~CrVinBLAST*$^{C51A,H56A}$ | 2µ; *LEU2*; *AmpR*; *ADH1p-AD~CrVinBLAST*$^{C51A,H56A}$*-ADH1t* |
| pGADT7-LEU-*AD~CrVinBLAST*$^{M299E,V299E}$ | 2µ; *LEU2*; *AmpR*; *ADH1p-AD~CrVinBLAST*$^{M298E,V299E}$*-ADH1t* |
| pGADT7-LEU-*AD~CrCAD1* | 2µ; *LEU2*; *AmpR*; *ADH1p-AD~CrCAD1-ADH1t* |
| pGBKT7-TRP-*BD~GS* | 2µ; *TRP1*; *KanR*; *ADH1p-BD~GS-ADH1t* |
| pGBKT7-TRP-*BD~GS*$^{I301E}$ | 2µ; *TRP1*; *KanR*; *ADH1p-BD~GS*$^{I301E}$*-ADH1t* |
| pGBKT7-TRP-*BD~CrVinBLAST* | 2µ; *TRP1*; *KanR*; *ADH1p-BD~CrVinBLAST-ADH1t* |
| pGBKT7-TRP-*BD~CrVinBLAST*$^{C51A,H56A}$ | 2µ; *TRP1*; *KanR*; *ADH1p-BD~CrVinBLAST*$^{C51A,H56A}$*-ADH1t* |
| pGBKT7-TRP-*BD~CrVinBLAST*$^{M298E,V299E}$ | 2µ; *TRP1*; *KanR*; *ADH1p-BD~CrVinBLAST*$^{M298E,V299E}$*-ADH1t* |
| pGBKT7-TRP-*BD~CrVinBLAST*$^{K359G}$ | 2µ; *TRP1*; *KanR*; *ADH1p-BD~CrVinBLAST*$^{K359G}$*-ADH1t* |
| pGBKT7-TRP-*BD~CrCAD1* | 2µ; *TRP1*; *KanR*; *ADH1p-BD~CrCAD1-ADH1t* |
| pET30b(+)-*CrGS* | *KanR*; *T7p-CrGS-T7t* |
| pET30b(+)-*CrGS*$^{I301E}$ | *KanR*; *T7p-CrGS*$^{I301E}$*-T7t* |
| pET30b(+)-*CrGS*$^{P290H,A291S,I295L}$ | *KanR*; *T7p-CrGS*$^{P290H,A291S,I295L}$*-T7t* |
| pET30b(+)-*GsGS* | *KanR*; *T7p-GsGS-T7t* |
| pET30b(+)-*SspGS* | *KanR*; *T7p-SspGS-T7t* |
| pET30b(+)-*CrVinBLAST* | *KanR*; *T7p-CrVinBLAST-T7t* |
| pET30b(+)-*CrVinBLAST*$^{C51A,H56A}$ | *KanR*; *T7p-CrVinBLAST*$^{C51A, H56A}$*-T7t* |
| pET30b(+)-*CrVinBLAST*$^{M298E,V299E}$ | *KanR*; *T7p-CrVinBLAST*$^{M298E,V299E}$*-T7t* |
| pET30b(+)-*CrVinBLAST*$^{H288P,S289A, I295L}$ | *KanR*; *T7p-CrVinBLAST*$^{H288P,S289A,L293I}$*-T7t* |
| pET30b(+)-*CrVinBLAST*$^{K359G}$ | *KanR*; *T7p-CrVinBLAST*$^{K359G}$*-T7t* |
| pET30b(+)-*GsVinBLAST* | *KanR*; *T7p-GsVinBLAST-T7t* |
| pET30b(+)-*SspVinBLAST* | *KanR*; *T7p-SspVinBLAST-T7t* |
| pET30b(+)-*MsVinBLAST* | *KanR*; *T7p-MsVinBLAST-T7t* |
| pET30b(+)-*VvVinBLAST* | *KanR*; *T7p-VvVinBLAST-T7t* |
| pET30b(+)-*NbVinBLAST* | *KanR*; *T7p-NbVinBLAST-T7t* |
| pET30b(+)-*CrSGD* | *KanR*; *T7p-CrSGD-T7t* |
| pGTQL1211YN-*CrVinBLAST~mVenus*$^{N}$ | *KanR*; *35Sp-CrVinBLAST~mVenus*$^{N}$*-OCSt* |
| pGTQL1211YN-*mVenus*$^{C}$*~CrSGD* | *KanR*; *35Sp-mVenus*$^{C}$*~CrSGD-OCSt* |
| pGTQL1211YN-*mVenus*$^{N}$*~CrGS* | *KanR*; *35Sp-mVenus*$^{N}$*~CrGS-OCSt* |
| pGTQL1211YN-*mVenus*$^{N}$*~CrVinBLAST* | *KanR*; *35Sp-mVenus*$^{N}$*~CrVinBLAST-OCSt* |
| pGTQL1211YN-*mVenus*$^{C}$*~CrVinBLAST* | *KanR*; *35Sp-mVenus*$^{C}$*~CrVinBLAST-OCSt* |
| pGTQL1211YN-*mVenus*$^{C}$*~NbVinBLAST* | *KanR*; *35Sp-mVenus*$^{C}$*~NbVinBLAST-OCSt* |

**Table S6.** List of the complete sequences of all yeast BiFC plasmids.

> pESC-HIS-*CrVinBLAST~mVenus*$^{C}$

TCGCGCGTTTCGGTGATGACGGTGAAAACCTCTGACACATGCAGCTCCCGGAGACGGTCACAGCT
TGTCTGTAAGCGGATGCCGGGAGCAGACAAGCCCGTCAGGGCGCGTCAGCGGGTGTTGGCGGGT
GTCGGGGCTGGCTTAACTATGCGGCATCAGAGCAGATTGTACTGAGAGTGCACCATAAATTCCCG
TTTTAAGAGCTTGGTGAGCGCTAGGAGTCACTGCCAGGTATCGTTTGAACACGGCATTAGTCAGG
GAAGTCATAACACAGTCCTTTCCCGCAATTTTCTTTTTCTATTACTCTTGGCCTCCTCTAGTACACT
CTATATTTTTTTATGCCTCGGTAATGATTTTCATTTTTTTTTTTCCCCTAGCGGATGACTCTTTTTTT
TTCTTAGCGATTGGCATTATCACATAATGAATTATACATTATATAAAGTAATGTGATTTCTTCGAA
GAATATACTAAAAAATGAGCAGGCAAGATAAACGAAGGCAAAGATGACAGAGCAGAAAGCCCT
AGTAAAGCGTATTACAAATGAAACCAAGATTCAGATTGCGATCTCTTTAAAGGGTGGTCCCCTAG
CGATAGAGCACTCGATCTTCCCAGAAAAAGAGGCAGAAGCAGTAGCAGAACAGGCCACACAATC
GCAAGTGATTAACGTCCACACAGGTATAGGGTTTCTGGACCATATGATACATGCTCTGGCCAAGC
ATTCCGGCTGGTCGCTAATCGTTGAGTGCATTGGTGACTTACACATAGACGACCATCACACCACT
GAAGACTGCGGGATTGCTCTCGGTCAAGCTTTTAAAGAGGCCCTACTGGCGCGTGGAGTAAAAA
GGTTTGGATCAGGATTTGCGCCTTTGGATGAGGCACTTTCCAGAGCGGTGGTAGATCTTTCGAAC
AGGCCGTACGCAGTTGTCGAACTTGGTTTGCAAAGGGAGAAAGTAGGAGATCTCTCTTGCGAGAT
GATCCCGCATTTTCTTGAAAGCTTTGCAGAGGCTAGCAGAATTACCCTCCACGTTGATTGTCTGCG
AGGCAAGAATGATCATCACCGTAGTGAGAGTGCGTTCAAGGCTCTTGCGGTTGCCATAAGAGAA
GCCACCTCGCCCAATGGTACCAACGATGTTCCCTCCACCAAAGGTGTTCTTATGTAGTGACACCG
ATTATTTAAAGCTGCAGCATACGATATATATACATGTGTATATATGTATACCTATGAATGTCAGTA
AGTATGTATACGAACAGTATGATACTGAAGATGACAAGGTAATGCATCATTCTATACGTGTCATT
CTGAACGAGGCGCGCTTTCCTTTTTTCTTTTTGCTTTTTCTTTTTTTTTCTCTTGAACTCGACGGATC
TATGCGGTGTGAAATACCGCACAGATGCGTAAGGAGAAAATACCGCATCAGGAAATTGTAAACG
TTAATATTTTGTTAAAATTCGCGTTAAATTTTTGTTAAATCAGCTCATTTTTTAACCAATAGGCCG
AAATCGGCAAAATCCCTTATAAATCAAAAGAATAGACCGAGATAGGGTTGAGTGTTGTTCCAGTT
TGGAACAAGAGTCCACTATTAAAGAACGTGGACTCCAACGTCAAAGGGCGAAAAACCGTCTATC
AGGGCGATGGCCCACTACGTGAACCATCACCCTAATCAAGTTTTTTGGGGTCGAGGTGCCGTAAA
GCACTAAATCGGAACCCTAAAGGGAGCCCCCGATTTAGAGCTTGACGGGGAAAGCCGGCGAACG
TGGCGAGAAAGGAAGGGAAGAAAGCGAAAGGAGCGGGCGCTAGGGCGCTGGCAAGTGTAGCGG
TCACGCTGCGCGTAACCACCACACCCGCCGCGCTTAATGCGCCGCTACAGGGCGCGTCGCGCCAT
TCGCCATTCAGGCTGCGCAACTGTTGGGAAGGGCGATCGGTGCGGGCCTCTTCGCTATTACGCCA
GCTGAATTGGAGCGACCTCATGCTATACCTGAGAAAGCAACCTGACCTACAGGAAAGAGTTACT
CAAGAATAAGAATTTTCGTTTTAAAACCTAAGAGTCACTTTAAAATTTGTATACACTTATTTTTTT
TATAACTTATTTAATAATAAAAATCATAAATCATAAGAAATTCGCTTATTTAGAAGTGTCAACAA
CGTATCTACCAACGATTTGACCCTTTTCCATCTTTTCGTAAATTTCTGGCAAGGTAGACAAGCCGA
CAACCTTGATTGGAGACTTGACCAAACCTCTGGCGAAGAATTGTTAATTAAGAGCTCAGATCTTA
TCGTCGTCATCCTTGTAATCCATCGATACTAGTGCTTACTTGTACAGCTCGTCCATGCCGAGAGTG
ATCCCGGCGGCGGTCACGAACTCCAGCAGGACCATGTGATCGCGCTTCTCGTTGGGGTCTTTGCT
CAGCTTGGACTGGTAGCTCAGGTAGTGGTTGTCGGGCAGCAGCACGGGGCCGTCGCCGATGGGG
GTGTTCTGCTGGTAGTGGTCGGCGAGCTGCACGCCGCCGTCCTCGATGTTGTGGCGGATCTTGAA
GTTGGCCTTGATGCCGTTCTTCTGCTTGTCCATAGTACCACCAGAACCAGGAGCTTTCAAGGTCTT
TGCAACGTCAAGGACAAAGCGATATCTAACATCACCCTTGGCCAGACGCTCCAAAGCTGTGTTGA
TATTGTCCGCGGAAATGAGTTCTATATCTGCAGTAATGTTATGCTTTCCGGCAAAATCAAGCATCT
CTTGGGTCTCCTTCAATCCTCCAATGCTACTTCCAGCAACCATCTTCCTCCCCATAAGCAAAGGAG
CAGAGTGAAGATCAAGTGGTTCCGGGGGTGCCCCAAGAAGAACAAGCTTGCCATGAGACTTTAG
TAGACCAAGCAATGGGATAATAGCGTGCTTAGCAGAAATAGTATTAAGTATGCCATCAAATGTG
CCTGTTGCAGCCTGCAGTGCTTCTGGATTACTGCTCAACAAAAATGCATCTGCACCCAAACGATT
GAGGGCATCGTCTTTCTTGCCCTCAGATGTACTTATAACTGTCACTTTTGCTCCCATGGCCTTTGC
AAACTTAACAGCCACATGGCCAAGTCCACCAAGACCATTAACAGCTATGTGGCTCCCGGGTTTGG
CAAAGCCATAGTATTTCATTGGACTGTACACAGTAATACCGGCACAAAGCAATGGTGCCCCACCA
TCAAGTGGTAGGTTCTCTGGGAAACGAACAATAAAGTGTTCATTGCATACCATCTCATTGGAATA
GCCTCCATAGGTAATCGTTCCATCAACGTTTGGACTTGCATAGGTTAGCACCATTTTGGGACAAT
AGTTCTCAAGATCTGCACGACAATTATCACAAGTGCGGCATGAACCAACCAAGCAGCCAACACC

AACTTTATCTCCAACCTTGACCTTTGTAACTTTGCCGCCGACCTCTGTAACTTCCCCTACGATTTCA
TGTCCTGGTACAAGAGGATAGGTCGAAATACCCCACTCATTCTTAGCGAAATGAAGGTCAGTATG
ACAAATCCCACAATATAGCACCTTGAACCTCACATCATCCTCAAGAGTTGCCCTCCTGGAGAATT
TGAACGGAGATAAAACCCCAGATGAATCATGAGCAGCCAATCCATAGGTCTTGACTGGGTGCTC
CTCTTCTGGTGATTTTCCGGCCATGCCCTTTAGTGAGGGTTGAATTCGAATTTTCAAAAATTCTTA
CTTTTTTTTTGGATGGACGCAAAGAAGTTTAATAATCATATTACATGGCATTACCACCATATACAT
ATCCATATACATATCCATATCTAATCTTACTTATATGTTGTGGAAATGTAAAGAGCCCCATTATCT
TAGCCTAAAAAAACCTTCTCTTTGGAACTTTCAGTAATACGCTTAACTGCTCATTGCTATATTGAA
GTACGGATTAGAAGCCGCCGAGCGGGTGACAGCCCTCCGAAGGAAGACTCTCCTCCGTGCGTCCT
CGTCTTCACCGGTCGCGTTCCTGAAACGCAGATGTGCCTCGCGCCGCACTGCTCCGAACAATAAA
GATTCTACAATACTAGCTTTTATGGTTATGAAGAGGAAAAATTGGCAGTAACCTGGCCCCACAAA
CCTTCAAATGAACGAATCAAATTAACAACCATAGGATGATAATGCGATTAGTTTTTTAGCCTTAT
TTCTGGGGTAATTAATCAGCGAAGCGATGATTTTTGATCTATTAACAGATATATAAATGCAAAAA
CTGCATAACCACTTTAACTAATACTTTCAACATTTTCGGTTTGTATTACTTCTTATTCAAATGTAAT
AAAAGTATCAACAAAAAATTGTTAATATACCTCTATACTTTAACGTCAAGGAGAAAAAACCCCG
GATCCGTAATACGACTCACTATAGGGCCCGGGCGTCGACATGGAACAGAAGTTGATTTCCGAAG
AAGACCTCGAGTAAGCTTGGTACCGCGGCTAGCTAAGATCCGCTCTAACCGAAAAGGAAGGAGT
TAGACAACCTGAAGTCTAGGTCCCTATTTATTTTTTTATAGTTATGTTAGTATTAAGAACGTTATT
TATATTTCAAATTTTTCTTTTTTTTCTGTACAGACGCGTGTACGCATGTAACATTATACTGAAAAC
CTTGCTTGAGAAGGTTTTGGGACGCTCGAAGATCCAGCTGCATTAATGAATCGGCCAACGCGCGG
GGAGAGGCGGTTTGCGTATTGGGCGCTCTTCCGCTTCCTCGCTCACTGACTCGCTGCGCTCGGTCG
TTCGGCTGCGGCGAGCGGTATCAGCTCACTCAAAGGCGGTAATACGGTTATCCACAGAATCAGG
GGATAACGCAGGAAAGAACATGTGAGCAAAAGGCCAGCAAAAGGCCAGGAACCGTAAAAAGGC
CGCGTTGCTGGCGTTTTTCCATAGGCTCCGCCCCCCTGACGAGCATCACAAAAATCGACGCTCAA
GTCAGAGGTGGCGAAACCCGACAGGACTATAAAGATACCAGGCGTTTCCCCCTGGAAGCTCCCT
CGTGCGCTCTCCTGTTCCGACCCTGCCGCTTACCGGATACCTGTCCGCCTTTCTCCCTTCGGGAAG
CGTGGCGCTTTCTCATAGCTCACGCTGTAGGTATCTCAGTTCGGTGTAGGTCGTTCGCTCCAAGCT
GGGCTGTGTGCACGAACCCCCCGTTCAGCCCGACCGCTGCGCCTTATCCGGTAACTATCGTCTTG
AGTCCAACCCGGTAAGACACGACTTATCGCCACTGGCAGCAGCCACTGGTAACAGGATTAGCAG
AGCGAGGTATGTAGGCGGTGCTACAGAGTTCTTGAAGTGGTGGCCTAACTACGGCTACACTAGA
AGGACAGTATTTGGTATCTGCGCTCTGCTGAAGCCAGTTACCTTCGGAAAAAGAGTTGGTAGCTC
TTGATCCGGCAAACAAACCACCGCTGGTAGCGGTGGTTTTTTTGTTTGCAAGCAGCAGATTACGC
GCAGAAAAAAAGGATCTCAAGAAGATCCTTTGATCTTTTCTACGGGGTCTGACGCTCAGTGGAAC
GAAAACTCACGTTAAGGGATTTTGGTCATGAGATTATCAAAAAGGATCTTCACCTAGATCCTTTT
AAATTAAAAATGAAGTTTTAAATCAATCTAAAGTATATATGAGTAAACTTGGTCTGACAGTTACC
AATGCTTAATCAGTGAGGCACCTATCTCAGCGATCTGTCTATTTCGTTCATCCATAGTTGCCTGAC
TCCCCGTCGTGTAGATAACTACGATACGGGAGGGCTTACCATCTGGCCCCAGTGCTGCAATGATA
CCGCGAGACCCACGCTCACCGGCTCCAGATTTATCAGCAATAAACCAGCCAGCCGGAAGGGCCG
AGCGCAGAAGTGGTCCTGCAACTTTATCCGCCTCCATCCAGTCTATTAATTGTTGCCGGGAAGCT
AGAGTAAGTAGTTCGCCAGTTAATAGTTTGCGCAACGTTGTTGCCATTGCTACAGGCATCGTGGT
GTCACGCTCGTCGTTTGGTATGGCTTCATTCAGCTCCGGTTCCCAACGATCAAGGCGAGTTACATG
ATCCCCCATGTTGTGCAAAAAAGCGGTTAGCTCCTTCGGTCCTCCGATCGTTGTCAGAAGTAAGT
TGGCCGCAGTGTTATCACTCATGGTTATGGCAGCACTGCATAATTCTCTTACTGTCATGCCATCCG
TAAGATGCTTTTCTGTGACTGGTGAGTACTCAACCAAGTCATTCTGAGAATAGTGTATGCGGCGA
CCGAGTTGCTCTTGCCCGGCGTCAATACGGGATAATACCGCGCCACATAGCAGAACTTTAAAAGT
GCTCATCATTGGAAAACGTTCTTCGGGGCGAAAACTCTCAAGGATCTTACCGCTGTTGAGATCCA
GTTCGATGTAACCCACTCGTGCACCCAACTGATCTTCAGCATCTTTTACTTTCACCAGCGTTTCTG
GGTGAGCAAAAACAGGAAGGCAAAATGCCGCAAAAAAGGGAATAAGGGCGACACGGAAATGTT
GAATACTCATACTCTTCCTTTTTCAATATTATTGAAGCATTTATCAGGGTTATTGTCTCATGAGCG
GATACATATTTGAATGTATTTAGAAAAATAAACAAATAGGGGTTCCGCGCACATTTCCCCGAAAA
GTGCCACCTGAACGAAGCATCTGTGCTTCATTTTGTAGAACAAAAATGCAACGCGAGAGCGCTAA
TTTTTCAAACAAAGAATCTGAGCTGCATTTTTACAGAACAGAAATGCAACGCGAAAGCGCTATTT
TACCAACGAAGAATCTGTGCTTCATTTTTGTAAAACAAAAATGCAACGCGAGAGCGCTAATTTTT
CAAACAAAGAATCTGAGCTGCATTTTTACAGAACAGAAATGCAACGCGAGAGCGCTATTTTACC
AACAAAGAATCTATACTTCTTTTTTGTTCTACAAAAATGCATCCCGAGAGCGCTATTTTTCTAACA
AAGCATCTTAGATTACTTTTTTTCTCCTTTGTGCGCTCTATAATGCAGTCTCTTGATAACTTTTTGC

ACTGTAGGTCCGTTAAGGTTAGAAGAAGGCTACTTTGGTGTCTATTTTCTCTTCCATAAAAAAAG
CCTGACTCCACTTCCCGCGTTTACTGATTACTAGCGAAGCTGCGGGTGCATTTTTTCAAGATAAAG
GCATCCCCGATTATATTCTATACCGATGTGGATTGCGCATACTTTGTGAACAGAAAGTGATAGCG
TTGATGATTCTTCATTGGTCAGAAAATTATGAACGGTTTCTTCTATTTTGTCTCTATATACTACGTA
TAGGAAATGTTTACATTTTCGTATTGTTTTCGATTCACTCTATGAATAGTTCTTACTACAATTTTTT
TGTCTAAAGAGTAATACTAGAGATAAACATAAAAAATGTAGAGGTCGAGTTTAGATGCAAGTTC
AAGGAGCGAAAGGTGGATGGGTAGGTTATATAGGGATATAGCACAGAGATATATAGCAAAGAG
ATACTTTTGAGCAATGTTTGTGGAAGCGGTATTCGCAATATTTTAGTAGCTCGTTACAGTCCGGTG
CGTTTTTGGTTTTTTGAAAGTGCGTCTTCAGAGCGCTTTTGGTTTTCAAAAGCGCTCTGAAGTTCC
TATACTTTCTAGAGAATAGGAACTTCGGAATAGGAACTTCAAAGCGTTTCCGAAAACGAGCGCTT
CCGAAAATGCAACGCGAGCTGCGCACATACAGCTCACTGTTCACGTCGCACCTATATCTGCGTGT
TGCCTGTATATATATATACATGAGAAGAACGGCATAGTGCGTGTTTATGCTTAAATGCGTACTTA
TATGCGTCTATTTATGTAGGATGAAAGGTAGTCTAGTACCTCCTGTGATATTATCCCATTCCATGC
GGGGTATCGTATGCTTCCTTCAGCACTACCCTTTAGCTGTTCTATATGCTGCCACTCCTCAATTGG
ATTAGTCTCATCCTTCAATGCTATCATTTCCTTTGATATTGGATCATCTAAGAAACCATTATTATC
ATGACATTAACCTATAAAAATAGGCGTATCACGAGGCCCTTTCGTC

> pESC-HIS-*CrVinBLAST*$^{C51A,\ H56A}$~*mVenus*$^{C}$

TCGCGCGTTTCGGTGATGACGGTGAAAACCTCTGACACATGCAGCTCCCGGAGACGGTCACAGCTTGT
CTGTAAGCGGATGCCGGGAGCAGACAAGCCCGTCAGGGCGCGTCAGCGGGTGTTGGCGGGTGTCGGG
GCTGGCTTAACTATGCGGCATCAGAGCAGATTGTACTGAGAGTGCACCATAAATTCCCGTTTTAAGAG
CTTGGTGAGCGCTAGGAGTCACTGCCAGGTATCGTTTGAACACGGCATTAGTCAGGGAAGTCATAACA
CAGTCCTTTCCCGCAATTTTCTTTTTCTATTACTCTTGGCCTCCTCTAGTACACTCTATATTTTTTTATGC
CTCGGTAATGATTTTCATTTTTTTTTTTCCCCTAGCGGATGACTCTTTTTTTTTCTTAGCGATTGGCATTA
TCACATAATGAATTATACATTATATAAAGTAATGTGATTTCTTCGAAGAATATACTAAAAAATGAGCA
GGCAAGATAAACGAAGGCAAAGATGACAGAGCAGAAAGCCCTAGTAAAGCGTATTACAAATGAAAC
CAAGATTCAGATTGCGATCTCTTTAAAGGGTGGTCCCCTAGCGATAGAGCACTCGATCTTCCCAGAAA
AAGAGGCAGAAGCAGTAGCAGAACAGGCCACACAATCGCAAGTGATTAACGTCCACACAGGTATAGG
GTTTCTGGACCATATGATACATGCTCTGGCCAAGCATTCCGGCTGGTCGCTAATCGTTGAGTGCATTGG
TGACTTACACATAGACGACCATCACACCACTGAAGACTGCGGGATTGCTCTCGGTCAAGCTTTTAAAG
AGGCCCTACTGGCGCGTGGAGTAAAAAGGTTTGGATCAGGATTTGCGCCTTTGGATGAGGCACTTTCC
AGAGCGGTGGTAGATCTTTCGAACAGGCCGTACGCAGTTGTCGAACTTGGTTTGCAAAGGGAGAAAGT
AGGAGATCTCTCTTGCGAGATGATCCCGCATTTTCTTGAAAGCTTTGCAGAGGCTAGCAGAATTACCCT
CCACGTTGATTGTCTGCGAGGCAAGAATGATCATCACCGTAGTGAGAGTGCGTTCAAGGCTCTTGCGG
TTGCCATAAGAGAAGCCACCTCGCCCAATGGTACCAACGATGTTCCCTCCACCAAAGGTGTTCTTATGT
AGTGACACCGATTATTTAAAGCTGCAGCATACGATATATATACATGTGTATATATGTATACCTATGAAT
GTCAGTAAGTATGTATACGAACAGTATGATACTGAAGATGACAAGGTAATGCATCATTCTATACGTGT
CATTCTGAACGAGGCGCGCTTTCCTTTTTTCTTTTTGCTTTTTCTTTTTTTTTCTCTTGAACTCGACGGAT
CTATGCGGTGTGAAATACCGCACAGATGCGTAAGGAGAAAATACCGCATCAGGAAATTGTAAACGTT
AATATTTTGTTAAAATTCGCGTTAAATTTTTGTTAAATCAGCTCATTTTTTAACCAATAGGCCGAAATC
GGCAAAATCCCTTATAAATCAAAAGAATAGACCGAGATAGGGTTGAGTGTTGTTCCAGTTTGGAACAA
GAGTCCACTATTAAAGAACGTGGACTCCAACGTCAAAGGGCGAAAAACCGTCTATCAGGGCGATGGC
CCACTACGTGAACCATCACCCTAATCAAGTTTTTTGGGGTCGAGGTGCCGTAAAGCACTAAATCGGAA
CCCTAAAGGGAGCCCCCGATTTAGAGCTTGACGGGGAAAGCCGGCGAACGTGGCGAGAAAGGAAGGG
AAGAAAGCGAAAGGAGCGGGCGCTAGGGCGCTGGCAAGTGTAGCGGTCACGCTGCGCGTAACCACCA
CACCCGCCGCGCTTAATGCGCCGCTACAGGGCGCGTCGCGCCATTCGCCATTCAGGCTGCGCAACTGT
TGGGAAGGGCGATCGGTGCGGGCCTCTTCGCTATTACGCCAGCTGAATTGGAGCGACCTCATGCTATA
CCTGAGAAAGCAACCTGACCTACAGGAAAGAGTTACTCAAGAATAAGAATTTTCGTTTTAAAACCTAA
GAGTCACTTTAAAATTTGTATACACTTATTTTTTTTATAACTTATTTAATAATAAAAATCATAAATCATA
AGAAATTCGCTTATTTAGAAGTGTCAACAACGTATCTACCAACGATTTGACCCTTTTCCATCTTTTCGT
AAATTTCTGGCAAGGTAGACAAGCCGACAACCTTGATTGGAGACTTGACCAAACCTCTGGCGAAGAAT
TGTTAATTAAGAGCTCAGATCTTATCGTCGTCATCCTTGTAATCCATCGATACTAGTGCTTACTTGTAC
AGCTCGTCCATGCCGAGAGTGATCCCGGCGGCGGTCACGAACTCCAGCAGGACCATGTGATCGCGCTT
CTCGTTGGGGTCTTTGCTCAGCTTGGACTGGTAGCTCAGGTAGTGGTTGTCGGGCAGCAGCACGGGGC
CGTCGCCGATGGGGGTGTTCTGCTGGTAGTGGTCGGCGAGCTGCACGCCGCCGTCCTCGATGTTGTGG

CGGATCTTGAAGTTGGCCTTGATGCCGTTCTTCTGCTTGTCCATAGTACCACCAGAACCAGGAGCTTTC
AAGGTCTTTGCAACGTCAAGGACAAAGCGATATCTAACATCACCCTTGGCCAGACGCTCCAAAGCTGT
GTTGATATTGTCCGCGGAAATGAGTTCTATATCTGCAGTAATGTTATGCTTTCCGGCAAAATCAAGCAT
CTCTTGGGTCTCCTTCAATCCTCCAATGCTACTTCCAGCAACCATCTTCCTCCCCATAAGCAAAGGAGC
AGAGTGAAGATCAAGTGGTTCCGGGGGTGCCCCAAGAAGAACAAGCTTGCCATGAGACTTTAGTAGA
CCAAGCAATGGGATAATAGCGTGCTTAGCAGAAATAGTATTAAGTATGCCATCAAATGTGCCTGTTGC
AGCCTGCAGTGCTTCTGGATTACTGCTCAACAAAAATGCATCTGCACCCAAACGATTGAGGGCATCGT
CTTTCTTGCCCTCAGATGTACTTATAACTGTCACTTTTGCTCCCATGGCCTTTGCAAACTTAACAGCCAC
ATGGCCAAGTCCACCAAGACCATTAACAGCTATGTGGCTCCCGGGTTTGGCAAAGCCATAGTATTTCA
TTGGACTGTACACAGTAATACCGGCACAAAGCAATGGTGCCCCACCATCAAGTGGTAGGTTCTCTGGG
AAACGAACAATAAAGTGTTCATTGCATACCATCTCATTGGAATAGCCTCCATAGGTAATCGTTCCATC
AACGTTTGGACTTGCATAGGTTAGCACCATTTTGGGACAATAGTTCTCAAGATCTGCACGACAATTATC
ACAAGTGCGGCATGAACCAACCAAGCAGCCAACACCAACTTTATCTCCAACCTTGACCTTTGTAACTT
TGCCGCCGACCTCTGTAACTTCCCCTACGATTTCATGTCCTGGTACAAGAGGATAGGTCGAAATACCCC
ACTCATTCTTAGCGAAAGCAAGGTCAGTATGAGCAATCCCACAATATAGCACCTTGAACCTCACATCA
TCCTCAAGAGTTGCCCTCCTGGAGAATTTGAACGGAGATAAAACCCCAGATGAATCATGAGCAGCCAA
TCCATAGGTCTTGACTGGGTGCTCCTCTTCTGGTGATTTTCCGGCCATGCCCTTTAGTGAGGGTTGAATT
CGAATTTTCAAAAATTCTTACTTTTTTTTTGGATGGACGCAAAGAAGTTTAATAATCATATTACATGGC
ATTACCACCATATACATATCCATATACATATCCATATCTAATCTTACTTATATGTTGTGGAAATGTAAA
GAGCCCCATTATCTTAGCCTAAAAAAACCTTCTCTTTGGAACTTTCAGTAATACGCTTAACTGCTCATT
GCTATATTGAAGTACGGATTAGAAGCCGCCGAGCGGGTGACAGCCCTCCGAAGGAAGACTCTCCTCCG
TGCGTCCTCGTCTTCACCGGTCGCGTTCCTGAAACGCAGATGTGCCTCGCGCCGCACTGCTCCGAACAA
TAAAGATTCTACAATACTAGCTTTTATGGTTATGAAGAGGAAAAATTGGCAGTAACCTGGCCCCACAA
ACCTTCAAATGAACGAATCAAATTAACAACCATAGGATGATAATGCGATTAGTTTTTTAGCCTTATTTC
TGGGGTAATTAATCAGCGAAGCGATGATTTTTGATCTATTAACAGATATATAAATGCAAAAACTGCAT
AACCACTTTAACTAATACTTTCAACATTTTCGGTTTGTATTACTTCTTATTCAAATGTAATAAAAGTATC
AACAAAAAATTGTTAATATACCTCTATACTTTAACGTCAAGGAGAAAAAACCCCGGATCCGTAATACG
ACTCACTATAGGGCCCGGGCGTCGACATGGAACAGAAGTTGATTTCCGAAGAAGACCTCGAGTAAGCT
TGGTACCGCGGCTAGCTAAGATCCGCTCTAACCGAAAAGGAAGGAGTTAGACAACCTGAAGTCTAGG
TCCCTATTTATTTTTTTATAGTTATGTTAGTATTAAGAACGTTATTTATATTTCAAATTTTTCTTTTTTTT
CTGTACAGACGCGTGTACGCATGTAACATTATACTGAAAACCTTGCTTGAGAAGGTTTTGGGACGCTC
GAAGATCCAGCTGCATTAATGAATCGGCCAACGCGCGGGGAGAGGCGGTTTGCGTATTGGGCGCTCTT
CCGCTTCCTCGCTCACTGACTCGCTGCGCTCGGTCGTTCGGCTGCGGCGAGCGGTATCAGCTCACTCAA
AGGCGGTAATACGGTTATCCACAGAATCAGGGGATAACGCAGGAAAGAACATGTGAGCAAAAGGCCA
GCAAAAGGCCAGGAACCGTAAAAAGGCCGCGTTGCTGGCGTTTTTCCATAGGCTCCGCCCCCCTGACG
AGCATCACAAAAATCGACGCTCAAGTCAGAGGTGGCGAAACCCGACAGGACTATAAAGATACCAGGC
GTTTCCCCCTGGAAGCTCCCTCGTGCGCTCTCCTGTTCCGACCCTGCCGCTTACCGGATACCTGTCCGC
CTTTCTCCCTTCGGGAAGCGTGGCGCTTTCTCATAGCTCACGCTGTAGGTATCTCAGTTCGGTGTAGGT
CGTTCGCTCCAAGCTGGGCTGTGTGCACGAACCCCCCGTTCAGCCCGACCGCTGCGCCTTATCCGGTAA
CTATCGTCTTGAGTCCAACCCGGTAAGACACGACTTATCGCCACTGGCAGCAGCCACTGGTAACAGGA
TTAGCAGAGCGAGGTATGTAGGCGGTGCTACAGAGTTCTTGAAGTGGTGGCCTAACTACGGCTACACT
AGAAGGACAGTATTTGGTATCTGCGCTCTGCTGAAGCCAGTTACCTTCGGAAAAAGAGTTGGTAGCTC
TTGATCCGGCAAACAAACCACCGCTGGTAGCGGTGGTTTTTTTGTTTGCAAGCAGCAGATTACGCGCA
GAAAAAAAGGATCTCAAGAAGATCCTTTGATCTTTTCTACGGGGTCTGACGCTCAGTGGAACGAAAAC
TCACGTTAAGGGATTTTGGTCATGAGATTATCAAAAAGGATCTTCACCTAGATCCTTTTAAATTAAAAA
TGAAGTTTTAAATCAATCTAAAGTATATATGAGTAAACTTGGTCTGACAGTTACCAATGCTTAATCAGT
GAGGCACCTATCTCAGCGATCTGTCTATTTCGTTCATCCATAGTTGCCTGACTCCCCGTCGTGTAGATA
ACTACGATACGGGAGGGCTTACCATCTGGCCCCAGTGCTGCAATGATACCGCGAGACCCACGCTCACC
GGCTCCAGATTTATCAGCAATAAACCAGCCAGCCGGAAGGGCCGAGCGCAGAAGTGGTCCTGCAACT
TTATCCGCCTCCATCCAGTCTATTAATTGTTGCCGGGAAGCTAGAGTAAGTAGTTCGCCAGTTAATAGT
TTGCGCAACGTTGTTGCCATTGCTACAGGCATCGTGGTGTCACGCTCGTCGTTTGGTATGGCTTCATTC
AGCTCCGGTTCCCAACGATCAAGGCGAGTTACATGATCCCCCATGTTGTGCAAAAAAGCGGTTAGCTC
CTTCGGTCCTCCGATCGTTGTCAGAAGTAAGTTGGCCGCAGTGTTATCACTCATGGTTATGGCAGCACT
GCATAATTCTCTTACTGTCATGCCATCCGTAAGATGCTTTTCTGTGACTGGTGAGTACTCAACCAAGTC
ATTCTGAGAATAGTGTATGCGGCGACCGAGTTGCTCTTGCCCGGCGTCAATACGGGATAATACCGCGC
CACATAGCAGAACTTTAAAAGTGCTCATCATTGGAAAACGTTCTTCGGGGCGAAAACTCTCAAGGATC

TTACCGCTGTTGAGATCCAGTTCGATGTAACCCACTCGTGCACCCAACTGATCTTCAGCATCTTTTACT
TTCACCAGCGTTTCTGGGTGAGCAAAAACAGGAAGGCAAAATGCCGCAAAAAAGGGAATAAGGGCGA
CACGGAAATGTTGAATACTCATACTCTTCCTTTTTCAATATTATTGAAGCATTTATCAGGGTTATTGTCT
CATGAGCGGATACATATTTGAATGTATTTAGAAAAATAAACAAATAGGGGTTCCGCGCACATTTCCCC
GAAAAGTGCCACCTGAACGAAGCATCTGTGCTTCATTTTGTAGAACAAAAATGCAACGCGAGAGCGCT
AATTTTTCAAACAAAGAATCTGAGCTGCATTTTTACAGAACAGAAATGCAACGCGAAAGCGCTATTTT
ACCAACGAAGAATCTGTGCTTCATTTTTGTAAAACAAAAATGCAACGCGAGAGCGCTAATTTTTCAAA
CAAAGAATCTGAGCTGCATTTTTACAGAACAGAAATGCAACGCGAGAGCGCTATTTTACCAACAAAGA
ATCTATACTTCTTTTTTGTTCTACAAAAATGCATCCCGAGAGCGCTATTTTTCTAACAAAGCATCTTAG
ATTACTTTTTTTCTCCTTTGTGCGCTCTATAATGCAGTCTCTTGATAACTTTTTGCACTGTAGGTCCGTTA
AGGTTAGAAGAAGGCTACTTTGGTGTCTATTTTCTCTTCCATAAAAAAAGCCTGACTCCACTTCCCGCG
TTTACTGATTACTAGCGAAGCTGCGGGTGCATTTTTTCAAGATAAAGGCATCCCCGATTATATTCTATA
CCGATGTGGATTGCGCATACTTTGTGAACAGAAAGTGATAGCGTTGATGATTCTTCATTGGTCAGAAA
ATTATGAACGGTTTCTTCTATTTTGTCTCTATATACTACGTATAGGAAATGTTTACATTTTCGTATTGTT
TTCGATTCACTCTATGAATAGTTCTTACTACAATTTTTTTGTCTAAAGAGTAATACTAGAGATAAACAT
AAAAAATGTAGAGGTCGAGTTTAGATGCAAGTTCAAGGAGCGAAAGGTGGATGGGTAGGTTATATAG
GGATATAGCACAGAGATATATAGCAAAGAGATACTTTTGAGCAATGTTTGTGGAAGCGGTATTCGCAA
TATTTTAGTAGCTCGTTACAGTCCGGTGCGTTTTTGGTTTTTTGAAAGTGCGTCTTCAGAGCGCTTTTGG
TTTTCAAAAGCGCTCTGAAGTTCCTATACTTTCTAGAGAATAGGAACTTCGGAATAGGAACTTCAAAG
CGTTTCCGAAAACGAGCGCTTCCGAAAATGCAACGCGAGCTGCGCACATACAGCTCACTGTTCACGTC
GCACCTATATCTGCGTGTTGCCTGTATATATATATACATGAGAAGAACGGCATAGTGCGTGTTTATGCT
TAAATGCGTACTTATATGCGTCTATTTATGTAGGATGAAAGGTAGTCTAGTACCTCCTGTGATATTATC
CCATTCCATGCGGGGTATCGTATGCTTCCTTCAGCACTACCCTTTAGCTGTTCTATATGCTGCCACTCCT
CAATTGGATTAGTCTCATCCTTCAATGCTATCATTTCCTTTGATATTGGATCATCTAAGAAACCATTATT
ATCATGACATTAACCTATAAAAATAGGCGTATCACGAGGCCCTTTCGTC

> pESC-HIS-*CrVinBLAST*$^{M298E, V299E}$~*mVenus*$^{C}$

TCGCGCGTTTCGGTGATGACGGTGAAAACCTCTGACACATGCAGCTCCCGGAGACGGTCACAGCT
TGTCTGTAAGCGGATGCCGGGAGCAGACAAGCCCGTCAGGGCGCGTCAGCGGGTGTTGGCGGGT
GTCGGGGCTGGCTTAACTATGCGGCATCAGAGCAGATTGTACTGAGAGTGCACCATAAATTCCCG
TTTTAAGAGCTTGGTGAGCGCTAGGAGTCACTGCCAGGTATCGTTTGAACACGGCATTAGTCAGG
GAAGTCATAACACAGTCCTTTCCCGCAATTTTCTTTTTCTATTACTCTTGGCCTCCTCTAGTACACT
CTATATTTTTTTATGCCTCGGTAATGATTTTCATTTTTTTTTTTCCCCTAGCGGATGACTCTTTTTTT
TTCTTAGCGATTGGCATTATCACATAATGAATTATACATTATATAAAGTAATGTGATTTCTTCGAA
GAATATACTAAAAAATGAGCAGGCAAGATAAACGAAGGCAAAGATGACAGAGCAGAAAGCCCT
AGTAAAGCGTATTACAAATGAAACCAAGATTCAGATTGCGATCTCTTTAAAGGGTGGTCCCCTAG
CGATAGAGCACTCGATCTTCCCAGAAAAAGAGGCAGAAGCAGTAGCAGAACAGGCCACACAATC
GCAAGTGATTAACGTCCACACAGGTATAGGGTTTCTGGACCATATGATACATGCTCTGGCCAAGC
ATTCCGGCTGGTCGCTAATCGTTGAGTGCATTGGTGACTTACACATAGACGACCATCACACCACT
GAAGACTGCGGGATTGCTCTCGGTCAAGCTTTTAAAGAGGCCCTACTGGCGCGTGGAGTAAAAA
GGTTTGGATCAGGATTTGCGCCTTTGGATGAGGCACTTTCCAGAGCGGTGGTAGATCTTTCGAAC
AGGCCGTACGCAGTTGTCGAACTTGGTTTGCAAAGGGAGAAAGTAGGAGATCTCTCTTGCGAGAT
GATCCCGCATTTTCTTGAAAGCTTTGCAGAGGCTAGCAGAATTACCCTCCACGTTGATTGTCTGCG
AGGCAAGAATGATCATCACCGTAGTGAGAGTGCGTTCAAGGCTCTTGCGGTTGCCATAAGAGAA
GCCACCTCGCCCAATGGTACCAACGATGTTCCCTCCACCAAAGGTGTTCTTATGTAGTGACACCG
ATTATTTAAAGCTGCAGCATACGATATATATACATGTGTATATATGTATACCTATGAATGTCAGTA
AGTATGTATACGAACAGTATGATACTGAAGATGACAAGGTAATGCATCATTCTATACGTGTCATT
CTGAACGAGGCGCGCTTTCCTTTTTTCTTTTTGCTTTTTCTTTTTTTTTTCTCTTGAACTCGACGGATC
TATGCGGTGTGAAATACCGCACAGATGCGTAAGGAGAAAATACCGCATCAGGAAATTGTAAACG
TTAATATTTTGTTAAAATTCGCGTTAAATTTTTGTTAAATCAGCTCATTTTTTAACCAATAGGCCG
AAATCGGCAAAATCCCTTATAAATCAAAAGAATAGACCGAGATAGGGTTGAGTGTTGTTCCAGTT
TGGAACAAGAGTCCACTATTAAAGAACGTGGACTCCAACGTCAAAGGGCGAAAAACCGTCTATC
AGGGCGATGGCCCACTACGTGAACCATCACCCTAATCAAGTTTTTTGGGGTCGAGGTGCCGTAAA
GCACTAAATCGGAACCCTAAAGGGAGCCCCCGATTTAGAGCTTGACGGGGAAAGCCGGCGAACG
TGGCGAGAAAGGAAGGGAAGAAAGCGAAAGGAGCGGGCGCTAGGGCGCTGGCAAGTGTAGCGG

TCACGCTGCGCGTAACCACCACACCCGCCGCGCTTAATGCGCCGCTACAGGGCGCGTCGCGCCAT
TCGCCATTCAGGCTGCGCAACTGTTGGGAAGGGCGATCGGTGCGGGCCTCTTCGCTATTACGCCA
GCTGAATTGGAGCGACCTCATGCTATACCTGAGAAAGCAACCTGACCTACAGGAAAGAGTTACT
CAAGAATAAGAATTTTCGTTTTAAAACCTAAGAGTCACTTTAAAATTTGTATACACTTATTTTTT
TATAACTTATTTAATAATAAAAATCATAAATCATAAGAAATTCGCTTATTTAGAAGTGTCAACAA
CGTATCTACCAACGATTTGACCCTTTTCCATCTTTTCGTAAATTTCTGGCAAGGTAGACAAGCCGA
CAACCTTGATTGGAGACTTGACCAAACCTCTGGCGAAGAATTGTTAATTAAGAGCTCAGATCTTA
TCGTCGTCATCCTTGTAATCCATCGATACTAGTGCTTACTTGTACAGCTCGTCCATGCCGAGAGTG
ATCCCGGCGGCGGTCACGAACTCCAGCAGGACCATGTGATCGCGCTTCTCGTTGGGGTCTTTGCT
CAGCTTGGACTGGTAGCTCAGGTAGTGGTTGTCGGGCAGCAGCACGGGGCCGTCGCCGATGGGG
GTGTTCTGCTGGTAGTGGTCGGCGAGCTGCACGCCGCCGTCCTCGATGTTGTGGCGGATCTTGAA
GTTGGCCTTGATGCCGTTCTTCTGCTTGTCCATAGTACCACCAGAACCAGGAGCTTTCAAGGTCTT
TGCAACGTCAAGGACAAAGCGATATCTAACATCACCCTTGGCCAGACGCTCCAAAGCTGTGTTGA
TATTGTCCGCGGAAATGAGTTCTATATCTGCAGTAATGTTATGCTTTCCGGCAAAATCAAGCATCT
CTTGGGTCTCCTTCAATCCTCCAATGCTACTTCCAGCTTCTTCCTTCCTCCCCATAAGCAAAGGAG
CAGAGTGAAGATCAAGTGGTTCCGGGGGTGCCCCAAGAAGAACAAGCTTGCCATGAGACTTTAG
TAGACCAAGCAATGGGATAATAGCGTGCTTAGCAGAAATAGTATTAAGTATGCCATCAAATGTG
CCTGTTGCAGCCTGCAGTGCTTCTGGATTACTGCTCAACAAAAATGCATCTGCACCCAAACGATT
GAGGGCATCGTCTTTCTTGCCCTCAGATGTACTTATAACTGTCACTTTTGCTCCCATGGCCTTTGC
AAACTTAACAGCCACATGGCCAAGTCCACCAAGACCATTAACAGCTATGTGGCTCCCGGGTTTGG
CAAAGCCATAGTATTTCATTGGACTGTACACAGTAATACCGGCACAAAGCAATGGTGCCCCACCA
TCAAGTGGTAGGTTCTCTGGGAAACGAACAATAAAGTGTTCATTGCATACCATCTCATTGGAATA
GCCTCCATAGGTAATCGTTCCATCAACGTTTGGACTTGCATAGGTTAGCACCATTTTGGGACAAT
AGTTCTCAAGATCTGCACGACAATTATCACAAGTGCGGCATGAACCAACCAAGCAGCCAACACC
AACTTTATCTCCAACCTTGACCTTTGTAACTTTGCCGCCGACCTCTGTAACTTCCCCTACGATTTCA
TGTCCTGGTACAAGAGGATAGGTCGAAATACCCCACTCATTCTTAGCGAAATGAAGGTCAGTATG
ACAAATCCCACAATATAGCACCTTGAACCTCACATCATCCTCAAGAGTTGCCCTCCTGGAGAATT
TGAACGGAGATAAAACCCCAGATGAATCATGAGCAGCCAATCCATAGGTCTTGACTGGGTGCTC
CTCTTCTGGTGATTTTCCGGCCATGCCCTTTAGTGAGGGTTGAATTCGAATTTTCAAAAATTCTTA
CTTTTTTTTTGGATGGACGCAAAGAAGTTTAATAATCATATTACATGGCATTACCACCATATACAT
ATCCATATACATATCCATATCTAATCTTACTTATATGTTGTGGAAATGTAAAGAGCCCCATTATCT
TAGCCTAAAAAAACCTTCTCTTTGGAACTTTCAGTAATACGCTTAACTGCTCATTGCTATATTGAA
GTACGGATTAGAAGCCGCCGAGCGGGTGACAGCCCTCCGAAGGAAGACTCTCCTCCGTGCGTCCT
CGTCTTCACCGGTCGCGTTCCTGAAACGCAGATGTGCCTCGCGCCGCACTGCTCCGAACAATAAA
GATTCTACAATACTAGCTTTTATGGTTATGAAGAGGAAAAATTGGCAGTAACCTGGCCCCACAAA
CCTTCAAATGAACGAATCAAATTAACAACCATAGGATGATAATGCGATTAGTTTTTTAGCCTTAT
TTCTGGGGTAATTAATCAGCGAAGCGATGATTTTTGATCTATTAACAGATATATAAATGCAAAAA
CTGCATAACCACTTTAACTAATACTTTCAACATTTTCGGTTTGTATTACTTCTTATTCAAATGTAAT
AAAAGTATCAACAAAAAATTGTTAATATACCTCTATACTTTAACGTCAAGGAGAAAAAACCCCG
GATCCGTAATACGACTCACTATAGGGCCCGGGCGTCGACATGGAACAGAAGTTGATTTCCGAAG
AAGACCTCGAGTAAGCTTGGTACCGCGGCTAGCTAAGATCCGCTCTAACCGAAAAGGAAGGAGT
TAGACAACCTGAAGTCTAGGTCCCTATTTATTTTTTTATAGTTATGTTAGTATTAAGAACGTTATT
TATATTTCAAATTTTTCTTTTTTTTCTGTACAGACGCGTGTACGCATGTAACATTATACTGAAAAC
CTTGCTTGAGAAGGTTTTGGGACGCTCGAAGATCCAGCTGCATTAATGAATCGGCCAACGCGCGG
GGAGAGGCGGTTTGCGTATTGGGCGCTCTTCCGCTTCCTCGCTCACTGACTCGCTGCGCTCGGTCG
TTCGGCTGCGGCGAGCGGTATCAGCTCACTCAAAGGCGGTAATACGGTTATCCACAGAATCAGG
GGATAACGCAGGAAAGAACATGTGAGCAAAAGGCCAGCAAAAGGCCAGGAACCGTAAAAAGGC
CGCGTTGCTGGCGTTTTTCCATAGGCTCCGCCCCCCTGACGAGCATCACAAAAATCGACGCTCAA
GTCAGAGGTGGCGAAACCCGACAGGACTATAAAGATACCAGGCGTTTCCCCCTGGAAGCTCCCT
CGTGCGCTCTCCTGTTCCGACCCTGCCGCTTACCGGATACCTGTCCGCCTTTCTCCCTTCGGGAAG
CGTGGCGCTTTCTCATAGCTCACGCTGTAGGTATCTCAGTTCGGTGTAGGTCGTTCGCTCCAAGCT
GGGCTGTGTGCACGAACCCCCCGTTCAGCCCGACCGCTGCGCCTTATCCGGTAACTATCGTCTTG
AGTCCAACCCGGTAAGACACGACTTATCGCCACTGGCAGCAGCCACTGGTAACAGGATTAGCAG
AGCGAGGTATGTAGGCGGTGCTACAGAGTTCTTGAAGTGGTGGCCTAACTACGGCTACACTAGA
AGGACAGTATTTGGTATCTGCGCTCTGCTGAAGCCAGTTACCTTCGGAAAAAGAGTTGGTAGCTC
TTGATCCGGCAAACAAACCACCGCTGGTAGCGGTGGTTTTTTTGTTTGCAAGCAGCAGATTACGC

GCAGAAAAAAAGGATCTCAAGAAGATCCTTTGATCTTTTCTACGGGGTCTGACGCTCAGTGGAAC
GAAAACTCACGTTAAGGGATTTTGGTCATGAGATTATCAAAAAGGATCTTCACCTAGATCCTTTT
AAATTAAAAATGAAGTTTTAAATCAATCTAAAGTATATATGAGTAAACTTGGTCTGACAGTTACC
AATGCTTAATCAGTGAGGCACCTATCTCAGCGATCTGTCTATTTCGTTCATCCATAGTTGCCTGAC
TCCCCGTCGTGTAGATAACTACGATACGGGAGGGCTTACCATCTGGCCCCAGTGCTGCAATGATA
CCGCGAGACCCACGCTCACCGGCTCCAGATTTATCAGCAATAAACCAGCCAGCCGGAAGGGCCG
AGCGCAGAAGTGGTCCTGCAACTTTATCCGCCTCCATCCAGTCTATTAATTGTTGCCGGGAAGCT
AGAGTAAGTAGTTCGCCAGTTAATAGTTTGCGCAACGTTGTTGCCATTGCTACAGGCATCGTGGT
GTCACGCTCGTCGTTTGGTATGGCTTCATTCAGCTCCGGTTCCCAACGATCAAGGCGAGTTACATG
ATCCCCCATGTTGTGCAAAAAAGCGGTTAGCTCCTTCGGTCCTCCGATCGTTGTCAGAAGTAAGT
TGGCCGCAGTGTTATCACTCATGGTTATGGCAGCACTGCATAATTCTCTTACTGTCATGCCATCCG
TAAGATGCTTTTCTGTGACTGGTGAGTACTCAACCAAGTCATTCTGAGAATAGTGTATGCGGCGA
CCGAGTTGCTCTTGCCCGGCGTCAATACGGGATAATACCGCGCCACATAGCAGAACTTTAAAAGT
GCTCATCATTGGAAAACGTTCTTCGGGGCGAAAACTCTCAAGGATCTTACCGCTGTTGAGATCCA
GTTCGATGTAACCCACTCGTGCACCCAACTGATCTTCAGCATCTTTTACTTTCACCAGCGTTTCTG
GGTGAGCAAAAACAGGAAGGCAAAATGCCGCAAAAAAGGGAATAAGGGCGACACGGAAATGTT
GAATACTCATACTCTTCCTTTTTCAATATTATTGAAGCATTTATCAGGGTTATTGTCTCATGAGCG
GATACATATTTGAATGTATTTAGAAAAATAAACAAATAGGGGTTCCGCGCACATTTCCCCGAAAA
GTGCCACCTGAACGAAGCATCTGTGCTTCATTTTGTAGAACAAAAATGCAACGCGAGAGCGCTAA
TTTTTCAAACAAAGAATCTGAGCTGCATTTTTACAGAACAGAAATGCAACGCGAAAGCGCTATTT
TACCAACGAAGAATCTGTGCTTCATTTTTGTAAAACAAAAATGCAACGCGAGAGCGCTAATTTTT
CAAACAAAGAATCTGAGCTGCATTTTTACAGAACAGAAATGCAACGCGAGAGCGCTATTTTACC
AACAAAGAATCTATACTTCTTTTTTGTTCTACAAAAATGCATCCCGAGAGCGCTATTTTTCTAACA
AAGCATCTTAGATTACTTTTTTTCTCCTTTGTGCGCTCTATAATGCAGTCTCTTGATAACTTTTTGC
ACTGTAGGTCCGTTAAGGTTAGAAGAAGGCTACTTTGGTGTCTATTTTCTCTTCCATAAAAAAAG
CCTGACTCCACTTCCCGCGTTTACTGATTACTAGCGAAGCTGCGGGTGCATTTTTTCAAGATAAAG
GCATCCCCGATTATATTCTATACCGATGTGGATTGCGCATACTTTGTGAACAGAAAGTGATAGCG
TTGATGATTCTTCATTGGTCAGAAAATTATGAACGGTTTCTTCTATTTTGTCTCTATATACTACGTA
TAGGAAATGTTTACATTTTCGTATTGTTTTCGATTCACTCTATGAATAGTTCTTACTACAATTTTTT
TGTCTAAAGAGTAATACTAGAGATAAACATAAAAAAATGTAGAGGTCGAGTTTAGATGCAAGTTC
AAGGAGCGAAAGGTGGATGGGTAGGTTATATAGGGATATAGCACAGAGATATATAGCAAAGAG
ATACTTTTGAGCAATGTTTGTGGAAGCGGTATTCGCAATATTTTAGTAGCTCGTTACAGTCCGGTG
CGTTTTTGGTTTTTTGAAAGTGCGTCTTCAGAGCGCTTTTGGTTTTCAAAAGCGCTCTGAAGTTCC
TATACTTTCTAGAGAATAGGAACTTCGGAATAGGAACTTCAAAGCGTTTCCGAAAACGAGCGCTT
CCGAAAATGCAACGCGAGCTGCGCACATACAGCTCACTGTTCACGTCGCACCTATATCTGCGTGT
TGCCTGTATATATATATACATGAGAAGAACGGCATAGTGCGTGTTTATGCTTAAATGCGTACTTA
TATGCGTCTATTTATGTAGGATGAAAGGTAGTCTAGTACCTCCTGTGATATTATCCCATTCCATGC
GGGGTATCGTATGCTTCCTTCAGCACTACCCTTTAGCTGTTCTATATGCTGCCACTCCTCAATTGG
ATTAGTCTCATCCTTCAATGCTATCATTTCCTTTGATATTGGATCATCTAAGAAACCATTATTATC
ATGACATTAACCTATAAAAATAGGCGTATCACGAGGCCCTTTCGTC

> pESC-HIS-*GS~mVenus*$^C$

TCGCGCGTTTCGGTGATGACGGTGAAAACCTCTGACACATGCAGCTCCCGGAGACGGTCACAGCT
TGTCTGTAAGCGGATGCCGGGAGCAGACAAGCCCGTCAGGGCGCGTCAGCGGGTGTTGGCGGGT
GTCGGGGCTGGCTTAACTATGCGGCATCAGAGCAGATTGTACTGAGAGTGCACCATAAATTCCCG
TTTTAAGAGCTTGGTGAGCGCTAGGAGTCACTGCCAGGTATCGTTTGAACACGGCATTAGTCAGG
GAAGTCATAACACAGTCCTTTCCCGCAATTTTCTTTTTCTATTACTCTTGGCCTCCTCTAGTACACT
CTATATTTTTTTATGCCTCGGTAATGATTTTCATTTTTTTTTTTCCCCTAGCGGATGACTCTTTTTTT
TTCTTAGCGATTGGCATTATCACATAATGAATTATACATTATATAAAGTAATGTGATTTCTTCGAA
GAATATACTAAAAAATGAGCAGGCAAGATAAACGAAGGCAAAGATGACAGAGCAGAAAGCCCT
AGTAAAGCGTATTACAAATGAAACCAAGATTCAGATTGCGATCTCTTTAAAGGGTGGTCCCCTAG
CGATAGAGCACTCGATCTTCCCAGAAAAAGAGGCAGAAGCAGTAGCAGAACAGGCCACACAATC
GCAAGTGATTAACGTCCACACAGGTATAGGGTTTCTGGACCATATGATACATGCTCTGGCCAAGC
ATTCCGGCTGGTCGCTAATCGTTGAGTGCATTGGTGACTTACACATAGACGACCATCACACCACT
GAAGACTGCGGGATTGCTCTCGGTCAAGCTTTTAAAGAGGCCCTACTGGCGCGTGGAGTAAAAA

GGTTTGGATCAGGATTTGCGCCTTTGGATGAGGCACTTTCCAGAGCGGTGGTAGATCTTTCGAAC
AGGCCGTACGCAGTTGTCGAACTTGGTTTGCAAAGGGAGAAAGTAGGAGATCTCTCTTGCGAGAT
GATCCCGCATTTTCTTGAAAGCTTTGCAGAGGCTAGCAGAATTACCCTCCACGTTGATTGTCTGCG
AGGCAAGAATGATCATCACCGTAGTGAGAGTGCGTTCAAGGCTCTTGCGGTTGCCATAAGAGAA
GCCACCTCGCCCAATGGTACCAACGATGTTCCCTCCACCAAAGGTGTTCTTATGTAGTGACACCG
ATTATTTAAAGCTGCAGCATACGATATATATACATGTGTATATATGTATACCTATGAATGTCAGTA
AGTATGTATACGAACAGTATGATACTGAAGATGACAAGGTAATGCATCATTCTATACGTGTCATT
CTGAACGAGGCGCGCTTTCCTTTTTTCTTTTTGCTTTTTCTTTTTTTTTCTCTTGAACTCGACGGATC
TATGCGGTGTGAAATACCGCACAGATGCGTAAGGAGAAAATACCGCATCAGGAAATTGTAAACG
TTAATATTTTGTTAAAATTCGCGTTAAATTTTTGTTAAATCAGCTCATTTTTTAACCAATAGGCCG
AAATCGGCAAAATCCCTTATAAATCAAAAGAATAGACCGAGATAGGGTTGAGTGTTGTTCCAGTT
TGGAACAAGAGTCCACTATTAAAGAACGTGGACTCCAACGTCAAAGGGCGAAAAACCGTCTATC
AGGGCGATGGCCCACTACGTGAACCATCACCCTAATCAAGTTTTTTGGGGTCGAGGTGCCGTAAA
GCACTAAATCGGAACCCTAAAGGGAGCCCCCGATTTAGAGCTTGACGGGGAAAGCCGGCGAACG
TGGCGAGAAAGGAAGGGAAGAAAGCGAAAGGAGCGGGCGCTAGGGCGCTGGCAAGTGTAGCGG
TCACGCTGCGCGTAACCACCACACCCGCCGCGCTTAATGCGCCGCTACAGGGCGCGTCGCGCCAT
TCGCCATTCAGGCTGCGCAACTGTTGGGAAGGGCGATCGGTGCGGGCCTCTTCGCTATTACGCCA
GCTGAATTGGAGCGACCTCATGCTATACCTGAGAAAGCAACCTGACCTACAGGAAAGAGTTACT
CAAGAATAAGAATTTTCGTTTTAAAACCTAAGAGTCACTTTAAAATTTGTATACACTTATTTTTTT
TATAACTTATTTAATAATAAAAATCATAAATCATAAGAAATTCGCTTATTTAGAAGTGTCAACAA
CGTATCTACCAACGATTTGACCCTTTTCCATCTTTTCGTAAATTTCTGGCAAGGTAGACAAGCCGA
CAACCTTGATTGGAGACTTGACCAAACCTCTGGCGAAGAATTGTTAATTAAGAGCTCAGATCTTA
TCGTCGTCATCCTTGTAATCCATCGATACTAGTGCTTACTTGTACAGCTCGTCCATGCCGAGAGTG
ATCCCGGCGGCGGTCACGAACTCCAGCAGGACCATGTGATCGCGCTTCTCGTTGGGGTCTTTGCT
CAGCTTGGACTGGTAGCTCAGGTAGTGGTTGTCGGGCAGCAGCACGGGGCCGTCGCCGATGGGG
GTGTTCTGCTGGTAGTGGTCGGCGAGCTGCACGCCGCCGTCCTCGATGTTGTGGCGGATCTTGAA
GTTGGCCTTGATGCCGTTCTTCTGCTTGTCCATAGTACCACCAGAACCTTCCTCAAATTTCAATGT
ATTTCCAATGTCAATCGCAAAACGGTACTTGACATCCAAATTCTTGATACGTTCCATAGCAGTGCT
GAGATAGTCAATCCCAATAACTTCAGTATCACATACAATGTTATGTTTGGCTGCGAAATCAAGCA
TTTCTTGGTACTCTTTGAGACCTCCAGTGGAACTTCCGATTATCTTTTTCCTTCCCATAATGAGAG
GTGCCGCAGGGAGCTCAAATAGCGACTCCGGTGCACCTACGAGCATAACCGCGCCGTCAAACTT
GAGCAAATTGAGCATAAGTGACATAGGAGTGCGGCCACCTGGGGTGGTGTCCACAACACCATCC
ATAGTACCTGCCAGAGCCTTCAATTGCTCAGAGTCAGTGTTGACAACAAAAGCATCAGCACCATG
TTCTTCAATGGCTTCCTTCTCCTTACGCCTTGATGTACTAATAACAGTAGCCTTACCACCAAAAGC
CTTAATAAACTTAACAGCAACAGAACCAAGACCTCCCAGCCCGAAAACCCCAATATGCTTTCCTG
GCTTATCGAGTCCCAAATGTTTCATTGGGCTATAAACAACAACCCCAGCACAAAGGAGAGCAAC
CCCTTTATCTTGAGGCAAGTTTTCGGGCCATCGGAGGACGAACTTTTCATCAACAACCATCACATT
TGAACAACCCCCATAGGATCGTTCCCCTTGCTCACGGTAAACAGATCCATCAGCCATATTGGGCT
CTGGGCAGTAATTCTCCATTCCACTTTGACAATTATAACATTGACCACAAGATCCGACCATACAT
CCCACAGCTACCTTGTCTCCAACCTTGAATTTCTCTACTTTGCTGCCAACTTCTACCACCTCACCG
GCAGTTTCATGTCCAAACACATAAGGATATCTGGTGAAACCCCACTTGTTTCTGACCATTTCCATA
TCGAAATTGCAAACACCAGAGTACAAAACTCTAATCTTCACATCCCGTTCACCAGGGACTCTTCT
ATAGAACTTGATGGGCTGAAGGACACCAGATGCATCTGCAGCACCCCATCCCACAGCCTTCACTG
AAAGGTCGAGTTTGGTTGTTTCTCCGGCCATGCCCTTTAGTGAGGGTTGAATTCGAATTTTCAAAA
ATTCTTACTTTTTTTTTGGATGGACGCAAAGAAGTTTAATAATCATATTACATGGCATTACCACCA
TATACATATCCATATACATATCCATATCTAATCTTACTTATATGTTGTGGAAATGTAAAGAGCCCC
ATTATCTTAGCCTAAAAAAACCTTCTCTTTGGAACTTTCAGTAATACGCTTAACTGCTCATTGCTA
TATTGAAGTACGGATTAGAAGCCGCCGAGCGGGTGACAGCCCTCCGAAGGAAGACTCTCCTCCG
TGCGTCCTCGTCTTCACCGGTCGCGTTCCTGAAACGCAGATGTGCCTCGCGCCGCACTGCTCCGA
ACAATAAAGATTCTACAATACTAGCTTTTATGGTTATGAAGAGGAAAAATTGGCAGTAACCTGGC
CCCACAAACCTTCAAATGAACGAATCAAATTAACAACCATAGGATGATAATGCGATTAGTTTTTT
AGCCTTATTTCTGGGGTAATTAATCAGCGAAGCGATGATTTTTGATCTATTAACAGATATATAAAT
GCAAAAACTGCATAACCACTTTAACTAATACTTTCAACATTTTCGGTTTGTATTACTTCTTATTCA
AATGTAATAAAAGTATCAACAAAAAAATTGTTAATATACCTCTATACTTTAACGTCAAGGAGAAAA
AACCCCGGATCCGTAATACGACTCACTATAGGGCCCGGGCGTCGACATGGAACAGAAGTTGATTT
CCGAAGAAGACCTCGAGTAAGCTTGGTACCGCGGCTAGCTAAGATCCGCTCTAACCGAAAAGGA

AGGAGTTAGACAACCTGAAGTCTAGGTCCCTATTTATTTTTTTATAGTTATGTTAGTATTAAGAAC
GTTATTTATATTTCAAATTTTTCTTTTTTTTCTGTACAGACGCGTGTACGCATGTAACATTATACTG
AAAACCTTGCTTGAGAAGGTTTTGGGACGCTCGAAGATCCAGCTGCATTAATGAATCGGCCAACG
CGCGGGGAGAGGCGGTTTGCGTATTGGGCGCTCTTCCGCTTCCTCGCTCACTGACTCGCTGCGCTC
GGTCGTTCGGCTGCGGCGAGCGGTATCAGCTCACTCAAAGGCGGTAATACGGTTATCCACAGAAT
CAGGGGATAACGCAGGAAAGAACATGTGAGCAAAAGGCCAGCAAAAGGCCAGGAACCGTAAAA
AGGCCGCGTTGCTGGCGTTTTTCCATAGGCTCCGCCCCCCTGACGAGCATCACAAAAATCGACGC
TCAAGTCAGAGGTGGCGAAACCCGACAGGACTATAAAGATACCAGGCGTTTCCCCCTGGAAGCT
CCCTCGTGCGCTCTCCTGTTCCGACCCTGCCGCTTACCGGATACCTGTCCGCCTTTCTCCCTTCGG
GAAGCGTGGCGCTTTCTCATAGCTCACGCTGTAGGTATCTCAGTTCGGTGTAGGTCGTTCGCTCCA
AGCTGGGCTGTGTGCACGAACCCCCCGTTCAGCCCGACCGCTGCGCCTTATCCGGTAACTATCGT
CTTGAGTCCAACCCGGTAAGACACGACTTATCGCCACTGGCAGCAGCCACTGGTAACAGGATTAG
CAGAGCGAGGTATGTAGGCGGTGCTACAGAGTTCTTGAAGTGGTGGCCTAACTACGGCTACACTA
GAAGGACAGTATTTGGTATCTGCGCTCTGCTGAAGCCAGTTACCTTCGGAAAAAGAGTTGGTAGC
TCTTGATCCGGCAAACAAACCACCGCTGGTAGCGGTGGTTTTTTTGTTTGCAAGCAGCAGATTAC
GCGCAGAAAAAAAGGATCTCAAGAAGATCCTTTGATCTTTTCTACGGGGTCTGACGCTCAGTGGA
ACGAAAACTCACGTTAAGGGATTTTGGTCATGAGATTATCAAAAAGGATCTTCACCTAGATCCTT
TTAAATTAAAAATGAAGTTTTAAATCAATCTAAAGTATATATGAGTAAACTTGGTCTGACAGTTA
CCAATGCTTAATCAGTGAGGCACCTATCTCAGCGATCTGTCTATTTCGTTCATCCATAGTTGCCTG
ACTCCCCGTCGTGTAGATAACTACGATACGGGAGGGCTTACCATCTGGCCCCAGTGCTGCAATGA
TACCGCGAGACCCACGCTCACCGGCTCCAGATTTATCAGCAATAAACCAGCCAGCCGGAAGGGC
CGAGCGCAGAAGTGGTCCTGCAACTTTATCCGCCTCCATCCAGTCTATTAATTGTTGCCGGGAAG
CTAGAGTAAGTAGTTCGCCAGTTAATAGTTTGCGCAACGTTGTTGCCATTGCTACAGGCATCGTG
GTGTCACGCTCGTCGTTTGGTATGGCTTCATTCAGCTCCGGTTCCCAACGATCAAGGCGAGTTACA
TGATCCCCCATGTTGTGCAAAAAAGCGGTTAGCTCCTTCGGTCCTCCGATCGTTGTCAGAAGTAA
GTTGGCCGCAGTGTTATCACTCATGGTTATGGCAGCACTGCATAATTCTCTTACTGTCATGCCATC
CGTAAGATGCTTTTCTGTGACTGGTGAGTACTCAACCAAGTCATTCTGAGAATAGTGTATGCGGC
GACCGAGTTGCTCTTGCCCGGCGTCAATACGGGATAATACCGCGCCACATAGCAGAACTTTAAAA
GTGCTCATCATTGGAAAACGTTCTTCGGGGCGAAAACTCTCAAGGATCTTACCGCTGTTGAGATC
CAGTTCGATGTAACCCACTCGTGCACCCAACTGATCTTCAGCATCTTTTACTTTCACCAGCGTTTC
TGGGTGAGCAAAAACAGGAAGGCAAAATGCCGCAAAAAAGGGAATAAGGGCGACACGGAAATG
TTGAATACTCATACTCTTCCTTTTTCAATATTATTGAAGCATTTATCAGGGTTATTGTCTCATGAGC
GGATACATATTTGAATGTATTTAGAAAAATAAACAAATAGGGGTTCCGCGCACATTTCCCCGAAA
AGTGCCACCTGAACGAAGCATCTGTGCTTCATTTTGTAGAACAAAAATGCAACGCGAGAGCGCTA
ATTTTTCAAACAAAGAATCTGAGCTGCATTTTTACAGAACAGAAATGCAACGCGAAAGCGCTATT
TTACCAACGAAGAATCTGTGCTTCATTTTTGTAAAACAAAAATGCAACGCGAGAGCGCTAATTTT
TCAAACAAAGAATCTGAGCTGCATTTTTACAGAACAGAAATGCAACGCGAGAGCGCTATTTTACC
AACAAAGAATCTATACTTCTTTTTTGTTCTACAAAAATGCATCCCGAGAGCGCTATTTTTCTAACA
AAGCATCTTAGATTACTTTTTTTCTCCTTTGTGCGCTCTATAATGCAGTCTCTTGATAACTTTTTGC
ACTGTAGGTCCGTTAAGGTTAGAAGAAGGCTACTTTGGTGTCTATTTTCTCTTCCATAAAAAAAG
CCTGACTCCACTTCCCGCGTTTACTGATTACTAGCGAAGCTGCGGGTGCATTTTTTCAAGATAAAG
GCATCCCCGATTATATTCTATACCGATGTGGATTGCGCATACTTTGTGAACAGAAAGTGATAGCG
TTGATGATTCTTCATTGGTCAGAAAATTATGAACGGTTTCTTCTATTTTGTCTCTATATACTACGTA
TAGGAAATGTTTACATTTTCGTATTGTTTTCGATTCACTCTATGAATAGTTCTTACTACAATTTTTT
TGTCTAAAGAGTAATACTAGAGATAAACATAAAAAAATGTAGAGGTCGAGTTTAGATGCAAGTTC
AAGGAGCGAAAGGTGGATGGGTAGGTTATATAGGGATATAGCACAGAGATATATAGCAAAGAG
ATACTTTTGAGCAATGTTTGTGGAAGCGGTATTCGCAATATTTTAGTAGCTCGTTACAGTCCGGTG
CGTTTTTGGTTTTTTGAAAGTGCGTCTTCAGAGCGCTTTTGGTTTTCAAAAGCGCTCTGAAGTTCC
TATACTTTCTAGAGAATAGGAACTTCGGAATAGGAACTTCAAAGCGTTTCCGAAAACGAGCGCTT
CCGAAAATGCAACGCGAGCTGCGCACATACAGCTCACTGTTCACGTCGCACCTATATCTGCGTGT
TGCCTGTATATATATATACATGAGAAGAACGGCATAGTGCGTGTTTATGCTTAAATGCGTACTTA
TATGCGTCTATTTATGTAGGATGAAAGGTAGTCTAGTACCTCCTGTGATATTATCCCATTCCATGC
GGGGTATCGTATGCTTCCTTCAGCACTACCCTTTAGCTGTTCTATATGCTGCCACTCCTCAATTGG
ATTAGTCTCATCCTTCAATGCTATCATTTCCTTTGATATTGGATCATCTAAGAAACCATTATTATC
ATGACATTAACCTATAAAAATAGGCGTATCACGAGGCCCTTTCGTC

> pESC-HIS-*GS*$^{I301E}$*~mVenus*$^{C}$

TCGCGCGTTTCGGTGATGACGGTGAAAACCTCTGACACATGCAGCTCCCGGAGACGGTCACAGCTTGT
CTGTAAGCGGATGCCGGGAGCAGACAAGCCCGTCAGGGCGCGTCAGCGGGTGTTGGCGGGTGTCGGG
GCTGGCTTAACTATGCGGCATCAGAGCAGATTGTACTGAGAGTGCACCATAAATTCCCGTTTTAAGAG
CTTGGTGAGCGCTAGGAGTCACTGCCAGGTATCGTTTGAACACGGCATTAGTCAGGGAAGTCATAACA
CAGTCCTTTCCCGCAATTTTCTTTTTCTATTACTCTTGGCCTCCTCTAGTACACTCTATATTTTTTTATGC
CTCGGTAATGATTTTCATTTTTTTTTTTCCCCTAGCGGATGACTCTTTTTTTTTCTTAGCGATTGGCATTA
TCACATAATGAATTATACATTATATAAAGTAATGTGATTTCTTCGAAGAATATACTAAAAAATGAGCA
GGCAAGATAAACGAAGGCAAAGATGACAGAGCAGAAAGCCCTAGTAAAGCGTATTACAAATGAAAC
CAAGATTCAGATTGCGATCTCTTTAAAGGGTGGTCCCCTAGCGATAGAGCACTCGATCTTCCCAGAAA
AAGAGGCAGAAGCAGTAGCAGAACAGGCCACACAATCGCAAGTGATTAACGTCCACACAGGTATAGG
GTTTCTGGACCATATGATACATGCTCTGGCCAAGCATTCCGGCTGGTCGCTAATCGTTGAGTGCATTGG
TGACTTACACATAGACGACCATCACACCACTGAAGACTGCGGGATTGCTCTCGGTCAAGCTTTTAAAG
AGGCCCTACTGGCGCGTGGAGTAAAAAGGTTTGGATCAGGATTTGCGCCTTTGGATGAGGCACTTTCC
AGAGCGGTGGTAGATCTTTCGAACAGGCCGTACGCAGTTGTCGAACTTGGTTTGCAAAGGGAGAAAGT
AGGAGATCTCTCTTGCGAGATGATCCCGCATTTTCTTGAAAGCTTTGCAGAGGCTAGCAGAATTACCCT
CCACGTTGATTGTCTGCGAGGCAAGAATGATCATCACCGTAGTGAGAGTGCGTTCAAGGCTCTTGCGG
TTGCCATAAGAGAAGCCACCTCGCCCAATGGTACCAACGATGTTCCCTCCACCAAAGGTGTTCTTATGT
AGTGACACCGATTATTTAAAGCTGCAGCATACGATATATATACATGTGTATATATGTATACCTATGAAT
GTCAGTAAGTATGTATACGAACAGTATGATACTGAAGATGACAAGGTAATGCATCATTCTATACGTGT
CATTCTGAACGAGGCGCGCTTTCCTTTTTTCTTTTTGCTTTTTCTTTTTTTTTCTCTTGAACTCGACGGAT
CTATGCGGTGTGAAATACCGCACAGATGCGTAAGGAGAAAATACCGCATCAGGAAATTGTAAACGTT
AATATTTTGTTAAAATTCGCGTTAAATTTTTGTTAAATCAGCTCATTTTTTAACCAATAGGCCGAAATC
GGCAAAATCCCTTATAAATCAAAAGAATAGACCGAGATAGGGTTGAGTGTTGTTCCAGTTTGGAACAA
GAGTCCACTATTAAAGAACGTGGACTCCAACGTCAAAGGGCGAAAAACCGTCTATCAGGGCGATGGC
CCACTACGTGAACCATCACCCTAATCAAGTTTTTTGGGGTCGAGGTGCCGTAAAGCACTAAATCGGAA
CCCTAAAGGGAGCCCCCGATTTAGAGCTTGACGGGGAAAGCCGGCGAACGTGGCGAGAAAGGAAGGG
AAGAAAGCGAAAGGAGCGGGCGCTAGGGCGCTGGCAAGTGTAGCGGTCACGCTGCGCGTAACCACCA
CACCCGCCGCGCTTAATGCGCCGCTACAGGGCGCGTCGCGCCATTCGCCATTCAGGCTGCGCAACTGT
TGGGAAGGGCGATCGGTGCGGGCCTCTTCGCTATTACGCCAGCTGAATTGGAGCGACCTCATGCTATA
CCTGAGAAAGCAACCTGACCTACAGGAAAGAGTTACTCAAGAATAAGAATTTTCGTTTTAAAACCTAA
GAGTCACTTTAAAATTTGTATACACTTATTTTTTTTATAACTTATTTAATAATAAAAATCATAAATCATA
AGAAATTCGCTTATTTAGAAGTGTCAACAACGTATCTACCAACGATTTGACCCTTTTCCATCTTTTCGT
AAATTTCTGGCAAGGTAGACAAGCCGACAACCTTGATTGGAGACTTGACCAAACCTCTGGCGAAGAAT
TGTTAATTAAGAGCTCAGATCTTATCGTCGTCATCCTTGTAATCCATCGATACTAGTGCTTACTTGTAC
AGCTCGTCCATGCCGAGAGTGATCCCGGCGGCGGTCACGAACTCCAGCAGGACCATGTGATCGCGCTT
CTCGTTGGGGTCTTTGCTCAGCTTGGACTGGTAGCTCAGGTAGTGGTTGTCGGGCAGCAGCACGGGGC
CGTCGCCGATGGGGGTGTTCTGCTGGTAGTGGTCGGCGAGCTGCACGCCGCCGTCCTCGATGTTGTGG
CGGATCTTGAAGTTGGCCTTGATGCCGTTCTTCTGCTTGTCCATAGTACCACCAGAACCTTCCTCAAAT
TTCAATGTATTTCCAATGTCAATCGCAAAACGGTACTTGACATCCAAATTCTTGATACGTTCCATAGCA
GTGCTGAGATAGTCAATCCCAATAACTTCAGTATCACATACAATGTTATGTTTGGCTGCGAAATCAAG
CATTTCTTGGTACTCTTTGAGACCTCCAGTGGAACTTCCGATTTCCTTTTTCCTTCCCATAATGAGAGGT
GCCGCAGGGAGCTCAAATAGCGACTCCGGTGCACCTACGAGCATAACCGCGCCGTCAAACTTGAGCA
AATTGAGCATAAGTGACATAGGAGTGCGGCCACCTGGGGTGGTGTCCACAACACCATCCATAGTACCT
GCCAGAGCCTTCAATTGCTCAGAGTCAGTGTTGACAACAAAAGCATCAGCACCATGTTCTTCAATGGC
TTCCTTCTCCTTACGCCTTGATGTACTAATAACAGTAGCCTTACCACCAAAAGCCTTAATAAACTTAAC
AGCAACAGAACCAAGACCTCCCAGCCCGAAAACCCCAATATGCTTTCCTGGCTTATCGAGTCCCAAAT
GTTTCATTGGGCTATAAACAACAACCCCAGCACAAAGGAGAGCAACCCCTTTATCTTGAGGCAAGTTT
TCGGGCCATCGGAGGACGAACTTTTCATCAACAACCATCACATTTGAACAACCCCCATAGGATCGTTC
CCCTTGCTCACGGTAAACAGATCCATCAGCCATATTGGGCTCTGGGCAGTAATTCTCCATTCCACTTTG
ACAATTATAACATTGACCACAAGATCCGACCATACATCCCACAGCTACCTTGTCTCCAACCTTGAATTT
CTCTACTTTGCTGCCAACTTCTACCACCTCACCGGCAGTTTCATGTCCAAACACATAAGGATATCTGGT
GAAACCCCACTTGTTTCTGACCATTTCCATATCGAAATTGCAAACACCAGAGTACAAAACTCTAATCTT
CACATCCCGTTCACCAGGGACTCTTCTATAGAACTTGATGGGCTGAAGGACACCAGATGCATCTGCAG
CACCCCATCCCACAGCCTTCACTGAAAGGTCGAGTTTGGTTGTTTCTCCGGCCATGCCCTTTAGTGAGG

GTTGAATTCGAATTTTCAAAAATTCTTACTTTTTTTTTGGATGGACGCAAAGAAGTTTAATAATCATAT
TACATGGCATTACCACCATATACATATCCATATACATATCCATATCTAATCTTACTTATATGTTGTGGA
AATGTAAAGAGCCCCATTATCTTAGCCTAAAAAAACCTTCTCTTTGGAACTTTCAGTAATACGCTTAAC
TGCTCATTGCTATATTGAAGTACGGATTAGAAGCCGCCGAGCGGGTGACAGCCCTCCGAAGGAAGACT
CTCCTCCGTGCGTCCTCGTCTTCACCGGTCGCGTTCCTGAAACGCAGATGTGCCTCGCGCCGCACTGCT
CCGAACAATAAAGATTCTACAATACTAGCTTTTATGGTTATGAAGAGGAAAAATTGGCAGTAACCTGG
CCCCACAAACCTTCAAATGAACGAATCAAATTAACAACCATAGGATGATAATGCGATTAGTTTTTTAG
CCTTATTTCTGGGGTAATTAATCAGCGAAGCGATGATTTTTGATCTATTAACAGATATATAAATGCAAA
AACTGCATAACCACTTTAACTAATACTTTCAACATTTTCGGTTTGTATTACTTCTTATTCAAATGTAATA
AAAGTATCAACAAAAAATTGTTAATATACCTCTATACTTTAACGTCAAGGAGAAAAAACCCCGGATCC
GTAATACGACTCACTATAGGGCCCGGGCGTCGACATGGAACAGAAGTTGATTTCCGAAGAAGACCTCG
AGTAAGCTTGGTACCGCGGCTAGCTAAGATCCGCTCTAACCGAAAAGGAAGGAGTTAGACAACCTGA
AGTCTAGGTCCCTATTTATTTTTTTATAGTTATGTTAGTATTAAGAACGTTATTTATATTTCAAATTTTTC
TTTTTTTTCTGTACAGACGCGTGTACGCATGTAACATTATACTGAAAACCTTGCTTGAGAAGGTTTTGG
GACGCTCGAAGATCCAGCTGCATTAATGAATCGGCCAACGCGCGGGGAGAGGCGGTTTGCGTATTGG
GCGCTCTTCCGCTTCCTCGCTCACTGACTCGCTGCGCTCGGTCGTTCGGCTGCGGCGAGCGGTATCAGC
TCACTCAAAGGCGGTAATACGGTTATCCACAGAATCAGGGGATAACGCAGGAAAGAACATGTGAGCA
AAAGGCCAGCAAAAGGCCAGGAACCGTAAAAAGGCCGCGTTGCTGGCGTTTTTCCATAGGCTCCGCCC
CCCTGACGAGCATCACAAAAATCGACGCTCAAGTCAGAGGTGGCGAAACCCGACAGGACTATAAAGA
TACCAGGCGTTTCCCCCTGGAAGCTCCCTCGTGCGCTCTCCTGTTCCGACCCTGCCGCTTACCGGATAC
CTGTCCGCCTTTCTCCCTTCGGGAAGCGTGGCGCTTTCTCATAGCTCACGCTGTAGGTATCTCAGTTCG
GTGTAGGTCGTTCGCTCCAAGCTGGGCTGTGTGCACGAACCCCCCGTTCAGCCCGACCGCTGCGCCTTA
TCCGGTAACTATCGTCTTGAGTCCAACCCGGTAAGACACGACTTATCGCCACTGGCAGCAGCCACTGG
TAACAGGATTAGCAGAGCGAGGTATGTAGGCGGTGCTACAGAGTTCTTGAAGTGGTGGCCTAACTACG
GCTACACTAGAAGGACAGTATTTGGTATCTGCGCTCTGCTGAAGCCAGTTACCTTCGGAAAAAGAGTT
GGTAGCTCTTGATCCGGCAAACAAACCACCGCTGGTAGCGGTGGTTTTTTTGTTTGCAAGCAGCAGATT
ACGCGCAGAAAAAAAGGATCTCAAGAAGATCCTTTGATCTTTTCTACGGGGTCTGACGCTCAGTGGAA
CGAAAACTCACGTTAAGGGATTTTGGTCATGAGATTATCAAAAAGGATCTTCACCTAGATCCTTTTAA
ATTAAAAATGAAGTTTTAAATCAATCTAAAGTATATATGAGTAAACTTGGTCTGACAGTTACCAATGC
TTAATCAGTGAGGCACCTATCTCAGCGATCTGTCTATTTCGTTCATCCATAGTTGCCTGACTCCCCGTC
GTGTAGATAACTACGATACGGGAGGGCTTACCATCTGGCCCCAGTGCTGCAATGATACCGCGAGACCC
ACGCTCACCGGCTCCAGATTTATCAGCAATAAACCAGCCAGCCGGAAGGGCCGAGCGCAGAAGTGGT
CCTGCAACTTTATCCGCCTCCATCCAGTCTATTAATTGTTGCCGGGAAGCTAGAGTAAGTAGTTCGCCA
GTTAATAGTTTGCGCAACGTTGTTGCCATTGCTACAGGCATCGTGGTGTCACGCTCGTCGTTTGGTATG
GCTTCATTCAGCTCCGGTTCCCAACGATCAAGGCGAGTTACATGATCCCCCATGTTGTGCAAAAAAGC
GGTTAGCTCCTTCGGTCCTCCGATCGTTGTCAGAAGTAAGTTGGCCGCAGTGTTATCACTCATGGTTAT
GGCAGCACTGCATAATTCTCTTACTGTCATGCCATCCGTAAGATGCTTTTCTGTGACTGGTGAGTACTC
AACCAAGTCATTCTGAGAATAGTGTATGCGGCGACCGAGTTGCTCTTGCCCGGCGTCAATACGGGATA
ATACCGCGCCACATAGCAGAACTTTAAAAGTGCTCATCATTGGAAAACGTTCTTCGGGGCGAAAACTC
TCAAGGATCTTACCGCTGTTGAGATCCAGTTCGATGTAACCCACTCGTGCACCCAACTGATCTTCAGCA
TCTTTTACTTTCACCAGCGTTTCTGGGTGAGCAAAAACAGGAAGGCAAAATGCCGCAAAAAAGGGAAT
AAGGGCGACACGGAAATGTTGAATACTCATACTCTTCCTTTTTCAATATTATTGAAGCATTTATCAGGG
TTATTGTCTCATGAGCGGATACATATTTGAATGTATTTAGAAAAATAAACAAATAGGGGTTCCGCGCA
CATTTCCCCGAAAAGTGCCACCTGAACGAAGCATCTGTGCTTCATTTTGTAGAACAAAAATGCAACGC
GAGAGCGCTAATTTTTCAAACAAAGAATCTGAGCTGCATTTTTACAGAACAGAAATGCAACGCGAAAG
CGCTATTTTACCAACGAAGAATCTGTGCTTCATTTTTGTAAAACAAAAATGCAACGCGAGAGCGCTAA
TTTTTCAAACAAAGAATCTGAGCTGCATTTTTACAGAACAGAAATGCAACGCGAGAGCGCTATTTTAC
CAACAAAGAATCTATACTTCTTTTTTGTTCTACAAAAATGCATCCCGAGAGCGCTATTTTTCTAACAAA
GCATCTTAGATTACTTTTTTTCTCCTTTGTGCGCTCTATAATGCAGTCTCTTGATAACTTTTTGCACTGTA
GGTCCGTTAAGGTTAGAAGAAGGCTACTTTGGTGTCTATTTTCTCTTCCATAAAAAAAGCCTGACTCCA
CTTCCCGCGTTTACTGATTACTAGCGAAGCTGCGGGTGCATTTTTTCAAGATAAAGGCATCCCCGATTA
TATTCTATACCGATGTGGATTGCGCATACTTTGTGAACAGAAAGTGATAGCGTTGATGATTCTTCATTG
GTCAGAAAATTATGAACGGTTTCTTCTATTTTGTCTCTATATACTACGTATAGGAAATGTTTACATTTTC
GTATTGTTTTCGATTCACTCTATGAATAGTTCTTACTACAATTTTTTTGTCTAAAGAGTAATACTAGAGA
TAAACATAAAAAATGTAGAGGTCGAGTTTAGATGCAAGTTCAAGGAGCGAAAGGTGGATGGGTAGGT
TATATAGGGATATAGCACAGAGATATATAGCAAAGAGATACTTTTGAGCAATGTTTGTGGAAGCGGTA

TTCGCAATATTTTAGTAGCTCGTTACAGTCCGGTGCGTTTTTGGTTTTTTGAAAGTGCGTCTTCAGAGCG
CTTTTGGTTTTCAAAAGCGCTCTGAAGTTCCTATACTTTCTAGAGAATAGGAACTTCGGAATAGGAACT
TCAAAGCGTTTCCGAAAACGAGCGCTTCCGAAAATGCAACGCGAGCTGCGCACATACAGCTCACTGTT
CACGTCGCACCTATATCTGCGTGTTGCCTGTATATATATATACATGAGAAGAACGGCATAGTGCGTGTT
TATGCTTAAATGCGTACTTATATGCGTCTATTTATGTAGGATGAAAGGTAGTCTAGTACCTCCTGTGAT
ATTATCCCATTCCATGCGGGGTATCGTATGCTTCCTTCAGCACTACCCTTTAGCTGTTCTATATGCTGCC
ACTCCTCAATTGGATTAGTCTCATCCTTCAATGCTATCATTTCCTTTGATATTGGATCATCTAAGAAACC
ATTATTATCATGACATTAACCTATAAAAATAGGCGTATCACGAGGCCCTTTCGTC

> pESC-HIS-*mVenus*$^{C}$*~SGD*

TCGCGCGTTTCGGTGATGACGGTGAAAACCTCTGACACATGCAGCTCCCGGAGACGGTCACAGCTTGT
CTGTAAGCGGATGCCGGGAGCAGACAAGCCCGTCAGGGCGCGTCAGCGGGTGTTGGCGGGTGTCGGG
GCTGGCTTAACTATGCGGCATCAGAGCAGATTGTACTGAGAGTGCACCATAAATTCCCGTTTTAAGAG
CTTGGTGAGCGCTAGGAGTCACTGCCAGGTATCGTTTGAACACGGCATTAGTCAGGGAAGTCATAACA
CAGTCCTTTCCCGCAATTTTCTTTTTCTATTACTCTTGGCCTCCTCTAGTACACTCTATATTTTTTTATGC
CTCGGTAATGATTTTCATTTTTTTTTTTCCCCTAGCGGATGACTCTTTTTTTTTCTTAGCGATTGGCATTA
TCACATAATGAATTATACATTATATAAAGTAATGTGATTTCTTCGAAGAATATACTAAAAAATGAGCA
GGCAAGATAAACGAAGGCAAAGATGACAGAGCAGAAAGCCCTAGTAAAGCGTATTACAAATGAAAC
CAAGATTCAGATTGCGATCTCTTTAAAGGGTGGTCCCCTAGCGATAGAGCACTCGATCTTCCCAGAAA
AAGAGGCAGAAGCAGTAGCAGAACAGGCCACACAATCGCAAGTGATTAACGTCCACACAGGTATAGG
GTTTCTGGACCATATGATACATGCTCTGGCCAAGCATTCCGGCTGGTCGCTAATCGTTGAGTGCATTGG
TGACTTACACATAGACGACCATCACACCACTGAAGACTGCGGGATTGCTCTCGGTCAAGCTTTTAAAG
AGGCCCTACTGGCGCGTGGAGTAAAAAGGTTTGGATCAGGATTTGCGCCTTTGGATGAGGCACTTTCC
AGAGCGGTGGTAGATCTTTCGAACAGGCCGTACGCAGTTGTCGAACTTGGTTTGCAAAGGGAGAAAGT
AGGAGATCTCTCTTGCGAGATGATCCCGCATTTTCTTGAAAGCTTTGCAGAGGCTAGCAGAATTACCCT
CCACGTTGATTGTCTGCGAGGCAAGAATGATCATCACCGTAGTGAGAGTGCGTTCAAGGCTCTTGCGG
TTGCCATAAGAGAAGCCACCTCGCCCAATGGTACCAACGATGTTCCCTCCACCAAAGGTGTTCTTATGT
AGTGACACCGATTATTTAAAGCTGCAGCATACGATATATATACATGTGTATATATGTATACCTATGAAT
GTCAGTAAGTATGTATACGAACAGTATGATACTGAAGATGACAAGGTAATGCATCATTCTATACGTGT
CATTCTGAACGAGGCGCGCTTTCCTTTTTTCTTTTTGCTTTTTCTTTTTTTTTCTCTTGAACTCGACGGAT
CTATGCGGTGTGAAATACCGCACAGATGCGTAAGGAGAAAATACCGCATCAGGAAATTGTAAACGTT
AATATTTTGTTAAAATTCGCGTTAAATTTTTGTTAAATCAGCTCATTTTTTAACCAATAGGCCGAAATC
GGCAAAATCCCTTATAAATCAAAAGAATAGACCGAGATAGGGTTGAGTGTTGTTCCAGTTTGGAACAA
GAGTCCACTATTAAAGAACGTGGACTCCAACGTCAAAGGGCGAAAAACCGTCTATCAGGGCGATGGC
CCACTACGTGAACCATCACCCTAATCAAGTTTTTTGGGGTCGAGGTGCCGTAAAGCACTAAATCGGAA
CCCTAAAGGGAGCCCCCGATTTAGAGCTTGACGGGGAAAGCCGGCGAACGTGGCGAGAAAGGAAGGG
AAGAAAGCGAAAGGAGCGGGCGCTAGGGCGCTGGCAAGTGTAGCGGTCACGCTGCGCGTAACCACCA
CACCCGCCGCGCTTAATGCGCCGCTACAGGGCGCGTCGCGCCATTCGCCATTCAGGCTGCGCAACTGT
TGGGAAGGGCGATCGGTGCGGGCCTCTTCGCTATTACGCCAGCTGAATTGGAGCGACCTCATGCTATA
CCTGAGAAAGCAACCTGACCTACAGGAAAGAGTTACTCAAGAATAAGAATTTTCGTTTTAAAACCTAA
GAGTCACTTTAAAATTTGTATACACTTATTTTTTTTATAACTTATTTAATAATAAAAATCATAAATCATA
AGAAATTCGCTTATTTAGAAGTGTCAACAACGTATCTACCAACGATTTGACCCTTTTCCATCTTTTCGT
AAATTTCTGGCAAGGTAGACAAGCCGACAACCTTGATTGGAGACTTGACCAAACCTCTGGCGAAGAAT
TGTTAATTAAGAGCTCAGATCTTATCGTCGTCATCCTTGTAATCCATCGATACTAGTGCTTAATATTTTT
GTTTTTTAACCAATTCAACCAATTTATCTTCTTCTCTAAATCTTTTTTTAGCAGTATTAGTAACAAAACC
TTCAGAAATAAAATTTTTATACCAAATAGCAGAATCTTTTGGATATCTTTGAAAAGTTTTATAATCAAC
ATGAATAATACCATATCTACAAATATAACCCAAATTCCATTCAAAATTATCAAAAAAAGACCAAACAA
AAAAACCTTTAACATTAACACCATCATCAATAGCATCTCTAACAGAAGCCAAATGAGATTGCAAAAAA
TCAACTCTCAATTTATCATGTCTAGCTTCAGTCAACAAAATATTAGTTTTACCTTCAGTCAACAAAATA
TTAGTTCTATTTTCTTCAACAACACCACATTCAGAAACATAAATAACTGGAACATGATATTTTTCTTTA
GTATAAACCAACAAATTATACAAACCAGATGGAACAACATGTTGCCAACCACCATAACATGGTTCACC
AATTCTAACTTCTTTACCATCAACTTTTTTAACAAAAATATTTTTATTAATTCTAGCATCAGTTTCATAA
CCTGGAGTATCTGGAATTTTATCAGCATTAGAAACATAAGTAGTAGTATAATAATTCATACCAATAAA
ATCATAACAACCAGTCAATTTTTCAGAATCTTCAGTAGAAAATTCTGGCAATCTAGAACCAACCAAAG
CTCTCATAGATTTTGGATATTCACCAGTAGTCAATGGTTCAATAAACCAACCCAACATAAAATCTGGA

CCTCTTTCTCTAGCATCAATATCTTCTTTAGTTTCATTCAATGGTTCCATCCACATAGAATTCAAAACAA
TACCAATTTCACCACCTTGACATTTTTGAAAATTTTTTCTATAAACTTCAACAGCAGCTTTATGAGACA
ACAACAAATTGTGTGTAGCAATATATGGTTCTTTACCTGGATTACCCTTACCATCAGCACCACCTCTAC
CTGGAGCAAATTCACCTGTAGCATAACCAGAAGCAACATAAGTATGTGGTTCGTTAAAAGTAGTCCAG
AATTTAACCTTATCACCAAATTCCCAAAAACAAAATTCAGCGTATTCAGTGAAATCTTCAACAATTCTA
TCAGACAAAAAACCACCATATTCATCTTCTAGAGCTTGAGGCAAATCCCAATGAAACAAAGTAGCAAA
AGGCTTGATACCATTAGCCAACAATTCATCTATAAAATCGTGGTAAAACTTAACACCATCTTTATTAAC
ACCACCAGACAAATTACCACCTGGCAAAACTCTAGACCATGAGATAGAAAATCTATAAGATTCCAAAC
CTGTTTGCTTCATGATTTTGATATCTTCCTTGTACAAGTTGTAAGAATTAATAGCTTGATTACCATTTGA
ACCATCAGCAATCTTAGCTGGGTATCTATTAGTAAAGGTATCCCATATTGATGGACCTCTGTTACCTTC
GTTATAAGCACCTTCACATTGATAAGCAGAACCACCGGCACCCAAAATAAAATCAGATGGAAAATCTC
TTCTGTGAACAATTGGCTTGTTATGTTTTCTTGGTTGAATTGGAATAGATGGATAAGCAAATGGAATTG
GAACGGAGTGATTACCATTTGGTTCAGCTGCTGGGGATATAGCAACAACCAAAGATTGATCATCTTTG
GAACCCATAGTACCACCAGAACCCTTGTACAGCTCGTCCATGCCGAGAGTGATCCCGGCGGCGGTCAC
GAACTCCAGCAGGACCATGTGATCGCGCTTCTCGTTGGGGTCTTTGCTCAGCTTGGACTGGTAGCTCAG
GTAGTGGTTGTCGGGCAGCAGCACGGGGCCGTCGCCGATGGGGGTGTTCTGCTGGTAGTGGTCGGCGA
GCTGCACGCCGCCGTCCTCGATGTTGTGGCGGATCTTGAAGTTGGCCTTGATGCCGTTCTTCTGCTTGT
CCATGCCCTTTAGTGAGGGTTGAATTCGAATTTTCAAAAATTCTTACTTTTTTTTTGGATGGACGCAAA
GAAGTTTAATAATCATATTACATGGCATTACCACCATATACATATCCATATACATATCCATATCTAATC
TTACTTATATGTTGTGGAAATGTAAAGAGCCCCATTATCTTAGCCTAAAAAAACCTTCTCTTTGGAACT
TTCAGTAATACGCTTAACTGCTCATTGCTATATTGAAGTACGGATTAGAAGCCGCCGAGCGGGTGACA
GCCCTCCGAAGGAAGACTCTCCTCCGTGCGTCCTCGTCTTCACCGGTCGCGTTCCTGAAACGCAGATGT
GCCTCGCGCCGCACTGCTCCGAACAATAAAGATTCTACAATACTAGCTTTTATGGTTATGAAGAGGAA
AAATTGGCAGTAACCTGGCCCCACAAACCTTCAAATGAACGAATCAAATTAACAACCATAGGATGATA
ATGCGATTAGTTTTTTAGCCTTATTTCTGGGGTAATTAATCAGCGAAGCGATGATTTTTGATCTATTAA
CAGATATATAAATGCAAAAACTGCATAACCACTTTAACTAATACTTTCAACATTTTCGGTTTGTATTAC
TTCTTATTCAAATGTAATAAAAGTATCAACAAAAAATTGTTAATATACCTCTATACTTTAACGTCAAGG
AGAAAAAACCCCGGATCCGTAATACGACTCACTATAGGGCCCGGGCGTCGACATGGAACAGAAGTTG
ATTTCCGAAGAAGACCTCGAGTAAGCTTGGTACCGCGGCTAGCTAAGATCCGCTCTAACCGAAAAGGA
AGGAGTTAGACAACCTGAAGTCTAGGTCCCTATTTATTTTTTTATAGTTATGTTAGTATTAAGAACGTT
ATTTATATTTCAAATTTTTCTTTTTTTTCTGTACAGACGCGTGTACGCATGTAACATTATACTGAAAACC
TTGCTTGAGAAGGTTTTGGGACGCTCGAAGATCCAGCTGCATTAATGAATCGGCCAACGCGCGGGGAG
AGGCGGTTTGCGTATTGGGCGCTCTTCCGCTTCCTCGCTCACTGACTCGCTGCGCTCGGTCGTTCGGCT
GCGGCGAGCGGTATCAGCTCACTCAAAGGCGGTAATACGGTTATCCACAGAATCAGGGGATAACGCA
GGAAAGAACATGTGAGCAAAAGGCCAGCAAAAGGCCAGGAACCGTAAAAAGGCCGCGTTGCTGGCG
TTTTTCCATAGGCTCCGCCCCCCTGACGAGCATCACAAAAATCGACGCTCAAGTCAGAGGTGGCGAAA
CCCGACAGGACTATAAAGATACCAGGCGTTTCCCCCTGGAAGCTCCCTCGTGCGCTCTCCTGTTCCGAC
CCTGCCGCTTACCGGATACCTGTCCGCCTTTCTCCCTTCGGGAAGCGTGGCGCTTTCTCATAGCTCACG
CTGTAGGTATCTCAGTTCGGTGTAGGTCGTTCGCTCCAAGCTGGGCTGTGTGCACGAACCCCCCGTTCA
GCCCGACCGCTGCGCCTTATCCGGTAACTATCGTCTTGAGTCCAACCCGGTAAGACACGACTTATCGCC
ACTGGCAGCAGCCACTGGTAACAGGATTAGCAGAGCGAGGTATGTAGGCGGTGCTACAGAGTTCTTG
AAGTGGTGGCCTAACTACGGCTACACTAGAAGGACAGTATTTGGTATCTGCGCTCTGCTGAAGCCAGT
TACCTTCGGAAAAAGAGTTGGTAGCTCTTGATCCGGCAAACAAACCACCGCTGGTAGCGGTGGTTTTT
TTGTTTGCAAGCAGCAGATTACGCGCAGAAAAAAAGGATCTCAAGAAGATCCTTTGATCTTTTCTACG
GGGTCTGACGCTCAGTGGAACGAAAACTCACGTTAAGGGATTTTGGTCATGAGATTATCAAAAAGGAT
CTTCACCTAGATCCTTTTAAATTAAAAATGAAGTTTTAAATCAATCTAAAGTATATATGAGTAAACTTG
GTCTGACAGTTACCAATGCTTAATCAGTGAGGCACCTATCTCAGCGATCTGTCTATTTCGTTCATCCAT
AGTTGCCTGACTCCCCGTCGTGTAGATAACTACGATACGGGAGGGCTTACCATCTGGCCCCAGTGCTG
CAATGATACCGCGAGACCCACGCTCACCGGCTCCAGATTTATCAGCAATAAACCAGCCAGCCGGAAG
GGCCGAGCGCAGAAGTGGTCCTGCAACTTTATCCGCCTCCATCCAGTCTATTAATTGTTGCCGGGAAG
CTAGAGTAAGTAGTTCGCCAGTTAATAGTTTGCGCAACGTTGTTGCCATTGCTACAGGCATCGTGGTGT
CACGCTCGTCGTTTGGTATGGCTTCATTCAGCTCCGGTTCCCAACGATCAAGGCGAGTTACATGATCCC
CCATGTTGTGCAAAAAAGCGGTTAGCTCCTTCGGTCCTCCGATCGTTGTCAGAAGTAAGTTGGCCGCA
GTGTTATCACTCATGGTTATGGCAGCACTGCATAATTCTCTTACTGTCATGCCATCCGTAAGATGCTTTT
CTGTGACTGGTGAGTACTCAACCAAGTCATTCTGAGAATAGTGTATGCGGCGACCGAGTTGCTCTTGC
CCGGCGTCAATACGGGATAATACCGCGCCACATAGCAGAACTTTAAAAGTGCTCATCATTGGAAAACG

TTCTTCGGGGCGAAAACTCTCAAGGATCTTACCGCTGTTGAGATCCAGTTCGATGTAACCCACTCGTGC
ACCCAACTGATCTTCAGCATCTTTTACTTTCACCAGCGTTTCTGGGTGAGCAAAAACAGGAAGGCAAA
ATGCCGCAAAAAAGGGAATAAGGGCGACACGGAAATGTTGAATACTCATACTCTTCCTTTTTCAATAT
TATTGAAGCATTTATCAGGGTTATTGTCTCATGAGCGGATACATATTTGAATGTATTTAGAAAAATAAA
CAAATAGGGGTTCCGCGCACATTTCCCCGAAAAGTGCCACCTGAACGAAGCATCTGTGCTTCATTTTGT
AGAACAAAAATGCAACGCGAGAGCGCTAATTTTTCAAACAAAGAATCTGAGCTGCATTTTTACAGAAC
AGAAATGCAACGCGAAAGCGCTATTTTACCAACGAAGAATCTGTGCTTCATTTTTGTAAAACAAAAAT
GCAACGCGAGAGCGCTAATTTTTCAAACAAAGAATCTGAGCTGCATTTTTACAGAACAGAAATGCAAC
GCGAGAGCGCTATTTTACCAACAAAGAATCTATACTTCTTTTTTGTTCTACAAAAATGCATCCCGAGAG
CGCTATTTTTCTAACAAAGCATCTTAGATTACTTTTTTTCTCCTTTGTGCGCTCTATAATGCAGTCTCTT
GATAACTTTTTGCACTGTAGGTCCGTTAAGGTTAGAAGAAGGCTACTTTGGTGTCTATTTTCTCTTCCAT
AAAAAAAGCCTGACTCCACTTCCCGCGTTTACTGATTACTAGCGAAGCTGCGGGTGCATTTTTTCAAGA
TAAAGGCATCCCCGATTATATTCTATACCGATGTGGATTGCGCATACTTTGTGAACAGAAAGTGATAG
CGTTGATGATTCTTCATTGGTCAGAAAATTATGAACGGTTTCTTCTATTTTGTCTCTATATACTACGTAT
AGGAAATGTTTACATTTTCGTATTGTTTTCGATTCACTCTATGAATAGTTCTTACTACAATTTTTTTGTC
TAAAGAGTAATACTAGAGATAAACATAAAAAATGTAGAGGTCGAGTTTAGATGCAAGTTCAAGGAGC
GAAAGGTGGATGGGTAGGTTATATAGGGATATAGCACAGAGATATATAGCAAAGAGATACTTTTGAG
CAATGTTTGTGGAAGCGGTATTCGCAATATTTTAGTAGCTCGTTACAGTCCGGTGCGTTTTTGGTTTTTT
GAAAGTGCGTCTTCAGAGCGCTTTTGGTTTTCAAAAGCGCTCTGAAGTTCCTATACTTTCTAGAGAATA
GGAACTTCGGAATAGGAACTTCAAAGCGTTTCCGAAAACGAGCGCTTCCGAAAATGCAACGCGAGCT
GCGCACATACAGCTCACTGTTCACGTCGCACCTATATCTGCGTGTTGCCTGTATATATATATACATGAG
AAGAACGGCATAGTGCGTGTTTATGCTTAAATGCGTACTTATATGCGTCTATTTATGTAGGATGAAAG
GTAGTCTAGTACCTCCTGTGATATTATCCCATTCCATGCGGGGTATCGTATGCTTCCTTCAGCACTACC
CTTTAGCTGTTCTATATGCTGCCACTCCTCAATTGGATTAGTCTCATCCTTCAATGCTATCATTTCCTTT
GATATTGGATCATCTAAGAAACCATTATTATCATGACATTAACCTATAAAAATAGGCGTATCACGAGG
CCCTTTCGTC

> pESC-LEU-*CrVinBLAST~mVenus*$^{N}$

TCGCGCGTTTCGGTGATGACGGTGAAAACCTCTGACACATGCAGCTCCCGGAGACGGTCACAGCT
TGTCTGTAAGCGGATGCCGGGAGCAGACAAGCCCGTCAGGGCGCGTCAGCGGGTGTTGGCGGGT
GTCGGGGCTGGCTTAACTATGCGGCATCAGAGCAGATTGTACTGAGAGTGCACCATATCGACTAC
GTCGTAAGGCCGTTTCTGACAGAGTAAAATTCTTGAGGGAACTTTCACCATTATGGGAAATGCTT
CAAGAAGGTATTGACTTAAACTCCATCAAATGGTCAGGTCATTGAGTGTTTTTTATTTGTTGTATT
TTTTTTTTTTTAGAGAAAATCCTCCAATATCAAATTAGGAATCGTAGTTTCATGATTTTCTGTTACA
CCTAACTTTTTGTGTGGTGCCCTCCTCCTTGTCAATATTAATGTTAAAGTGCAATTCTTTTTCCTTA
TCACGTTGAGCCATTAGTATCAATTTGCTTACCTGTATTCCTTTACTATCCTCCTTTTTCTCCTTCTT
GATAAATGTATGTAGATTGCGTATATAGTTTCGTCTACCCTATGAACATATTCCATTTTGTAATTT
CGTGTCGTTTCTATTATGAATTTCATTTATAAAGTTTATGTACAAATATCATAAAAAAAGAGAATC
TTTTTAAGCAAGGATTTTCTTAACTTCTTCGGCGACAGCATCACCGACTTCGGTGGTACTGTTGGA
ACCACCTAAATCACCAGTTCTGATACCTGCATCCAAAACCTTTTTAACTGCATCTTCAATGGCCTT
ACCTTCTTCAGGCAAGTTCAATGACAATTTCAACATCATTGCAGCAGACAAGATAGTGGCGATAG
GGTCAACCTTATTCTTTGGCAAATCTGGAGCAGAACCGTGGCATGGTTCGTACAAACCAAATGCG
GTGTTCTTGTCTGGCAAAGAGGCCAAGGACGCAGATGGCAACAAACCCAAGGAACCTGGGATAA
CGGAGGCTTCATCGGAGATGATATCACCAAACATGTTGCTGGTGATTATAATACCATTTAGGTGG
GTTGGGTTCTTAACTAGGATCATGGCGGCAGAATCAATCAATTGATGTTGAACCTTCAATGTAGG
GAATTCGTTCTTGATGGTTTCCTCCACAGTTTTTCTCCATAATCTTGAAGAGGCCAAAAGATTAGC
TTTATCCAAGGACCAAATAGGCAATGGTGGCTCATGTTGTAGGGCCATGAAAGCGGCCATTCTTG
TGATTCTTTGCACTTCTGGAACGGTGTATTGTTCACTATCCCAAGCGACACCATCACCATCGTCTT
CCTTTCTCTTACCAAAGTAAATACCTCCCACTAATTCTCTGACAACAACGAAGTCAGTACCTTTAG
CAAATTGTGGCTTGATTGGAGATAAGTCTAAAAGAGAGTCGGATGCAAAGTTACATGGTCTTAAG
TTGGCGTACAATTGAAGTTCTTTACGGATTTTTAGTAAACCTTGTTCAGGTCTAACACTACCGGTA
CCCCATTTAGGACCAGCCACAGCACCTAACAAAACGGCATCAACCTTCTTGGAGGCTTCCAGCGC
CTCATCTGGAAGTGGGACACCTGTAGCATCGATAGCAGCACCACCAATTAAATGATTTTCGAAAT
CGAACTTGACATTGGAACGAACATCAGAAATAGCTTTAAGAACCTTAATGGCTTCGGCTGTGATT
TCTTGACCAACGTGGTCACCTGGCAAAACGACGATCTTCTTAGGGGCAGACATAGGGGCAGACA

TTAGAATGGTATATCCTTGAAATATATATATATATTGCTGAAATGTAAAAGGTAAGAAAAGTTAG
AAAGTAAGACGATTGCTAACCACCTATTGGAAAAAACAATAGGTCCTTAAATAATATTGTCAACT
TCAAGTATTGTGATGCAAGCATTTAGTCATGAACGCTTCTCTATTCTATATGAAAAGCCGGTTCCG
GCCTCTCACCTTTCCTTTTTCTCCCAATTTTTCAGTTGAAAAAGGTATATGCGTCAGGCGACCTCT
GAAATTAACAAAAAATTTCCAGTCATCGAATTTGATTCTGTGCGATAGCGCCCCTGTGTGTTCTCG
TTATGTTGAGGAAAAAAATAATGGTTGCTAAGAGATTCGAACTCTTGCATCTTACGATACCTGAG
TATTCCCACAGTTAACTGCGGTCAAGATATTTCTTGAATCAGGCGCCTTAGACCGCTCGGCCAAA
CAACCAATTACTTGTTGAGAAATAGAGTATAATTATCCTATAAATATAACGTTTTTGAACACACA
TGAACAAGGAAGTACAGGACAATTGATTTTGAAGAGAATGTGGATTTTGATGTAATTGTTGGGAT
TCCATTTTTAATAAGGCAATAATATTAGGTATGTGGATATACTAGAAGTTCTCCTCGACCGTCGAT
ATGCGGTGTGAAATACCGCACAGATGCGTAAGGAGAAAATACCGCATCAGGAAATTGTAAACGT
TAATATTTTGTTAAAATTCGCGTTAAATTTTTGTTAAATCAGCTCATTTTTTAACCAATAGGCCGA
AATCGGCAAAATCCCTTATAAATCAAAAGAATAGACCGAGATAGGGTTGAGTGTTGTTCCAGTTT
GGAACAAGAGTCCACTATTAAAGAACGTGGACTCCAACGTCAAAGGGCGAAAAACCGTCTATCA
GGGCGATGGCCCACTACGTGAACCATCACCCTAATCAAGTTTTTTGGGGTCGAGGTGCCGTAAAG
CACTAAATCGGAACCCTAAAGGGAGCCCCCGATTTAGAGCTTGACGGGGAAAGCCGGCGAACGT
GGCGAGAAAGGAAGGGAAGAAAGCGAAAGGAGCGGGCGCTAGGGCGCTGGCAAGTGTAGCGGT
CACGCTGCGCGTAACCACCACACCCGCCGCGCTTAATGCGCCGCTACAGGGCGCGTCGCGCCATT
CGCCATTCAGGCTGCGCAACTGTTGGGAAGGGCGATCGGTGCGGGCCTCTTCGCTATTACGCCAG
CTGAATTGGAGCGACCTCATGCTATACCTGAGAAAGCAACCTGACCTACAGGAAAGAGTTACTC
AAGAATAAGAATTTTCGTTTTAAAACCTAAGAGTCACTTTAAAATTTGTATACACTTATTTTTTT
ATAACTTATTTAATAATAAAAATCATAAATCATAAGAAATTCGCTTATTTAGAAGTGTCAACAAC
GTATCTACCAACGATTTGACCCTTTTCCATCTTTTCGTAAATTTCTGGCAAGGTAGACAAGCCGAC
AACCTTGATTGGAGACTTGACCAAACCTCTGGCGAAGAATTGTTAATTAAGAGCTCAGATCTTAT
CGTCGTCATCCTTGTAATCCATCGATACTAGTGCTTACTCGATGTTGTGGCGGATCTTGAAGTTGG
CCTTGATGCCGTTCTTCTGCTTGTCGGCGGTGATATAGACGTTGTGGCTGTTGTAGTTGTACTCCA
GCTTGTGCCCCAGGATGTTGCCGTCCTCCTTGAAGTCGATGCCCTTCAGCTCGATGCGGTTCACCA
GGGTGTCGCCCTCGAACTTCACCTCGGCGCGGGTCTTGTAGTTGCCGTCGTCCTTGAAGAAGATG
GTGCGCTCCTGGACGTAGCCTTCGGGCATGGCGGACTTGAAGAAGTCGTGCTGCTTCATGTGGTC
GGGGTAGCGGGCGAAGCACTGCAGGCCGTAGCCCAGGGTGGTCACGAGGGTGGGCCAGGGCAC
GGGCAGCTTGCCGGTGGTGCAGATCAGCTTCAGGGTCAGCTTGCCGTAGGTGGCATCGCCCTCGC
CCTCGCCGGACACGCTGAACTTGTGGCCGTTTACGTCGCCGTCCAGCTCGACCAGGATGGGCACC
ACCCCGGTGAACAGCTCCTCGCCCTTGCTCACGAATTCCATAGTACCACCAGAACCAGGAGCTTT
CAAGGTCTTTGCAACGTCAAGGACAAAGCGATATCTAACATCACCCTTGGCCAGACGCTCCAAAG
CTGTGTTGATATTGTCCGCGGAAATGAGTTCTATATCTGCAGTAATGTTATGCTTTCCGGCAAAAT
CAAGCATCTCTTGGGTCTCCTTCAATCCTCCAATGCTACTTCCAGCAACCATCTTCCTCCCCATAA
GCAAAGGAGCAGAGTGAAGATCAAGTGGTTCCGGGGGTGCCCCAAGAAGAACAAGCTTGCCATG
AGACTTTAGTAGACCAAGCAATGGGATAATAGCGTGCTTAGCAGAAATAGTATTAAGTATGCCAT
CAAATGTGCCTGTTGCAGCCTGCAGTGCTTCTGGATTACTGCTCAACAAAAATGCATCTGCACCC
AAACGATTGAGGGCATCGTCTTTCTTGCCCTCAGATGTACTTATAACTGTCACTTTTGCTCCCATG
GCCTTTGCAAACTTAACAGCCACATGGCCAAGTCCACCAAGACCATTAACAGCTATGTGGCTCCC
GGGTTTGGCAAAGCCATAGTATTTCATTGGACTGTACACAGTAATACCGGCACAAAGCAATGGTG
CCCCACCATCAAGTGGTAGGTTCTCTGGGAAACGAACAATAAAGTGTTCATTGCATACCATCTCA
TTGGAATAGCCTCCATAGGTAATCGTTCCATCAACGTTTGGACTTGCATAGGTTAGCACCATTTTG
GGACAATAGTTCTCAAGATCTGCACGACAATTATCACAAGTGCGGCATGAACCAACCAAGCAGC
CAACACCAACTTTATCTCCAACCTTGACCTTTGTAACTTTGCCGCCGACCTCTGTAACTTCCCCTA
CGATTTCATGTCCTGGTACAAGAGGATAGGTCGAAATACCCCACTCATTCTTAGCGAAATGAAGG
TCAGTATGACAAATCCCACAATATAGCACCTTGAACCTCACATCATCCTCAAGAGTTGCCCTCCT
GGAGAATTTGAACGGAGATAAAACCCCAGATGAATCATGAGCAGCCAATCCATAGGTCTTGACT
GGGTGCTCCTCTTCTGGTGATTTTCCGGCCATGCCCTTTAGTGAGGGTTGAATTCGAATTTTCAAA
AATTCTTACTTTTTTTTTGGATGGACGCAAAGAAGTTTAATAATCATATTACATGGCATTACCACC
ATATACATATCCATATACATATCCATATCTAATCTTACTTATATGTTGTGGAAATGTAAAGAGCCC
CATTATCTTAGCCTAAAAAAACCTTCTCTTTGGAACTTTCAGTAATACGCTTAACTGCTCATTGCT
ATATTGAAGTACGGATTAGAAGCCGCCGAGCGGGTGACAGCCCTCCGAAGGAAGACTCTCCTCC
GTGCGTCCTCGTCTTCACCGGTCGCGTTCCTGAAACGCAGATGTGCCTCGCGCCGCACTGCTCCG
AACAATAAAGATTCTACAATACTAGCTTTTATGGTTATGAAGAGGAAAAATTGGCAGTAACCTGG

CCCCACAAACCTTCAAATGAACGAATCAAATTAACAACCATAGGATGATAATGCGATTAGTTTTT
TAGCCTTATTTCTGGGGTAATTAATCAGCGAAGCGATGATTTTTGATCTATTAACAGATATATAAA
TGCAAAAACTGCATAACCACTTTAACTAATACTTTCAACATTTTCGGTTTGTATTACTTCTTATTC
AAATGTAATAAAAGTATCAACAAAAAATTGTTAATATACCTCTATACTTTAACGTCAAGGAGAAA
AAACCCCGGATCCGTAATACGACTCACTATAGGGCCCGGGCGTCGACATGGAACAGAAGTTGAT
TTCCGAAGAAGACCTCGAGTAAGCTTGGTACCGCGGCTAGCTAAGATCCGCTCTAACCGAAAAG
GAAGGAGTTAGACAACCTGAAGTCTAGGTCCCTATTTATTTTTTTATAGTTATGTTAGTATTAAGA
ACGTTATTTATATTTCAAATTTTTCTTTTTTTTCTGTACAGACGCGTGTACGCATGTAACATTATAC
TGAAAACCTTGCTTGAGAAGGTTTTGGGACGCTCGAAGATCCAGCTGCATTAATGAATCGGCCAA
CGCGCGGGGAGAGGCGGTTTGCGTATTGGGCGCTCTTCCGCTTCCTCGCTCACTGACTCGCTGCG
CTCGGTCGTTCGGCTGCGGCGAGCGGTATCAGCTCACTCAAAGGCGGTAATACGGTTATCCACAG
AATCAGGGGATAACGCAGGAAAGAACATGTGAGCAAAAGGCCAGCAAAAGGCCAGGAACCGTA
AAAAGGCCGCGTTGCTGGCGTTTTTCCATAGGCTCCGCCCCCCTGACGAGCATCACAAAAATCGA
CGCTCAAGTCAGAGGTGGCGAAACCCGACAGGACTATAAAGATACCAGGCGTTTCCCCCTGGAA
GCTCCCTCGTGCGCTCTCCTGTTCCGACCCTGCCGCTTACCGGATACCTGTCCGCCTTTCTCCCTTC
GGGAAGCGTGGCGCTTTCTCATAGCTCACGCTGTAGGTATCTCAGTTCGGTGTAGGTCGTTCGCTC
CAAGCTGGGCTGTGTGCACGAACCCCCCGTTCAGCCCGACCGCTGCGCCTTATCCGGTAACTATC
GTCTTGAGTCCAACCCGGTAAGACACGACTTATCGCCACTGGCAGCAGCCACTGGTAACAGGATT
AGCAGAGCGAGGTATGTAGGCGGTGCTACAGAGTTCTTGAAGTGGTGGCCTAACTACGGCTACA
CTAGAAGGACAGTATTTGGTATCTGCGCTCTGCTGAAGCCAGTTACCTTCGGAAAAAGAGTTGGT
AGCTCTTGATCCGGCAAACAAACCACCGCTGGTAGCGGTGGTTTTTTTGTTTGCAAGCAGCAGAT
TACGCGCAGAAAAAAAGGATCTCAAGAAGATCCTTTGATCTTTTCTACGGGGTCTGACGCTCAGT
GGAACGAAAACTCACGTTAAGGGATTTTGGTCATGAGATTATCAAAAAGGATCTTCACCTAGATC
CTTTTAAATTAAAAATGAAGTTTTAAATCAATCTAAAGTATATATGAGTAAACTTGGTCTGACAG
TTACCAATGCTTAATCAGTGAGGCACCTATCTCAGCGATCTGTCTATTTCGTTCATCCATAGTTGC
CTGACTCCCCGTCGTGTAGATAACTACGATACGGGAGGGCTTACCATCTGGCCCCAGTGCTGCAA
TGATACCGCGAGACCCACGCTCACCGGCTCCAGATTTATCAGCAATAAACCAGCCAGCCGGAAG
GGCCGAGCGCAGAAGTGGTCCTGCAACTTTATCCGCCTCCATCCAGTCTATTAATTGTTGCCGGG
AAGCTAGAGTAAGTAGTTCGCCAGTTAATAGTTTGCGCAACGTTGTTGCCATTGCTACAGGCATC
GTGGTGTCACGCTCGTCGTTTGGTATGGCTTCATTCAGCTCCGGTTCCCAACGATCAAGGCGAGTT
ACATGATCCCCCATGTTGTGCAAAAAAGCGGTTAGCTCCTTCGGTCCTCCGATCGTTGTCAGAAG
TAAGTTGGCCGCAGTGTTATCACTCATGGTTATGGCAGCACTGCATAATTCTCTTACTGTCATGCC
ATCCGTAAGATGCTTTTCTGTGACTGGTGAGTACTCAACCAAGTCATTCTGAGAATAGTGTATGC
GGCGACCGAGTTGCTCTTGCCCGGCGTCAATACGGGATAATACCGCGCCACATAGCAGAACTTTA
AAAGTGCTCATCATTGGAAAACGTTCTTCGGGGCGAAAACTCTCAAGGATCTTACCGCTGTTGAG
ATCCAGTTCGATGTAACCCACTCGTGCACCCAACTGATCTTCAGCATCTTTTACTTTCACCAGCGT
TTCTGGGTGAGCAAAAACAGGAAGGCAAAATGCCGCAAAAAAGGGAATAAGGGCGACACGGAA
ATGTTGAATACTCATACTCTTCCTTTTTCAATATTATTGAAGCATTTATCAGGGTTATTGTCTCATG
AGCGGATACATATTTGAATGTATTTAGAAAAATAAACAAATAGGGGTTCCGCGCACATTTCCCCG
AAAAGTGCCACCTGAACGAAGCATCTGTGCTTCATTTTGTAGAACAAAAATGCAACGCGAGAGC
GCTAATTTTTCAAACAAAGAATCTGAGCTGCATTTTTACAGAACAGAAATGCAACGCGAAAGCGC
TATTTTACCAACGAAGAATCTGTGCTTCATTTTTGTAAAACAAAAATGCAACGCGAGAGCGCTAA
TTTTTCAAACAAAGAATCTGAGCTGCATTTTTACAGAACAGAAATGCAACGCGAGAGCGCTATTT
TACCAACAAAGAATCTATACTTCTTTTTTGTTCTACAAAAATGCATCCCGAGAGCGCTATTTTTCT
AACAAAGCATCTTAGATTACTTTTTTTCTCCTTTGTGCGCTCTATAATGCAGTCTCTTGATAACTTT
TTGCACTGTAGGTCCGTTAAGGTTAGAAGAAGGCTACTTTGGTGTCTATTTTCTCTTCCATAAAAA
AAGCCTGACTCCACTTCCCGCGTTTACTGATTACTAGCGAAGCTGCGGGTGCATTTTTTCAAGATA
AAGGCATCCCCGATTATATTCTATACCGATGTGGATTGCGCATACTTTGTGAACAGAAAGTGATA
GCGTTGATGATTCTTCATTGGTCAGAAAATTATGAACGGTTTCTTCTATTTTGTCTCTATATACTAC
GTATAGGAAATGTTTACATTTTCGTATTGTTTTCGATTCACTCTATGAATAGTTCTTACTACAATTT
TTTTGTCTAAAGAGTAATACTAGAGATAAACATAAAAAAATGTAGAGGTCGAGTTTAGATGCAAGT
TCAAGGAGCGAAAGGTGGATGGGTAGGTTATATAGGGATATAGCACAGAGATATATAGCAAAGA
GATACTTTTGAGCAATGTTTGTGGAAGCGGTATTCGCAATATTTTAGTAGCTCGTTACAGTCCGGT
GCGTTTTTGGTTTTTTGAAAGTGCGTCTTCAGAGCGCTTTTGGTTTTCAAAAGCGCTCTGAAGTTC
CTATACTTTCTAGAGAATAGGAACTTCGGAATAGGAACTTCAAAGCGTTTCCGAAAACGAGCGCT
TCCGAAAATGCAACGCGAGCTGCGCACATACAGCTCACTGTTCACGTCGCACCTATATCTGCGTG

TTGCCTGTATATATATATACATGAGAAGAACGGCATAGTGCGTGTTTATGCTTAAATGCGTACTT
ATATGCGTCTATTTATGTAGGATGAAAGGTAGTCTAGTACCTCCTGTGATATTATCCCATTCCATG
CGGGGTATCGTATGCTTCCTTCAGCACTACCCTTTAGCTGTTCTATATGCTGCCACTCCTCAATTG
GATTAGTCTCATCCTTCAATGCTATCATTTCCTTTGATATTGGATCATACTAAGAAACCATTATTA
TCATGACATTAACCTATAAAAATAGGCGTATCACGAGGCCCTTTCGTC

> pESC-LEU-*CrVinBLAST*$^{C51A,\ H56A}$~*mVenus*$^{N}$

TCGCGCGTTTCGGTGATGACGGTGAAAACCTCTGACACATGCAGCTCCCGGAGACGGTCACAGCTTGT
CTGTAAGCGGATGCCGGGAGCAGACAAGCCCGTCAGGGCGCGTCAGCGGGTGTTGGCGGGTGTCGGG
GCTGGCTTAACTATGCGGCATCAGAGCAGATTGTACTGAGAGTGCACCATATCGACTACGTCGTAAGG
CCGTTTCTGACAGAGTAAAATTCTTGAGGGAACTTTCACCATTATGGGAAATGCTTCAAGAAGGTATT
GACTTAAACTCCATCAAATGGTCAGGTCATTGAGTGTTTTTTATTTGTTGTATTTTTTTTTTTTTAGAGA
AAATCCTCCAATATCAAATTAGGAATCGTAGTTTCATGATTTTCTGTTACACCTAACTTTTTGTGTGGTG
CCCTCCTCCTTGTCAATATTAATGTTAAAGTGCAATTCTTTTTCCTTATCACGTTGAGCCATTAGTATCA
ATTTGCTTACCTGTATTCCTTTACTATCCTCCTTTTTCTCCTTCTTGATAAATGTATGTAGATTGCGTATA
TAGTTTCGTCTACCCTATGAACATATTCCATTTTGTAATTTCGTGTCGTTTCTATTATGAATTTCATTTAT
AAAGTTTATGTACAAATATCATAAAAAAAGAGAATCTTTTTAAGCAAGGATTTTCTTAACTTCTTCGGC
GACAGCATCACCGACTTCGGTGGTACTGTTGGAACCACCTAAATCACCAGTTCTGATACCTGCATCCA
AAACCTTTTTAACTGCATCTTCAATGGCCTTACCTTCTTCAGGCAAGTTCAATGACAATTTCAACATCA
TTGCAGCAGACAAGATAGTGGCGATAGGGTCAACCTTATTCTTTGGCAAATCTGGAGCAGAACCGTGG
CATGGTTCGTACAAACCAAATGCGGTGTTCTTGTCTGGCAAAGAGGCCAAGGACGCAGATGGCAACA
AACCCAAGGAACCTGGGATAACGGAGGCTTCATCGGAGATGATATCACCAAACATGTTGCTGGTGATT
ATAATACCATTTAGGTGGGTTGGGTTCTTAACTAGGATCATGGCGGCAGAATCAATCAATTGATGTTG
AACCTTCAATGTAGGGAATTCGTTCTTGATGGTTTCCTCCACAGTTTTTCTCCATAATCTTGAAGAGGC
CAAAAGATTAGCTTTATCCAAGGACCAAATAGGCAATGGTGGCTCATGTTGTAGGGCCATGAAAGCGG
CCATTCTTGTGATTCTTTGCACTTCTGGAACGGTGTATTGTTCACTATCCCAAGCGACACCATCACCAT
CGTCTTCCTTTCTCTTACCAAAGTAAATACCTCCCACTAATTCTCTGACAACAACGAAGTCAGTACCTT
TAGCAAATTGTGGCTTGATTGGAGATAAGTCTAAAAGAGAGTCGGATGCAAAGTTACATGGTCTTAAG
TTGGCGTACAATTGAAGTTCTTTACGGATTTTTAGTAAACCTTGTTCAGGTCTAACACTACCGGTACCC
CATTTAGGACCAGCCACAGCACCTAACAAAACGGCATCAACCTTCTTGGAGGCTTCCAGCGCCTCATC
TGGAAGTGGGACACCTGTAGCATCGATAGCAGCACCACCAATTAAATGATTTTCGAAATCGAACTTGA
CATTGGAACGAACATCAGAAATAGCTTTAAGAACCTTAATGGCTTCGGCTGTGATTTCTTGACCAACG
TGGTCACCTGGCAAAACGACGATCTTCTTAGGGGCAGACATAGGGGCAGACATTAGAATGGTATATCC
TTGAAATATATATATATATTGCTGAAATGTAAAAGGTAAGAAAAGTTAGAAAGTAAGACGATTGCTAA
CCACCTATTGGAAAAAAACAATAGGTCCTTAAATAATATTGTCAACTTCAAGTATTGTGATGCAAGCAT
TTAGTCATGAACGCTTCTCTATTCTATATGAAAAGCCGGTTCCGGCCTCTCACCTTTCCTTTTTCTCCCA
ATTTTTCAGTTGAAAAAGGTATATGCGTCAGGCGACCTCTGAAATTAACAAAAAATTTCCAGTCATCG
AATTTGATTCTGTGCGATAGCGCCCCTGTGTGTTCTCGTTATGTTGAGGAAAAAAATAATGGTTGCTAA
GAGATTCGAACTCTTGCATCTTACGATACCTGAGTATTCCCACAGTTAACTGCGGTCAAGATATTTCTT
GAATCAGGCGCCTTAGACCGCTCGGCCAAACAACCAATTACTTGTTGAGAAATAGAGTATAATTATCC
TATAAATATAACGTTTTTGAACACACATGAACAAGGAAGTACAGGACAATTGATTTTGAAGAGAATGT
GGATTTTGATGTAATTGTTGGGATTCCATTTTTAATAAGGCAATAATATTAGGTATGTGGATATACTAG
AAGTTCTCCTCGACCGTCGATATGCGGTGTGAAATACCGCACAGATGCGTAAGGAGAAAATACCGCAT
CAGGAAATTGTAAACGTTAATATTTTGTTAAAATTCGCGTTAAATTTTTGTTAAATCAGCTCATTTTTTA
ACCAATAGGCCGAAATCGGCAAAATCCCTTATAAATCAAAAGAATAGACCGAGATAGGGTTGAGTGT
TGTTCCAGTTTGGAACAAGAGTCCACTATTAAAGAACGTGGACTCCAACGTCAAAGGGCGAAAAACC
GTCTATCAGGGCGATGGCCCACTACGTGAACCATCACCCTAATCAAGTTTTTTGGGGTCGAGGTGCCG
TAAAGCACTAAATCGGAACCCTAAAGGGAGCCCCCGATTTAGAGCTTGACGGGGAAAGCCGGCGAAC
GTGGCGAGAAAGGAAGGGAAGAAAGCGAAAGGAGCGGGCGCTAGGGCGCTGGCAAGTGTAGCGGTC
ACGCTGCGCGTAACCACCACACCCGCCGCGCTTAATGCGCCGCTACAGGGCGCGTCGCGCCATTCGCC
ATTCAGGCTGCGCAACTGTTGGGAAGGGCGATCGGTGCGGGCCTCTTCGCTATTACGCCAGCTGAATT
GGAGCGACCTCATGCTATACCTGAGAAAGCAACCTGACCTACAGGAAAGAGTTACTCAAGAATAAGA
ATTTTCGTTTTAAAACCTAAGAGTCACTTTAAAATTTGTATACACTTATTTTTTTTATAACTTATTTAAT
AATAAAAATCATAAATCATAAGAAATTCGCTTATTTAGAAGTGTCAACAACGTATCTACCAACGATTT
GACCCTTTTCCATCTTTTCGTAAATTTCTGGCAAGGTAGACAAGCCGACAACCTTGATTGGAGACTTGA

CCAAACCTCTGGCGAAGAATTGTTAATTAAGAGCTCAGATCTTATCGTCGTCATCCTTGTAATCCATCG
ATACTAGTGCTTACTCGATGTTGTGGCGGATCTTGAAGTTGGCCTTGATGCCGTTCTTCTGCTTGTCGG
CGGTGATATAGACGTTGTGGCTGTTGTAGTTGTACTCCAGCTTGTGCCCCAGGATGTTGCCGTCCTCCT
TGAAGTCGATGCCCTTCAGCTCGATGCGGTTCACCAGGGTGTCGCCCTCGAACTTCACCTCGGCGCGG
GTCTTGTAGTTGCCGTCGTCCTTGAAGAAGATGGTGCGCTCCTGGACGTAGCCTTCGGGCATGGCGGA
CTTGAAGAAGTCGTGCTGCTTCATGTGGTCGGGGTAGCGGGCGAAGCACTGCAGGCCGTAGCCCAGGG
TGGTCACGAGGGTGGGCCAGGGCACGGGCAGCTTGCCGGTGGTGCAGATCAGCTTCAGGGTCAGCTTG
CCGTAGGTGGCATCGCCCTCGCCCTCGCCGGACACGCTGAACTTGTGGCCGTTTACGTCGCCGTCCAGC
TCGACCAGGATGGGCACCACCCCGGTGAACAGCTCCTCGCCCTTGCTCACGAATTCCATAGTACCACC
AGAACCAGGAGCTTTCAAGGTCTTTGCAACGTCAAGGACAAAGCGATATCTAACATCACCCTTGGCCA
GACGCTCCAAAGCTGTGTTGATATTGTCCGCGGAAATGAGTTCTATATCTGCAGTAATGTTATGCTTTC
CGGCAAAATCAAGCATCTCTTGGGTCTCCTTCAATCCTCCAATGCTACTTCCAGCAACCATCTTCCTCC
CCATAAGCAAAGGAGCAGAGTGAAGATCAAGTGGTTCCGGGGGTGCCCCAAGAAGAACAAGCTTGCC
ATGAGACTTTAGTAGACCAAGCAATGGGATAATAGCGTGCTTAGCAGAAATAGTATTAAGTATGCCAT
CAAATGTGCCTGTTGCAGCCTGCAGTGCTTCTGGATTACTGCTCAACAAAAATGCATCTGCACCCAAA
CGATTGAGGGCATCGTCTTTCTTGCCCTCAGATGTACTTATAACTGTCACTTTTGCTCCCATGGCCTTTG
CAAACTTAACAGCCACATGGCCAAGTCCACCAAGACCATTAACAGCTATGTGGCTCCCGGGTTTGGCA
AAGCCATAGTATTTCATTGGACTGTACACAGTAATACCGGCACAAAGCAATGGTGCCCCACCATCAAG
TGGTAGGTTCTCTGGGAAACGAACAATAAAGTGTTCATTGCATACCATCTCATTGGAATAGCCTCCAT
AGGTAATCGTTCCATCAACGTTTGGACTTGCATAGGTTAGCACCATTTTGGGACAATAGTTCTCAAGAT
CTGCACGACAATTATCACAAGTGCGGCATGAACCAACCAAGCAGCCAACACCAACTTTATCTCCAACC
TTGACCTTTGTAACTTTGCCGCCGACCTCTGTAACTTCCCCTACGATTTCATGTCCTGGTACAAGAGGA
TAGGTCGAAATACCCCACTCATTCTTAGCGAAAGCAAGGTCAGTATGAGCAATCCCACAATATAGCAC
CTTGAACCTCACATCATCCTCAAGAGTTGCCCTCCTGGAGAATTTGAACGGAGATAAAACCCCAGATG
AATCATGAGCAGCCAATCCATAGGTCTTGACTGGGTGCTCCTCTTCTGGTGATTTTCCGGCCATGCCCT
TTAGTGAGGGTTGAATTCGAATTTTCAAAAATTCTTACTTTTTTTTTGGATGGACGCAAAGAAGTTTAA
TAATCATATTACATGGCATTACCACCATATACATATCCATATACATATCCATATCTAATCTTACTTATAT
GTTGTGGAAATGTAAAGAGCCCCATTATCTTAGCCTAAAAAAACCTTCTCTTTGGAACTTTCAGTAATA
CGCTTAACTGCTCATTGCTATATTGAAGTACGGATTAGAAGCCGCCGAGCGGGTGACAGCCCTCCGAA
GGAAGACTCTCCTCCGTGCGTCCTCGTCTTCACCGGTCGCGTTCCTGAAACGCAGATGTGCCTCGCGCC
GCACTGCTCCGAACAATAAAGATTCTACAATACTAGCTTTTATGGTTATGAAGAGGAAAAATTGGCAG
TAACCTGGCCCCACAAACCTTCAAATGAACGAATCAAATTAACAACCATAGGATGATAATGCGATTAG
TTTTTTAGCCTTATTTCTGGGGTAATTAATCAGCGAAGCGATGATTTTTGATCTATTAACAGATATATA
AATGCAAAAACTGCATAACCACTTTAACTAATACTTTCAACATTTTCGGTTTGTATTACTTCTTATTCAA
ATGTAATAAAAGTATCAACAAAAAATTGTTAATATACCTCTATACTTTAACGTCAAGGAGAAAAAACC
CCGGATCCGTAATACGACTCACTATAGGGCCCGGGCGTCGACATGGAACAGAAGTTGATTTCCGAAGA
AGACCTCGAGTAAGCTTGGTACCGCGGCTAGCTAAGATCCGCTCTAACCGAAAAGGAAGGAGTTAGA
CAACCTGAAGTCTAGGTCCCTATTTATTTTTTTATAGTTATGTTAGTATTAAGAACGTTATTTATATTTC
AAATTTTTCTTTTTTTTCTGTACAGACGCGTGTACGCATGTAACATTATACTGAAAACCTTGCTTGAGA
AGGTTTTGGGACGCTCGAAGATCCAGCTGCATTAATGAATCGGCCAACGCGCGGGGAGAGGCGGTTTG
CGTATTGGGCGCTCTTCCGCTTCCTCGCTCACTGACTCGCTGCGCTCGGTCGTTCGGCTGCGGCGAGCG
GTATCAGCTCACTCAAAGGCGGTAATACGGTTATCCACAGAATCAGGGGATAACGCAGGAAAGAACA
TGTGAGCAAAAGGCCAGCAAAAGGCCAGGAACCGTAAAAAGGCCGCGTTGCTGGCGTTTTTCCATAG
GCTCCGCCCCCCTGACGAGCATCACAAAAATCGACGCTCAAGTCAGAGGTGGCGAAACCCGACAGGA
CTATAAAGATACCAGGCGTTTCCCCCTGGAAGCTCCCTCGTGCGCTCTCCTGTTCCGACCCTGCCGCTT
ACCGGATACCTGTCCGCCTTTCTCCCTTCGGGAAGCGTGGCGCTTTCTCATAGCTCACGCTGTAGGTAT
CTCAGTTCGGTGTAGGTCGTTCGCTCCAAGCTGGGCTGTGTGCACGAACCCCCCGTTCAGCCCGACCGC
TGCGCCTTATCCGGTAACTATCGTCTTGAGTCCAACCCGGTAAGACACGACTTATCGCCACTGGCAGC
AGCCACTGGTAACAGGATTAGCAGAGCGAGGTATGTAGGCGGTGCTACAGAGTTCTTGAAGTGGTGG
CCTAACTACGGCTACACTAGAAGGACAGTATTTGGTATCTGCGCTCTGCTGAAGCCAGTTACCTTCGG
AAAAAGAGTTGGTAGCTCTTGATCCGGCAAACAAACCACCGCTGGTAGCGGTGGTTTTTTTGTTTGCA
AGCAGCAGATTACGCGCAGAAAAAAAGGATCTCAAGAAGATCCTTTGATCTTTTCTACGGGGTCTGAC
GCTCAGTGGAACGAAAACTCACGTTAAGGGATTTTGGTCATGAGATTATCAAAAAGGATCTTCACCTA
GATCCTTTTAAATTAAAAATGAAGTTTTAAATCAATCTAAAGTATATATGAGTAAACTTGGTCTGACAG
TTACCAATGCTTAATCAGTGAGGCACCTATCTCAGCGATCTGTCTATTTCGTTCATCCATAGTTGCCTG
ACTCCCCGTCGTGTAGATAACTACGATACGGGAGGGCTTACCATCTGGCCCCAGTGCTGCAATGATAC

CGCGAGACCCACGCTCACCGGCTCCAGATTTATCAGCAATAAACCAGCCAGCCGGAAGGGCCGAGCG
CAGAAGTGGTCCTGCAACTTTATCCGCCTCCATCCAGTCTATTAATTGTTGCCGGGAAGCTAGAGTAAG
TAGTTCGCCAGTTAATAGTTTGCGCAACGTTGTTGCCATTGCTACAGGCATCGTGGTGTCACGCTCGTC
GTTTGGTATGGCTTCATTCAGCTCCGGTTCCCAACGATCAAGGCGAGTTACATGATCCCCCATGTTGTG
CAAAAAAGCGGTTAGCTCCTTCGGTCCTCCGATCGTTGTCAGAAGTAAGTTGGCCGCAGTGTTATCACT
CATGGTTATGGCAGCACTGCATAATTCTCTTACTGTCATGCCATCCGTAAGATGCTTTTCTGTGACTGG
TGAGTACTCAACCAAGTCATTCTGAGAATAGTGTATGCGGCGACCGAGTTGCTCTTGCCCGGCGTCAA
TACGGGATAATACCGCGCCACATAGCAGAACTTTAAAAGTGCTCATCATTGGAAAACGTTCTTCGGGG
CGAAAACTCTCAAGGATCTTACCGCTGTTGAGATCCAGTTCGATGTAACCCACTCGTGCACCCAACTG
ATCTTCAGCATCTTTTACTTTCACCAGCGTTTCTGGGTGAGCAAAAACAGGAAGGCAAAATGCCGCAA
AAAAGGGAATAAGGGCGACACGGAAATGTTGAATACTCATACTCTTCCTTTTTCAATATTATTGAAGC
ATTTATCAGGGTTATTGTCTCATGAGCGGATACATATTTGAATGTATTTAGAAAAATAAACAAATAGG
GGTTCCGCGCACATTTCCCCGAAAAGTGCCACCTGAACGAAGCATCTGTGCTTCATTTTGTAGAACAA
AAATGCAACGCGAGAGCGCTAATTTTTCAAACAAAGAATCTGAGCTGCATTTTTACAGAACAGAAATG
CAACGCGAAAGCGCTATTTTACCAACGAAGAATCTGTGCTTCATTTTTGTAAAACAAAAATGCAACGC
GAGAGCGCTAATTTTTCAAACAAAGAATCTGAGCTGCATTTTTACAGAACAGAAATGCAACGCGAGAG
CGCTATTTTACCAACAAAGAATCTATACTTCTTTTTTGTTCTACAAAAATGCATCCCGAGAGCGCTATT
TTTCTAACAAAGCATCTTAGATTACTTTTTTTCTCCTTTGTGCGCTCTATAATGCAGTCTCTTGATAACT
TTTTGCACTGTAGGTCCGTTAAGGTTAGAAGAAGGCTACTTTGGTGTCTATTTTCTCTTCCATAAAAAA
AGCCTGACTCCACTTCCCGCGTTTACTGATTACTAGCGAAGCTGCGGGTGCATTTTTTCAAGATAAAGG
CATCCCCGATTATATTCTATACCGATGTGGATTGCGCATACTTTGTGAACAGAAAGTGATAGCGTTGAT
GATTCTTCATTGGTCAGAAAATTATGAACGGTTTCTTCTATTTTGTCTCTATATACTACGTATAGGAAAT
GTTTACATTTTCGTATTGTTTTCGATTCACTCTATGAATAGTTCTTACTACAATTTTTTTGTCTAAAGAG
TAATACTAGAGATAAACATAAAAAATGTAGAGGTCGAGTTTAGATGCAAGTTCAAGGAGCGAAAGGT
GGATGGGTAGGTTATATAGGGATATAGCACAGAGATATATAGCAAAGAGATACTTTTGAGCAATGTTT
GTGGAAGCGGTATTCGCAATATTTTAGTAGCTCGTTACAGTCCGGTGCGTTTTTGGTTTTTTGAAAGTG
CGTCTTCAGAGCGCTTTTGGTTTTCAAAAGCGCTCTGAAGTTCCTATACTTTCTAGAGAATAGGAACTT
CGGAATAGGAACTTCAAAGCGTTTCCGAAAACGAGCGCTTCCGAAAATGCAACGCGAGCTGCGCACA
TACAGCTCACTGTTCACGTCGCACCTATATCTGCGTGTTGCCTGTATATATATATACATGAGAAGAACG
GCATAGTGCGTGTTTATGCTTAAATGCGTACTTATATGCGTCTATTTATGTAGGATGAAAGGTAGTCTA
GTACCTCCTGTGATATTATCCCATTCCATGCGGGGTATCGTATGCTTCCTTCAGCACTACCCTTTAGCTG
TTCTATATGCTGCCACTCCTCAATTGGATTAGTCTCATCCTTCAATGCTATCATTTCCTTTGATATTGGA
TCATACTAAGAAACCATTATTATCATGACATTAACCTATAAAAATAGGCGTATCACGAGGCCCTTTCGT
C

> pESC-LEU-*CrVinBLAST*$^{M298E,\ V299E}$~*mVenus*$^{N}$

TCGCGCGTTTCGGTGATGACGGTGAAAACCTCTGACACATGCAGCTCCCGGAGACGGTCACAGCTTGT
CTGTAAGCGGATGCCGGGAGCAGACAAGCCCGTCAGGGCGCGTCAGCGGGTGTTGGCGGGTGTCGGG
GCTGGCTTAACTATGCGGCATCAGAGCAGATTGTACTGAGAGTGCACCATATCGACTACGTCGTAAGG
CCGTTTCTGACAGAGTAAAATTCTTGAGGGAACTTTCACCATTATGGGAAATGCTTCAAGAAGGTATT
GACTTAAACTCCATCAAATGGTCAGGTCATTGAGTGTTTTTTATTTGTTGTATTTTTTTTTTTTTAGAGA
AAATCCTCCAATATCAAATTAGGAATCGTAGTTTCATGATTTTCTGTTACACCTAACTTTTTGTGTGGTG
CCCTCCTCCTTGTCAATATTAATGTTAAAGTGCAATTCTTTTTCCTTATCACGTTGAGCCATTAGTATCA
ATTTGCTTACCTGTATTCCTTTACTATCCTCCTTTTTCTCCTTCTTGATAAATGTATGTAGATTGCGTATA
TAGTTTCGTCTACCCTATGAACATATTCCATTTTGTAATTTCGTGTCGTTTCTATTATGAATTTCATTTAT
AAAGTTTATGTACAAATATCATAAAAAAAGAGAATCTTTTTAAGCAAGGATTTTCTTAACTTCTTCGGC
GACAGCATCACCGACTTCGGTGGTACTGTTGGAACCACCTAAATCACCAGTTCTGATACCTGCATCCA
AAACCTTTTTAACTGCATCTTCAATGGCCTTACCTTCTTCAGGCAAGTTCAATGACAATTTCAACATCA
TTGCAGCAGACAAGATAGTGGCGATAGGGTCAACCTTATTCTTTGGCAAATCTGGAGCAGAACCGTGG
CATGGTTCGTACAAACCAAATGCGGTGTTCTTGTCTGGCAAAGAGGCCAAGGACGCAGATGGCAACA
AACCCAAGGAACCTGGGATAACGGAGGCTTCATCGGAGATGATATCACCAAACATGTTGCTGGTGATT
ATAATACCATTTAGGTGGGTTGGGTTCTTAACTAGGATCATGGCGGCAGAATCAATCAATTGATGTTG
AACCTTCAATGTAGGGAATTCGTTCTTGATGGTTTCCTCCACAGTTTTTCTCCATAATCTTGAAGAGGC
CAAAAGATTAGCTTTATCCAAGGACCAAATAGGCAATGGTGGCTCATGTTGTAGGGCCATGAAAGCGG
CCATTCTTGTGATTCTTTGCACTTCTGGAACGGTGTATTGTTCACTATCCCAAGCGACACCATCACCAT

CGTCTTCCTTTCTCTTACCAAAGTAAATACCTCCCACTAATTCTCTGACAACAACGAAGTCAGTACCTT
TAGCAAATTGTGGCTTGATTGGAGATAAGTCTAAAAGAGAGTCGGATGCAAAGTTACATGGTCTTAAG
TTGGCGTACAATTGAAGTTCTTTACGGATTTTTAGTAAACCTTGTTCAGGTCTAACACTACCGGTACCC
CATTTAGGACCAGCCACAGCACCTAACAAAACGGCATCAACCTTCTTGGAGGCTTCCAGCGCCTCATC
TGGAAGTGGGACACCTGTAGCATCGATAGCAGCACCACCAATTAAATGATTTTCGAAATCGAACTTGA
CATTGGAACGAACATCAGAAATAGCTTTAAGAACCTTAATGGCTTCGGCTGTGATTTCTTGACCAACG
TGGTCACCTGGCAAAACGACGATCTTCTTAGGGGCAGACATAGGGGCAGACATTAGAATGGTATATCC
TTGAAATATATATATATATTGCTGAAATGTAAAAGGTAAGAAAAGTTAGAAAGTAAGACGATTGCTAA
CCACCTATTGGAAAAAACAATAGGTCCTTAAATAATATTGTCAACTTCAAGTATTGTGATGCAAGCAT
TTAGTCATGAACGCTTCTCTATTCTATATGAAAAGCCGGTTCCGGCCTCTCACCTTTCCTTTTTCTCCCA
ATTTTTCAGTTGAAAAAGGTATATGCGTCAGGCGACCTCTGAAATTAACAAAAAATTTCCAGTCATCG
AATTTGATTCTGTGCGATAGCGCCCCTGTGTGTTCTCGTTATGTTGAGGAAAAAAATAATGGTTGCTAA
GAGATTCGAACTCTTGCATCTTACGATACCTGAGTATTCCCACAGTTAACTGCGGTCAAGATATTTCTT
GAATCAGGCGCCTTAGACCGCTCGGCCAAACAACCAATTACTTGTTGAGAAATAGAGTATAATTATCC
TATAAATATAACGTTTTTGAACACACATGAACAAGGAAGTACAGGACAATTGATTTTGAAGAGAATGT
GGATTTTGATGTAATTGTTGGGATTCCATTTTTAATAAGGCAATAATATTAGGTATGTGGATATACTAG
AAGTTCTCCTCGACCGTCGATATGCGGTGTGAAATACCGCACAGATGCGTAAGGAGAAAATACCGCAT
CAGGAAATTGTAAACGTTAATATTTTGTTAAAATTCGCGTTAAATTTTTGTTAAATCAGCTCATTTTTTA
ACCAATAGGCCGAAATCGGCAAAATCCCTTATAAATCAAAAGAATAGACCGAGATAGGGTTGAGTGT
TGTTCCAGTTTGGAACAAGAGTCCACTATTAAAGAACGTGGACTCCAACGTCAAAGGGCGAAAAACC
GTCTATCAGGGCGATGGCCCACTACGTGAACCATCACCCTAATCAAGTTTTTTGGGGTCGAGGTGCCG
TAAAGCACTAAATCGGAACCCTAAAGGGAGCCCCCGATTTAGAGCTTGACGGGGAAAGCCGGCGAAC
GTGGCGAGAAAGGAAGGGAAGAAAGCGAAAGGAGCGGGCGCTAGGGCGCTGGCAAGTGTAGCGGTC
ACGCTGCGCGTAACCACCACACCCGCCGCGCTTAATGCGCCGCTACAGGGCGCGTCGCGCCATTCGCC
ATTCAGGCTGCGCAACTGTTGGGAAGGGCGATCGGTGCGGGCCTCTTCGCTATTACGCCAGCTGAATT
GGAGCGACCTCATGCTATACCTGAGAAAGCAACCTGACCTACAGGAAAGAGTTACTCAAGAATAAGA
ATTTTCGTTTTAAAACCTAAGAGTCACTTTAAAATTTGTATACACTTATTTTTTTTATAACTTATTTAAT
AATAAAAATCATAAATCATAAGAAATTCGCTTATTTAGAAGTGTCAACAACGTATCTACCAACGATTT
GACCCTTTTCCATCTTTTCGTAAATTTCTGGCAAGGTAGACAAGCCGACAACCTTGATTGGAGACTTGA
CCAAACCTCTGGCGAAGAATTGTTAATTAAGAGCTCAGATCTTATCGTCGTCATCCTTGTAATCCATCG
ATACTAGTGCTTACTCGATGTTGTGGCGGATCTTGAAGTTGGCCTTGATGCCGTTCTTCTGCTTGTCGG
CGGTGATATAGACGTTGTGGCTGTTGTAGTTGTACTCCAGCTTGTGCCCCAGGATGTTGCCGTCCTCCT
TGAAGTCGATGCCCTTCAGCTCGATGCGGTTCACCAGGGTGTCGCCCTCGAACTTCACCTCGGCGCGG
GTCTTGTAGTTGCCGTCGTCCTTGAAGAAGATGGTGCGCTCCTGGACGTAGCCTTCGGGCATGGCGGA
CTTGAAGAAGTCGTGCTGCTTCATGTGGTCGGGGTAGCGGGCGAAGCACTGCAGGCCGTAGCCCAGGG
TGGTCACGAGGGTGGGCCAGGGCACGGGCAGCTTGCCGGTGGTGCAGATCAGCTTCAGGGTCAGCTTG
CCGTAGGTGGCATCGCCCTCGCCCTCGCCGGACACGCTGAACTTGTGGCCGTTTACGTCGCCGTCCAGC
TCGACCAGGATGGGCACCACCCCGGTGAACAGCTCCTCGCCCTTGCTCACGAATTCCATAGTACCACC
AGAACCAGGAGCTTTCAAGGTCTTTGCAACGTCAAGGACAAAGCGATATCTAACATCACCCTTGGCCA
GACGCTCCAAAGCTGTGTTGATATTGTCCGCGGAAATGAGTTCTATATCTGCAGTAATGTTATGCTTTC
CGGCAAAATCAAGCATCTCTTGGGTCTCCTTCAATCCTCCAATGCTACTTCCAGCTTCTTCCTTCCTCCC
CATAAGCAAAGGAGCAGAGTGAAGATCAAGTGGTTCCGGGGGTGCCCCAAGAAGAACAAGCTTGCCA
TGAGACTTTAGTAGACCAAGCAATGGGATAATAGCGTGCTTAGCAGAAATAGTATTAAGTATGCCATC
AAATGTGCCTGTTGCAGCCTGCAGTGCTTCTGGATTACTGCTCAACAAAAATGCATCTGCACCCAAAC
GATTGAGGGCATCGTCTTTCTTGCCCTCAGATGTACTTATAACTGTCACTTTTGCTCCCATGGCCTTTGC
AAACTTAACAGCCACATGGCCAAGTCCACCAAGACCATTAACAGCTATGTGGCTCCCGGGTTTGGCAA
AGCCATAGTATTTCATTGGACTGTACACAGTAATACCGGCACAAAGCAATGGTGCCCCACCATCAAGT
GGTAGGTTCTCTGGGAAACGAACAATAAAGTGTTCATTGCATACCATCTCATTGGAATAGCCTCCATA
GGTAATCGTTCCATCAACGTTTGGACTTGCATAGGTTAGCACCATTTTGGGACAATAGTTCTCAAGATC
TGCACGACAATTATCACAAGTGCGGCATGAACCAACCAAGCAGCCAACACCAACTTTATCTCCAACCT
TGACCTTTGTAACTTTGCCGCCGACCTCTGTAACTTCCCCTACGATTTCATGTCCTGGTACAAGAGGAT
AGGTCGAAATACCCCACTCATTCTTAGCGAAATGAAGGTCAGTATGACAAATCCCACAATATAGCACC
TTGAACCTCACATCATCCTCAAGAGTTGCCCTCCTGGAGAATTTGAACGGAGATAAAACCCCAGATGA
ATCATGAGCAGCCAATCCATAGGTCTTGACTGGGTGCTCCTCTTCTGGTGATTTTCCGGCCATGCCCTT
TAGTGAGGGTTGAATTCGAATTTTCAAAAATTCTTACTTTTTTTTTGGATGGACGCAAAGAAGTTTAAT
AATCATATTACATGGCATTACCACCATATACATATCCATATACATATCCATATCTAATCTTACTTATAT

GTTGTGGAAATGTAAAGAGCCCCATTATCTTAGCCTAAAAAAACCTTCTCTTTGGAACTTTCAGTAATA
CGCTTAACTGCTCATTGCTATATTGAAGTACGGATTAGAAGCCGCCGAGCGGGTGACAGCCCTCCGAA
GGAAGACTCTCCTCCGTGCGTCCTCGTCTTCACCGGTCGCGTTCCTGAAACGCAGATGTGCCTCGCGCC
GCACTGCTCCGAACAATAAAGATTCTACAATACTAGCTTTTATGGTTATGAAGAGGAAAAATTGGCAG
TAACCTGGCCCCACAAACCTTCAAATGAACGAATCAAATTAACAACCATAGGATGATAATGCGATTAG
TTTTTTAGCCTTATTTCTGGGGTAATTAATCAGCGAAGCGATGATTTTTGATCTATTAACAGATATATA
AATGCAAAAACTGCATAACCACTTTAACTAATACTTTCAACATTTTCGGTTTGTATTACTTCTTATTCAA
ATGTAATAAAAGTATCAACAAAAAATTGTTAATATACCTCTATACTTTAACGTCAAGGAGAAAAAACC
CCGGATCCGTAATACGACTCACTATAGGGCCCGGGCGTCGACATGGAACAGAAGTTGATTTCCGAAGA
AGACCTCGAGTAAGCTTGGTACCGCGGCTAGCTAAGATCCGCTCTAACCGAAAAGGAAGGAGTTAGA
CAACCTGAAGTCTAGGTCCCTATTTATTTTTTTATAGTTATGTTAGTATTAAGAACGTTATTTATATTTC
AAATTTTTCTTTTTTTTCTGTACAGACGCGTGTACGCATGTAACATTATACTGAAAACCTTGCTTGAGA
AGGTTTTGGGACGCTCGAAGATCCAGCTGCATTAATGAATCGGCCAACGCGCGGGGAGAGGCGGTTTG
CGTATTGGGCGCTCTTCCGCTTCCTCGCTCACTGACTCGCTGCGCTCGGTCGTTCGGCTGCGGCGAGCG
GTATCAGCTCACTCAAAGGCGGTAATACGGTTATCCACAGAATCAGGGGATAACGCAGGAAAGAACA
TGTGAGCAAAAGGCCAGCAAAAGGCCAGGAACCGTAAAAAGGCCGCGTTGCTGGCGTTTTTCCATAG
GCTCCGCCCCCCTGACGAGCATCACAAAAATCGACGCTCAAGTCAGAGGTGGCGAAACCCGACAGGA
CTATAAAGATACCAGGCGTTTCCCCCTGGAAGCTCCCTCGTGCGCTCTCCTGTTCCGACCCTGCCGCTT
ACCGGATACCTGTCCGCCTTTCTCCCTTCGGGAAGCGTGGCGCTTTCTCATAGCTCACGCTGTAGGTAT
CTCAGTTCGGTGTAGGTCGTTCGCTCCAAGCTGGGCTGTGTGCACGAACCCCCCGTTCAGCCCGACCGC
TGCGCCTTATCCGGTAACTATCGTCTTGAGTCCAACCCGGTAAGACACGACTTATCGCCACTGGCAGC
AGCCACTGGTAACAGGATTAGCAGAGCGAGGTATGTAGGCGGTGCTACAGAGTTCTTGAAGTGGTGG
CCTAACTACGGCTACACTAGAAGGACAGTATTTGGTATCTGCGCTCTGCTGAAGCCAGTTACCTTCGG
AAAAAGAGTTGGTAGCTCTTGATCCGGCAAACAAACCACCGCTGGTAGCGGTGGTTTTTTTGTTTGCA
AGCAGCAGATTACGCGCAGAAAAAAAGGATCTCAAGAAGATCCTTTGATCTTTTCTACGGGGTCTGAC
GCTCAGTGGAACGAAAACTCACGTTAAGGGATTTTGGTCATGAGATTATCAAAAAGGATCTTCACCTA
GATCCTTTTAAATTAAAAATGAAGTTTTAAATCAATCTAAAGTATATATGAGTAAACTTGGTCTGACAG
TTACCAATGCTTAATCAGTGAGGCACCTATCTCAGCGATCTGTCTATTTCGTTCATCCATAGTTGCCTG
ACTCCCCGTCGTGTAGATAACTACGATACGGGAGGGCTTACCATCTGGCCCCAGTGCTGCAATGATAC
CGCGAGACCCACGCTCACCGGCTCCAGATTTATCAGCAATAAACCAGCCAGCCGGAAGGGCCGAGCG
CAGAAGTGGTCCTGCAACTTTATCCGCCTCCATCCAGTCTATTAATTGTTGCCGGGAAGCTAGAGTAAG
TAGTTCGCCAGTTAATAGTTTGCGCAACGTTGTTGCCATTGCTACAGGCATCGTGGTGTCACGCTCGTC
GTTTGGTATGGCTTCATTCAGCTCCGGTTCCCAACGATCAAGGCGAGTTACATGATCCCCCATGTTGTG
CAAAAAAGCGGTTAGCTCCTTCGGTCCTCCGATCGTTGTCAGAAGTAAGTTGGCCGCAGTGTTATCACT
CATGGTTATGGCAGCACTGCATAATTCTCTTACTGTCATGCCATCCGTAAGATGCTTTTCTGTGACTGG
TGAGTACTCAACCAAGTCATTCTGAGAATAGTGTATGCGGCGACCGAGTTGCTCTTGCCCGGCGTCAA
TACGGGATAATACCGCGCCACATAGCAGAACTTTAAAAGTGCTCATCATTGGAAAACGTTCTTCGGGG
CGAAAACTCTCAAGGATCTTACCGCTGTTGAGATCCAGTTCGATGTAACCCACTCGTGCACCCAACTG
ATCTTCAGCATCTTTTACTTTCACCAGCGTTTCTGGGTGAGCAAAAACAGGAAGGCAAAATGCCGCAA
AAAAGGGAATAAGGGCGACACGGAAATGTTGAATACTCATACTCTTCCTTTTTCAATATTATTGAAGC
ATTTATCAGGGTTATTGTCTCATGAGCGGATACATATTTGAATGTATTTAGAAAAATAAACAAATAGG
GGTTCCGCGCACATTTCCCCGAAAAGTGCCACCTGAACGAAGCATCTGTGCTTCATTTTGTAGAACAA
AAATGCAACGCGAGAGCGCTAATTTTTCAAACAAAGAATCTGAGCTGCATTTTTACAGAACAGAAATG
CAACGCGAAAGCGCTATTTTACCAACGAAGAATCTGTGCTTCATTTTTGTAAAACAAAAATGCAACGC
GAGAGCGCTAATTTTTCAAACAAAGAATCTGAGCTGCATTTTTACAGAACAGAAATGCAACGCGAGAG
CGCTATTTTACCAACAAAGAATCTATACTTCTTTTTTGTTCTACAAAAATGCATCCCGAGAGCGCTATT
TTTCTAACAAAGCATCTTAGATTACTTTTTTTCTCCTTTGTGCGCTCTATAATGCAGTCTCTTGATAACT
TTTTGCACTGTAGGTCCGTTAAGGTTAGAAGAAGGCTACTTTGGTGTCTATTTTCTCTTCCATAAAAAA
AGCCTGACTCCACTTCCCGCGTTTACTGATTACTAGCGAAGCTGCGGGTGCATTTTTTCAAGATAAAGG
CATCCCCGATTATATTCTATACCGATGTGGATTGCGCATACTTTGTGAACAGAAAGTGATAGCGTTGAT
GATTCTTCATTGGTCAGAAAATTATGAACGGTTTCTTCTATTTTGTCTCTATATACTACGTATAGGAAAT
GTTTACATTTTCGTATTGTTTTCGATTCACTCTATGAATAGTTCTTACTACAATTTTTTTGTCTAAAGAG
TAATACTAGAGATAAACATAAAAAATGTAGAGGTCGAGTTTAGATGCAAGTTCAAGGAGCGAAAGGT
GGATGGGTAGGTTATATAGGGATATAGCACAGAGATATATAGCAAAGAGATACTTTTGAGCAATGTTT
GTGGAAGCGGTATTCGCAATATTTTAGTAGCTCGTTACAGTCCGGTGCGTTTTTGGTTTTTTGAAAGTG
CGTCTTCAGAGCGCTTTTGGTTTTCAAAAGCGCTCTGAAGTTCCTATACTTTCTAGAGAATAGGAACTT

CGGAATAGGAACTTCAAAGCGTTTCCGAAAACGAGCGCTTCCGAAAATGCAACGCGAGCTGCGCACA
TACAGCTCACTGTTCACGTCGCACCTATATCTGCGTGTTGCCTGTATATATATATACATGAGAAGAACG
GCATAGTGCGTGTTTATGCTTAAATGCGTACTTATATGCGTCTATTTATGTAGGATGAAAGGTAGTCTA
GTACCTCCTGTGATATTATCCCATTCCATGCGGGGTATCGTATGCTTCCTTCAGCACTACCCTTTAGCTG
TTCTATATGCTGCCACTCCTCAATTGGATTAGTCTCATCCTTCAATGCTATCATTTCCTTTGATATTGGA
TCATACTAAGAAACCATTATTATCATGACATTAACCTATAAAAATAGGCGTATCACGAGGCCCTTTCGT
C

>pESC-LEU-*CrVinBLAST*$^{K359G}$~*mVenus*$^{N}$

TCGCGCGTTTCGGTGATGACGGTGAAAACCTCTGACACATGCAGCTCCCGGAGACGGTCACAGCTTGT
CTGTAAGCGGATGCCGGGAGCAGACAAGCCCGTCAGGGCGCGTCAGCGGGTGTTGGCGGGTGTCGGG
GCTGGCTTAACTATGCGGCATCAGAGCAGATTGTACTGAGAGTGCACCATATCGACTACGTCGTAAGG
CCGTTTCTGACAGAGTAAAATTCTTGAGGGAACTTTCACCATTATGGGAAATGCTTCAAGAAGGTATT
GACTTAAACTCCATCAAATGGTCAGGTCATTGAGTGTTTTTTATTTGTTGTATTTTTTTTTTTTTAGAGA
AAATCCTCCAATATCAAATTAGGAATCGTAGTTTCATGATTTTCTGTTACACCTAACTTTTTGTGTGGTG
CCCTCCTCCTTGTCAATATTAATGTTAAAGTGCAATTCTTTTTCCTTATCACGTTGAGCCATTAGTATCA
ATTTGCTTACCTGTATTCCTTTACTATCCTCCTTTTTCTCCTTCTTGATAAATGTATGTAGATTGCGTATA
TAGTTTCGTCTACCCTATGAACATATTCCATTTTGTAATTTCGTGTCGTTTCTATTATGAATTTCATTTAT
AAAGTTTATGTACAAATATCATAAAAAAAGAGAATCTTTTTAAGCAAGGATTTTCTTAACTTCTTCGGC
GACAGCATCACCGACTTCGGTGGTACTGTTGGAACCACCTAAATCACCAGTTCTGATACCTGCATCCA
AAACCTTTTTAACTGCATCTTCAATGGCCTTACCTTCTTCAGGCAAGTTCAATGACAATTTCAACATCA
TTGCAGCAGACAAGATAGTGGCGATAGGGTCAACCTTATTCTTTGGCAAATCTGGAGCAGAACCGTGG
CATGGTTCGTACAAACCAAATGCGGTGTTCTTGTCTGGCAAAGAGGCCAAGGACGCAGATGGCAACA
AACCCAAGGAACCTGGGATAACGGAGGCTTCATCGGAGATGATATCACCAAACATGTTGCTGGTGATT
ATAATACCATTTAGGTGGGTTGGGTTCTTAACTAGGATCATGGCGGCAGAATCAATCAATTGATGTTG
AACCTTCAATGTAGGGAATTCGTTCTTGATGGTTTCCTCCACAGTTTTTCTCCATAATCTTGAAGAGGC
CAAAAGATTAGCTTTATCCAAGGACCAAATAGGCAATGGTGGCTCATGTTGTAGGGCCATGAAAGCGG
CCATTCTTGTGATTCTTTGCACTTCTGGAACGGTGTATTGTTCACTATCCCAAGCGACACCATCACCAT
CGTCTTCCTTTCTCTTACCAAAGTAAATACCTCCCACTAATTCTCTGACAACAACGAAGTCAGTACCTT
TAGCAAATTGTGGCTTGATTGGAGATAAGTCTAAAAGAGAGTCGGATGCAAAGTTACATGGTCTTAAG
TTGGCGTACAATTGAAGTTCTTTACGGATTTTTAGTAAACCTTGTTCAGGTCTAACACTACCGGTACCC
CATTTAGGACCAGCCACAGCACCTAACAAAACGGCATCAACCTTCTTGGAGGCTTCCAGCGCCTCATC
TGGAAGTGGGACACCTGTAGCATCGATAGCAGCACCACCAATTAAATGATTTTCGAAATCGAACTTGA
CATTGGAACGAACATCAGAAATAGCTTTAAGAACCTTAATGGCTTCGGCTGTGATTTCTTGACCAACG
TGGTCACCTGGCAAAACGACGATCTTCTTAGGGGCAGACATAGGGGCAGACATTAGAATGGTATATCC
TTGAAATATATATATATATATTGCTGAAATGTAAAAGGTAAGAAAAGTTAGAAAGTAAGACGATTGCTAA
CCACCTATTGGAAAAAAACAATAGGTCCTTAAATAATATTGTCAACTTCAAGTATTGTGATGCAAGCAT
TTAGTCATGAACGCTTCTCTATTCTATATGAAAAGCCGGTTCCGGCCTCTCACCTTTCCTTTTTCTCCCA
ATTTTTCAGTTGAAAAAGGTATATGCGTCAGGCGACCTCTGAAATTAACAAAAAATTTCCAGTCATCG
AATTTGATTCTGTGCGATAGCGCCCCTGTGTGTTCTCGTTATGTTGAGGAAAAAAATAATGGTTGCTAA
GAGATTCGAACTCTTGCATCTTACGATACCTGAGTATTCCCACAGTTAACTGCGGTCAAGATATTTCTT
GAATCAGGCGCCTTAGACCGCTCGGCCAAACAACCAATTACTTGTTGAGAAATAGAGTATAATTATCC
TATAAATATAACGTTTTTGAACACACATGAACAAGGAAGTACAGGACAATTGATTTTGAAGAGAATGT
GGATTTTGATGTAATTGTTGGGATTCCATTTTTAATAAGGCAATAATATTAGGTATGTGGATATACTAG
AAGTTCTCCTCGACCGTCGATATGCGGTGTGAAATACCGCACAGATGCGTAAGGAGAAAATACCGCAT
CAGGAAATTGTAAACGTTAATATTTTGTTAAAATTCGCGTTAAATTTTTGTTAAATCAGCTCATTTTTTA
ACCAATAGGCCGAAATCGGCAAAATCCCTTATAAATCAAAAGAATAGACCGAGATAGGGTTGAGTGT
TGTTCCAGTTTGGAACAAGAGTCCACTATTAAAGAACGTGGACTCCAACGTCAAAGGGCGAAAAACC
GTCTATCAGGGCGATGGCCCACTACGTGAACCATCACCCTAATCAAGTTTTTTGGGGTCGAGGTGCCG
TAAAGCACTAAATCGGAACCCTAAAGGGAGCCCCCGATTTAGAGCTTGACGGGGAAAGCCGGCGAAC
GTGGCGAGAAAGGAAGGGAAGAAAGCGAAAGGAGCGGGCGCTAGGGCGCTGGCAAGTGTAGCGGTC
ACGCTGCGCGTAACCACCACACCCGCCGCGCTTAATGCGCCGCTACAGGGCGCGTCGCGCCATTCGCC
ATTCAGGCTGCGCAACTGTTGGGAAGGGCGATCGGTGCGGGCCTCTTCGCTATTACGCCAGCTGAATT
GGAGCGACCTCATGCTATACCTGAGAAAGCAACCTGACCTACAGGAAAGAGTTACTCAAGAATAAGA
ATTTTCGTTTTAAAACCTAAGAGTCACTTTAAAATTTGTATACACTTATTTTTTTTATAACTTATTTAAT

AATAAAAATCATAAATCATAAGAAATTCGCTTATTTAGAAGTGTCAACAACGTATCTACCAACGATTT
GACCCTTTTCCATCTTTTCGTAAATTTCTGGCAAGGTAGACAAGCCGACAACCTTGATTGGAGACTTGA
CCAAACCTCTGGCGAAGAATTGTTAATTAAGAGCTCAGATCTTATCGTCGTCATCCTTGTAATCCATCG
ATACTAGTGCTTACTCGATGTTGTGGCGGATCTTGAAGTTGGCCTTGATGCCGTTCTTCTGCTTGTCGG
CGGTGATATAGACGTTGTGGCTGTTGTAGTTGTACTCCAGCTTGTGCCCCAGGATGTTGCCGTCCTCCT
TGAAGTCGATGCCCTTCAGCTCGATGCGGTTCACCAGGGTGTCGCCCTCGAACTTCACCTCGGCGCGG
GTCTTGTAGTTGCCGTCGTCCTTGAAGAAGATGGTGCGCTCCTGGACGTAGCCTTCGGGCATGGCGGA
CTTGAAGAAGTCGTGCTGCTTCATGTGGTCGGGGTAGCGGGCGAAGCACTGCAGGCCGTAGCCCAGGG
TGGTCACGAGGGTGGGCCAGGGCACGGGCAGCTTGCCGGTGGTGCAGATCAGCTTCAGGGTCAGCTTG
CCGTAGGTGGCATCGCCCTCGCCCTCGCCGGACACGCTGAACTTGTGGCCGTTTACGTCGCCGTCCAGC
TCGACCAGGATGGGCACCACCCCGGTGAACAGCTCCTCGCCCTTGCTCACGAATTCCATAGTACCACC
AGAACCAGGAGCACCCAAGGTCTTTGCAACGTCAAGGACAAAGCGATATCTAACATCACCCTTGGCCA
GACGCTCCAAAGCTGTGTTGATATTGTCCGCGGAAATGAGTTCTATATCTGCAGTAATGTTATGCTTTC
CGGCAAAATCAAGCATCTCTTGGGTCTCCTTCAATCCTCCAATGCTACTTCCAGCAACCATCTTCCTCC
CCATAAGCAAAGGAGCAGAGTGAAGATCAAGTGGTTCCGGGGGTGCCCCAAGAAGAACAAGCTTGCC
ATGAGACTTTAGTAGACCAAGCAATGGGATAATAGCGTGCTTAGCAGAAATAGTATTAAGTATGCCAT
CAAATGTGCCTGTTGCAGCCTGCAGTGCTTCTGGATTACTGCTCAACAAAAATGCATCTGCACCCAAA
CGATTGAGGGCATCGTCTTTCTTGCCCTCAGATGTACTTATAACTGTCACTTTTGCTCCCATGGCCTTTG
CAAACTTAACAGCCACATGGCCAAGTCCACCAAGACCATTAACAGCTATGTGGCTCCCGGGTTTGGCA
AAGCCATAGTATTTCATTGGACTGTACACAGTAATACCGGCACAAAGCAATGGTGCCCCACCATCAAG
TGGTAGGTTCTCTGGGAAACGAACAATAAAGTGTTCATTGCATACCATCTCATTGGAATAGCCTCCAT
AGGTAATCGTTCCATCAACGTTTGGACTTGCATAGGTTAGCACCATTTTGGGACAATAGTTCTCAAGAT
CTGCACGACAATTATCACAAGTGCGGCATGAACCAACCAAGCAGCCAACACCAACTTTATCTCCAACC
TTGACCTTTGTAACTTTGCCGCCGACCTCTGTAACTTCCCCTACGATTTCATGTCCTGGTACAAGAGGA
TAGGTCGAAATACCCCACTCATTCTTAGCGAAATGAAGGTCAGTATGACAAATCCCACAATATAGCAC
CTTGAACCTCACATCATCCTCAAGAGTTGCCCTCCTGGAGAATTTGAACGGAGATAAAACCCCAGATG
AATCATGAGCAGCCAATCCATAGGTCTTGACTGGGTGCTCCTCTTCTGGTGATTTTCCGGCCATGCCCT
TTAGTGAGGGTTGAATTCGAATTTTCAAAAATTCTTACTTTTTTTTTGGATGGACGCAAAGAAGTTTAA
TAATCATATTACATGGCATTACCACCATATACATATCCATATACATATCCATATCTAATCTTACTTATAT
GTTGTGGAAATGTAAAGAGCCCCATTATCTTAGCCTAAAAAAACCTTCTCTTTGGAACTTTCAGTAATA
CGCTTAACTGCTCATTGCTATATTGAAGTACGGATTAGAAGCCGCCGAGCGGGTGACAGCCCTCCGAA
GGAAGACTCTCCTCCGTGCGTCCTCGTCTTCACCGGTCGCGTTCCTGAAACGCAGATGTGCCTCGCGCC
GCACTGCTCCGAACAATAAAGATTCTACAATACTAGCTTTTATGGTTATGAAGAGGAAAAATTGGCAG
TAACCTGGCCCCACAAACCTTCAAATGAACGAATCAAATTAACAACCATAGGATGATAATGCGATTAG
TTTTTTAGCCTTATTTCTGGGGTAATTAATCAGCGAAGCGATGATTTTTGATCTATTAACAGATATATA
AATGCAAAAACTGCATAACCACTTTAACTAATACTTTCAACATTTTCGGTTTGTATTACTTCTTATTCAA
ATGTAATAAAAGTATCAACAAAAAATTGTTAATATACCTCTATACTTTAACGTCAAGGAGAAAAAACC
CCGGATCCGTAATACGACTCACTATAGGGCCCGGGCGTCGACATGGAACAGAAGTTGATTTCCGAAGA
AGACCTCGAGTAAGCTTGGTACCGCGGCTAGCTAAGATCCGCTCTAACCGAAAAGGAAGGAGTTAGA
CAACCTGAAGTCTAGGTCCCTATTTATTTTTTTATAGTTATGTTAGTATTAAGAACGTTATTTATATTTC
AAATTTTTCTTTTTTTTCTGTACAGACGCGTGTACGCATGTAACATTATACTGAAAACCTTGCTTGAGA
AGGTTTTGGGACGCTCGAAGATCCAGCTGCATTAATGAATCGGCCAACGCGCGGGGAGAGGCGGTTTG
CGTATTGGGCGCTCTTCCGCTTCCTCGCTCACTGACTCGCTGCGCTCGGTCGTTCGGCTGCGGCGAGCG
GTATCAGCTCACTCAAAGGCGGTAATACGGTTATCCACAGAATCAGGGGATAACGCAGGAAAGAACA
TGTGAGCAAAAGGCCAGCAAAAGGCCAGGAACCGTAAAAAGGCCGCGTTGCTGGCGTTTTTCCATAG
GCTCCGCCCCCCTGACGAGCATCACAAAAATCGACGCTCAAGTCAGAGGTGGCGAAACCCGACAGGA
CTATAAAGATACCAGGCGTTTCCCCCTGGAAGCTCCCTCGTGCGCTCTCCTGTTCCGACCCTGCCGCTT
ACCGGATACCTGTCCGCCTTTCTCCCTTCGGGAAGCGTGGCGCTTTCTCATAGCTCACGCTGTAGGTAT
CTCAGTTCGGTGTAGGTCGTTCGCTCCAAGCTGGGCTGTGTGCACGAACCCCCCGTTCAGCCCGACCGC
TGCGCCTTATCCGGTAACTATCGTCTTGAGTCCAACCCGGTAAGACACGACTTATCGCCACTGGCAGC
AGCCACTGGTAACAGGATTAGCAGAGCGAGGTATGTAGGCGGTGCTACAGAGTTCTTGAAGTGGTGG
CCTAACTACGGCTACACTAGAAGGACAGTATTTGGTATCTGCGCTCTGCTGAAGCCAGTTACCTTCGG
AAAAAGAGTTGGTAGCTCTTGATCCGGCAAACAAACCACCGCTGGTAGCGGTGGTTTTTTTGTTTGCA
AGCAGCAGATTACGCGCAGAAAAAAAGGATCTCAAGAAGATCCTTTGATCTTTTCTACGGGGTCTGAC
GCTCAGTGGAACGAAAACTCACGTTAAGGGATTTTGGTCATGAGATTATCAAAAAGGATCTTCACCTA
GATCCTTTTAAATTAAAAATGAAGTTTTAAATCAATCTAAAGTATATATGAGTAAACTTGGTCTGACAG

TTACCAATGCTTAATCAGTGAGGCACCTATCTCAGCGATCTGTCTATTTCGTTCATCCATAGTTGCCTG
ACTCCCCGTCGTGTAGATAACTACGATACGGGAGGGCTTACCATCTGGCCCCAGTGCTGCAATGATAC
CGCGAGACCCACGCTCACCGGCTCCAGATTTATCAGCAATAAACCAGCCAGCCGGAAGGGCCGAGCG
CAGAAGTGGTCCTGCAACTTTATCCGCCTCCATCCAGTCTATTAATTGTTGCCGGGAAGCTAGAGTAAG
TAGTTCGCCAGTTAATAGTTTGCGCAACGTTGTTGCCATTGCTACAGGCATCGTGGTGTCACGCTCGTC
GTTTGGTATGGCTTCATTCAGCTCCGGTTCCCAACGATCAAGGCGAGTTACATGATCCCCCATGTTGTG
CAAAAAAGCGGTTAGCTCCTTCGGTCCTCCGATCGTTGTCAGAAGTAAGTTGGCCGCAGTGTTATCACT
CATGGTTATGGCAGCACTGCATAATTCTCTTACTGTCATGCCATCCGTAAGATGCTTTTCTGTGACTGG
TGAGTACTCAACCAAGTCATTCTGAGAATAGTGTATGCGGCGACCGAGTTGCTCTTGCCCGGCGTCAA
TACGGGATAATACCGCGCCACATAGCAGAACTTTAAAAGTGCTCATCATTGGAAAACGTTCTTCGGGG
CGAAAACTCTCAAGGATCTTACCGCTGTTGAGATCCAGTTCGATGTAACCCACTCGTGCACCCAACTG
ATCTTCAGCATCTTTTACTTTCACCAGCGTTTCTGGGTGAGCAAAAACAGGAAGGCAAAATGCCGCAA
AAAAGGGAATAAGGGCGACACGGAAATGTTGAATACTCATACTCTTCCTTTTTCAATATTATTGAAGC
ATTTATCAGGGTTATTGTCTCATGAGCGGATACATATTTGAATGTATTTAGAAAAATAAACAAATAGG
GGTTCCGCGCACATTTCCCCGAAAAGTGCCACCTGAACGAAGCATCTGTGCTTCATTTTGTAGAACAA
AAATGCAACGCGAGAGCGCTAATTTTTCAAACAAAGAATCTGAGCTGCATTTTTACAGAACAGAAATG
CAACGCGAAAGCGCTATTTTACCAACGAAGAATCTGTGCTTCATTTTTGTAAAACAAAAATGCAACGC
GAGAGCGCTAATTTTTCAAACAAAGAATCTGAGCTGCATTTTTACAGAACAGAAATGCAACGCGAGAG
CGCTATTTTACCAACAAAGAATCTATACTTCTTTTTTGTTCTACAAAAATGCATCCCGAGAGCGCTATT
TTTCTAACAAAGCATCTTAGATTACTTTTTTTCTCCTTTGTGCGCTCTATAATGCAGTCTCTTGATAACT
TTTTGCACTGTAGGTCCGTTAAGGTTAGAAGAAGGCTACTTTGGTGTCTATTTTCTCTTCCATAAAAAA
AGCCTGACTCCACTTCCCGCGTTTACTGATTACTAGCGAAGCTGCGGGTGCATTTTTTCAAGATAAAGG
CATCCCCGATTATATTCTATACCGATGTGGATTGCGCATACTTTGTGAACAGAAAGTGATAGCGTTGAT
GATTCTTCATTGGTCAGAAAATTATGAACGGTTTCTTCTATTTTGTCTCTATATACTACGTATAGGAAAT
GTTTACATTTTCGTATTGTTTTCGATTCACTCTATGAATAGTTCTTACTACAATTTTTTTGTCTAAAGAG
TAATACTAGAGATAAACATAAAAAATGTAGAGGTCGAGTTTAGATGCAAGTTCAAGGAGCGAAAGGT
GGATGGGTAGGTTATATAGGGATATAGCACAGAGATATATAGCAAAGAGATACTTTTGAGCAATGTTT
GTGGAAGCGGTATTCGCAATATTTTAGTAGCTCGTTACAGTCCGGTGCGTTTTTGGTTTTTTGAAAGTG
CGTCTTCAGAGCGCTTTTGGTTTTCAAAAGCGCTCTGAAGTTCCTATACTTTCTAGAGAATAGGAACTT
CGGAATAGGAACTTCAAAGCGTTTCCGAAAACGAGCGCTTCCGAAAATGCAACGCGAGCTGCGCACA
TACAGCTCACTGTTCACGTCGCACCTATATCTGCGTGTTGCCTGTATATATATATACATGAGAAGAACG
GCATAGTGCGTGTTTATGCTTAAATGCGTACTTATATGCGTCTATTTATGTAGGATGAAAGGTAGTCTA
GTACCTCCTGTGATATTATCCCATTCCATGCGGGGTATCGTATGCTTCCTTCAGCACTACCCTTTAGCTG
TTCTATATGCTGCCACTCCTCAATTGGATTAGTCTCATCCTTCAATGCTATCATTTCCTTTGATATTGGA
TCATACTAAGAAACCATTATTATCATGACATTAACCTATAAAAATAGGCGTATCACGAGGCCCTTTCGT
C

> pESC-LEU-*GS~mVenus*$^{N}$

TCGCGCGTTTCGGTGATGACGGTGAAAACCTCTGACACATGCAGCTCCCGGAGACGGTCACAGCTTGT
CTGTAAGCGGATGCCGGGAGCAGACAAGCCCGTCAGGGCGCGTCAGCGGGTGTTGGCGGGTGTCGGG
GCTGGCTTAACTATGCGGCATCAGAGCAGATTGTACTGAGAGTGCACCATATCGACTACGTCGTAAGG
CCGTTTCTGACAGAGTAAAATTCTTGAGGGAACTTTCACCATTATGGGAAATGCTTCAAGAAGGTATT
GACTTAAACTCCATCAAATGGTCAGGTCATTGAGTGTTTTTTATTTGTTGTATTTTTTTTTTTTTAGAGA
AAATCCTCCAATATCAAATTAGGAATCGTAGTTTCATGATTTTCTGTTACACCTAACTTTTTGTGTGGTG
CCCTCCTCCTTGTCAATATTAATGTTAAAGTGCAATTCTTTTTCCTTATCACGTTGAGCCATTAGTATCA
ATTTGCTTACCTGTATTCCTTTACTATCCTCCTTTTTCTCCTTCTTGATAAATGTATGTAGATTGCGTATA
TAGTTTCGTCTACCCTATGAACATATTCCATTTTGTAATTTCGTGTCGTTTCTATTATGAATTTCATTTAT
AAAGTTTATGTACAAATATCATAAAAAAAGAGAATCTTTTTAAGCAAGGATTTTCTTAACTTCTTCGGC
GACAGCATCACCGACTTCGGTGGTACTGTTGGAACCACCTAAATCACCAGTTCTGATACCTGCATCCA
AAACCTTTTTAACTGCATCTTCAATGGCCTTACCTTCTTCAGGCAAGTTCAATGACAATTTCAACATCA
TTGCAGCAGACAAGATAGTGGCGATAGGGTCAACCTTATTCTTTGGCAAATCTGGAGCAGAACCGTGG
CATGGTTCGTACAAACCAAATGCGGTGTTCTTGTCTGGCAAAGAGGCCAAGGACGCAGATGGCAACA
AACCCAAGGAACCTGGGATAACGGAGGCTTCATCGGAGATGATATCACCAAACATGTTGCTGGTGATT
ATAATACCATTTAGGTGGGTTGGGTTCTTAACTAGGATCATGGCGGCAGAATCAATCAATTGATGTTG
AACCTTCAATGTAGGGAATTCGTTCTTGATGGTTTCCTCCACAGTTTTTCTCCATAATCTTGAAGAGGC

CAAAAGATTAGCTTTATCCAAGGACCAAATAGGCAATGGTGGCTCATGTTGTAGGGCCATGAAAGCGG
CCATTCTTGTGATTCTTTGCACTTCTGGAACGGTGTATTGTTCACTATCCCAAGCGACACCATCACCAT
CGTCTTCCTTTCTCTTACCAAAGTAAATACCTCCCACTAATTCTCTGACAACAACGAAGTCAGTACCTT
TAGCAAATTGTGGCTTGATTGGAGATAAGTCTAAAAGAGAGTCGGATGCAAAGTTACATGGTCTTAAG
TTGGCGTACAATTGAAGTTCTTTACGGATTTTTAGTAAACCTTGTTCAGGTCTAACACTACCGGTACCC
CATTTAGGACCAGCCACAGCACCTAACAAAACGGCATCAACCTTCTTGGAGGCTTCCAGCGCCTCATC
TGGAAGTGGGACACCTGTAGCATCGATAGCAGCACCACCAATTAAATGATTTTCGAAATCGAACTTGA
CATTGGAACGAACATCAGAAATAGCTTTAAGAACCTTAATGGCTTCGGCTGTGATTTCTTGACCAACG
TGGTCACCTGGCAAAACGACGATCTTCTTAGGGGCAGACATAGGGGCAGACATTAGAATGGTATATCC
TTGAAATATATATATATATTGCTGAAATGTAAAAGGTAAGAAAAGTTAGAAAGTAAGACGATTGCTAA
CCACCTATTGGAAAAAACAATAGGTCCTTAAATAATATTGTCAACTTCAAGTATTGTGATGCAAGCAT
TTAGTCATGAACGCTTCTCTATTCTATATGAAAAGCCGGTTCCGGCCTCTCACCTTTCCTTTTTCTCCCA
ATTTTTCAGTTGAAAAAGGTATATGCGTCAGGCGACCTCTGAAATTAACAAAAAATTTCCAGTCATCG
AATTTGATTCTGTGCGATAGCGCCCCTGTGTGTTCTCGTTATGTTGAGGAAAAAAATAATGGTTGCTAA
GAGATTCGAACTCTTGCATCTTACGATACCTGAGTATTCCCACAGTTAACTGCGGTCAAGATATTTCTT
GAATCAGGCGCCTTAGACCGCTCGGCCAAACAACCAATTACTTGTTGAGAAATAGAGTATAATTATCC
TATAAATATAACGTTTTTGAACACACATGAACAAGGAAGTACAGGACAATTGATTTTGAAGAGAATGT
GGATTTTGATGTAATTGTTGGGATTCCATTTTTAATAAGGCAATAATATTAGGTATGTGGATATACTAG
AAGTTCTCCTCGACCGTCGATATGCGGTGTGAAATACCGCACAGATGCGTAAGGAGAAAATACCGCAT
CAGGAAATTGTAAACGTTAATATTTTGTTAAAATTCGCGTTAAATTTTTGTTAAATCAGCTCATTTTTTA
ACCAATAGGCCGAAATCGGCAAAATCCCTTATAAATCAAAAGAATAGACCGAGATAGGGTTGAGTGT
TGTTCCAGTTTGGAACAAGAGTCCACTATTAAAGAACGTGGACTCCAACGTCAAAGGGCGAAAAACC
GTCTATCAGGGCGATGGCCCACTACGTGAACCATCACCCTAATCAAGTTTTTTGGGGTCGAGGTGCCG
TAAAGCACTAAATCGGAACCCTAAAGGGAGCCCCCGATTTAGAGCTTGACGGGGAAAGCCGGCGAAC
GTGGCGAGAAAGGAAGGGAAGAAAGCGAAAGGAGCGGGCGCTAGGGCGCTGGCAAGTGTAGCGGTC
ACGCTGCGCGTAACCACCACACCCGCCGCGCTTAATGCGCCGCTACAGGGCGCGTCGCGCCATTCGCC
ATTCAGGCTGCGCAACTGTTGGGAAGGGCGATCGGTGCGGGCCTCTTCGCTATTACGCCAGCTGAATT
GGAGCGACCTCATGCTATACCTGAGAAAGCAACCTGACCTACAGGAAAGAGTTACTCAAGAATAAGA
ATTTTCGTTTTAAAACCTAAGAGTCACTTTAAAATTTGTATACACTTATTTTTTTTATAACTTATTTAAT
AATAAAAATCATAAATCATAAGAAATTCGCTTATTTAGAAGTGTCAACAACGTATCTACCAACGATTT
GACCCTTTTCCATCTTTTCGTAAATTTCTGGCAAGGTAGACAAGCCGACAACCTTGATTGGAGACTTGA
CCAAACCTCTGGCGAAGAATTGTTAATTAAGAGCTCAGATCTTATCGTCGTCATCCTTGTAATCCATCG
ATACTAGTGCTTACTCGATGTTGTGGCGGATCTTGAAGTTGGCCTTGATGCCGTTCTTCTGCTTGTCGG
CGGTGATATAGACGTTGTGGCTGTTGTAGTTGTACTCCAGCTTGTGCCCCAGGATGTTGCCGTCCTCCT
TGAAGTCGATGCCCTTCAGCTCGATGCGGTTCACCAGGGTGTCGCCCTCGAACTTCACCTCGGCGCGG
GTCTTGTAGTTGCCGTCGTCCTTGAAGAAGATGGTGCGCTCCTGGACGTAGCCTTCGGGCATGGCGGA
CTTGAAGAAGTCGTGCTGCTTCATGTGGTCGGGGTAGCGGGCGAAGCACTGCAGGCCGTAGCCCAGGG
TGGTCACGAGGGTGGGCCAGGGCACGGGCAGCTTGCCGGTGGTGCAGATCAGCTTCAGGGTCAGCTTG
CCGTAGGTGGCATCGCCCTCGCCCTCGCCGGACACGCTGAACTTGTGGCCGTTTACGTCGCCGTCCAGC
TCGACCAGGATGGGCACCACCCCGGTGAACAGCTCCTCGCCCTTGCTCACGAATTCCATAGTACCACC
AGAACCTTCCTCAAATTTCAATGTATTTCCAATGTCAATCGCAAAACGGTACTTGACATCCAAATTCTT
GATACGTTCCATAGCAGTGCTGAGATAGTCAATCCCAATAACTTCAGTATCACATACAATGTTATGTTT
GGCTGCGAAATCAAGCATTTCTTGGTACTCTTTGAGACCTCCAGTGGAACTTCCGATTATCTTTTTCCTT
CCCATAATGAGAGGTGCCGCAGGGAGCTCAAATAGCGACTCCGGTGCACCTACGAGCATAACCGCGC
CGTCAAACTTGAGCAAATTGAGCATAAGTGACATAGGAGTGCGGCCACCTGGGGTGGTGTCCACAAC
ACCATCCATAGTACCTGCCAGAGCCTTCAATTGCTCAGAGTCAGTGTTGACAACAAAAGCATCAGCAC
CATGTTCTTCAATGGCTTCCTTCTCCTTACGCCTTGATGTACTAATAACAGTAGCCTTACCACCAAAAG
CCTTAATAAACTTAACAGCAACAGAACCAAGACCTCCCAGCCCGAAAACCCCAATATGCTTTCCTGGC
TTATCGAGTCCCAAATGTTTCATTGGGCTATAAACAACAACCCCAGCACAAAGGAGAGCAACCCCTTT
ATCTTGAGGCAAGTTTTCGGGCCATCGGAGGACGAACTTTTCATCAACAACCATCACATTTGAACAAC
CCCCATAGGATCGTTCCCCTTGCTCACGGTAAACAGATCCATCAGCCATATTGGGCTCTGGGCAGTAAT
TCTCCATTCCACTTTGACAATTATAACATTGACCACAAGATCCGACCATACATCCCACAGCTACCTTGT
CTCCAACCTTGAATTTCTCTACTTTGCTGCCAACTTCTACCACCTCACCGGCAGTTTCATGTCCAAACAC
ATAAGGATATCTGGTGAAACCCCACTTGTTTCTGACCATTTCCATATCGAAATTGCAAACACCAGAGT
ACAAAACTCTAATCTTCACATCCCGTTCACCAGGGACTCTTCTATAGAACTTGATGGGCTGAAGGACA
CCAGATGCATCTGCAGCACCCCATCCCACAGCCTTCACTGAAAGGTCGAGTTTGGTTGTTTCTCCGGCC

ATGCCCTTTAGTGAGGGTTGAATTCGAATTTTCAAAAATTCTTACTTTTTTTTTGGATGGACGCAAAGA
AGTTTAATAATCATATTACATGGCATTACCACCATATACATATCCATATACATATCCATATCTAATCTT
ACTTATATGTTGTGGAAATGTAAAGAGCCCCATTATCTTAGCCTAAAAAAACCTTCTCTTTGGAACTTT
CAGTAATACGCTTAACTGCTCATTGCTATATTGAAGTACGGATTAGAAGCCGCCGAGCGGGTGACAGC
CCTCCGAAGGAAGACTCTCCTCCGTGCGTCCTCGTCTTCACCGGTCGCGTTCCTGAAACGCAGATGTGC
CTCGCGCCGCACTGCTCCGAACAATAAAGATTCTACAATACTAGCTTTTATGGTTATGAAGAGGAAAA
ATTGGCAGTAACCTGGCCCCACAAACCTTCAAATGAACGAATCAAATTAACAACCATAGGATGATAAT
GCGATTAGTTTTTTAGCCTTATTTCTGGGGTAATTAATCAGCGAAGCGATGATTTTTGATCTATTAACA
GATATATAAATGCAAAAACTGCATAACCACTTTAACTAATACTTTCAACATTTTCGGTTTGTATTACTT
CTTATTCAAATGTAATAAAAGTATCAACAAAAAAATTGTTAATATACCTCTATACTTTAACGTCAAGGA
GAAAAAACCCCGGATCCGTAATACGACTCACTATAGGGCCCGGGCGTCGACATGGAACAGAAGTTGA
TTTCCGAAGAAGACCTCGAGTAAGCTTGGTACCGCGGCTAGCTAAGATCCGCTCTAACCGAAAAGGAA
GGAGTTAGACAACCTGAAGTCTAGGTCCCTATTTATTTTTTTATAGTTATGTTAGTATTAAGAACGTTA
TTTATATTTCAAATTTTTCTTTTTTTTCTGTACAGACGCGTGTACGCATGTAACATTATACTGAAAACCT
TGCTTGAGAAGGTTTTGGGACGCTCGAAGATCCAGCTGCATTAATGAATCGGCCAACGCGCGGGGAGA
GGCGGTTTGCGTATTGGGCGCTCTTCCGCTTCCTCGCTCACTGACTCGCTGCGCTCGGTCGTTCGGCTG
CGGCGAGCGGTATCAGCTCACTCAAAGGCGGTAATACGGTTATCCACAGAATCAGGGGATAACGCAG
GAAAGAACATGTGAGCAAAAGGCCAGCAAAAGGCCAGGAACCGTAAAAAGGCCGCGTTGCTGGCGTT
TTTCCATAGGCTCCGCCCCCCTGACGAGCATCACAAAAATCGACGCTCAAGTCAGAGGTGGCGAAACC
CGACAGGACTATAAAGATACCAGGCGTTTCCCCCTGGAAGCTCCCTCGTGCGCTCTCCTGTTCCGACCC
TGCCGCTTACCGGATACCTGTCCGCCTTTCTCCCTTCGGGAAGCGTGGCGCTTTCTCATAGCTCACGCT
GTAGGTATCTCAGTTCGGTGTAGGTCGTTCGCTCCAAGCTGGGCTGTGTGCACGAACCCCCCGTTCAGC
CCGACCGCTGCGCCTTATCCGGTAACTATCGTCTTGAGTCCAACCCGGTAAGACACGACTTATCGCCAC
TGGCAGCAGCCACTGGTAACAGGATTAGCAGAGCGAGGTATGTAGGCGGTGCTACAGAGTTCTTGAA
GTGGTGGCCTAACTACGGCTACACTAGAAGGACAGTATTTGGTATCTGCGCTCTGCTGAAGCCAGTTA
CCTTCGGAAAAAGAGTTGGTAGCTCTTGATCCGGCAAACAAACCACCGCTGGTAGCGGTGGTTTTTTT
GTTTGCAAGCAGCAGATTACGCGCAGAAAAAAAGGATCTCAAGAAGATCCTTTGATCTTTTCTACGGG
GTCTGACGCTCAGTGGAACGAAAACTCACGTTAAGGGATTTTGGTCATGAGATTATCAAAAAGGATCT
TCACCTAGATCCTTTTAAATTAAAAATGAAGTTTTAAATCAATCTAAAGTATATATGAGTAAACTTGGT
CTGACAGTTACCAATGCTTAATCAGTGAGGCACCTATCTCAGCGATCTGTCTATTTCGTTCATCCATAG
TTGCCTGACTCCCCGTCGTGTAGATAACTACGATACGGGAGGGCTTACCATCTGGCCCCAGTGCTGCA
ATGATACCGCGAGACCCACGCTCACCGGCTCCAGATTTATCAGCAATAAACCAGCCAGCCGGAAGGG
CCGAGCGCAGAAGTGGTCCTGCAACTTTATCCGCCTCCATCCAGTCTATTAATTGTTGCCGGGAAGCTA
GAGTAAGTAGTTCGCCAGTTAATAGTTTGCGCAACGTTGTTGCCATTGCTACAGGCATCGTGGTGTCAC
GCTCGTCGTTTGGTATGGCTTCATTCAGCTCCGGTTCCCAACGATCAAGGCGAGTTACATGATCCCCCA
TGTTGTGCAAAAAAGCGGTTAGCTCCTTCGGTCCTCCGATCGTTGTCAGAAGTAAGTTGGCCGCAGTGT
TATCACTCATGGTTATGGCAGCACTGCATAATTCTCTTACTGTCATGCCATCCGTAAGATGCTTTTCTGT
GACTGGTGAGTACTCAACCAAGTCATTCTGAGAATAGTGTATGCGGCGACCGAGTTGCTCTTGCCCGG
CGTCAATACGGGATAATACCGCGCCACATAGCAGAACTTTAAAAGTGCTCATCATTGGAAAACGTTCT
TCGGGGCGAAAACTCTCAAGGATCTTACCGCTGTTGAGATCCAGTTCGATGTAACCCACTCGTGCACC
CAACTGATCTTCAGCATCTTTTACTTTCACCAGCGTTTCTGGGTGAGCAAAAACAGGAAGGCAAAATG
CCGCAAAAAAGGGAATAAGGGCGACACGGAAATGTTGAATACTCATACTCTTCCTTTTTCAATATTAT
TGAAGCATTTATCAGGGTTATTGTCTCATGAGCGGATACATATTTGAATGTATTTAGAAAAATAAACA
AATAGGGGTTCCGCGCACATTTCCCCGAAAAGTGCCACCTGAACGAAGCATCTGTGCTTCATTTTGTA
GAACAAAAATGCAACGCGAGAGCGCTAATTTTTCAAACAAAGAATCTGAGCTGCATTTTTACAGAACA
GAAATGCAACGCGAAAGCGCTATTTTACCAACGAAGAATCTGTGCTTCATTTTTGTAAAACAAAAATG
CAACGCGAGAGCGCTAATTTTTCAAACAAAGAATCTGAGCTGCATTTTTACAGAACAGAAATGCAACG
CGAGAGCGCTATTTTACCAACAAAGAATCTATACTTCTTTTTTGTTCTACAAAAATGCATCCCGAGAGC
GCTATTTTTCTAACAAAGCATCTTAGATTACTTTTTTTCTCCTTTGTGCGCTCTATAATGCAGTCTCTTG
ATAACTTTTTGCACTGTAGGTCCGTTAAGGTTAGAAGAAGGCTACTTTGGTGTCTATTTTCTCTTCCATA
AAAAAAGCCTGACTCCACTTCCCGCGTTTACTGATTACTAGCGAAGCTGCGGGTGCATTTTTTCAAGAT
AAAGGCATCCCCGATTATATTCTATACCGATGTGGATTGCGCATACTTTGTGAACAGAAAGTGATAGC
GTTGATGATTCTTCATTGGTCAGAAAATTATGAACGGTTTCTTCTATTTTGTCTCTATATACTACGTATA
GGAAATGTTTACATTTTCGTATTGTTTTCGATTCACTCTATGAATAGTTCTTACTACAATTTTTTTGTCT
AAAGAGTAATACTAGAGATAAACATAAAAAATGTAGAGGTCGAGTTTAGATGCAAGTTCAAGGAGCG
AAAGGTGGATGGGTAGGTTATATAGGGATATAGCACAGAGATATATAGCAAAGAGATACTTTTGAGC

AATGTTTGTGGAAGCGGTATTCGCAATATTTTAGTAGCTCGTTACAGTCCGGTGCGTTTTTGGTTTTTTG
AAAGTGCGTCTTCAGAGCGCTTTTGGTTTTCAAAAGCGCTCTGAAGTTCCTATACTTTCTAGAGAATAG
GAACTTCGGAATAGGAACTTCAAAGCGTTTCCGAAAACGAGCGCTTCCGAAAATGCAACGCGAGCTG
CGCACATACAGCTCACTGTTCACGTCGCACCTATATCTGCGTGTTGCCTGTATATATATATACATGAGA
AGAACGGCATAGTGCGTGTTTATGCTTAAATGCGTACTTATATGCGTCTATTTATGTAGGATGAAAGGT
AGTCTAGTACCTCCTGTGATATTATCCCATTCCATGCGGGGTATCGTATGCTTCCTTCAGCACTACCCTT
TAGCTGTTCTATATGCTGCCACTCCTCAATTGGATTAGTCTCATCCTTCAATGCTATCATTTCCTTTGAT
ATTGGATCATACTAAGAAACCATTATTATCATGACATTAACCTATAAAAATAGGCGTATCACGAGGCC
CTTTCGTC

> pESC-LEU-*GS*$^{I301E}$~*mVenus*$^{N}$

TCGCGCGTTTCGGTGATGACGGTGAAAACCTCTGACACATGCAGCTCCCGGAGACGGTCACAGCT
TGTCTGTAAGCGGATGCCGGGAGCAGACAAGCCCGTCAGGGCGCGTCAGCGGGTGTTGGCGGGT
GTCGGGGCTGGCTTAACTATGCGGCATCAGAGCAGATTGTACTGAGAGTGCACCATATCGACTAC
GTCGTAAGGCCGTTTCTGACAGAGTAAAATTCTTGAGGGAACTTTCACCATTATGGGAAATGCTT
CAAGAAGGTATTGACTTAAACTCCATCAAATGGTCAGGTCATTGAGTGTTTTTTATTTGTTGTATT
TTTTTTTTTTTAGAGAAAATCCTCCAATATCAAATTAGGAATCGTAGTTTCATGATTTTCTGTTACA
CCTAACTTTTTGTGTGGTGCCCTCCTCCTTGTCAATATTAATGTTAAAGTGCAATTCTTTTTCCTTA
TCACGTTGAGCCATTAGTATCAATTTGCTTACCTGTATTCCTTTACTATCCTCCTTTTTCTCCTTCTT
GATAAATGTATGTAGATTGCGTATATAGTTTCGTCTACCCTATGAACATATTCCATTTTGTAATTT
CGTGTCGTTTCTATTATGAATTTCATTTATAAAGTTTATGTACAAATATCATAAAAAAAGAGAATC
TTTTTAAGCAAGGATTTTCTTAACTTCTTCGGCGACAGCATCACCGACTTCGGTGGTACTGTTGGA
ACCACCTAAATCACCAGTTCTGATACCTGCATCCAAAACCTTTTTAACTGCATCTTCAATGGCCTT
ACCTTCTTCAGGCAAGTTCAATGACAATTTCAACATCATTGCAGCAGACAAGATAGTGGCGATAG
GGTCAACCTTATTCTTTGGCAAATCTGGAGCAGAACCGTGGCATGGTTCGTACAAACCAAATGCG
GTGTTCTTGTCTGGCAAAGAGGCCAAGGACGCAGATGGCAACAAACCCAAGGAACCTGGGATAA
CGGAGGCTTCATCGGAGATGATATCACCAAACATGTTGCTGGTGATTATAATACCATTTAGGTGG
GTTGGGTTCTTAACTAGGATCATGGCGGCAGAATCAATCAATTGATGTTGAACCTTCAATGTAGG
GAATTCGTTCTTGATGGTTTCCTCCACAGTTTTTCTCCATAATCTTGAAGAGGCCAAAAGATTAGC
TTTATCCAAGGACCAAATAGGCAATGGTGGCTCATGTTGTAGGGCCATGAAAGCGGCCATTCTTG
TGATTCTTTGCACTTCTGGAACGGTGTATTGTTCACTATCCCAAGCGACACCATCACCATCGTCTT
CCTTTCTCTTACCAAAGTAAATACCTCCCACTAATTCTCTGACAACAACGAAGTCAGTACCTTTAG
CAAATTGTGGCTTGATTGGAGATAAGTCTAAAAGAGAGTCGGATGCAAAGTTACATGGTCTTAAG
TTGGCGTACAATTGAAGTTCTTTACGGATTTTTAGTAAACCTTGTTCAGGTCTAACACTACCGGTA
CCCCATTTAGGACCAGCCACAGCACCTAACAAAACGGCATCAACCTTCTTGGAGGCTTCCAGCGC
CTCATCTGGAAGTGGGACACCTGTAGCATCGATAGCAGCACCACCAATTAAATGATTTTCGAAAT
CGAACTTGACATTGGAACGAACATCAGAAATAGCTTTAAGAACCTTAATGGCTTCGGCTGTGATT
TCTTGACCAACGTGGTCACCTGGCAAAACGACGATCTTCTTAGGGGCAGACATAGGGGCAGACA
TTAGAATGGTATATCCTTGAAATATATATATATATATTGCTGAAATGTAAAAGGTAAGAAAAGTTAG
AAAGTAAGACGATTGCTAACCACCTATTGGAAAAAACAATAGGTCCTTAAATAATATTGTCAACT
TCAAGTATTGTGATGCAAGCATTTAGTCATGAACGCTTCTCTATTCTATATGAAAAGCCGGTTCCG
GCCTCTCACCTTTCCTTTTTCTCCCAATTTTTCAGTTGAAAAAGGTATATGCGTCAGGCGACCTCT
GAAATTAACAAAAAATTTCCAGTCATCGAATTTGATTCTGTGCGATAGCGCCCCTGTGTGTTCTCG
TTATGTTGAGGAAAAAAATAATGGTTGCTAAGAGATTCGAACTCTTGCATCTTACGATACCTGAG
TATTCCCACAGTTAACTGCGGTCAAGATATTTCTTGAATCAGGCGCCTTAGACCGCTCGGCCAAA
CAACCAATTACTTGTTGAGAAATAGAGTATAATTATCCTATAAATATAACGTTTTTGAACACACA
TGAACAAGGAAGTACAGGACAATTGATTTTGAAGAGAATGTGGATTTTGATGTAATTGTTGGGAT
TCCATTTTTAATAAGGCAATAATATTAGGTATGTGGATATACTAGAAGTTCTCCTCGACCGTCGAT
ATGCGGTGTGAAATACCGCACAGATGCGTAAGGAGAAAATACCGCATCAGGAAATTGTAAACGT
TAATATTTTGTTAAAATTCGCGTTAAATTTTTGTTAAATCAGCTCATTTTTTAACCAATAGGCCGA
AATCGGCAAAATCCCTTATAAATCAAAAGAATAGACCGAGATAGGGTTGAGTGTTGTTCCAGTTT
GGAACAAGAGTCCACTATTAAAGAACGTGGACTCCAACGTCAAAGGGCGAAAAACCGTCTATCA
GGGCGATGGCCCACTACGTGAACCATCACCCTAATCAAGTTTTTTGGGGTCGAGGTGCCGTAAAG
CACTAAATCGGAACCCTAAAGGGAGCCCCCGATTTAGAGCTTGACGGGGAAAGCCGGCGAACGT
GGCGAGAAAGGAAGGGAAGAAAGCGAAAGGAGCGGGCGCTAGGGCGCTGGCAAGTGTAGCGGT

CACGCTGCGCGTAACCACCACACCCGCCGCGCTTAATGCGCCGCTACAGGGCGCGTCGCGCCATT
CGCCATTCAGGCTGCGCAACTGTTGGGAAGGGCGATCGGTGCGGGCCTCTTCGCTATTACGCCAG
CTGAATTGGAGCGACCTCATGCTATACCTGAGAAAGCAACCTGACCTACAGGAAAGAGTTACTC
AAGAATAAGAATTTTCGTTTTAAAACCTAAGAGTCACTTTAAAATTTGTATACACTTATTTTTTTT
ATAACTTATTTAATAATAAAAATCATAAATCATAAGAAATTCGCTTATTTAGAAGTGTCAACAAC
GTATCTACCAACGATTTGACCCTTTTCCATCTTTTCGTAAATTTCTGGCAAGGTAGACAAGCCGAC
AACCTTGATTGGAGACTTGACCAAACCTCTGGCGAAGAATTGTTAATTAAGAGCTCAGATCTTAT
CGTCGTCATCCTTGTAATCCATCGATACTAGTGCTTACTCGATGTTGTGGCGGATCTTGAAGTTGG
CCTTGATGCCGTTCTTCTGCTTGTCGGCGGTGATATAGACGTTGTGGCTGTTGTAGTTGTACTCCA
GCTTGTGCCCCAGGATGTTGCCGTCCTCCTTGAAGTCGATGCCCTTCAGCTCGATGCGGTTCACCA
GGGTGTCGCCCTCGAACTTCACCTCGGCGCGGGTCTTGTAGTTGCCGTCGTCCTTGAAGAAGATG
GTGCGCTCCTGGACGTAGCCTTCGGGCATGGCGGACTTGAAGAAGTCGTGCTGCTTCATGTGGTC
GGGGTAGCGGGCGAAGCACTGCAGGCCGTAGCCCAGGGTGGTCACGAGGGTGGGCCAGGGCAC
GGGCAGCTTGCCGGTGGTGCAGATCAGCTTCAGGGTCAGCTTGCCGTAGGTGGCATCGCCCTCGC
CCTCGCCGGACACGCTGAACTTGTGGCCGTTTACGTCGCCGTCCAGCTCGACCAGGATGGGCACC
ACCCCGGTGAACAGCTCCTCGCCCTTGCTCACGAATTCCATAGTACCACCAGAACCTTCCTCAAA
TTTCAATGTATTTCCAATGTCAATCGCAAAACGGTACTTGACATCCAAATTCTTGATACGTTCCAT
AGCAGTGCTGAGATAGTCAATCCCAATAACTTCAGTATCACATACAATGTTATGTTTGGCTGCGA
AATCAAGCATTTCTTGGTACTCTTTGAGACCTCCAGTGGAACTTCCGATTTCCTTTTTCCTTCCCAT
AATGAGAGGTGCCGCAGGGAGCTCAAATAGCGACTCCGGTGCACCTACGAGCATAACCGCGCCG
TCAAACTTGAGCAAATTGAGCATAAGTGACATAGGAGTGCGGCCACCTGGGGTGGTGTCCACAA
CACCATCCATAGTACCTGCCAGAGCCTTCAATTGCTCAGAGTCAGTGTTGACAACAAAAGCATCA
GCACCATGTTCTTCAATGGCTTCCTTCTCCTTACGCCTTGATGTACTAATAACAGTAGCCTTACCA
CCAAAAGCCTTAATAAACTTAACAGCAACAGAACCAAGACCTCCCAGCCCGAAAACCCCAATAT
GCTTTCCTGGCTTATCGAGTCCCAAATGTTTCATTGGGCTATAAACAACAACCCCAGCACAAAGG
AGAGCAACCCCTTTATCTTGAGGCAAGTTTTCGGGCCATCGGAGGACGAACTTTTCATCAACAAC
CATCACATTTGAACAACCCCCATAGGATCGTTCCCCTTGCTCACGGTAAACAGATCCATCAGCCA
TATTGGGCTCTGGGCAGTAATTCTCCATTCCACTTTGACAATTATAACATTGACCACAAGATCCGA
CCATACATCCCACAGCTACCTTGTCTCCAACCTTGAATTTCTCTACTTTGCTGCCAACTTCTACCA
CCTCACCGGCAGTTTCATGTCCAAACACATAAGGATATCTGGTGAAACCCCACTTGTTTCTGACC
ATTTCCATATCGAAATTGCAAACACCAGAGTACAAAACTCTAATCTTCACATCCCGTTCACCAGG
GACTCTTCTATAGAACTTGATGGGCTGAAGGACACCAGATGCATCTGCAGCACCCCATCCCACAG
CCTTCACTGAAAGGTCGAGTTTGGTTGTTTCTCCGGCCATGCCCTTTAGTGAGGGTTGAATTCGAA
TTTTCAAAAATTCTTACTTTTTTTTTTGGATGGACGCAAAGAAGTTTAATAATCATATTACATGGCA
TTACCACCATATACATATCCATATACATATCCATATCTAATCTTACTTATATGTTGTGGAAATGTA
AAGAGCCCCATTATCTTAGCCTAAAAAAACCTTCTCTTTGGAACTTTCAGTAATACGCTTAACTGC
TCATTGCTATATTGAAGTACGGATTAGAAGCCGCCGAGCGGGTGACAGCCCTCCGAAGGAAGAC
TCTCCTCCGTGCGTCCTCGTCTTCACCGGTCGCGTTCCTGAAACGCAGATGTGCCTCGCGCCGCAC
TGCTCCGAACAATAAAGATTCTACAATACTAGCTTTTATGGTTATGAAGAGGAAAAATTGGCAGT
AACCTGGCCCCACAAACCTTCAAATGAACGAATCAAATTAACAACCATAGGATGATAATGCGAT
TAGTTTTTTAGCCTTATTTCTGGGGTAATTAATCAGCGAAGCGATGATTTTTGATCTATTAACAGA
TATATAAATGCAAAAACTGCATAACCACTTTAACTAATACTTTCAACATTTTCGGTTTGTATTACT
TCTTATTCAAATGTAATAAAAGTATCAACAAAAAATTGTTAATATACCTCTATACTTTAACGTCAA
GGAGAAAAAACCCCGGATCCGTAATACGACTCACTATAGGGCCCGGGCGTCGACATGGAACAGA
AGTTGATTTCCGAAGAAGACCTCGAGTAAGCTTGGTACCGCGGCTAGCTAAGATCCGCTCTAACC
GAAAAGGAAGGAGTTAGACAACCTGAAGTCTAGGTCCCTATTTATTTTTTTATAGTTATGTTAGT
ATTAAGAACGTTATTTATATTTCAAATTTTTCTTTTTTTTCTGTACAGACGCGTGTACGCATGTAAC
ATTATACTGAAAACCTTGCTTGAGAAGGTTTTGGGACGCTCGAAGATCCAGCTGCATTAATGAAT
CGGCCAACGCGCGGGGAGAGGCGGTTTGCGTATTGGGCGCTCTTCCGCTTCCTCGCTCACTGACT
CGCTGCGCTCGGTCGTTCGGCTGCGGCGAGCGGTATCAGCTCACTCAAAGGCGGTAATACGGTTA
TCCACAGAATCAGGGGATAACGCAGGAAAGAACATGTGAGCAAAAGGCCAGCAAAAGGCCAGG
AACCGTAAAAAGGCCGCGTTGCTGGCGTTTTTCCATAGGCTCCGCCCCCCTGACGAGCATCACAA
AAATCGACGCTCAAGTCAGAGGTGGCGAAACCCGACAGGACTATAAAGATACCAGGCGTTTCCC
CCTGGAAGCTCCCTCGTGCGCTCTCCTGTTCCGACCCTGCCGCTTACCGGATACCTGTCCGCCTTT
CTCCCTTCGGGAAGCGTGGCGCTTTCTCATAGCTCACGCTGTAGGTATCTCAGTTCGGTGTAGGTC
GTTCGCTCCAAGCTGGGCTGTGTGCACGAACCCCCCGTTCAGCCCGACCGCTGCGCCTTATCCGG

TAACTATCGTCTTGAGTCCAACCCGGTAAGACACGACTTATCGCCACTGGCAGCAGCCACTGGTA
ACAGGATTAGCAGAGCGAGGTATGTAGGCGGTGCTACAGAGTTCTTGAAGTGGTGGCCTAACTA
CGGCTACACTAGAAGGACAGTATTTGGTATCTGCGCTCTGCTGAAGCCAGTTACCTTCGGAAAAA
GAGTTGGTAGCTCTTGATCCGGCAAACAAACCACCGCTGGTAGCGGTGGTTTTTTTGTTTGCAAG
CAGCAGATTACGCGCAGAAAAAAAGGATCTCAAGAAGATCCTTTGATCTTTTCTACGGGGTCTGA
CGCTCAGTGGAACGAAAACTCACGTTAAGGGATTTTGGTCATGAGATTATCAAAAAGGATCTTCA
CCTAGATCCTTTTAAATTAAAAATGAAGTTTTAAATCAATCTAAAGTATATATGAGTAAACTTGG
TCTGACAGTTACCAATGCTTAATCAGTGAGGCACCTATCTCAGCGATCTGTCTATTTCGTTCATCC
ATAGTTGCCTGACTCCCCGTCGTGTAGATAACTACGATACGGGAGGGCTTACCATCTGGCCCCAG
TGCTGCAATGATACCGCGAGACCCACGCTCACCGGCTCCAGATTTATCAGCAATAAACCAGCCAG
CCGGAAGGGCCGAGCGCAGAAGTGGTCCTGCAACTTTATCCGCCTCCATCCAGTCTATTAATTGT
TGCCGGGAAGCTAGAGTAAGTAGTTCGCCAGTTAATAGTTTGCGCAACGTTGTTGCCATTGCTAC
AGGCATCGTGGTGTCACGCTCGTCGTTTGGTATGGCTTCATTCAGCTCCGGTTCCCAACGATCAAG
GCGAGTTACATGATCCCCCATGTTGTGCAAAAAAGCGGTTAGCTCCTTCGGTCCTCCGATCGTTGT
CAGAAGTAAGTTGGCCGCAGTGTTATCACTCATGGTTATGGCAGCACTGCATAATTCTCTTACTGT
CATGCCATCCGTAAGATGCTTTTCTGTGACTGGTGAGTACTCAACCAAGTCATTCTGAGAATAGT
GTATGCGGCGACCGAGTTGCTCTTGCCCGGCGTCAATACGGGATAATACCGCGCCACATAGCAGA
ACTTTAAAAGTGCTCATCATTGGAAAACGTTCTTCGGGGCGAAAACTCTCAAGGATCTTACCGCT
GTTGAGATCCAGTTCGATGTAACCCACTCGTGCACCCAACTGATCTTCAGCATCTTTTACTTTCAC
CAGCGTTTCTGGGTGAGCAAAAACAGGAAGGCAAAATGCCGCAAAAAAGGGAATAAGGGCGAC
ACGGAAATGTTGAATACTCATACTCTTCCTTTTTCAATATTATTGAAGCATTTATCAGGGTTATTG
TCTCATGAGCGGATACATATTTGAATGTATTTAGAAAAATAAACAAATAGGGGTTCCGCGCACAT
TTCCCCGAAAAGTGCCACCTGAACGAAGCATCTGTGCTTCATTTTGTAGAACAAAAATGCAACGC
GAGAGCGCTAATTTTTCAAACAAAGAATCTGAGCTGCATTTTTACAGAACAGAAATGCAACGCG
AAAGCGCTATTTTACCAACGAAGAATCTGTGCTTCATTTTTGTAAAACAAAAATGCAACGCGAGA
GCGCTAATTTTTCAAACAAAGAATCTGAGCTGCATTTTTACAGAACAGAAATGCAACGCGAGAGC
GCTATTTTACCAACAAAGAATCTATACTTCTTTTTTGTTCTACAAAAATGCATCCCGAGAGCGCTA
TTTTTCTAACAAAGCATCTTAGATTACTTTTTTTCTCCTTTGTGCGCTCTATAATGCAGTCTCTTGA
TAACTTTTTGCACTGTAGGTCCGTTAAGGTTAGAAGAAGGCTACTTTGGTGTCTATTTTCTCTTCC
ATAAAAAAAGCCTGACTCCACTTCCCGCGTTTACTGATTACTAGCGAAGCTGCGGGTGCATTTTTT
CAAGATAAAGGCATCCCCGATTATATTCTATACCGATGTGGATTGCGCATACTTTGTGAACAGAA
AGTGATAGCGTTGATGATTCTTCATTGGTCAGAAAATTATGAACGGTTTCTTCTATTTTGTCTCTA
TATACTACGTATAGGAAATGTTTACATTTTCGTATTGTTTTCGATTCACTCTATGAATAGTTCTTAC
TACAATTTTTTTGTCTAAAGAGTAATACTAGAGATAAACATAAAAAATGTAGAGGTCGAGTTTAG
ATGCAAGTTCAAGGAGCGAAAGGTGGATGGGTAGGTTATATAGGGATATAGCACAGAGATATAT
AGCAAAGAGATACTTTTGAGCAATGTTTGTGGAAGCGGTATTCGCAATATTTTAGTAGCTCGTTA
CAGTCCGGTGCGTTTTTGGTTTTTTGAAAGTGCGTCTTCAGAGCGCTTTTGGTTTTCAAAAGCGCT
CTGAAGTTCCTATACTTTCTAGAGAATAGGAACTTCGGAATAGGAACTTCAAAGCGTTTCCGAAA
ACGAGCGCTTCCGAAAATGCAACGCGAGCTGCGCACATACAGCTCACTGTTCACGTCGCACCTAT
ATCTGCGTGTTGCCTGTATATATATATACATGAGAAGAACGGCATAGTGCGTGTTTATGCTTAAA
TGCGTACTTATATGCGTCTATTTATGTAGGATGAAAGGTAGTCTAGTACCTCCTGTGATATTATCC
CATTCCATGCGGGGTATCGTATGCTTCCTTCAGCACTACCCTTTAGCTGTTCTATATGCTGCCACT
CCTCAATTGGATTAGTCTCATCCTTCAATGCTATCATTTCCTTTGATATTGGATCATACTAAGAAA
CCATTATTATCATGACATTAACCTATAAAAATAGGCGTATCACGAGGCCCTTTCGTC

> pESC-LEU-*mVenus*$^{N}$*~SGD*

TCGCGCGTTTCGGTGATGACGGTGAAAACCTCTGACACATGCAGCTCCCGGAGACGGTCACAGCT
TGTCTGTAAGCGGATGCCGGGAGCAGACAAGCCCGTCAGGGCGCGTCAGCGGGTGTTGGCGGGT
GTCGGGGCTGGCTTAACTATGCGGCATCAGAGCAGATTGTACTGAGAGTGCACCATATCGACTAC
GTCGTAAGGCCGTTTCTGACAGAGTAAAATTCTTGAGGGAACTTTCACCATTATGGGAAATGCTT
CAAGAAGGTATTGACTTAAACTCCATCAAATGGTCAGGTCATTGAGTGTTTTTTATTTGTTGTATT
TTTTTTTTTTTAGAGAAAATCCTCCAATATCAAATTAGGAATCGTAGTTTCATGATTTTCTGTTACA
CCTAACTTTTTGTGTGGTGCCCTCCTCCTTGTCAATATTAATGTTAAAGTGCAATTCTTTTTCCTTA
TCACGTTGAGCCATTAGTATCAATTTGCTTACCTGTATTCCTTTACTATCCTCCTTTTTCTCCTTCTT
GATAAATGTATGTAGATTGCGTATATAGTTTCGTCTACCCTATGAACATATTCCATTTTGTAATTT

CGTGTCGTTTCTATTATGAATTTCATTTATAAAGTTTATGTACAAATATCATAAAAAAAGAGAATC
TTTTTAAGCAAGGATTTTCTTAACTTCTTCGGCGACAGCATCACCGACTTCGGTGGTACTGTTGGA
ACCACCTAAATCACCAGTTCTGATACCTGCATCCAAAACCTTTTTAACTGCATCTTCAATGGCCTT
ACCTTCTTCAGGCAAGTTCAATGACAATTTCAACATCATTGCAGCAGACAAGATAGTGGCGATAG
GGTCAACCTTATTCTTTGGCAAATCTGGAGCAGAACCGTGGCATGGTTCGTACAAACCAAATGCG
GTGTTCTTGTCTGGCAAAGAGGCCAAGGACGCAGATGGCAACAAACCCAAGGAACCTGGGATAA
CGGAGGCTTCATCGGAGATGATATCACCAAACATGTTGCTGGTGATTATAATACCATTTAGGTGG
GTTGGGTTCTTAACTAGGATCATGGCGGCAGAATCAATCAATTGATGTTGAACCTTCAATGTAGG
GAATTCGTTCTTGATGGTTTCCTCCACAGTTTTTCTCCATAATCTTGAAGAGGCCAAAAGATTAGC
TTTATCCAAGGACCAAATAGGCAATGGTGGCTCATGTTGTAGGGCCATGAAAGCGGCCATTCTTG
TGATTCTTTGCACTTCTGGAACGGTGTATTGTTCACTATCCCAAGCGACACCATCACCATCGTCTT
CCTTTCTCTTACCAAAGTAAATACCTCCCACTAATTCTCTGACAACAACGAAGTCAGTACCTTTAG
CAAATTGTGGCTTGATTGGAGATAAGTCTAAAAGAGAGTCGGATGCAAAGTTACATGGTCTTAAG
TTGGCGTACAATTGAAGTTCTTTACGGATTTTTAGTAAACCTTGTTCAGGTCTAACACTACCGGTA
CCCCATTTAGGACCAGCCACAGCACCTAACAAAACGGCATCAACCTTCTTGGAGGCTTCCAGCGC
CTCATCTGGAAGTGGGACACCTGTAGCATCGATAGCAGCACCACCAATTAAATGATTTTCGAAAT
CGAACTTGACATTGGAACGAACATCAGAAATAGCTTTAAGAACCTTAATGGCTTCGGCTGTGATT
TCTTGACCAACGTGGTCACCTGGCAAAACGACGATCTTCTTAGGGGCAGACATAGGGGCAGACA
TTAGAATGGTATATCCTTGAAATATATATATATATTGCTGAAATGTAAAAGGTAAGAAAAGTTAG
AAAGTAAGACGATTGCTAACCACCTATTGGAAAAAACAATAGGTCCTTAAATAATATTGTCAACT
TCAAGTATTGTGATGCAAGCATTTAGTCATGAACGCTTCTCTATTCTATATGAAAAGCCGGTTCCG
GCCTCTCACCTTTCCTTTTTCTCCCAATTTTTCAGTTGAAAAAGGTATATGCGTCAGGCGACCTCT
GAAATTAACAAAAAAATTTCCAGTCATCGAATTTGATTCTGTGCGATAGCGCCCCTGTGTGTTCTCG
TTATGTTGAGGAAAAAAATAATGGTTGCTAAGAGATTCGAACTCTTGCATCTTACGATACCTGAG
TATTCCCACAGTTAACTGCGGTCAAGATATTTCTTGAATCAGGCGCCTTAGACCGCTCGGCCAAA
CAACCAATTACTTGTTGAGAAATAGAGTATAATTATCCTATAAATATAACGTTTTTGAACACACA
TGAACAAGGAAGTACAGGACAATTGATTTTGAAGAGAATGTGGATTTTGATGTAATTGTTGGGAT
TCCATTTTTAATAAGGCAATAATATTAGGTATGTGGATATACTAGAAGTTCTCCTCGACCGTCGAT
ATGCGGTGTGAAATACCGCACAGATGCGTAAGGAGAAAATACCGCATCAGGAAATTGTAAACGT
TAATATTTTGTTAAAATTCGCGTTAAATTTTTGTTAAATCAGCTCATTTTTTAACCAATAGGCCGA
AATCGGCAAAATCCCTTATAAATCAAAAGAATAGACCGAGATAGGGTTGAGTGTTGTTCCAGTTT
GGAACAAGAGTCCACTATTAAAGAACGTGGACTCCAACGTCAAAGGGCGAAAAACCGTCTATCA
GGGCGATGGCCCACTACGTGAACCATCACCCTAATCAAGTTTTTTGGGGTCGAGGTGCCGTAAAG
CACTAAATCGGAACCCTAAAGGGAGCCCCCGATTTAGAGCTTGACGGGGAAAGCCGGCGAACGT
GGCGAGAAAGGAAGGGAAGAAAGCGAAAGGAGCGGGCGCTAGGGCGCTGGCAAGTGTAGCGGT
CACGCTGCGCGTAACCACCACACCCGCCGCGCTTAATGCGCCGCTACAGGGCGCGTCGCGCCATT
CGCCATTCAGGCTGCGCAACTGTTGGGAAGGGCGATCGGTGCGGGCCTCTTCGCTATTACGCCAG
CTGAATTGGAGCGACCTCATGCTATACCTGAGAAAGCAACCTGACCTACAGGAAAGAGTTACTC
AAGAATAAGAATTTTCGTTTTAAAACCTAAGAGTCACTTTAAAATTTGTATACACTTATTTTTTTT
ATAACTTATTTAATAATAAAAATCATAAATCATAAGAAATTCGCTTATTTAGAAGTGTCAACAAC
GTATCTACCAACGATTTGACCCTTTTCCATCTTTTCGTAAATTTCTGGCAAGGTAGACAAGCCGAC
AACCTTGATTGGAGACTTGACCAAACCTCTGGCGAAGAATTGTTAATTAAGAGCTCAGATCTTAT
CGTCGTCATCCTTGTAATCCATCGATACTAGTGCTTAATATTTTTGTTTTTTAACCAATTCAACCAA
TTTATCTTCTTCTCTAAATCTTTTTTTAGCAGTATTAGTAACAAAACCTTCAGAAATAAAATTTTTA
TACCAAATAGCAGAATCTTTTGGATATCTTTGAAAAGTTTTATAATCAACATGAATAATACCATA
TCTACAAATATAACCCAAATTCCATTCAAAATTATCAAAAAAAGACCAAACAAAAAAACCTTTA
ACATTAACACCATCATCAATAGCATCTCTAACAGAAGCCAAATGAGATTGCAAAAAATCAACTCT
CAATTTATCATGTCTAGCTTCAGTCAACAAAATATTAGTTTTACCTTCAGTCAACAAAATATTAGT
TCTATTTTCTTCAACAACACCACATTCAGAAACATAAATAACTGGAACATGATATTTTTCTTTAGT
ATAAACCAACAAATTATACAAACCAGATGGAACAACATGTTGCCAACCACCATAACATGGTTCA
CCAATTCTAACTTCTTTACCATCAACTTTTTTAACAAAAATATTTTTATTAATTCTAGCATCAGTTT
CATAACCTGGAGTATCTGGAATTTTATCAGCATTAGAAACATAAGTAGTAGTATAATAATTCATA
CCAATAAAATCATAACAACCAGTCAATTTTTCAGAATCTTCAGTAGAAAATTCTGGCAATCTAGA
ACCAACCAAAGCTCTCATAGATTTTGGATATTCACCAGTAGTCAATGGTTCAATAAACCAACCCA
ACATAAAATCTGGACCTCTTTCTCTAGCATCAATATCTTCTTTAGTTTCATTCAATGGTTCCATCCA
CATAGAATTCAAAACAATACCAATTTCACCACCTTGACATTTTTGAAAATTTTTTCTATAAACTTC

AACAGCAGCTTTATGAGACAACAACAAATTGTGTGTAGCAATATATGGTTCTTTACCTGGATTAC
CCTTACCATCAGCACCACCTCTACCTGGAGCAAATTCACCTGTAGCATAACCAGAAGCAACATAA
GTATGTGGTTCGTTAAAAGTAGTCCAGAATTTAACCTTATCACCAAATTCCCAAAAACAAAATTC
AGCGTATTCAGTGAAATCTTCAACAATTCTATCAGACAAAAAACCACCATATTCATCTTCTAGAG
CTTGAGGCAAATCCCAATGAAACAAAGTAGCAAAAGGCTTGATACCATTAGCCAACAATTCATCT
ATAAAATCGTGGTAAAACTTAACACCATCTTTATTAACACCACCAGACAAATTACCACCTGGCAA
AACTCTAGACCATGAGATAGAAAATCTATAAGATTCCAAACCTGTTTGCTTCATGATTTTGATATC
TTCCTTGTACAAGTTGTAAGAATTAATAGCTTGATTACCATTTGAACCATCAGCAATCTTAGCTGG
GTATCTATTAGTAAAGGTATCCCATATTGATGGACCTCTGTTACCTTCGTTATAAGCACCTTCACA
TTGATAAGCAGAACCACCGGCACCCAAAATAAAATCAGATGGAAAATCTCTTCTGTGAACAATT
GGCTTGTTATGTTTTCTTGGTTGAATTGGAATAGATGGATAAGCAAATGGAATTGGAACGGAGTG
ATTACCATTTGGTTCAGCTGCTGGGGATATAGCAACAACCAAAGATTGATCATCTTTGGAACCCA
TAGTACCACCAGAACCCTCGATGTTGTGGCGGATCTTGAAGTTGGCCTTGATGCCGTTCTTCTGCT
TGTCGGCGGTGATATAGACGTTGTGGCTGTTGTAGTTGTACTCCAGCTTGTGCCCCAGGATGTTGC
CGTCCTCCTTGAAGTCGATGCCCTTCAGCTCGATGCGGTTCACCAGGGTGTCGCCCTCGAACTTCA
CCTCGGCGCGGGTCTTGTAGTTGCCGTCGTCCTTGAAGAAGATGGTGCGCTCCTGGACGTAGCCT
TCGGGCATGGCGGACTTGAAGAAGTCGTGCTGCTTCATGTGGTCGGGGTAGCGGGCGAAGCACT
GCAGGCCGTAGCCCAGGGTGGTCACGAGGGTGGGCCAGGGCACGGGCAGCTTGCCGGTGGTGCA
GATCAGCTTCAGGGTCAGCTTGCCGTAGGTGGCATCGCCCTCGCCCTCGCCGGACACGCTGAACT
TGTGGCCGTTTACGTCGCCGTCCAGCTCGACCAGGATGGGCACCACCCCGGTGAACAGCTCCTCG
CCCTTGCTCACGAATTCCATGCCCTTTAGTGAGGGTTGAATTCGAATTTTCAAAAATTCTTACTTT
TTTTTTGGATGGACGCAAAGAAGTTTAATAATCATATTACATGGCATTACCACCATATACATATCC
ATATACATATCCATATCTAATCTTACTTATATGTTGTGGAAATGTAAAGAGCCCCATTATCTTAGC
CTAAAAAAACCTTCTCTTTGGAACTTTCAGTAATACGCTTAACTGCTCATTGCTATATTGAAGTAC
GGATTAGAAGCCGCCGAGCGGGTGACAGCCCTCCGAAGGAAGACTCTCCTCCGTGCGTCCTCGTC
TTCACCGGTCGCGTTCCTGAAACGCAGATGTGCCTCGCGCCGCACTGCTCCGAACAATAAAGATT
CTACAATACTAGCTTTTATGGTTATGAAGAGGAAAAATTGGCAGTAACCTGGCCCCACAAACCTT
CAAATGAACGAATCAAATTAACAACCATAGGATGATAATGCGATTAGTTTTTTAGCCTTATTTCT
GGGGTAATTAATCAGCGAAGCGATGATTTTTGATCTATTAACAGATATATAAATGCAAAAACTGC
ATAACCACTTTAACTAATACTTTCAACATTTTCGGTTTGTATTACTTCTTATTCAAATGTAATAAA
AGTATCAACAAAAAATTGTTAATATACCTCTATACTTTAACGTCAAGGAGAAAAAACCCCGGATC
CGTAATACGACTCACTATAGGGCCCGGGCGTCGACATGGAACAGAAGTTGATTTCCGAAGAAGA
CCTCGAGTAAGCTTGGTACCGCGGCTAGCTAAGATCCGCTCTAACCGAAAAGGAAGGAGTTAGA
CAACCTGAAGTCTAGGTCCCTATTTATTTTTTTATAGTTATGTTAGTATTAAGAACGTTATTTATAT
TTCAAATTTTTCTTTTTTTTCTGTACAGACGCGTGTACGCATGTAACATTATACTGAAAACCTTGCT
TGAGAAGGTTTTGGGACGCTCGAAGATCCAGCTGCATTAATGAATCGGCCAACGCGCGGGGAGA
GGCGGTTTGCGTATTGGGCGCTCTTCCGCTTCCTCGCTCACTGACTCGCTGCGCTCGGTCGTTCGG
CTGCGGCGAGCGGTATCAGCTCACTCAAAGGCGGTAATACGGTTATCCACAGAATCAGGGGATA
ACGCAGGAAAGAACATGTGAGCAAAAGGCCAGCAAAAGGCCAGGAACCGTAAAAAGGCCGCGT
TGCTGGCGTTTTTCCATAGGCTCCGCCCCCCTGACGAGCATCACAAAAATCGACGCTCAAGTCAG
AGGTGGCGAAACCCGACAGGACTATAAAGATACCAGGCGTTTCCCCCTGGAAGCTCCCTCGTGC
GCTCTCCTGTTCCGACCCTGCCGCTTACCGGATACCTGTCCGCCTTTCTCCCTTCGGGAAGCGTGG
CGCTTTCTCATAGCTCACGCTGTAGGTATCTCAGTTCGGTGTAGGTCGTTCGCTCCAAGCTGGGCT
GTGTGCACGAACCCCCCGTTCAGCCCGACCGCTGCGCCTTATCCGGTAACTATCGTCTTGAGTCC
AACCCGGTAAGACACGACTTATCGCCACTGGCAGCAGCCACTGGTAACAGGATTAGCAGAGCGA
GGTATGTAGGCGGTGCTACAGAGTTCTTGAAGTGGTGGCCTAACTACGGCTACACTAGAAGGAC
AGTATTTGGTATCTGCGCTCTGCTGAAGCCAGTTACCTTCGGAAAAAGAGTTGGTAGCTCTTGAT
CCGGCAAACAAACCACCGCTGGTAGCGGTGGTTTTTTTGTTTGCAAGCAGCAGATTACGCGCAGA
AAAAAAGGATCTCAAGAAGATCCTTTGATCTTTTCTACGGGGTCTGACGCTCAGTGGAACGAAAA
CTCACGTTAAGGGATTTTGGTCATGAGATTATCAAAAAGGATCTTCACCTAGATCCTTTTAAATTA
AAAATGAAGTTTTAAATCAATCTAAAGTATATATGAGTAAACTTGGTCTGACAGTTACCAATGCT
TAATCAGTGAGGCACCTATCTCAGCGATCTGTCTATTTCGTTCATCCATAGTTGCCTGACTCCCCG
TCGTGTAGATAACTACGATACGGGAGGGCTTACCATCTGGCCCCAGTGCTGCAATGATACCGCGA
GACCCACGCTCACCGGCTCCAGATTTATCAGCAATAAACCAGCCAGCCGGAAGGGCCGAGCGCA
GAAGTGGTCCTGCAACTTTATCCGCCTCCATCCAGTCTATTAATTGTTGCCGGGAAGCTAGAGTA
AGTAGTTCGCCAGTTAATAGTTTGCGCAACGTTGTTGCCATTGCTACAGGCATCGTGGTGTCACG

CTCGTCGTTTGGTATGGCTTCATTCAGCTCCGGTTCCCAACGATCAAGGCGAGTTACATGATCCCC
CATGTTGTGCAAAAAAGCGGTTAGCTCCTTCGGTCCTCCGATCGTTGTCAGAAGTAAGTTGGCCG
CAGTGTTATCACTCATGGTTATGGCAGCACTGCATAATTCTCTTACTGTCATGCCATCCGTAAGAT
GCTTTTCTGTGACTGGTGAGTACTCAACCAAGTCATTCTGAGAATAGTGTATGCGGCGACCGAGT
TGCTCTTGCCCGGCGTCAATACGGGATAATACCGCGCCACATAGCAGAACTTTAAAAGTGCTCAT
CATTGGAAAACGTTCTTCGGGGCGAAAACTCTCAAGGATCTTACCGCTGTTGAGATCCAGTTCGA
TGTAACCCACTCGTGCACCCAACTGATCTTCAGCATCTTTTACTTTCACCAGCGTTTCTGGGTGAG
CAAAAACAGGAAGGCAAAATGCCGCAAAAAAGGGAATAAGGGCGACACGGAAATGTTGAATAC
TCATACTCTTCCTTTTTCAATATTATTGAAGCATTTATCAGGGTTATTGTCTCATGAGCGGATACA
TATTTGAATGTATTTAGAAAAATAAACAAATAGGGGTTCCGCGCACATTTCCCCGAAAAGTGCCA
CCTGAACGAAGCATCTGTGCTTCATTTTGTAGAACAAAAATGCAACGCGAGAGCGCTAATTTTTC
AAACAAAGAATCTGAGCTGCATTTTTACAGAACAGAAATGCAACGCGAAAGCGCTATTTTACCA
ACGAAGAATCTGTGCTTCATTTTTGTAAAACAAAAATGCAACGCGAGAGCGCTAATTTTTCAAAC
AAAGAATCTGAGCTGCATTTTTACAGAACAGAAATGCAACGCGAGAGCGCTATTTTACCAACAA
AGAATCTATACTTCTTTTTTGTTCTACAAAAATGCATCCCGAGAGCGCTATTTTTCTAACAAAGCA
TCTTAGATTACTTTTTTTCTCCTTTGTGCGCTCTATAATGCAGTCTCTTGATAACTTTTTGCACTGT
AGGTCCGTTAAGGTTAGAAGAAGGCTACTTTGGTGTCTATTTTCTCTTCCATAAAAAAAGCCTGA
CTCCACTTCCCGCGTTTACTGATTACTAGCGAAGCTGCGGGTGCATTTTTTCAAGATAAAGGCATC
CCCGATTATATTCTATACCGATGTGGATTGCGCATACTTTGTGAACAGAAAGTGATAGCGTTGAT
GATTCTTCATTGGTCAGAAAATTATGAACGGTTTCTTCTATTTTGTCTCTATATACTACGTATAGG
AAATGTTTACATTTTCGTATTGTTTTCGATTCACTCTATGAATAGTTCTTACTACAATTTTTTTGTC
TAAAGAGTAATACTAGAGATAAACATAAAAAAATGTAGAGGTCGAGTTTAGATGCAAGTTCAAGG
AGCGAAAGGTGGATGGGTAGGTTATATAGGGATATAGCACAGAGATATATAGCAAAGAGATACT
TTTGAGCAATGTTTGTGGAAGCGGTATTCGCAATATTTTAGTAGCTCGTTACAGTCCGGTGCGTTT
TTGGTTTTTTGAAAGTGCGTCTTCAGAGCGCTTTTGGTTTTCAAAAGCGCTCTGAAGTTCCTATAC
TTTCTAGAGAATAGGAACTTCGGAATAGGAACTTCAAAGCGTTTCCGAAAACGAGCGCTTCCGAA
AATGCAACGCGAGCTGCGCACATACAGCTCACTGTTCACGTCGCACCTATATCTGCGTGTTGCCT
GTATATATATATACATGAGAAGAACGGCATAGTGCGTGTTTATGCTTAAATGCGTACTTATATGC
GTCTATTTATGTAGGATGAAAGGTAGTCTAGTACCTCCTGTGATATTATCCCATTCCATGCGGGGT
ATCGTATGCTTCCTTCAGCACTACCCTTTAGCTGTTCTATATGCTGCCACTCCTCAATTGGATTAGT
CTCATCCTTCAATGCTATCATTTCCTTTGATATTGGATCATACTAAGAAACCATTATTATCATGAC
ATTAACCTATAAAAATAGGCGTATCACGAGGCCCTTTCGTC

**Table S7.** List of primers used in this study.

| Primer Name | Sequences 5'-3' |
|---|---|
| CrCAD1/2-BamHI-F | ataggatccaatggccggaaaatcaccaga |
| CrCAD1/2-SalI-R | gttgtcgacttaaggagctttcaaggt |
| GsVinBLAST-BamHI-F | acaaggccatggcgatatcggatccaatggctggaaaatcaccagaaga |
| GsVinBLAST-SalI-R | cgagtgcggccgcaagcttgtcgacttaaggagctttcaaggtgttgcc |
| SspVinBLAST-BamHI-F | acaaggccatggcgatatcggatccaatggccggaaaatcgccaga |
| SspVinBLAST-SalI-R | cgagtgcggccgcaagcttgtcgacttaaggagctttcaatgtgttgcca |
| RsVinBLAST-BamHI-F | ataggatccgatggccggaaaatcacctg |
| RsVinBLAST-SalI-R | atagtcgacttaaggagctttcaaggtgttgg |
| attb-MsVinBLAST-F | ggggacaagtttgtacaaaaaagcaggcttcatgactggaaaatcaccagaag |
| attb-Ms1VinBLAST-R | ggggaccactttgtacaagaaagctgggtcaaggagctttcaaggtgttgg |
| attb-NbVinBLAST-F | ggggacaagtttgtacaaaaaagcaggcttcatggagaaatcactagaggaagtg |
| attb-Nb1VinBLAST-R | ggggaccactttgtacaagaaagctgggtcattctgctttgagggtcttag |
| CrGS-BamHI-F | tacgggatccatggctggagaaacaacaaaactagac |
| CrGS-SalI-R | aggcgtcgactcattcctcaaatttcaatgtatttcc |
| GsGS-BamHI-F | ataggatccaatggctgccaaatcaccagaa |
| GsGS-SacI-R | atagagctctcaatcatcctttaatgtatttgca |
| SspGS-BamHI-F | acaaggccatggcgatatcggatccaatggttggaaaaccagctggaga |
| SspGS-SalI-R | cgagtgcggccgcaagcttgtcgactacatgacactttcaagaacggt |
| pET30b+NdeI-F | ctttaagaaggagatatacatatgacaaggccatggcgatatcggatc |
| pET30b+R | cgagtgcggccgcaagcttgtcgac |
| TMV_fwd | ttcatttggagaggacacgcctcgagtataagagctctatttttac |
| TMV_rev-mVenus-N | tgctcaccatggtaattgtaaatgtaattgtaatg |
| TMV_rev-mVenus-C | cgctgcccatggtaattgtaaatgtaattgtaatg |
| mVenus-N_fwd | tacaattaccatggtgagcaagggcgag |
| mVenus-N_rev | tttctttcatgtcctcgatgttgtggcg |
| mVenus-C_fwd | tacaattaccatgggcagcgtgcagctc |
| mVenus-C_rev | gctgtacaagatgaaagaaaccgctgctg |
| pET30_fwd | catcgaggacatgaaagaaaccgctgctg |
| pET30_rev | ttaaagcaggactctagggactagtctcgagtgcggccgcaag |
| mVenus-NbVinBLAST_fwd | acaaggccatggcgatatcggatcccatggagaaatcactagaggaa |
| mVenus-NbVinBLAST_rev | ttaaagcaggactctagggactagtctcgagtgcggccgcaatcattctgctttgagggtcttag |
| VIGS-CrVinBLAST-F | cgcgaattcatgagatggtatgcaatgaacac |
| VIGS-CrVinBLAST-R | cacgaattccttctggattactgctcaaca |
| VIGS-CrCAD3-F | ttgtgaattcgttaccacgtacggtgcattgaga |
| VIGS-CrCAD3-R | tgagatggtgccccaatcaaca |
| VIGS-CrCAD4-F | gatgaattcatgcaggcaaggaaaaggagccta |
| VIGS-CrCAD4-R | ggagaattcttggcaagccgatcaagagcttcat |
| VIGS-CrCAD5-F | gtagaattctgttggtctaggtggacttggtca |
| VIGS-CrCAD5-R | ctaagaattccctgtagctccaaccaaaacaag |
| qPCR-CrCAD1-F | tgtgtggttgggggagtagcag |
| qPCR-CrCAD1-R | gtctccttcaatcctccaaggtcat |
| qPCR-CrCAD2-F | aagttggtgttggctgcttggtt |
| qPCR-CrCAD2-R | tggacttgcataggttagcaccat |
| qPCR-CrGS-F | tccctgcggcacctctcatta |
| qPCR-CrGS-R | acatacaatgttatgtttggctgcgaa |
| GAL-1-10-F | tttcaaaaattcttactttttttttggat |
| GAL-1-10-R | gttttttctccttgacgtt |
| M13R-48-R | agcggataacaatttcacacagga |
| ADH1-F | acaggaaagagttactcaagaata |

| Primer Name | Sequences 5'-3' |
|---|---|
| CYC1-R | actccttccttttcggttagag |
| IntG3-ADH1-F | ctctgtagatgccactatttgtcgctctccatgtagtaaggagcgacctcatgctatac |
| IntG3-CYC1-R | agcagatcaattgttccagcacctataactacagcttactcttcgagcgtcccaaaacc |
| IntG4-ADH1-F | tagatgactcagtttagtctgaccttctatagtatactacgagcgacctcatgctatac |
| IntG4-CYC1-R | tttcgccggtcccggaattttcgtttccgcaataaaagaacttcgagcgtcccaaaacc |
| IntG11-ADH1-F | agactattcccataatgtttacgtttcacagaaattgagggagcgacctcatgctatac |
| IntG11-CYC1-R | tattcgtatagcaaggtactagtggcgacactcaggtggtcttcgagcgtcccaaaacc |
| IntG13-ADH1-F | aaattttgcaaattagtgcgcggggtgaatgggtgcgtcagagcgacctcatgctatac |
| IntG13-CYC1-R | tgctatggcctgccttggacacgatacgtcgctgcttatgcttcgagcgtcccaaaacc |
| IntG16-ADH1-F | ttaagagggaataagaaaactaagggaaaatacgctatctgagcgacctcatgctatac |
| IntG16-CYC1-R | ttacgtactcgcatgtattcgaaaagcctctaaaaattgccttcgagcgtcccaaaacc |
| IntG17-ADH1-F | cgaaccagaattttcttattttttcactaaggataattggagcgacctcatgctatac |
| IntG17-CYC1-R | caatcttctattgtattggtggtatttaaggtgaactaatcttcgagcgtcccaaaacc |
| IntG21-ADH1-F | ctatagaggaaaaggttgtatattttagagattcgtccatgagcgacctcatgctatac |
| IntG21-CYC1-R | agggcaacgctttgaactagtattttccgttatagtatgtcttcgagcgtcccaaaacc |
| IntG22-ADH1-F | tttctatcattgatgacgggcattaccccgttaatgacctgagcgacctcatgctatac |
| IntG22-CYC1-R | tatttctcacttattgcacctctcgaagttctctcattagcttcgagcgtcccaaaacc |
| IntG26-ADH1-F | ataagagtggaaaaaaagtaacagattagtggctcccagtgagcgacctcatgctatac |
| IntG26-CYC1-R | aaaatactgtgttgtttgtttctctctagccgttgattggcttcgagcgtcccaaaacc |
| Conf-IntG3-F | acgaatccaacggtgctgttcttg |
| Conf-IntG3-R | aactgttccaccgttcttgacac |
| Conf-IntG4-F | agcggaaacagcgtgatgagtgagttaag |
| Conf-IntG4-R | agacactcaagatacacacttacatgaacg |
| Conf-IntG11-F | cagatattcagtggagtggggctt |
| Conf-IntG11-R | agtaccaatttcctgctaatgggca |
| Conf-IntG13-F | ttcaagtgcgaagtttccgcatatg |
| Conf-IntG13-R | gtacaagacctggttgtgtgccata |
| Conf-IntG16-F | attcaagcctgctgcaattgtgaag |
| Conf-IntG16-R | ttcagatgacaatagtctcttgcagaacac |
| Conf-IntG17-F | gctaacaatgtgaatacgcacaccgtata |
| Conf-IntG17-R | ggttagaaatcgctggtaacattactgatacc |
| Conf-IntG21-F | gaccaagctatgaaatgcgtaagatgaac |
| Conf-IntG21-R | attaaatgagtagatgctgccagagtactg |
| Conf-IntG22-F | tgtagattcaatatattttcgatcaatggcttctatcag |
| Conf-IntG22-R | tcgttcatatgtagcagcgatggtag |
| Conf-IntG26-F | cagacaaactagggtgaggattcttcg |
| Conf-IntG26-R | tcagtccaatgaatagatcggttaaagc |
| CrGS-F | gtcaaggagaaaaaaccccggatccatggccggagaaacaacc |
| CrGS-R | agccgcggtaccaagcttactcgagtcattcctcaaatttcaatgtattt |
| CrGS-I301E-F | ggcacctctcattatgggaaggaaaaaggaaatcggaagttccactgg |
| CrGS-I301E-R | tttccttcccataatgagaggtgccgcagggagctcaaat |
| CrGS-P290H-A291S-I295L-F | cggagtcgctatttgagctccactccgcacctctcctaatgggaaggaaaaaga |
| CrGS-P290H-A291S-I295L-R | gagctcaaatagcgactccggtgcacctacgagcata |
| CrVinBLAST-F | tcaaggagaaaaaaccccggatccaatggccggaaaatcaccaga |
| CrVinBLAST-R | aaatcaacttctgttccatgtcgacttaaggagctttcaaggtcttt |
| CrVinBLAST-C51A-H56A-F | ggttcaaggtgctatattgtgggattgctcatactgaccttgctttcgctaagaatgag |
| CrVinBLAST-C51A-H56A-R | aatcccacaatatagcaccttgaacctcacatcatcctca |
| CrVinBLAST-M298E-V299E-F | ctctgctcctttgcttatggggaggaaggaagaagctggaagtagcattgg |
| CrVinBLAST-M298E-V299E-R | cctccccataagcaaaggagcagagtgaagatcaagtggt |
| CrVinBLAST-H288P-S289A-L293I-F | gcacccccggaaccacttgatcttccagccgctcctttgataatggggaggaagatggt |
| CrVinBLAST-H288P-S289A-L293I-R | aagatcaagtggttccgggggtgccccaagaagaacaagctt |

| Primer Name | Sequences 5'-3' |
|---|---|
| CrVinBLAST-K359G-F | gctccttaagtcgacatggaacagaagttgatttccgaagaagacctc |
| CrVinBLAST-K359G-R | ttctgttccatgtcgacttaaggagcccccaaggtctttgcaacgtc |
| For-val-pGADT7 | nnnnnttacgctcatatggccatgg |
| Rev-val-pGADT7 | nnnnnttcagtatctacgattcatc |
| For-val-pGBKT7 | nnnnntgactgtatcgccggaatttgtaata |
| Rev-val-pGBKT7 | nnnnncataagaaattcgcccggaatta |
| Conf-AD-F | ctatctattcgatgatgaagataccccac |
| Conf-AD-R | ggccaagattgaaacttagaggag |
| Conf-BD-F | gacagttgactgtatcgccg |
| Conf-BD-R | ctcaagacccgtttagaggcc |
| AD-CrVinBLAST-F | atatggccatggaggccagtgaattcatggccggaaaatcaccagaag |
| AD-CrVinBLAST-R | tatctacgattcatctgcagctcgagttaaggagctttcaaggtctttgcaac |
| BD-CrVinBLAST-F | tgcatatggccatggaggccgaattcatggccggaaaatcaccag |
| BD-CrVinBLAST-R | tagttatgcggccgctgcaggtcgacttaaggagctttcaaggtctttgcaac |
| AD-CrGS-F | tatggccatggaggccagtgaattcatggccggagaaacaaccaa |
| AD-CrGS-R | atctacgattcatctgcagctcgagtcattcctcaaatttcaatgtattt |
| BD-CrGS-F | gcatatggccatggaggccgaattcatggccggagaaacaaccaa |
| BD-CrGS-R | agttatgcggccgctgcaggtcgactcattcctcaaatttcaatgtattt |
| Not1-SGD-F | ccttgtaatccatcgatactagtgcttaagtttttgtcttttaaccaattc |
| SGD-Not1-R | tcgaattcaaccctcactaaagggcatggataacacccaagctg |
| SGD-EGFP-R | tctcggcatggacgagctgtacaagggtagcggtagcggtagcatggataacacccaagctgaa |
| EGFP-SGD-F | actccatgctaccgctaccgctaccagtttttgtcttttaaccaattcaac |
| EGFP-linker-R | ggtagcggtagcggtagcatggagttcgtgagcaagg |
| Not1-EGFP-F | ccttgtaatccatcgatactagtgctcacttgtacagctcgtcc |
| EGFP-Not1-R | tcgaattcaaccctcactaaagggcatggagttcgtgagcaagg |
| EGFP-F | cttgtacagctcgtccatg |
| SGD-R | ttaagtttttgtcttttaaccaattc |
| IN-SGD-R | cagcttgaacatctgacaatgg |
| pESC-URA-CrGS-R | tcgaattcaaccctcactaaagggcatggccggagaaacaac |
| IN-CrGS-F | ctatggatggtgttgtggacac |
| EGFP-CrGS-F | actccatgctaccgctaccgctaccttcctcaaatttcaatgtatttccaa |
| CrGS-EGFP-R | tctcggcatggacgagctgtacaagggtagcggtagcggtagcatggccggagaaacaacc |
| Not1-CrGS-F | ccttgtaatccatcgatactagtgcttattcctcaaatttcaatgtatttcca |
| Nab2-Not1-R | tcgaattcaaccctcactaaagggcatgtctcaagaacagtacacag |
| IN-Nab2-F | cttggtagatcatgcccacatgc |
| IN-Nab2-R | gcatgtgggcatgatctaccaagaggacagtgaggaaaca |
| mCherry-NAB2-F | ccttgctcacaaattccatgctaccgctaccgctaccgttcatttccgtatcttgttct |
| mCherry-R | ggtagcatggaatttgtg |
| IN-mCherry-F | ttcaccttgtagatgaactcg |
| PESC-URA-mCherry-F | ccttgtaatccatcgatactagtgcttacttgtacagctcgtccat |
| CrCAD1-F | tcgaattcaaccctcactaaagggcatggccggaaaatcaccagag |
| CrCAD1-R | ccttgtaatccatcgatactagtgcttaaggagctttcaaggtgttggca |
| CrCAD3-F | tcgaattcaaccctcactaaagggcatggccagaaaatcaccaga |
| CrCAD3-R | ccttgtaatccatcgatactagtgctcacacctctgatggaagag |
| CrCAD5-F | tcgaattcaaccctcactaaagggcatggctggaaaatcaccaga |
| CrCAD5-R | ccttgtaatccatcgatactagtgcttaggactctggtggaggagtt |
| RsVinBLAST-F | tcgaattcaaccctcactaaagggcatggccggaaaatcacctgaa |
| RsVinBLAST-R | ccttgtaatccatcgatactagtgcttaaggagctttcaaggtgttggca |
| CoVinBLAST-F | tcgaattcaaccctcactaaagggcatggctggaaaatccccagaagag |
| CoVinBLAST-R | ccttgtaatccatcgatactagtgcttaaggagctttcaaggtgtt |
| MsVinBLAST-F | tcgaattcaaccctcactaaagggcatggctggaaaatcaccagaa |

| Primer Name | Sequences 5'-3' |
|---|---|
| MsVinBLAST-R | ccttgtaatccatcgatactagtgcttaaggagctttcaaggtgttggca |
| CrGS-Not1-R | tcgaattcaaccctcactaaagggcatggccggagaaacaaccaa |
| linker-CrGS-F | cgccaccgccgctgcctccgccaccttcctcaaatttcaatgtatttccaatg |
| SGD-linker-R | ggtggcggaggcagcggcggtggcggaagcggaggaggaggtagcatggataacacccaagctga |
| linker-RIDD-R | cggtggcggaagcggaggaggaggtagctgcggaagtttaagggaatg |
| Not1-RIDD-F | atccttgtaatccatcgatactagtgctcacttagcttcttccttttc |
| Not1-SpyTag-linker-F | atccttgtaatccatcgatactagtgctcacttagttggcttgtaagcgtcaaccatgacgatgtgggcgctac ctcctcctccgc |
| RIAD-Not1-R | tcgaattcaaccctcactaaagggcatgtgtggtctagaacaatatgcaaac |
| linker-RIAD-F | cgccaccgccgctgcctccgccaccacagccctcggtggcc |
| SpyCatcher-Not1-R | tcgaattcaaccctcactaaagggcatggctatggttgacactttgtcc |
| linker-SpyCatcher-F | cgccaccgccgctgcctccgccaccaatgtgagcatcacccttg |
| linker-SpyCatcher-R | tggcggaagcggaggaggaggtagcgctatggttgacactttgtcc |
| Not1-SpyCatcher-F | ccttgtaatccatcgatactagtgctcaaatgtgagcatcaccct |
| mVenusN-F | tcgaattcaaccctcactaaagggcatggaattcgtgagcaagggcgagg |
| mVenusN-R | tcatccttgtaatccatcgatactagtgcttactcgatgttgtggcggatcttga |
| Linker-mVenusN-R | agtaccaccagaaccctcgatgttgtggcggatcttgaagttggc |
| mVenusN-SGD-F | ggttctggtggtactatgggttccaaagatgatcaatctttggtt |
| CrGS-mVenusN-F | gaattccatagtaccaccagaaccttcctcaaatttcaatgtatttccaa |
| CrGS-mVenusN-R | ggaaggttctggtggtactatggaattcgtgagcaagggcgaggagctgttca |
| CrVinBLAST-mVenusN-F | ctcacgaattccatagtaccaccagaaccaggagctttcaaggtctttgcaacg |
| CrVinBLAST-mVenusN-R | aaagctcctggttctggtggtactatggaattcgtgagcaagggcgaggagctg |
| IN-mVenusN-R | gaccaggatgggcaccaccc |
| CrVinBLAST-mVenusC-F | gttcttctgcttgtccatagtaccaccagaaccaggagctttcaaggtctttgca |
| CrVinBLAST-mVenusC-R | ccttgaaagctcctggttctggtggtactatggacaagcagaagaacggcatc |
| CrGS-mVenusC-F | gcttgtccatagtaccaccagaaccttcctcaaatttcaatgtatttcca |
| CrGS-mVenusC-R | ttgaggaaggttctggtggtactatggacaagcagaagaacggcatcaag |
| mVenusC-SGD-F | tgatcatctttggaacccatagtaccaccagaacccttgtacagctcgtccatgccgagagtgat |
| mVenusC-SGD-R | agctgtacaagggttctggtggtactatgggttccaaagatgatcaatctttggttgttgc |
| mVenusC-F | atggacaagcagaagaacggcatcaag |
| mVenusC-R | ttacttgtacagctcgtccatgccga |

**Table S8.** List of genomic loci for yeast genome integrations.

| Integration sites | Spacer sequences (5'-3') | Chromosomal loci |
|---|---|---|
| IntG3 | GCCGTCCTAGCTGAAGTGTG | ChrVI: 48,452-48,471 |
| IntG4 | CCTGGCGCTATGATGATGAG | ChrVII: 478,898-478,917 |
| IntG11 | ATATGTCTCTAATTTTGGAA | ChrXI: 93,963-93,982 |
| IntG13 | GAGCATTTACTGACACCTGG | ChrVII: 1,011,308-1,011,327 |
| IntG14 | AATGGATAAAAAATACAACG | ChrVIII: 90,487-90,506 |
| IntG16 | TATATAATGAATACACATGG | ChrVIII: 121,948-121,967 |
| IntG17 | GAAATTATATAAAACACATG | ChrVIII: 147,100-147,119 |
| IntG21 | TCAAGGGGTTGCATATAGGG | ChrXV: 73,653-73,672 |
| IntG22 | TCACACGAATGAGAATTGGG | ChrXV: 371,041-371,060 |
| IntG26 | GAGAAAATAAAAAAAATATG | ChrXV: 724,860-724,879 |

**Table S9.** Percentage protein identity of enzymes in the syntenic regions.

| | g3232.t1_ g3233.t1 | SI8HD1_ SI43.28 | Sl8HD2_ SI43.27 | CrCAD2 (VinBLAST) | CrGS | CrGS2 | CrTHAS2 |
|---|---|---|---|---|---|---|---|
| g3232.t1_g3233.t1 | / | 65.664 | 71.161 | 68.434 | 57.434 | 55.66 | 57.27 |
| SI8HD1_SI43.28 | 65.664 | / | 72.191 | 58.449 | 47.419 | 46.281 | 51.075 |
| Sl8HD2_SI43.27 | 71.161 | 72.191 | / | 64.706 | 54.045 | 51.532 | 56.486 |
| CrCAD2 (VinBLAST) | 68.434 | 58.449 | 64.706 | / | 53.226 | 52.198 | 53.226 |
| CrGS | 57.434 | 47.419 | 54.045 | 53.226 | / | 91.64 | 50.311 |
| CrGS2 | 55.66 | 46.281 | 51.532 | 52.198 | 91.64 | / | 49.598 |
| CrTHAS2 | 57.27 | 51.075 | 56.486 | 53.226 | 50.311 | 49.598 | / |

**Table S10.** Details on genome information used in this study.

| Species | Version | Source | Citation |
|---|---|---|---|
| *Nepeta mussinii* | 2023 | Dryad Digital Repository (https://doi.org/10.5061/dryad.88tj450) | (*25*) |
| *Solanum lycopersicum* | SL28frl | CoGe (https://genomevolution.org/coge/) | (*50*) |
| *Mitragyna speciosa* | 2021 | Medicinal Plant Genomics project website (http://mpgr.uga.edu) | (*51*) |
| *Gelsemium sempervirens* | 2018 | Dryad Digital Repository (https://doi.org/10.5061/dryad.08vv50n) | (*52*) |
| *Calotropis gigantea* | 2018 | Dryad Digital Repository (https://doi.org/10.5061/dryad.fk41r) | (*53*) |
| *Catharanthus roseus* | 2023 | Dryad Digital Repository (https://doi.org/10.5061/dryad.d2547d851) | (*7*) |
| *Rauvolfia tetraphylla* | 2024 | Figshare https://doi.org/10.6084/m9.figshare.21679628 | (*54*) |
| *Eustoma grandiflorum* | 2023 | Plantgarden (https://plantgarden.jp/en/list/t52518) | (*55*) |

**Data S1. All updated figures with statistical information.** Quantitative comparisons and statistical analysis (p-values) for all figures in the manuscript.

**Data S2. The list of genes co-expressed with one or more baits from weighted gene co-expression network analysis (WGCNA) in Fig. 2A.**

**Data S3. The input files for MD simulations.** The files include force field parameters for all constructed substrate molecules and NADPH, as well as the final equilibrated conformations from MD simulations of all complexes.
